# Single-cell resolution of resection-dependent chromatin accessibility in response to DNA double strand breaks reveals specific gene expression programs

**DOI:** 10.64898/2026.04.21.719635

**Authors:** Sarah Collins, Anne-Laure Finoux, Coline Arnould, Romane Carette, Vincent Rocher, Gaëlle Legube, Thomas Clouaire

**Author notes:** these authors contributed equally (Anne-Laure Finoux, Coline Arnould).

## Abstract

Chromatin structure is extensively modified and remodeled to allow for efficient signaling and repair of DNA double-strand breaks (DSBs), a process which is influenced by the nature and properties of the damaged locus or the repair pathway involved. However, the extent of cell-to-cell variability in these DSB-induced chromatin modifications remains poorly characterized. Here, we report how chromatin accessibility is reshuffled at the damaged locus at high resolution. Notably, we identify a long-range increase in chromatin accessibility which is dependent on functional resection. We report that resection-dependent chromatin remodeling events at individual damaged loci display significant heterogeneity in single cells which are only partly explained by the cell cycle, and can frequently be detected as asymmetric and unidirectional. Moreover, we identify strong transcriptional phenotypes upon sustained DNA damage as well as subpopulations of S and G2/M cells for which similar resection-dependent chromatin accessibility patterns occur simultaneously at multiple DSBs. These coordinated chromatin accessibility patterns at multiple DSBs are associated with specific gene expression programs, including DNA damage checkpoint signaling, inflammation and apoptosis. Altogether, our results highlight the heterogeneity in chromatin remodeling during DSB repair and suggest unexpected links between DNA end resection and specific gene expression programs mounted in response to genotoxic stress.

## Introduction

DNA double-strand breaks (DSBs) represent highly toxic DNA lesions which can locally induce mutations (such as insertions or deletions) or trigger severe chromosomal rearrangements including translocations. Deficiencies in genes involved in DSB repair are associated with many human diseases including neurological defects, immune deficiencies, premature aging and cancer predisposition ^1^. Eukaryotic cells are equipped with two main pathways ensuring the repair of DSBs, Non-Homologous End Joining (NHEJ) and Homologous Recombination (HR)^2^. NHEJ can operate throughout the cell cycle and relies on the direct ligation of DSB ends by the XRCC4-DNA ligase IV complex^3^. HR repair is initiated by the nucleolytic processing of DSB ends, also known as resection, generating a stretch of single-stranded DNA^4^. This structure is first recognized by Replication Protein A (RPA) which is subsequently exchanged for RAD51. The RAD51 nucleofilament is responsible for homology search, allowing for repair synthesis using the undamaged copy of the chromosome as a template.

DNA end resection is crucial to determine the fate of a given DSB since it is mandatory for HR repair and concomitantly inhibits the execution of canonical NHEJ^4^. Resection is initiated by the MRE11-RAD50-NBS1 (MRN) complex (Mre11-Rad50-Xrs2 in yeast) in a two-step process involving a first incision of the 5’-terminated strand by the endonuclease activity of MRE11, followed by 3’ to 5’ exonucleolytic digestion of the same DNA strand via MRE11 exonuclease activity^5–7^. This “short-range” bidirectional resection can be extended by two nucleases, EXO1 and DNA2, allowing further degradation of the 5’ terminated strand, or “long-range” resection^4^. Both short-range and long-range resection are stimulated by CtIP/Sae2 which is itself activated upon phosphorylation by cyclin dependent kinases (CDKs)^6,8–11^. As a consequence, end resection, and thus HR, is more prevalent during the S and G2 phase of the cell cycle, coinciding with the availability of the sister chromatid as a template for repair. Yet, resection also occurs in G0 or G1 in some contexts^12–16^. Aside from this cell cycle regulation, the different mechanisms precisely controlling the extent and directionality of resection at a given DSB have not yet been fully characterized.

Chromatin is the natural substrate for DSB repair and is extensively modified at different scales following DSBs, contributing to damage sensing, signaling and repair (reviewed in^17^), including resection. For example, DSB-induced modifications of histones such as H2A ubiquitylation, H3K36me3 or H4K20 methylation facilitate the recruitment of positive or negative regulators of resection such as BRCA1, CtIP or 53BP1 (reviewed in^17,18^). Nucleosomes themselves also impede MRE11-dependent incisions *in vitro* and *in vivo* in yeast^19–22^, even though MRN is able to diffuse past nucleosomes *in vitro*^23^. Additionally, the activity of EXO1 or DNA2 is inhibited to different extents by nucleosomes *in vitro*^24–26^. Chromatin remodeling complexes can alleviate these local obstacles to resection through nucleosome eviction, sliding or editing via the incorporation of variants^27^. For example, RSC and SWI/SNF are required for efficient resection *in vivo* in yeast^28^ and Fun30, or its mammalian homolog SMARCAD1, facilitates resection in different contexts^22,29–34^. Yet, the precise molecular mechanisms by which these chromatin remodelers positively impact resection are not fully elucidated (discussed in^35^). Moreover, the interplay between chromatin remodeling and DSB repair appears more complex since, although controversial^28^, nucleosomes were also proposed to assemble on ssDNA generated by resection^36,37^ and resection-independent nucleosome eviction has been observed immediately at the break site in human and yeast systems^38,39^. Therefore, the type, and extent, of chromatin remodeling events necessary for efficient repair is expected to depend on the pathway involved but also on the local chromatin landscape established prior to damage as a consequence of the nature and properties of the locus^17^.

Here, we used ATAC-seq, a highly sensitive method to analyze chromatin accessibility *in vivo*, and identify that chromatin structure is extensively reshuffled at the damaged locus. First, chromatin accessibility decreases sharply at the break itself, likely reflecting the engagement of DNA end binding factors. Second, we identify a long-range increase in accessibility, extending up to 20 kb on each side of the DSB in conditions of sustained DNA damage by long-term DSB induction, which is dependent on functional resection. By single-cell ATAC-seq, we find that resection-dependent chromatin remodeling events display significant cell-to-cell heterogeneity at individual damaged loci, which is only partly explained by the cell cycle. In particular, we found that resection-dependent chromatin accessibility can frequently be asymmetric and unidirectional in single cells. Moreover, we identify that sustained DNA damage triggers extensive transcriptional phenotypes and the appearance of subpopulations of S and G2/M cells characterized by a simultaneous increase or decrease in resection-dependent chromatin accessibility at multiple DSBs. Integrating single-cell chromatin accessibility with transcriptomic data revealed that these coordinated chromatin accessibility patterns at multiple DSBs are associated with the activation of specific gene expression programs, including DNA damage checkpoint signaling, inflammation and immune response and apoptosis. Altogether, our results highlight the heterogeneity in chromatin remodeling during DSB repair and suggest unexpected links between DNA end resection and the cellular responses involving specific gene expression programs mounted upon genotoxic stress.

## Results

### Chromatin structure is drastically reconfigured at the damaged locus in response to DSB induction and the engagement of DSB repair factors

In order to probe for local modifications in chromatin structure in response to DSBs, we measured chromatin accessibility using ATAC-seq (Assay for Transposase Accessible Chromatin)^40,41^ in DIvA cells (for DSB inducible via AsiSI). In this cellular model, multiple localized DSBs can be induced on the human genome in a controlled manner following treatment with 4-hydroxytamoxifen (OHT)^42,43^. We performed ATAC-seq in absence of damage and following DSB induction by the AsiSI restriction enzyme for 4 and 24 hours (2 replicates per condition). As expected, we observed a good correspondence between ATAC-seq signal intensity and the accumulation of H3K4me3 and RNA pol II at gene promoters, as previously reported^40^, which was true across individual replicates (Supplementary Fig. 11). In order to precisely map how local chromatin accessibility is modified upon DSB induction, we focused on a set of 80 DSB sites which were previously shown to be robustly induced by AsiSI using Breaks Labeling and Enrichment on Streptavidin and Sequencing (BLESS)^43^. These sites were readily accessible prior to break induction (Fig. 1A-B and Supplementary Fig. 11-C), which was anticipated since 1- the AsiSI enzyme requires access to induce cleavage and 2- a significant fraction of these sites reside near active promoters^43,44^. Differential ATAC-seq profiles revealed that DSB induction led to a striking modification of the chromatin accessibility landscape around the break site (Fig. 1A-F). First, we detected a sharp local decrease in accessibility very close to the DSB site itself (Fig. 1A-D, see example Supplementary Fig. 11 top panel). This local decrease was accompanied by a prominent and symmetric increase in chromatin accessibility beginning from around 250 bp on each side of the break (Fig. 1C-D and Supplementary Fig. 11 bottom panel), spanning to around 2 kb on each side of the DSB after 4 hours of DSB induction (Fig. 1C), in good agreement with a recent report^45^ and data obtained 30min after Cas9-mediated DSB induction^46^. The increase in chromatin accessibility reached up to 20 kb in cells treated for 24 hours (Figure 1E-F). These patterns were consistent across individual replicates (Supplementary Fig. 11-F). While relatively wide, these continuous tracts of increased chromatin accessibility upon DSB induction did not extend over entire megabase-sized regions decorated by γH2AX or 53BP1 (Supplementary Fig. 11-H) corresponding to DNA Damage Response (DDR) foci^43,47^. Thus, we conclude that chromatin structure is substantially modified in response to DSBs, an event which remains confined to the relative proximity of the break itself and is likely related to the engagement of specific DSB repair factors.

**Figure 1:**
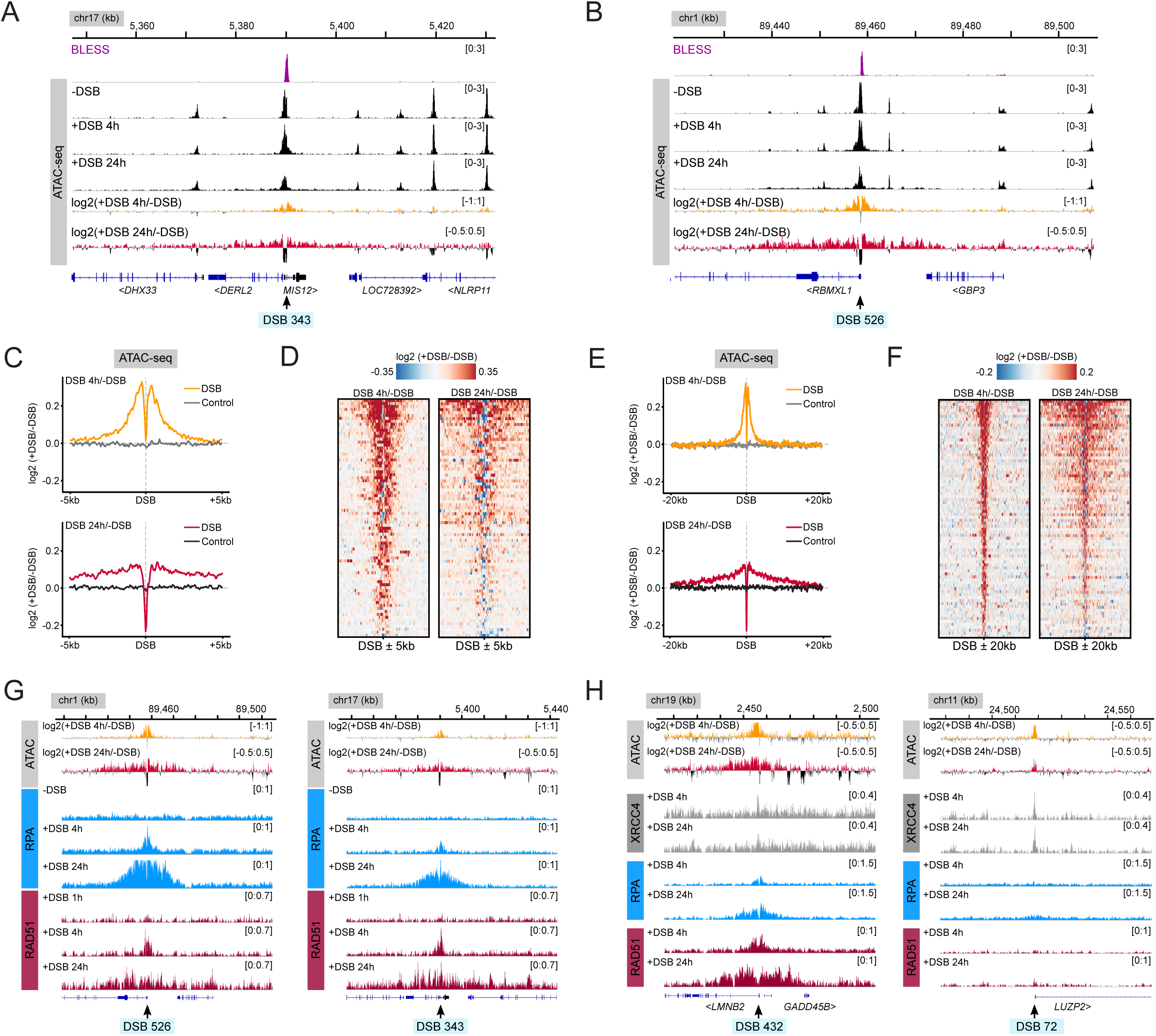
Chromatin accessibility is modified at the damaged locus following DSB induction and coincides with the binding profiles of single-strand DNA-binding proteins. (A) Genome browser screenshot representing signals for BLESS (purple), normalized ATAC-seq signals (Replicate 1, in black) as well as differential ATAC-seq profiles as log2(+DSB/-DSB) after 4 h or 24 h of DSB induction for a DSB located on chromosome 17 (DSB 343. Note that DSBs are numbered according to the predicted AsiSI sites in the genome, the majority of which are not cleaved efficiently, leading to discontinuous numbering between efficiently induced DSBs). (B) Same as (A), but for a DSB located on chromosome 1 (DSB 526). (C) Average differential ATAC-seq profile (Replicate 1) as log2(+DSB/-DSB) for 80 DSBs^43^ or 80 control regions on a 10 kb window (± 5 kb from the DSB) after 4 h of DSB induction (yellow, top panel) or 24 h of DSB induction (red, bottom panel). (D) Heatmap representing differential ATAC-seq profile (Replicate 1) as log2(+DSB/-DSB) for 80 DSBs on a 10 kb window (± 5 kb from the DSB) after 4h of DSB induction (left panel) or 24 h of DSB induction (right panel). (E) Same as (C) on a 40 kb window (± 20 kb from the DSB). (F) Same as (D) on a 40 kb window (± 20 kb from the DSB). (G) Genome browser screenshot representing differential ATAC-seq profiles as log2(+DSB/-DSB) after 4 h or 24 h of DSB induction as in Figure 1A, together with RPA ChIP seq in DIvA cells (blue), either untreated, treated for 4 h or 24 h and RAD51 ChIP-seq (dark red) for cells treated for 1 h, 4 h, or 24 h. Signals are displayed for a DSB located on chromosome 1 (DSB 526, left panel) and a DSB located on chromosome 17 (DSB 343, right panel). (H) Genome browser screenshot representing differential ATAC-seq profiles as log2(+DSB/-DSB) after 4 h or 24 h of DSB induction as in Figure 1A, XRCC4 ChIP-seq after 4 h and 24 h of DSB induction (from^43^, in grey), RPA ChIP-seq after 4 h and 24 h of DSB induction (from^44^ in blue) and RAD51 ChIP-seq after 4 h and 24 h of DSB induction (from^49^ in dark red). Signals are displayed for a DSB located on chromosome 19 (DSB 432, left panel), a HR-prone DSB, and a DSB located on chromosome 11 (DSB 72, right panel), a NHEJ-prone DSB.

Next, we compared these differential ATAC-seq signals with high resolution ChIP-seq profiles for specific DSB repair factors. The local decrease in chromatin accessibility, spanning about 250 bp on each side of the break, corresponded well with the point of maximum ChIP-seq signal intensity for MRE11 and was immediately adjacent to the localization of phosphorylated ATM (ATM S1981P, from^47^) (Supplementary Fig. 22, left panels), suggesting a link with the recruitment of DSB sensors to DNA ends. Of note, this local decrease also overlapped well with the maximal binding of XRCC4 and DNA ligase IV (Supplementary Fig. 22, right panels) and may therefore also be indicative of ongoing NHEJ repair. On the other hand, the symmetric and distal increase in chromatin accessibility following DSB showed good concordance with ChIP-seq profiles for the single-strand DNA binding proteins RPA and RAD51 (Fig. 1G). Indeed, both factors are recruited on both sides of DSBs after 4 hours of DSB induction^43,44,48^ and can spread further away after 24 hours due to the accumulation of DSBs with longer resection tracts within the cell population upon sustained DNA damage^44,49^. This trend could be detected by inspection of individual DSBs (Fig. 1G-H) and was confirmed on average profiles for the 80-best induced DSBs in DIvA cells (Supplementary Fig. 22- C).

Previous reports identified that a subset of AsiSI-induced DSBs localized in actively transcribed genes are particularly prone to undergo resection and accumulate RPA and RAD51 (^43,48^, see Supplementary Fig. 22). By opposition, NHEJ-prone DSBs can efficiently recruit the NHEJ factors XRCC4 or DNA ligase IV but exhibit low levels of RPA or RAD51 (Supplementary Fig. 22)^43,48^. Comparison of average differential ATAC-seq profiles (Supplementary Fig. 22-F), and individual examples (Fig. 1H, compare left and right panels), revealed that the distal increase in chromatin accessibility triggered upon damage is more prominent at HR-prone- compared to NHEJ-prone DSBs (Supplementary Fig. 22), suggesting that it is directly linked with resection and HR. To confirm that increased Tn5 transposition can indeed occur on resected DNA strands, we profiled single-strand DNA binding proteins by CUT&Tag, an antibody-targeted Tn5 transposition method^50^. As expected, we observed CUT&Tag enrichment for RAD51 (using two different antibodies), RPA phosphorylated on serine 33 (RPA S33P) (Supplementary Fig. 22) and RPA (Supplementary Fig. 22) spanning over more than 20 kb on each side of many AsiSI-induced DSBs after 24 hours of DSB induction, in agreement with the reported ability of Tn5 to transpose in single-strand DNA^51^. As a comparison, CUT&Tag signals for phosphorylated ATM (ATM S1981P) showed a strong enrichment near DSB sites, as previously observed by ChIP-seq (^47^ and Supplementary Fig. 22). Altogether, our data suggests that the reconfiguration of chromatin structure around DSBs detected by ATAC-seq represents a composite picture involving the recruitment of DSB sensing factors at the break itself as well as detectable remodeling of chromatin upon DSB end resection and HR repair.

### Distal and extended increase in DSB-induced chromatin accessibility requires DNA end resection

Next, we investigated whether the DSB-induced chromatin remodeling events identified above were indeed dependent on the execution of DSB repair and, particularly, DNA end resection. For this, we performed ATAC-Seq experiments following siRNA-mediated depletion of CtIP (Supplementary Fig. 33), which is required for efficient resection in many contexts including AsiSI-induced DSBs^52^. Differential ATAC-seq profiles revealed that CtIP depletion led to a significant reduction of chromatin accessibility on both sides of the break following 4 hours or 24 hours of DSB induction (Fig. 2A-B, Supplementary Fig. 33). Similarly, ATAC-seq experiments following inhibition of MRE11 endonuclease activity using PFM01^7^ revealed that impairing resection initiation lead to a complete abolition of DSB-induced chromatin accessibility distal to the break after either 4 or 24 hours of DSB induction (Fig. 2C-E, Supplementary Fig. 33). Moreover, inhibition of ATM, which is reported to favor resection of DNA breaks^53,54^, with KU-55933 also strongly affected the increase in accessibility following DSB induction (Fig. 2C-E, Supplementary Fig. 33). Conversely, inhibition of DNA-PKcs with NU7441, which was previously shown to increase resection in a similar context^16,55,56^, led to a sharp increase in chromatin accessibility at a distance from the break at the 24 hours time point (Fig. 2C,E and Supplementary Fig. 33). In agreement, we also observed a reduction in ATAC-seq signal following DSB induction upon co-depletion of EXO1 and DNA2 (Fig. 2F-G and Supplementary Fig. 33-E), even though the effect was not as pronounced as CtIP depletion or MRE11 inhibition.

**Figure 2:**
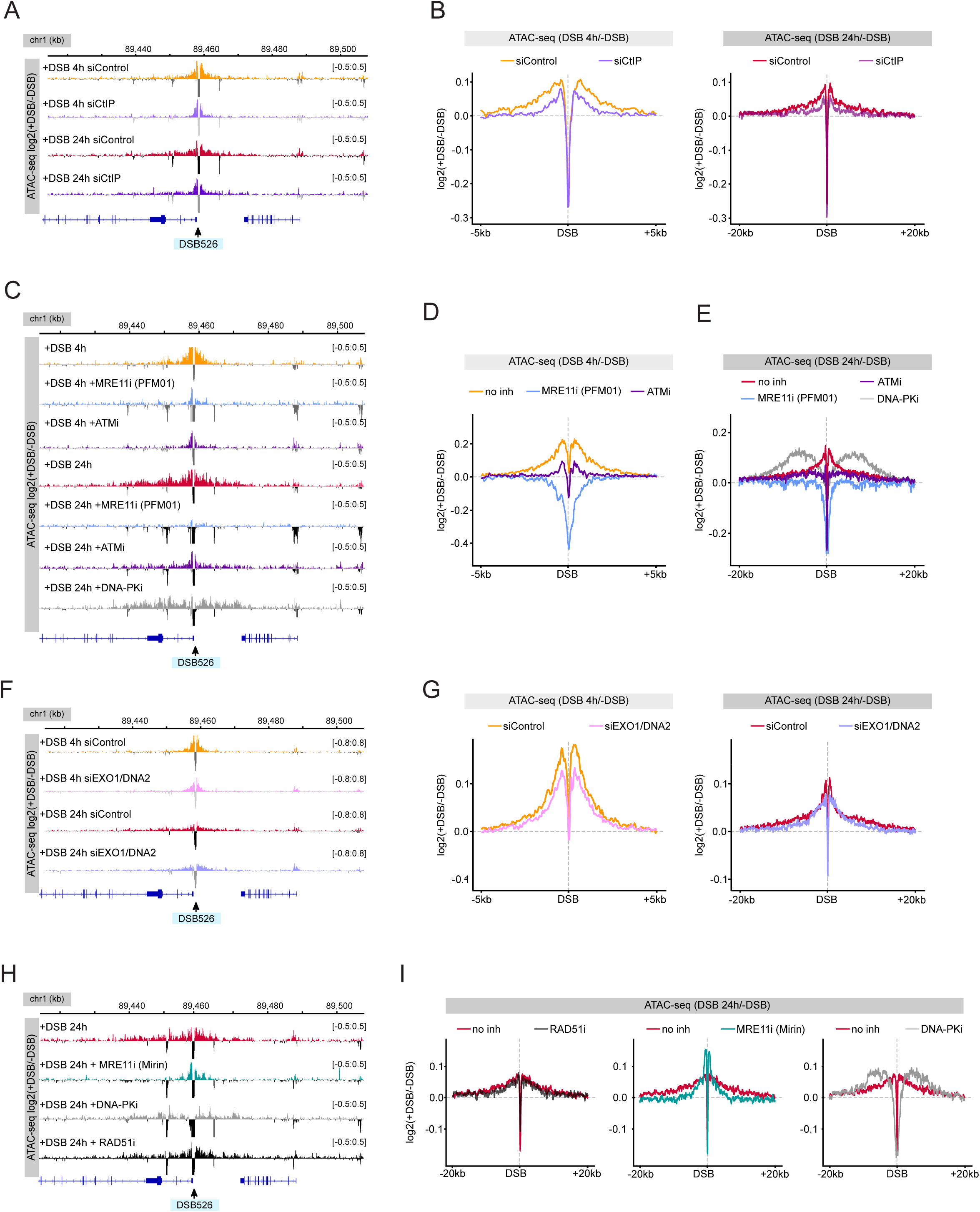
DSB-induced increase in chromatin accessibility depends on DNA end resection. (A) Genome browser screenshot representing differential ATAC-seq profiles as log2(+DSB/-DSB) after 4 h or 24 h of DSB induction in siControl- or siCtIP-treated cells for a DSB located on chromosome 1 (DSB 526). (B) Average profiles of differential ATAC-seq represented as log2(+DSB/-DSB) in siControl-treated cells after 4 h (yellow line, left panel) and 24 h (red line, right panel) of DSB induction, and in siCtIP-treated cells after 4 h (light purple, left panel) and 24 h (dark purple, right panel) at 80 DSBs on a 10 kb window (± 5 kb from the DSB, left panel) and on a 40 kb window (± 20 kb from the DSB, right panel). (C) Genome browser screenshot representing differential ATAC-seq profiles as log2(+DSB/-DSB) after 4 h or 24 h of DSB induction, as well in cells with additional treatment of inhibitors of MRE11 endonuclease activity (PFM01), ATM (KU-55933), and DNA-PKcs (NU7441) for a DSB located on chromosome 1 (DSB 526). (D) Average profiles of differential ATAC-seq represented as log2(+DSB/-DSB) in cells after 4 h (yellow line) of DSB induction, and in cells with additional treatment of inhibitors of MRE11 endonuclease activity (PFM01, blue line), and ATM (KU-55933, purple line,) at 80 DSBs on a 10 kb window (± 5 kb from the DSB). (E) Average profiles of differential ATAC-seq represented as log2(+DSB/-DSB) in cells after 24 h (red line) of DSB induction, and in cells with additional treatment of inhibitors of MRE11 endonuclease activity (PFM01, blue line), ATM (KU-55933, purple line), and DNA-PKcs (NU7441, grey line), at 80 DSBs on a 40 kb window (± 20 kb from the DSB). (F) Genome browser screenshot representing differential ATAC-seq profiles as log2(+DSB/-DSB) after 4 h or 24 h of DSB induction in siControl- or siEXO1+DNA2-treated cells for a DSB located on chromosome 1 (DSB 526). (G) Average profiles of differential ATAC-seq represented as log2(+DSB/-DSB) in siControl-treated cells after 4 h (yellow line, left panel) and 24 h (red line, right panels) of DSB induction, and in siEXO1+DNA2-treated cells after 4 h (light pink, left panel) and 24 h (light purple, right panels) at 80 DSBs on a 10 kb window (± 5 kb from the DSB, left panel) and on a 40 kb window (± 20 kb from the DSB, right panel). (H) Genome browser screenshot representing differential ATAC-seq profiles as log2(+DSB/-DSB) after 24 h of DSB induction, as well in cells with additional treatment of inhibitors of MRE11 exonuclease activity (Mirin), DNA-PKcs (NU7441), and RAD51 (B02) for a DSB located on chromosome 1 (DSB 526). (I) Average profiles of differential ATAC-seq represented as log2(+DSB/-DSB) in cells after 24 h (red lines) of DSB induction, and in cells with additional treatment of inhibitors of RAD51 (B02, black line, left panel), MRE11 exonuclease activity (Mirin, teal line, middle panel), and DNA-PKcs (NU7441, grey line, right panel), at 80 DSBs on a 40 kb window (± 20 kb from the DSB).

Finally, we investigated whether distal DSB-induced chromatin remodeling is occurring upstream or downstream of RAD51 nucleofilament assembly using B-02, an inhibitor of RAD51 association with DNA^57^. RAD51 immunofluorescence confirmed that B-02 treatment strongly reduced RAD51 nucleofilament assembly in cells, similar to the inhibition of MRE11 exonuclease activity by Mirin^58^ (Supplementary Fig. 33-G). We performed ATAC-seq experiments upon 24 hours of DSB induction in cells treated or not with B-02, as well as Mirin and NU7441 as controls. Differential accessibility profiles at DSBs in B-02-treated cells showed a very moderate effect at an individual site (Fig. 2H) or when averaged for 80 DSBs (Fig. 2I, left panel). While B-02 treatment had a small, but significant, impact within the 10-20 kb range (Supplementary Fig. 33-I), the defects were milder compared with MRE11 inhibition with Mirin or the increased accessibility triggered by inhibition of DNA-PKcs (Fig. 2H-I, Supplementary Fig. 33-I). Thus, while DSB-induced chromatin accessibility is largely dependent on resection, it appears to mostly occur prior to RAD51 nucleofilament assembly. Altogether, we conclude that distal long-range DSB-induced chromatin accessibility is occurring as a consequence of DNA end resection, either concomitantly with end processing and/or as a requirement to ensure access to resection nucleases and subsequent loading of single-strand DNA binding proteins.

### Deconvolution of DSB-induced chromatin remodeling at single-cell resolution identifies significant cell-to-cell heterogeneity at individual loci and frequent asymmetric events

Next, we set out to characterize DSB-induced and resection-dependent chromatin accessibility at the level of individual cells. For this, we performed single-nuclei ATAC-seq (10X Genomics, referred to as single-cell ATAC-seq for convenience) in untreated DIvA cells as well as following 4 hours and 24 hours of DSB induction to retrieve a total of more than 4000 cells (see Supplementary Table1). Pseudo-bulk differential ATAC-seq profiles for 80 DSBs, combining accessibility profiles from each individual cell in a given condition, were similar to those obtained from bulk data. Indeed, we observed an increased signal around 2 kb on each side of DSBs for the 4 hours treatment and extending further away after 24 hours of DSB induction (Supplementary Fig. 44, Fig. 3A, to be compared with Fig. 1C, E). Of note, the local decrease in accessibility immediately at DSBs on these pseudo-bulk average profiles was less striking compared to bulk experiments while the local increase (≈ 2 kb) was slightly more pronounced, especially after 24 hours, for reasons which remain unclear.

**Figure 3:**
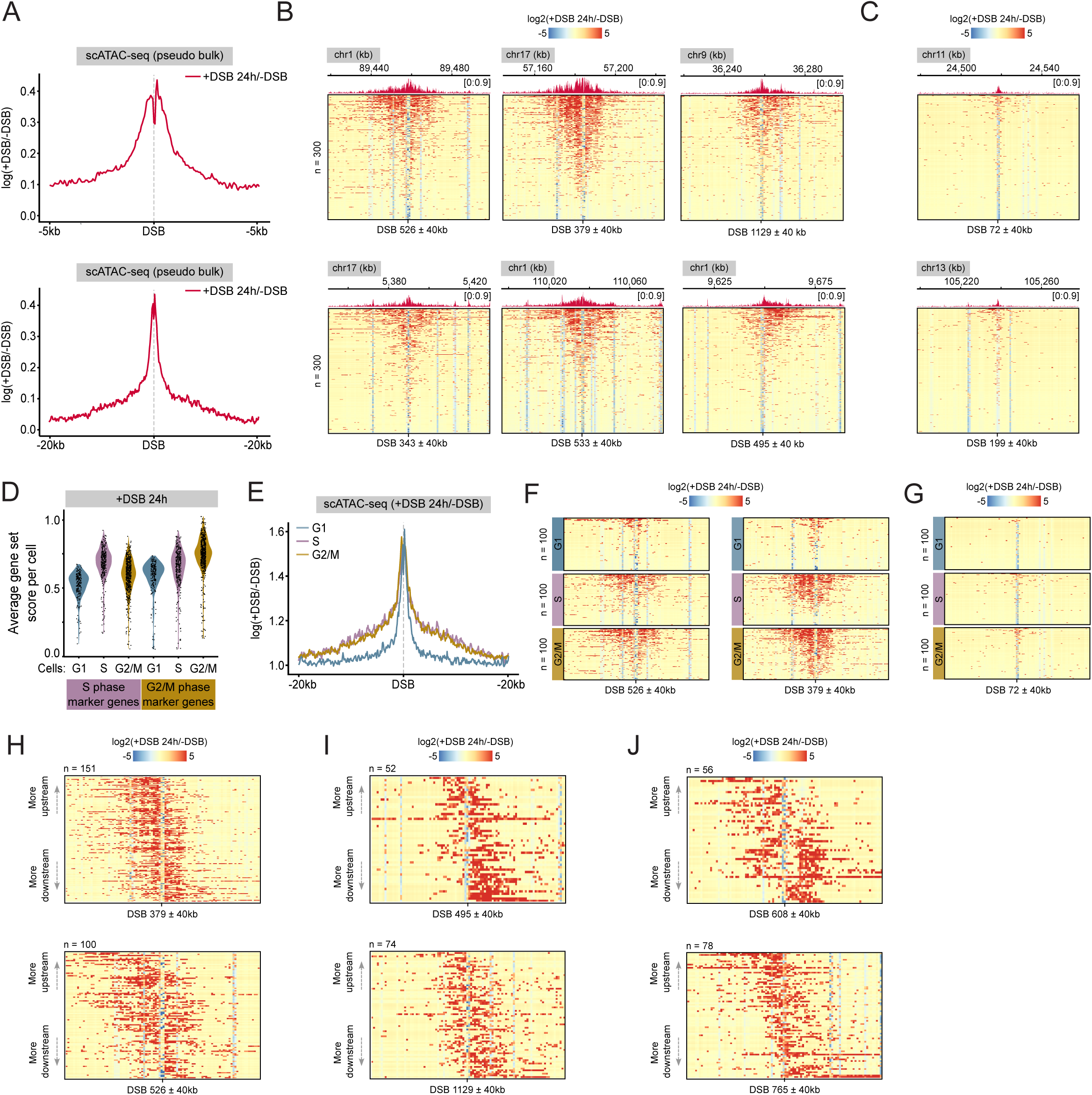
Resolving DSB-induced chromatin remodeling at single-cell resolution. (A) Pseudo-bulk average differential ATAC-seq profile as log2(+DSB/-DSB) for 80 DSBs after 24 h of DSB induction on a 10 kb window (± 5 kb from the DSB, top panel) and 40 kb window (± 20 kb from the DSB, bottom panel). (B) Single-cell differential chromatin accessibility profiles as log2(+DSB/-DSB) (n=300 cells) on an 80 kb window (± 40 kb from the DSB) after 24 h of DSB induction for DSBs 526, 379, 1129, 343, 533, and 495. (C) Same as (B), for DSBs 72 and 199. (D) Violin plots showing average gene set scores per cell for S phase and G2/M phase marker gene sets as defined in Seurat^60^ in cells after 24 h of DSB induction. (E) Pseudo-bulk average differential ATAC-seq profile as log2(+DSB/-DSB) for cells in G1 (blue line), S (pink line) or G2/M (gold line) for 80 DSBs after 24 h of DSB induction on a 40 kb window (± 20 kb from the DSB). (F) Single-cell differential chromatin accessibility profiles as log2(+DSB/-DSB) for cells in G1, S or G2/M on a 40 kb window (± 20 kb from the DSB) after 24 h of DSB induction for DSBs 526 and 379. (G) Same as (F), for DSB 72 (H) Single-cell differential chromatin accessibility profiles as log2(+DSB/-DSB) for cells after 24 h of DSB induction on an 80kb window for DSBs 379 (n=151 cells) and 526 (n=100 cells). Cells ordered by calculated upstream-to-downstream ratio (see methods). (I) Same as (H) for DSBs 495 (n=52 cells) and 1129 (n=74 cells). (J) Same as (J) for DSBs 608 (n=56 cells) and 765 (n=78 cells).

Next, we generated differential accessibility heatmaps for each DSB in individual cells after either 4 hours or 24 hours of DSB induction. For this, the local accessibility in a given genomic bin in each treated cell was divided by the average accessibility for the same bin computed across all untreated cells. We focused our analysis on sets of 300 cells with the highest overall coverage in each condition from our scATAC-seq experiment. This analysis highlighted that DSB-induced chromatin accessibility profiles exhibit substantial cell-to-cell variation (heatmaps for each individual DSB are available as Additional Figure 1). First, single-cell resolved heatmaps showed that the majority of cells exhibit a decrease in chromatin accessibility proximal to the break site after 4 hours of damage induction (Supplementary Fig. 44-C). This local decrease was observed at both HR- and NHEJ-prone DSBs (Supplementary Fig. 44-C), and, for HR-prone DSBs, appeared stronger for cells which did not display a concomitant distal increase in ATAC-seq signals (Supplementary Fig. 44). This suggests that these two patterns may represent separate events occurring sequentially. This trend was detectable, but less consistent, across individual DSBs for cells subjected to sustained DNA damage induction for 24 hours with AsiSI (Fig. 3B). At both timepoints, the length of DSB-induced accessibility tracts varied substantially between individual damaged cells when focusing on DSBs which are prone to undergo resection and HR (Fig. 3B and Supplementary Fig. 44). By comparison, distal chromatin remodeling was minimal at NHEJ-prone DSBs after either 4 or 24 hours of DSB induction, even when resolved at the level of individual cells (Fig. 3C and Supplementary Fig. 44). This indicates that resection-dependent increase in chromatin accessibility is rare and almost negligible for DSBs mostly relying on NHEJ for repair^48^, even in conditions of sustained DNA damage.

We next investigated how the cell cycle contributed to the observed cell-to-cell heterogeneity of DSB-induced, resection-dependent, chromatin accessibility. For this, each cell in our single-cell ATAC-seq dataset was assigned a specific cell cycle phase using the packages Signac and Seurat^59,60^, where gene activity of specific S and G2/M phase marker genes is inferred from accessibility (Fig. 3D and Supplementary Fig. 44). Cell-cycle-resolved differential ATAC-seq profiles after 24 hours of DSB induction indicated that long tracts of DSB-induced chromatin accessibility were mainly detected for cells in S and G2/M (Fig. 3E), while the local decrease in chromatin accessibility occurred independently of the cell cycle (Supplementary Fig. 44-F). Yet, these patterns remained heterogeneous and were not necessarily detected in each individual cell (Fig. 3F). Distal DSB-induced chromatin accessibility could be observed in a much smaller fraction of G1 cells and was reduced in size (Fig. 3F). Upon 4 hours of DSB induction, DSB-induced chromatin accessibility was increased for cells in G2/M while cells in G1 and S displayed generally similar patterns both in terms of frequency and regarding the length of chromatin accessibility tracts (Supplementary Fig. 44-F). Of note, we failed to detect cells with increased accessibility tracts above 2 kb upon 4 hours of DSB induction, even in G2/M cells, indicating that extended chromatin remodeling is rare in this condition. This suggests the existence of rate-limiting steps in resection-dependent chromatin remodeling and potentially resection itself. Finally, we observed that extensive chromatin remodeling upon damage is a very infrequent event at non-resection-prone DSBs, even when focusing on S and G2/M cells (Fig. 3G, Supplementary Fig. 44). This confirms the existence of substantial differences between damaged loci in their ability to undergo chromatin remodeling following damage, even in specific cell cycle phases in which resection is favored.

Of note, single-cell resolved differential ATAC-seq heatmaps also revealed discrete sites of locally decreased chromatin accessibility, located at various distances from DSBs, and appearing consistently across individual cells (materialized by vertical blue stripes in Fig. 3B-C). These sites of focal decrease in accessibility could be detected for cells G1, S and G2/M (Fig. 3F and Supplementary Fig. 44) and at DSBs which are not prone to undergo HR repair (DSB 199 in Figure 3C and Supplementary Fig. 44 right panel), suggesting that this process is independent of resection per se. Visual inspection suggested that most of these regions were associated with detectable ATAC-seq peaks in untreated cells (see examples in Supplementary Fig. 44), and therefore likely correspond to different classes of regulatory elements such as promoters, enhancers or insulators. This suggests that DSB induction has a broad impact on the chromatin accessibility landscape, not directly related with local repair events, and for which the precise underlying mechanism remains unclear.

Next, we further investigated the directionality of DSB-induced chromatin remodeling at specific sites in individual cells. For this, we sorted single-cell differential ATAC-seq heatmaps based on the upstream-to-downstream signal ratio relative to the DSB position. For practical reasons, this analysis was restricted to cells for which the overall accessibility signals were above an empirically determined threshold, favoring cells in either S or G2/M after 24 hours of DSB induction. Importantly, this revealed that the symmetric pattern observed in bulk experiments is not necessarily representing the most frequent situation at the level of individual cells. Indeed, while we could find examples of DSBs exhibiting multiple cells with largely symmetric chromatin accessibility patterns (see DSB 379 in Fig. 3H), this was not necessarily the case for other DSBs (see DSB 526 in Fig. 3H or DSB 533 in Supplementary Fig. 44). Moreover, we could also identify specific sites for which long range increased accessibility was more frequently unidirectional and biased towards one recurrent side of the DSB (DSB 495, 1129 in Fig. 3I, DSB 600 in Supplementary Fig. 44). Plotting the distribution of the upstream-to-downstream signal ratio at selected HR-prone DSBs indicated that one side of the DSB was slightly favored in many cases (Supplementary Fig. 44), suggesting that the initial chromatin context influences the directionality of the resection-associated increase in chromatin accessibility. Yet, the extent of cell-to-cell variability in directionality (represented by the absolute z-score value of the upstream-to-downstream signal ratio) varied between individual sites (Supplementary Fig. 44). Indeed, while chromatin accessibility profiles at DSB 379 were among the most homogeneous (Fig. 3H, Supplementary Fig. 44), other sites displayed a lot more heterogeneity in terms of directionality across individual cells (for example DSB 608 and 765 in Fig. 3J and DSB 343 in Supplementary Fig. 44). Altogether, our single-cell ATAC-seq data highlights that resection-dependent chromatin accessibility is (i) subjected to extensive cell-to-cell variability at individual damaged loci, which is not entirely explained by the cell cycle, and (ii) is frequently unidirectional, challenging the common view of resection being a generally bidirectional process.

### Sustained DNA damage strongly impacts gene expression leading to distinct switches in cell identity

Next, we set to determine if heterogeneous chromatin remodeling and DNA end resection observed at HR-prone DSBs are coordinated with underlying cell-specific gene expression programs. For this, we performed single-cell Multiome experiments allowing to simultaneously profile chromatin accessibility and gene expression within the same individual nuclei (which we refer to as single cells for convenience). We generated single-cell ATAC-seq and RNA-seq data for more than 11000 cells either untreated, after 4 and 24 hours of DSB induction and upon 24 hours of DSB induction together with MRE11 inhibition by Mirin to inhibit resection (Supplementary Table 2). Pseudo-bulk differential accessibility profiles at DSBs recapitulated the results obtained in previous bulk and single-cell ATAC-seq experiments (Supplementary Fig. 55, to be compared with Fig. 1C, Supplementary Fig. 44, Fig. 3A and Fig. 2I). The matching gene expression data allowed to detect the strong downregulation of genes directly damaged by AsiSI upon DSB induction (Supplementary Fig. 55, first column, genes are listed in Supplementary Table 3), as previously reported^44,61,62^. Moreover, genes previously identified as upregulated after 4 hours of DSB induction in DIvA cells by bulk RNA-seq^63^ (hereby referred to as “DIvA-Up genes”, n=286) and associated with specific Hallmark pathways^64^ such as inflammation and the p53 response (Supplementary Fig. 55), appeared upregulated in damaged cells (Supplementary Fig. 55, second column). Several candidate Gene Ontology (GO) or Pathcards gene sets including DNA damage checkpoint signaling, p53-regulated genes (p53 transcriptional gene network), immune response and inflammation (Inflammatory response, NF-κB signal transduction, Immune response), apoptosis and senescence (Apoptotic process, Cellular Senescence, Senescence and autophagy in cancer) or autophagy also showed transcriptional induction (Supplementary Fig. 55 columns 6 to 13).

Applying dimensionality reduction and Uniform Manifold Approximation and Projection (UMAP) to the single-cell Multiome expression data, we observed a significant overlap between untreated DIvA cells and those subjected to 4 hours of DSB induction, suggesting that mild, short-term, damage does not drastically alter the overall transcriptome (Fig. 4A-B). In contrast, nearly all cells treated for 24 hours clustered separately in low-dimensional space (Fig. 4A-B) indicating that sustained DNA damage substantially impacts overall gene expression. Additionally, MRE11 inhibition led to the appearance of two distinct subsets of cells not found in other conditions (see right-hand side of the UMAP on Fig. 4A, fourth panel on Fig. 4B). A parallel 3’ single-cell RNA-seq experiment (∼8000 cells, Supplementary Table 4) confirmed the downregulation of damaged genes and the concomitant upregulation of DIvA-Up genes (Supplementary Fig. 55, first 2 columns) and candidate Hallmark, Pathcards and GO gene sets (Supplementary Fig. 55, column 3 to 13) upon damage. Moreover, as in our scMultiome data, cells subjected to 24 hours of DSB induction were spatially separated in UMAP representations (Supplementary Fig. 55-G) while non-treated cells largely overlapped with cells exposed to 4 hours DSB induction using either AsiSI or 0.5 µM Etoposide, both generating a comparable number of γH2AX foci (Supplementary Fig. 55). Altogether, these findings indicate that prolonged DNA damage, associated with extended tracts of chromatin accessibility and resection in the cell population, substantially alters gene expression while short-term treatments induce a milder transcriptional response.

**Figure 4:**
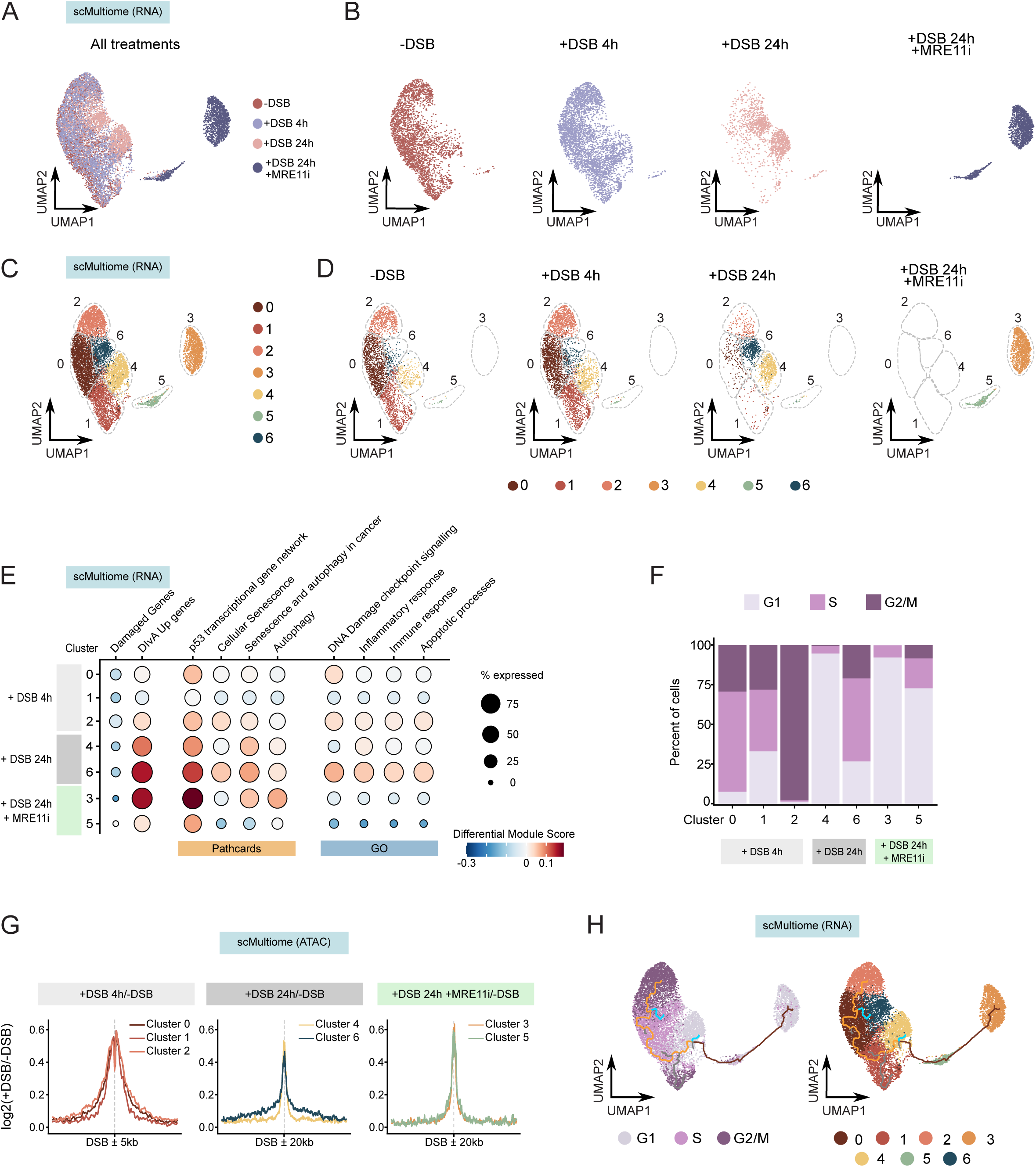
Single-cell analysis of the transcriptional response to DNA double-strand breaks. (A) UMAP projection of 10397 single-cell gene expression profiles from scMultiome experiments. Cells colored according to individual treatments. (B) Same as (A), with individual UMAP projections for each treatment. (C) Same as (A), cells colored according to identified clusters. (D) Same as (C), with individual UMAP projections for each treatment. (E) Dot plots representing the average activity of Pathcards (PC) and GO gene signatures per identified cluster relating to a specific treatment. The size of the dot reflects the percentage of cells expressing the genes of interest while the color bar encodes for up- or down-regulation compared with untreated cells. (F) Histogram representing the percentage of cells in each cell cycle phase per identified cluster relating to a specific treatment. (G) Pseudo-bulk average differential ATAC-seq profiles as log2(+DSB/-DSB) at 80 DSBs for cells in each identified cluster relating to a specific treatment (clusters 0, 1, and 2 on the left panel, clusters 4 and 6 on the middle panel, clusters 3 and 5 on the right panel) on a 10 kb window (± 5 kb from the DSB, left panel) and 40 kb window (± 20 kb from the DSB, middle and right panels). (H) UMAP projections of cells annotated by cell cycle phase (left panel) and cluster identity (right panel) with overlaid trajectory paths representing pseudotime inference highlighting potential cellular transitions across clusters.

Clustering analysis of our scMultiome data identified seven main clusters which were largely dependent on the nature of treatment (Fig. 4C-D). As described above, two of these clusters consisted almost exclusively of Mirin-treated cells (cluster 3 and 5 in Fig. 4C-D), confirming the strong transcriptional phenotype observed in this condition. Similarly, cells subjected to 24 hours of AsiSI-induced damage localized in two distinct clusters (cluster 4, in yellow, and 6, in blue, in Fig. 4C-D), suggesting that cells exposed to sustained DNA damage can primarily adopt one of two distinct cell states or fates. Similar results were obtained from clustering analysis of the scRNA-seq data yielding 6 clusters, two of which (clusters 3 in grey and 4 in dark blue) were largely specific to cells exposed to sustained DNA damage by AsiSI for 24 hours (Supplementary Fig. 66-B). The repression of genes damaged by AsiSI was detectable in each cluster (Fig. 4E and Supplementary Fig. 66), suggesting that distinct DSB-induced transcriptional responses indeed arise in different cell subpopulations experiencing comparable levels of damage.

Cell cycle deconvolution of these scMultiome expression data (Supplementary Fig. 66-E) revealed that each cluster displayed very different cell cycle distributions (Fig. 4F). Indeed, cluster 4, containing about half of cells subjected to 24 hours of DSB induction, as well as cluster 3 and 5, which are characteristic of Mirin-treated cells, were mostly composed of G1 cells (Fig. 4F). In contrast, the other half of 24 hours-treated cells, which were mostly localized in cluster 6, consisted of a majority of cells in S or G2/M (Fig. 4F). Interestingly, this cluster is also characterized by increased expression of several gene sets (DNA damage checkpoint signaling, immune response, inflammation, apoptosis or senescence, Fig. 4E). The other 3 clusters, which are largely composed of both untreated cells and cells treated for 4 hours, seem to represent different stages of a normal cell cycle progression with the largest fraction of G1 cells in cluster 1, a majority of S phase cells in cluster 0 and almost exclusively G2/M cells in cluster 2 (Fig. 4F). Pseudo-bulk differential accessibility at DSBs for treated cells in each individual cluster showed distal increased accessibility at DSBs in cluster 0, 2 or 6, as compared to cluster 1 and 4 (Fig. 4G), in agreement with these cell cycle distributions. Distal increased accessibility was also absent in cluster 3 and 5 in agreement with bulk ATAC-seq data obtained upon MRE11 inhibition (Fig. 2H-I). We could also observe similar cell cycle profiles in clusters identified from our scRNA-seq experiment (Supplementary Fig. 66). A possible interpretation is that cells subjected to sustained DNA damage for 24 hours can undergo cell cycle arrest in either G1 (cluster 4) or S-G2/M (cluster 6). Trajectory analysis of the scMultiome expression data identified a path originating in cluster 4 and ending in cluster 2 (orange line on Fig. 4H) reflecting the progression through a normal cell cycle in untreated cells (according to Fig. 4F). However, cells subjected to 24 hours of AsiSI-induced damage appear to diverge from this path at two different branching points (see blue lines in Fig. 4H) which could be interpreted as cell cycle arrest decisions in either G1 or S-G2/M. This analysis also identifies both clusters of Mirin-treated cells (3 and 5) as originating from G1-arrested cells in cluster 4 (purple line in Fig. 4H), suggesting that S and G2/M cells were lost under these conditions. However, other branching points in these trajectories are more difficult to explain (in dark grey in Fig. 4H). Thus, single-cell transcriptome data revealed that cells subjected to sustained DNA damage experience an acute DSB-induced transcriptional response and appear to undergo switches in cell identity. Moreover, the adoption of these two distinct cell fates, suggestive of cell cycle arrest decisions occurring in either G1 or S-G2/M, is associated with very different resection-dependent chromatin remodeling profiles at DSBs.

### Coordinated resection-dependent chromatin remodeling at multiple DSBs is associated with acute signaling and increased expression of inflammation-related genes

We next used our scATAC-seq data to explore the possibility that DSB-induced and resection-dependent chromatin remodeling could occur in a coordinated fashion across multiple DSBs in subsets of cells, a behavior which may contribute to some of the transcriptional phenotypes observed in single-cell transcriptome experiments. Ordering single-cell-resolved differential chromatin accessibility heatmaps by decreasing signal for a given DSB (DSB 526) indeed suggested that extensive increase in accessibility could occur concomitantly at several other DSBs at least in a fraction of cells (Figure 5A, Supplementary Fig. 77), especially those in S and G2/M (Supplementary Fig. 77).This behavior would also be in agreement with the previously reported simultaneous binding of RAD51 at multiple AsiSI-induced DSBs in individual cells^65^. To further explore this coordinated chromatin remodeling behavior, we computed a differential accessibility score over all 80 AsiSI-induced DSBs in each individual cell over a single genomic bin (± 5 kb at 4 hours and ± 40 kb at 24 hours) and the resulting matrices were subjected to hierarchical clustering. This differential accessibility matrix revealed that a significant fraction of cells (about ⅓) showed a simultaneous increase in accessibility at multiple DSBs after 24 hours of DSB induction (right side of Fig. 5B), mostly involving HR-prone DSBs (appearing in orange on Fig. 5B). We also found a second group of cells for which DSB-induced accessibility is much lower (left side of Fig. 5B) and a third one for which this was barely detectable (middle of Fig. 5B). A similar pattern could also be observed after 4 hours of DSB induction (Supplementary Fig. 77) but with a less clear distinction between the different subgroups. We subsequently focused on the differential accessibility after 24 hours of induction at a subset of 10 DSBs which are prone to undergo extensive resection and chromatin remodeling, focusing on S and G2/M cells to reduce the potential interference from the cell cycle. Doing so, we could indeed define subsets of cells characterized by a coordinated strong increase in resection-dependent chromatin accessibility at several DSBs (labeled as HAM for <u>H</u>igh <u>A</u>ccessibility at <u>M</u>ultiple DSBs) as well as a group of cells for which chromatin accessibility was either unchanged or decreased post DSB induction across all tested sites (labeled LAM for <u>L</u>ow <u>A</u>ccessibility at <u>M</u>ultiple DSBs) (Fig. 5C-D). Similar cell subpopulation of HAM and LAM cells could also be identified upon 4 hours of DSB induction (Supplementary Fig. 77).

**Figure 5:**
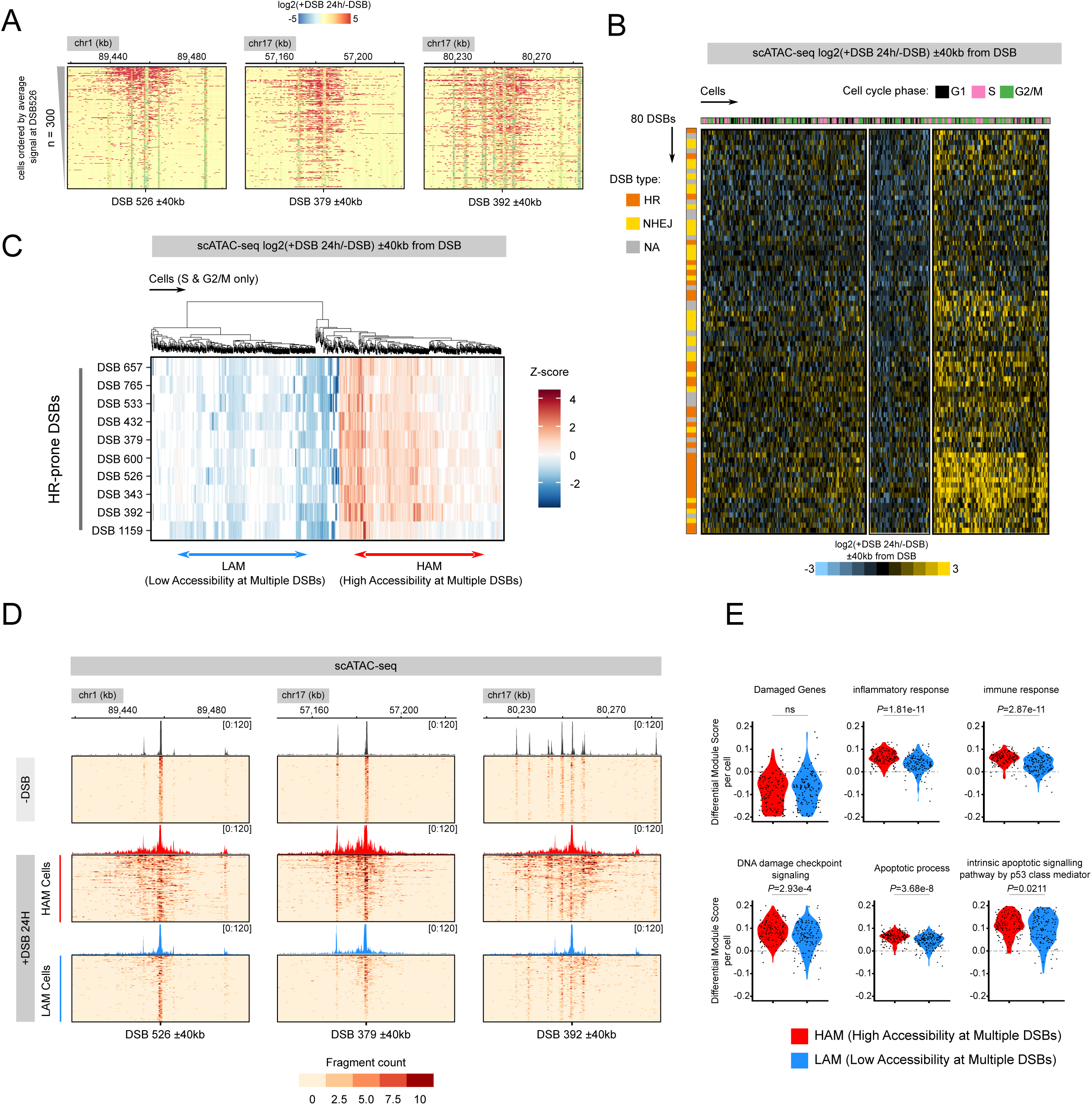
A subset of cells display coordinated DSB-induced chromatin remodeling associated with DNA end resection and specific gene expression programs. (A) Single-cell differential chromatin accessibility profiles as log2(+DSB/-DSB) on an 80 kb window (± 40 kb from the DSB) after 24 h of DSB induction for DSBs 526, 379, and 392. Each heatmap displays 300 cells that are ordered by the mean differential score at DSB 526. (B) Heatmap representing the mean differential log2(+DSB/-DSB) chromatin accessibility scores for cells after 24 h of DSB induction on an 80 kb window (± 40 kb from the DSB). Columns correspond to individual cells colored by cell cycle phase (G1 in black, S in pink, G2/M in green) and divided into three clusters visible from hierarchical clustering. Rows correspond to DSBs colored by preferred repair pathway where applicable (HR in orange, NHEJ in yellow, NA in grey). Color bar refers to the increase (yellow) or decrease (blue) in accessibility at a given DSB in a cell. (C) Hierarchical clustering of cells with High Accessibility at Multiple DSBs (HAM, in red, n=140 cells) or Low Accessibility at Multiple DSBs (LAM, in blue, n=143 cells) in chromatin accessibility among S and G2 cells at a subset of HR-prone DSBs after 24 h of DSB induction. Differential accessibility was computed as log2(+DSB/-DSB) in a single 100 kb bin and transformed into Z-scores (see methods). (D) Single-cell fragments profiles of non-treated, HAM and LAM sub-populations described in (C) (n=100) on an 80kb window (± 40 kb) at DSBs 526, 379, and 392. Color bar indicates sum of fragments in a given bin. (E) Violin plots representing the expression of specific Gene Ontology (GO) gene sets in individual cells in the HAM and LAM sub-populations from S and G2/M cells after 24 h of DSB induction. P-values were obtained from two-sided Wilcoxon tests.

We next aimed to determine if these coordinated patterns in DSB-induced chromatin accessibility could be associated with specific cellular responses using single-cell gene expression data. For this, we isolated specific subsets of S or G2/M cells characterized by simultaneous strong increase (HAM) or weak decrease (LAM) in DSB-induced chromatin accessibility from our Multiome experiments after 24 hours of DSB induction (Supplementary Fig. 88-B). Interestingly, we found that the downregulation of genes damaged by AsiSI is not significantly different between HAM and LAM cells (Fig. 5E, top left panel), suggesting that both subgroups experienced similar amounts of damage. GO pathway enrichment of differentially expressed genes between these two subpopulations suggested that HAM cells were characterized by increased expression of genes involved in lipid metabolism, intracellular membrane trafficking (organelle fusion, vesicle mediated transport, exocytic process) and regulators of autophagy (Supplementary Fig. 88). By comparison, LAM cells display increased expression of few genes associated with cell division and mitosis (Supplementary Fig. 88), implying that this subpopulation may have progressed further into the cell cycle compared to HAM cells. When comparing the expression of candidate genes sets between these two populations, we found that HAM cells displayed significantly higher expression for pathways related to inflammation, immune response and DNA damage checkpoint signaling (Fig. 5E). This was also true, to a lesser extent, for the activation of genes involved in apoptosis or senescence and p53 responsive genes (Fig. 5E, Supplementary Fig. 88). This effect appeared to be specific since no differences in gene expression were detected for control Hallmark pathways (Oxydative Phoshorylation or Unfolded Protein Response) (Supplementary Fig. 88). Therefore, our results identify two subsets of cells showing antagonistic chromatin accessibility patterns at multiple DSBs associated with specific gene expression programs. In particular, cells showing extended resection tracks at multiple DSBs display increased checkpoint signaling and expression of genes involved in inflammation as well as a potentially altered intracellular organization and internal membrane trafficking.

## Discussion

In this study, we employed high-resolution bulk and single-cell ATAC-seq approaches to chart the elaborate reshuffling of chromatin structure emerging upon DSB induction in mammals. Remarkably, we report a long-range increase in chromatin accessibility distal to DSBs which depends on functional resection, is mostly restricted to a subset of DSBs undergoing homologous recombination, and is dynamic across the cell cycle. Importantly, our single cell deconvolution revealed that resection-dependent chromatin accessibility displays substantial cell-to-cell heterogeneity at individual damaged loci and is frequently unidirectional, suggesting that resection itself is heterogenous from one cell to another. Moreover, we identify subpopulations of S and G2/M cells with similar resection-dependent chromatin accessibility patterns occurring simultaneously at multiple DSBs which are associated with specific gene expression programs, including DNA damage checkpoint signaling, inflammation and apoptosis. By conducting one of the very first investigations of DSB repair by single-cell genomics together with^65^, we provide results highlighting the intricacy and heterogeneity of chromatin remodeling events occurring upon DSB repair at individual DSBs, as well as the ability of cells to coordinate chromatin remodeling events at multiple DSBs in the same cell and revealing unexpected connections between resection and the transcriptional response to genotoxic insults.

### Multistep remodeling of chromatin accessibility in response to DSBs in mammals

Our chromatin accessibility data highlight a composite pattern in which chromatin accessibility is sharply reduced on ∼500bp in the immediate vicinity of DSB sites and increases distally in a resection-dependent manner. At this stage, the precise mechanism responsible for the decrease in chromatin accessibility at DSBs remains unclear. An attractive possibility, in agreement with our ChIP-seq data, is that decreased ATAC-seq signals represent footprints of the binding of factors such as MRE11, KU, DNA-PKcs or ATM to DNA ends, similar to those observed at transcription factor binding sites^40^. These footprints can also be interpreted as productive, or at least ongoing, attempts of NHEJ repair. Of note, such Tn5 transposition patterns would hinder our ability to detect DSB-proximal nucleosome depletion, as reported in yeast or mammalian systems^38,39,66,67^. By contrast, bulk ATAC-seq experiments using specific inhibitors showed that DSB-induced chromatin accessibility is initiated in an MRE11- and ATM-dependent manner, and is occurring on the damaged chromosome prior to RAD51 filament assembly. Altogether, this favors a two-step model in which chromatin accessibility first decreases at the break due to the binding of DSB sensors or NHEJ factors and is subsequently increased at a larger distance in cases where the resection machinery is engaged, which would be in agreement with a sequential hand-off model for DSB repair^68^.

At this stage, it cannot be excluded that increased ATAC-seq signals reflect altered nucleosomal structures, such as H2A–H2B dimer removal or H2AZ incorporation, which favor the activity of resection nucleases on chromatinized templates *in vitro*^24^, and may be more permissive to Tn5-mediated insertions. However, a simpler interpretation is that increased accessibility reflects nucleosome eviction occurring concomitantly with end processing, facilitating the access to resection nucleases or stimulating their activity, as reported in yeast^28^. Yet, the precise high-resolution dynamics of nucleosomes themselves upon resection at multiple DSBs in mammals remains to be formally established. Moreover, it remains unclear whether increasing chromatin accessibility distal to DSBs is a requirement for the access and/or activity of nucleases, or if it is occurring as a consequence of DNA end processing. In yeast, chromatin remodeling complexes appear largely dispensable for MRE11 incisions^21^, while nucleosome eviction concomitant with resection requires RSC and SWI/SNF^28^. Fun30 favors resection by both MRX/Sae2 and Exo1 during yeast meiosis, but has little impact on resection-dependent nucleosome eviction in vegetative cells^28^. Instead, Fun30, and its human homolog SMARCAD1, rather act by evicting anti-resection factors^29,31,33^. This suggests that the requirement for chromatin remodeling complexes might differ at each step of the resection process and can also be context dependent. This may render the precise identification of the remodeler(s) involved in DSB resection in mammals difficult due to higher complexity and potential redundancies between these complexes^27^.

### DSB-induced chromatin accessibility is restricted to a subset of DSBs, is dynamic across the cell cycle, and displays substantial cell-to-cell heterogeneity at individual damaged loci

ATAC-seq data revealed that, while the local, ∼ 500bp decrease in chromatin accessibility is visible at most AsiSI-induced DSBs, the extensive distal chromatin remodeling is only notably detectable at a subset of DSBs in cells, especially upon sustained DSB induction for 24 hours. This increased distal accessibility at specific DSBs was also evident at the single-cell level, allowing the detection of potentially rare events, particularly in S and G2/M cells, where resection is believed to be inherently favored. These results agree with previous work showing that DSBs localized in actively transcribed genes are more prone to undergo resection and HR repair, independent of DSB induction efficiency^43,48,69^, reinforcing the idea that specific mechanisms ensure the efficient repair of DSBs occurring in active genes, which represent physiological hotspots in a variety of contexts (reviewed in^70^). Our results also support the broader notion that chromatin reconfiguration events required for efficient DSB repair are strongly determined by the properties of the damaged locus.

Our single-cell ATAC-seq approach resolved DSB-induced chromatin accessibility dynamics across the cell cycle without the need for experimentally-induced synchronization. While this confirmed that long tracts of resection-dependent chromatin remodeling were more readily observed in cells in S and G2/M, we could also detect narrower and less frequent events of DSB-induced increases in chromatin accessibility in G1 cells. Indeed, several studies have reported that resection can occur in G0 or G1^12–15^. Another non-mutually exclusive possibility is that fill-in synthesis by CST-Polα (CTC1-STN1-TEN1-DNA polymerase α), which counteracts resection^71^ and is most prominent in G1^14^, may contribute to increased chromatin accessibility in G1, a hypothesis requiring further investigation.

Interestingly, resection-dependent chromatin accessibility tracts at HR-prone DSBs were heterogeneous in length across the cell population and increased noticeably with DSB induction time. These heterogeneous lengths for chromatin accessibility in individual cells could represent different steps in the progress of resection at DSBs induced at different times in each cell. Indeed, induction is not synchronous in the DIvA model, allowing AsiSI-induced DSBs to undergo several cycles of cleavage and repair. Alternatively, heterogeneous chromatin accessibility tracts could be explained by the presence of specific boundaries constraining the extent of resection *in vivo* which would need to be bypassed, a process which could occur at different rates in individual cells. Importantly, while bulk analyses suggested that resection and nucleofilament assembly at DSBs is mostly bidirectional, our single-cell data rather revealed that resection-dependent chromatin accessibility can frequently be unidirectional. Interestingly, individual DSBs display different behaviors, with some loci being strongly biased towards a specific side, while others did not exhibit side specificity, *i.e.* resection-dependent accessibility can occur with equal probabilities on one side or the other. Such a behavior is not necessarily in agreement with the existence of strong DNA-encoded static boundaries regulating DNA-end processing and rather suggest that specific factors dynamically localizing to the damaged locus before or after break induction, such as RNA Pol II or cohesin^44,47^, could constrain the activity of the resection machinery at different locations in individual cells. Importantly, this could affect the ability of each DNA end to engage the recombination process, potentially leading to mutagenic DNA repair outcomes, as previously reported in yeast using a synthetic substrate undergoing asymmetric resection and showing an increase in non-reciprocal translocation events^72^. Further technological developments are necessary to precisely chart resection endpoints in single cells and determine how other molecular machineries involved in DNA transactions may impact the progression of resection and the execution of HR at individual DNA ends.

### Coordination of extended resection-dependent chromatin accessibility, cell fate decisions and DSB cytotoxicity

Our transcriptome analysis revealed that DSB induction with a low dose of etoposide or AsiSI for 4 hours can activate several expected DNA damage responsive genes and pathways. Yet, the overall response remained relatively mild, with a moderate impact on cell cycle distribution in damaged cells. By contrast, DSB induction with AsiSI for 24 hours caused strong transcriptional phenotypes, with two distinct groups of cells accumulating either in G1 or in S/G2. While both subsets show strong activation of p53-responsive genes, the S/G2 subpopulation showed upregulation of many gene sets, suggestive of an acute signaling or checkpoint activation. This echoes previous work showing that breaks which are difficult to repair, combined with increased signaling, can trigger early decisions of permanent cell cycle withdrawal^73^. At this stage, it is not clear if cells in each subset have remained blocked in a specific phase due to unmanageable DSB loads or if some cell cycle transitions occurred during the course of the experiment. Moreover, these conclusions are based on gene expression measurements and do not assess absolute protein levels, subcellular localizations, post-translational modifications or cellular DNA content. Nevertheless, these data suggest that the cellular response to DNA damage differs across the cell cycle, which can potentially influence the cytotoxicity of DSBs^74^. More work is needed to understand whether damaged cell subpopulations have different abilities to recover from checkpoints and re-enter the cell cycle, and how this may impact the long-term viability of daughter cells.

Additionally, single-cell chromatin accessibility data highlighted a subpopulation of cells characterized by simultaneous long-range resection-dependent chromatin accessibility at multiple DSBs. This agrees with a recent study identifying similar simultaneous binding of RAD51 at multiple DSBs in single cells^65^ which suggested that repair might be coordinated at multiple DSB sites. Interestingly, while such coordination likely involves association with specific nuclear micro-environments^63,75^ and the formation of multi-DSB hubs^63,65^, it appears to be detectable as coordinated resection-dependent chromatin remodeling events occurring at the vicinity of the break. Interestingly, cells displaying coordinated resection-dependent chromatin accessibility at multiple DSBs also showed increased activation of genes involved in DNA damage checkpoint signaling, inflammation and immune response and, to a lesser extent, apoptosis or senescence, which could indicate cells facing unmanageable DSB loads rather than coordinated repair. This agrees with the previous observation that, in response to IR, cells permanently withdrawing from the cell cycle in G2 accumulate long stretches of resected DNA^73^, suggesting that excessive single-strand DNA might be intrinsically cytotoxic. However, during normal mouse meiosis, spermatocytes are estimated to accumulate 400-700 kb of resected DNA^76^, which is compatible with cell viability, at least in this context. Alternatively, secondary products of end processing may directly participate in activating intracellular signaling pathways. Indeed, in response to genotoxic treatments, the accumulation of cytosolic DNA fragments generated by EXO-1 and DNA2-dependent resection can lead to activation of type I interferon signaling^77^. Similarly, in specific contexts, MRE11 activity is associated with the accumulation of cytoplasmic DNA, micronuclei formation, and the expression of type I interferons and pro-inflammatory cytokines^78,79^. Further work is required to better understand the relationship between excessive resection, the activation of these signaling pathways, and the initiation of specific processes such as apoptosis or senescence. This would allow a better understanding of the basic mechanisms ensuring genome stability and to better design chemotherapeutic approaches relying on DSB induction.

## Acknowledgments

We thank the Genomics Core Facility (GeneCore) of EMBL for high-throughput sequencing. Funding in G.L laboratory is provided by grants from the European Research Council (ERC-AdG-101019963), the Agence Nationale pour la Recherche (ANR-21-CE12-0033-03), the Association Contre le Cancer (ARC), the association Robert Debré and the Fondation Bettencourt-Schueller. This work was also supported by a government grant managed by the Agence Nationale de la Recherche under the France 2030 program, with the reference numbers ANR-24-EXCI-0001, ANR-24-EXCI-0002, ANR-24-EXCI-0003, ANR-24-EXCI-0004, ANR-24-EXCI-0005”. S.C was a recipient of a PhD fellowship from the Joint Training and Research Programme on Chromatin Dynamics & the DNA Damage Response (H2020 ITN aDDRess, grant N° 812829). C.A was a recipient of a fellowship from the Fondation pour la Recherche Médicale (FRM FDT201904007941). R. C is a recipient of a PhD fellowship from the Fondation pour la Recherche Médicale (ECO202306017314). T.C is an INSERM researcher.

## Authors contributions Statement

S.C, A.L.F, C.A, R.C. and T.C performed and analyzed experiments. S.C and V.R performed bioinformatic analyses of high-throughput sequencing datasets. T.C and G.L conceived and supervised the study. T.C wrote the manuscript with the help of S.C and G.L. All authors commented and edited the manuscript. A.L.F and C.A contributed equally to this work.

## Methods

### Cell Culture and treatments

DIvA (AsiSI-ER-U2OS) cells were cultured in Dulbecco’s modified Eagle’s medium (DMEM) supplemented with antibiotics, 10% FCS (Invitrogen) and 1µg/mL puromycin at 37°C under a humidified atmosphere with 5% CO2. AsiSI-dependent DSBs were induced by the addition of 300 nM 4-OHT (Sigma, H7904) into the medium for 4 h or 24 h, or by the addition of etoposide (Sigma, E1383) at 0.5 µM for 4 h. Inhibitors were added to the medium 1 hour prior to 4-OHT treatment (see Supplementary Table 5 for catalog number and final concentrations). siRNA transfections were performed using 4D-Nucleofector from Lonza either with the SE-small kit (V4XC-1032) for 1 million cells or SE-large kit (V4XC-1024) for 5-20 million of cells, according to the manufacturer’s instruction. Cells were collected 48 h after siRNA transfection. The siRNA used are listed in Supplementary Table 6.

### Chromatin Immunoprecipitation

ChIP experiments were performed as in^42,48^. Briefly, cells were crosslinked with 1% formaldehyde (Sigma-Aldrich, F8775) for 15 min at room temperature and quenched with 0,125 M glycine for 5 min. Cells were washed with 1X PBS and harvested by scrapping. Cell pellets were resuspended in cellular lysis buffer (5 mM Pipes pH 8, 85 mM KCl, 0.5% NP-40) and homogenized with a Dounce homogenizer. Nuclei were harvested by centrifugation and resuspended in nuclear lysis buffer (50 mM Tris pH 8.1, 10 mM EDTA, 1% SDS) and sonicated using a Branson Sonifier 250 (at a power setting of 5 and 50% duty cycle) to obtain fragments of about 500 bp. Samples were diluted 10 times in dilution buffer (16.7 mM Tris pH 8, 167 mM NaCl, 1.2 mM EDTA, 1.1% Triton X-100, 0.01% SDS) and each aliquot of 200 µg of chromatin was precleared with 100 µL of 50% protein-A (Pierce) and protein-G (Sigma) beads, (previously blocked with 500 µg of BSA) for 2 hours at 4°C on a rotating wheel. 100 µL of pre-cleared chromatin was kept at −20°C as input sample and 10-200 µg was incubated with specific antibodies at 4°C on a rotating wheel overnight (antibodies used are outlined in Supplementary Table 7). The next day, 100 µL of pre-blocked protein A/G beads were added to each sample for 2 hours at 4°C on a rotating wheel. Beads were washed once with dialysis buffer (50 mM Tris pH 8.1, 2 mM EDTA, 0.2% Sarkosyl), five times with wash buffer (100 mM Tris pH 8.8, 500 mM LiCl, 1% NP-40, 1% NaDoc) and twice in TE buffer (Tris 10 mM pH 8, 1 mM EDTA). Immunoprecipitated complexes were resuspended in 200 µL of TE and input samples were adjusted to 200 µL. Next, 0.5 µL of RNase A (30mg/mL) was added during 30 min at 37°C. Following addition of 0.5% SDS, samples were incubated overnight with shaking at 70°C to reverse the crosslink. Next, proteinase K was added (200 µg/mL final) and samples were incubated for 1h30 at 45°C. DNA was purified by phenol:chlorophorm extraction followed by ethanol precipitation. For ChIP-Seq, multiple ChIP experiments were pooled, sonicated for 7 cycles (30 sec on, 30 sec off, high setting) with a Bioruptor (Diagenode), then concentrated with a vacuum concentrator (Eppendorf). About 10 ng of purified DNA was subjected to library preparation (New England Biolabs, NEBNext Ultra II DNA Library kit, E7645 and NEBNext Multiplex Oligos for Illumina, E6440) and sequenced with single-end 100 bp sequencing reads on a NextSeq2000 at EMBL GeneCore (Heidelberg, Germany).

### Bulk ATAC-seq

Following treatment with inhibitors and OHT, cells were washed twice with PBS and harvested with Trypsin and resuspended in 5 ml of medium. Cells were then counted and resuspended at 5×10^5^ cells/ml in 1X PBS. 1ml of suspension was then centrifuged at 600g for 5 minutes at room temperature and resuspended in 500 µl of ice cold 1X PBS before being centrifuged again at 600g for 5 minutes at 4°C. Nuclei were isolated by resuspending cells in 500 µl cold Lysis buffer (10mM Tris-HCl pH 7.5; 10mM NaCl, 3mM MgCl2, 0.1% NP-40, 0.1% Tween 20, 0.01% Digitonin). After a 3 minutes incubation on ice, 10 ml of cold lysis buffer lacking NP-40 and Digitonin was added to stop the reaction. Nuclei were collected by centrifugation at 600g for 10 minutes at 4°C and resuspended in 165 µl of ice cold 1X PBS. The equivalent of 50000 nuclei (16.5 µl) were resuspended in 50 μl of transposition mix consisting of 25 µl 2X TD buffer (Illumina #15027866), 2.5 µl Transposase TDE1 (Illumina #15027865), 0.1 µl Digitonin 5%, 0.5 µl Tween 20 10%, 5.4 µl H_2_0 and incubated at 37°C for 30 minutes in a thermomixer at 700rpm. Transposed DNA was then purified using the Qiagen MinElute PCR Purification Kit (Cat. No 28004) following manufacturer’s instructions. Library preamplification was performed by combining 20 µl of transposed DNA with 2.5 µL of 25 µM barcoded i7 and i5 primers (Supplementary Table 8) and 25 µL of NEBNext HiFi 2x PCR Master mix (New England Biolabs, M0541). Samples were subjected to PCR preamplification using the following conditions (72°C for 5 minutes, 98°C for 30 seconds then 5 cycles of 98°C for 30 s, 63°C for 10 s and 72°C for 1 minute). To estimate the number of cycles required for PCR amplification, 4 µL of the preamplification reaction were combined with 0.5 µL of 25 µM barcoded i7 and i5 primers and 6 µl Kapa HiFi Hot Start Real-Time PCR mix (KM2702) and subjected to 20 cycles of real time PCR using the following cycling conditions (98°C for 30 seconds then 20 cycles of 98°C for 30 s, 63°C for 10 s, 72°C for 1 minute). The number of required cycles were determined by taking the cycle number needed for ¼ of the maximum fluorescence intensity. Individual samples were further PCR amplified for the required amount of cycles (usually 5) with the following conditions (98°C for 30 seconds then cycles of 98°C for 30 s, 63°C for 10 s, 72°C for 1 minute, followed by 72°C for 1 minute). Libraries were purified using Ampure XP (Beckman Coulter, A63881) or SPRIselect (Beckman Coulter, B23318) as follows: large fragments were first removed with a 0.4X samples to beads ratio and the library was recovered with a 1.1X sample to beads ratio. Libraries were quantified with a Quantus Fluorometer (Promega) using the QuantiFluor® dsDNA System (Promega, E2670) and run on a High Sensitivity DNA chip on the Bioanalyzer 2100 instrument (Agilent Technologies). Finally, samples were sequenced at Genecore (EMBL) on a NextSeq500 in 2*75 base pairs mode or on a Nextseq 2000 instrument in 2*50 base pairs mode.

### CUT&Tag

CUT&Tag was performed as described in^80^ with minor modification. Following OHT treatment, cells were harvested using Trypsin and collected by centrifugation at 600g for 5 minutes at room temperature. Cell pellets were resuspended in 5 mL of ice-cold NE1 buffer (20 mM Hepes pH 7.9, 10 mM KCl, 20% Glycerol, 0.5 mM Spermidine, 0.1% Triton X-100) with 1X EDTA-free

Complete protease (Merck 11873580001) and phosphatase inhibitors (Sigma-Aldrich, P5726). After 10 minutes of incubation on ice, nuclei were collected by centrifugation at 1000g, 4°C for 5 minutes and resuspended in PBS containing 0.5 mM Spermidine and protease and phosphatase inhibitors. Nuclei were crosslinked by adding formaldehyde (Methanol Free, Electron Microscopy Science, 15710) at 0.2% final concentration for 2 minutes at RT. Formaldehyde was quenched by adding glycine (125 mM final) and nuclei were recovered by centrifugation at 1000g, 4°C for 10 minutes. Nuclei were resuspended in Antibody buffer (20 mM Hepes pH 7.5, 150 mM NaCl, 0.5 mM Spermidine, 0.1% BSA) with protease and phosphatase inhibitors at around 2.10^6^ nuclei/mL. For each individual experiment, about 100000 nuclei were bound to 3.5 µL of activated Concanavalin A-Conjugated Paramagnetic Beads (CUTANA, Epicypher, 21-1411) in Antibody buffer for 15 minutes at room temperature on a rotating wheel. Beads containing nuclei were washed once with Antibody buffer, resuspended in 200 µL of Antibody buffer, transferred to 8-tubes 0.2 mL PCR strips and primary antibodies were added (typically 1 µg, described in Supplementary Table 7). Samples were incubated overnight on a rotating wheel at 4°. The following day, 1 µg of the appropriate secondary antibody was added to the samples and incubated for 1 h at room temperature on a rotating wheel. Nuclei were washed in Wash Buffer 150 (20 mM Hepes pH 7.5, 150 mM NaCl, 0.5 mM Spermidine) with protease and phosphatase inhibitors and resuspended in 200 µL of Wash 300 (Wash Buffer 150 with 300 mM NaCl final). 2 µL of pAG-Tn5 (CUTANA, Epicypher, 15-1117) was added, and samples were placed for 1 h on a rotating wheel at room temperature. After washing with the Wash 300 solution, nuclei were resuspended in 50 µL of Tagmentation buffer (Wash 300 with 10 mM MgCl2). Samples were then incubated at 37°C for 1h in a Thermocycler. Following incubation, the liquid was removed, and 5 µL of SDS Release buffer (10mM TAPS pH 8.5, 0.1% SDS) was dropped on the nuclei/beads mixture and samples were incubated for 1h at 58°C. 15 µL of 10 mM TAPS pH 8.5 with 0.67% Triton-X100 was then added to the mixture to quench the SDS. Libraries were amplified using 2.5 µL of 10 µM barcoded i7 and i5 primers (Supplementary Table 8) together with 25 µL of NEBNext HiFi 2x PCR Master mix (New England Biolabs, M0541) using the following conditions (58°C for 5 minutes, 72°C for 5 minutes, 98°C for 5 minutes then 15-21 cycles of 98°C for 15 s, 63°C for 10 s and 72°C for 1 minute). Samples were purified using SPRISelect beads (Beckman Coulter, B23318). Large fragments were first removed with a 0.4X samples to beads ratio and the CUT&Tag library was recovered with a 1.3X sample to beads ratio. Libraries were quantified with a Quantus Fluorometer (Promega) using the QuantiFluor® dsDNA System (Promega, E2670) and run on a High Sensitivity DNA chip on the Bioanalyzer 2100 instrument (Agilent Technologies). Finally, samples were sequenced at Genecore (EMBL) on a Nextseq 2000 instrument in 2 x 50 base pairs mode.

### Single-cell ATAC-seq, single-cell RNA-seq and single-cell Multiome experiments

For single-cell Multiome and ATAC-seq experiments, nuclei were isolated using the cell suspension isolation protocol from 10X Genomics (protocol CG000124, Rev F) with a lysis time of 5 minutes. Nuclei were loaded at an expected recovery of 5000 nuclei. Both ATAC-seq libraries were generated using the Chromium Next GEM Single Cell ATAC Library and Gel Bead Kit v1.1 (PN-1000175) or Chromium Next GEM Single Cell Multiome ATAC + Gene Expression Reagent Bundle (PN-1000285) and were further purified by double-sided clean up using SPRISelect beads (Beckman Coulter, B23318). Cells for single-cell RNA-seq were loaded at an expected recovery of 10000 cells and libraries were generated using the Chromium Next GEM Single Cell 3ʹ Kit v3.1 (PN-1000269). Libraries were quantified with a Quantus Fluorometer (Promega) using the QuantiFluor® dsDNA System (Promega, E2670) and run on a High Sensitivity DNA chip on the Bioanalyzer 2100. Finally, samples were sequenced at Genecore (EMBL) on a Nextseq 2000 (scMultiome) or Nextseq500 (scRNA-seq and scATAC-seq) instrument. Sequencing was performed in paired-end, dual indexing mode according to instructions from 10X Genomics for single-cell ATAC-seq (Insert Read 1 : 71 bp, i7 Index : 8 bp, i5 Index : 16 bp, Insert Read 2 : 71 bp), single-cell Multiome ATAC-seq (Insert Read 1 : 53 bp, i7 Index : 8 bp, i5 Index : 24 bp, Insert Read 2 : 53 bp) and 3’ single-cell RNAseq/single-cell Multiome expression (Read 1 10X Barcode+UMI : 28 bp, i7 Index : 10 bp, i5 Index : 10 bp, Read 2 insert : 100 bp).

### Immunofluorescence

Cells were plated in glass coverslips, then submitted to a pre-extraction with ice cold buffer (20 mM HEPES pH 7.5; 20 mM NaCl; 5 mM MgCl2; 1 mM DTT; 0.5% NP40) for 20 min on ice. Cells were fixed with 4% paraformaldehyde for 15 min and blocked with PBS-BSA 3% for 30 min at room temperature. Cells were then incubated with antibody against γH2AX (JBW301, 05-636, Millipore, 1:1000) and RAD51 (Santa cruz sc8349, 1:200) overnight at 4 °C (antibodies are described in Supplementary Table 7). Cells were washed three times in PBS-BSA 3% and incubated with secondary antibody for 1 h. After three washes (one PBS-BSA 3% and two PBS), nuclei were stained with Hoechst 33342 (Sigma). Image acquisition was performed using MetaMorph on a wide-field microscope equipped with a cooled charge-coupled device camera (CoolSNAP HQ2), using a ×40. Quantification was performed using Columbus, an integrated software to the Operetta automated high-content screening microscope (PerkinElmer). Hoechst stained nuclei were selected according to the B method and appropriate parameters, such as the size and intensity of fluorescent objects, were applied to eliminate false-positives. Then γ-H2AX or RAD51 foci were detected with the D method with the following parameters: detection sensitivity, 1; splitting coefficient, 1; background correction, >0.5 to 0.9.

### Western Blots

For CtIP western blots, pellets from a single 10 cm plate were resuspended in 100 µL of LDS 1X NuPAGE LDS (ThermoFisher, NP0007) containing 5 µL of Benzonase (GENIUS™ Nuclease, SantaCruz, sc-202391) for 10 minutes at room temperature. 11 µL of 10X NuPAGE Sample Reducing agent (ThermoFisher, NP0009) were then added to each sample which were denatured for 10 min at 70°C prior to loading. For EXO1 and DNA2 western blots, pellets from a single 10 cm plate were resuspended in 50 µL of LDS 1X NuPAGE LDS for 10 minutes on ice, then incubated with 1 µL of Benzonase for 20 minutes at room temperature. 5 µL of NuPAGE Sample Reducing agent was added to each sample which were denatured for 5 min at 95°C prior to loading. 20 µL (CtIP) or 10 µL (EXO1/DNA2) of samples were resolved on NuPAGE Bis Tris 4-12% precast gels (ThermoFisher NP0321) with 1X MOPS buffer (ThermoFisher NP0001), and transferred on PVDF membranes using the Trans-Blot Turbo System (Bio-Rad) according to the manufacturer’s instructions. Membranes were blocked in TBS containing 0.1% Tween 20 and 4% BSA (CtIP) or 3% skimmed dried milk (EXO1/DNA2) for 1 hour at room temperature, followed by overnight incubation at 4 °C using primary antibody (Supplementary Table 7) diluted in blocking solution. Washed membranes were subsequently incubated with appropriate mouse or rabbit HRP-coupled secondary antibodies (Sigma, A2554 and A0545) at 1:10,000 in blocking solution for 1 hour at room temperature. Proteins were revealed by chemiluminescence (Super Signal West Dura Extended Duration Substrate, ThermoFisher, 34075 or Clarity Western ECL Substrate, Bio-Rad, 1705061), images were acquired with a ChemiDoc Touch Imaging System (Bio-Rad) and analyzed with Image Lab (Bio-Rad)

### High-throughput sequencing data processing

The quality of each raw sequencing file (fastq) was verified with FastQC. All sequencing data, unless specified, were aligned using bwa-mem (https://bio-bwa.sourceforge.net/bwa.shtml) to the human genome (hg19) and further sorted and duplicates removed using samtools (http://www.htslib.org/). Bigwig coverage tracks for each file was generated using *bamCoverage* from deeptools (https://deeptools.readthedocs.io/en/latest/) and, unless specified, were normalized by the total read count for each sample. Differential coverage between two conditions was performed using *bigwigCompare* from deeptools with setting the bin size parameter to 50 bp and the operation parameter as “log2”. Further data analysis specifications for each sequencing protocol are outlined below.

*Single-cell ATAC-seq*

Processing of individual single-cell ATAC-seq datasets was conducted using the Cell Ranger ATAC tool from 10X Genomics using default parameters (https://support.10xgenomics.com/single-cell-atac/software/pipelines/latest/what-is-cell-ranger-atac). This step involves alignment to the human genome (hg19), unique molecular identifier (UMI) identification, and low-quality cell filtering to output a peak-by-cell-matrix to be used for downstream analysis in R using the Signac package^59^. Individual matrices were imported to R and directly merged to generate a Signac object by identifying a common peak set as outlined in the tutorial (https://stuartlab.org/signac/articles/merging). Following this, dimensionality reduction and cluster identification was performed using the functions *RunTFIDF*, FindTopFeatures (min.cutoff =q0), *RunSVD*, *RunUMAP* (dims=2:20, reduction=’lsi’), *FindNeighbours* (dims=2:20, reduction=’lsi’), and *FindClusters* (resolution=0.01). To determine the cell cycle phase of each cell, a predicted gene activity matrix was generated using the function *GeneActivity* which was then used as input to the function *CellCycleScoring* which makes use of previously defined marker genes of S and G2/M phases to generate a cell cycle score for each cell. Cells assigned to each cell cycle phase were extracted to generate pseudobulk coverage files for each condition and phase using custom scripts to be used for visualization.

### Single-cell RNA-seq

Processing of individual single-cell RNA-seq datasets was conducted using the Cell Ranger tool from 10X Genomics using default parameters (https://support.10xgenomics.com/single-cell-gene-expression/software/pipelines/latest/what-is-cell-ranger) with alignment to the human genome (hg19). The outputted gene-by-cell matrix was further processed using the R package *Seurat*^60^. Individual matrices were imported to R and directly merged to generate a combined Seurat object. Following this, dimensionality reduction and cluster identification was performed using the functions *RunPCA*, *FindNeighbors* (dims=1:10), and *FindClusters* (resolution=0.3). Cell cycle phases were assigned using the *CellCycleScoring* function and cells assigned to each cell cycle phase were extracted to generate pseudobulk coverage files for each condition and cell cycle phase using custom scripts to be used for visualization.

*Single-cell Multiome*

Pre-processing of each sample of scMultiome experiments was conducted using the 10X genomics tool Cell Ranger Arc on default parameters and aligning to the human genome (hg19). Count matrices for both accessibility and gene expression were processed using the R packages Seurat and Signac. Merging of individual conditions and analysis was conducted as described above for single cell ATAC and RNA-seq datasets. Joint UMAP visualization of accessibility and gene expression matrices was conducted using the *FindMultiModalNeighbors* function from Signac (as described in the tutorial: https://stuartlab.org/signac/articles/pbmc_multiomic).

### Classification of DSB numbering

DSBs are labelled with specific numbers as defined in^43^. Briefly, all DSBs are labelled based on ordering of all identified AsiSI cut sites (n=1211) by chromosome, which are then subsetting to the 80-best cleaved AsIS1 sites based on signal from BLESS (breaks labeling, enrichment on streptavidin and next-generation sequencing) experiments.

### Data analysis and visualization

#### Average profiles, heatmaps and boxplots

Coverage matrices used for averaged profiles, heatmaps, and boxplots were generated using deeptools (https://deeptools.readthedocs.io/en/latest/) and further visualized using custom R scripts or deeptools. For each boxplot, the center line represents the median, box ends represent respectively the first and third quartiles, and whiskers represent the minimum and maximum values without outliers. Outliers were defined as below the first quartile −(1.5 × interquartile range) and above the third quartile + (1.5 × interquartile range).

#### Single-cell differential accessibility heatmaps for individual DSBs

For heatmaps representing chromatin accessibility tracks of individual cells, the top 300 cells with the highest fragment count at each DSB site were selected. DSB sites were extended to either 10 kb (± 5kb) or 80 kb (± 40kb) windows and the fragment count of each cell over these windows was divided into 100 bins. The log2 ratio of each bin for each cell from 4 h or 24 h OHT treatments was then calculated compared to the average value of each bin in non-treated cells. The resulting matrix was then plotted with *ggplot2* in R.

#### Single-cell differential accessibility heatmaps for all DSBs

For heatmaps representing differential chromatin accessibility for all DSB sites after 4 h or 24 h of DSB induction in each cell, we extracted fragment counts at each DSB for either 10 kb (± 5kb) or 80 kb (± 40kb) windows. To account for overall differences in coverage, cells were divided into 15 sub classes according to their total fragment count and the log2 ratio for each DSB in each cell was determined against the average value for the corresponding fragment count subclass of non-treated cells. The resulting matrix was then plotted using the *pheatmap* package in R.

#### Upstream-to-downstream ratio calculation

Upstream-to-downstream ratio calculations to measure asymmetry of DSB-induced chromatin accessibility changes were performed by first extracting fragment counts of the top 300 cells with the highest fragment count at all HR-prone DSBs (as defined in^43^) in non-treated cells or cells treated with OHT for 24 h. DSB sites were extended to an 80kb (± 40kb) window and the fragment count of each cell over these windows was divided into 100 bins. Following this, cells with a sum fragment count of less than 20 at a given DSB were excluded, and a sum value was taken for each cell at upstream (bins 1:50) or downstream (bins 51:100) at each DSB. The upstream-to-downstream ratio was then determined by calculating the log2 ratio of the upstream and downstream values. The standard deviation of all ratio values of cells at a given DSB was then calculated, converted into a Z-core and plotted as a barplot.

#### Differential module score dotplots

Dot plot visualizations of selected GO, Hallmark, and Pathcards gene sets (see Supplementary Table 9) were generated by extracting the list of genes associated with each gene set that were expressed in over 10% of cells per treatment. The expression of each gene in the gene set was then averaged per treated cell. The mean expression value of the gene set was then averaged across all non-treated cells, and this value was subtracted from the gene set value in each individual treated cell.

#### Identification of cells with High Accessibility at Multiple DSBs (HAM) or Low Accessibility at Multiple DSBs (LAM)

For heatmaps differentiating HAM and LAM cells, the top 300 cells with the highest fragment count at a subset of DSBs prone to undergo homologous recombination (DSB 379, DSB 526, DSB 533, DSB 600, DSB 657, DSB 432, DSB 392, DSB 1159, DSB 343, DSB 765) were selected from defined S & G2/M cell subsets in non-treated cells and cells subjected to 24 h of treatment with OHT. The fragment count for each cell was determined at each DSB site for a 100 kb (± 50 kb) window and the log2 ratio for each cell subjected to 24 h of OHT treatment over the combined average value of non-treated cells at the same DSB was calculated and further converted to a Z-score. The resulting matrix was then subjected to hierarchical clustering and plotted with *ggplot2* in R. HAM and LAM subsets were then manually extracted based on visibly unique clusters from the resulting heatmap.

#### UMAP visualization and trajectory analysis

UMAPs were generated using the *DimPlot* function of the *Seurat* package. Trajectory analysis was performed with *Monocle3*^81^ by first generating a *cell data set* object using the *as.cell_data_set* function with default parameters, followed by the *cluster_cells* function with parameters *reduction_method = “UMAP”, k = 8,* and *random_seed = 2222*. Reversed graph embedding was then performed with *learn_graph* with parameters *use_partition=F, close_loop = T,* with minimal branch length set as 17. Cells were then ordered according to pseudotime with root cells being set as G1 cells from non-treated samples, and plotted with the *plot_cells* function.

## Data and code availability

Data produced in this study are available on ArrayExpress under the accessions E-MTAB-15602 and E-MTAB-15367. All publicly available high throughput sequencing datasets that were used in this study are listed in Supplementary Table 10. Custom scripts and analysis performed in this study have been deposited in the GitHub repository: https://github.com/LegubeDNAREPAIR/scDSB_ATAC.

---

**Supplementary Figure 1:**
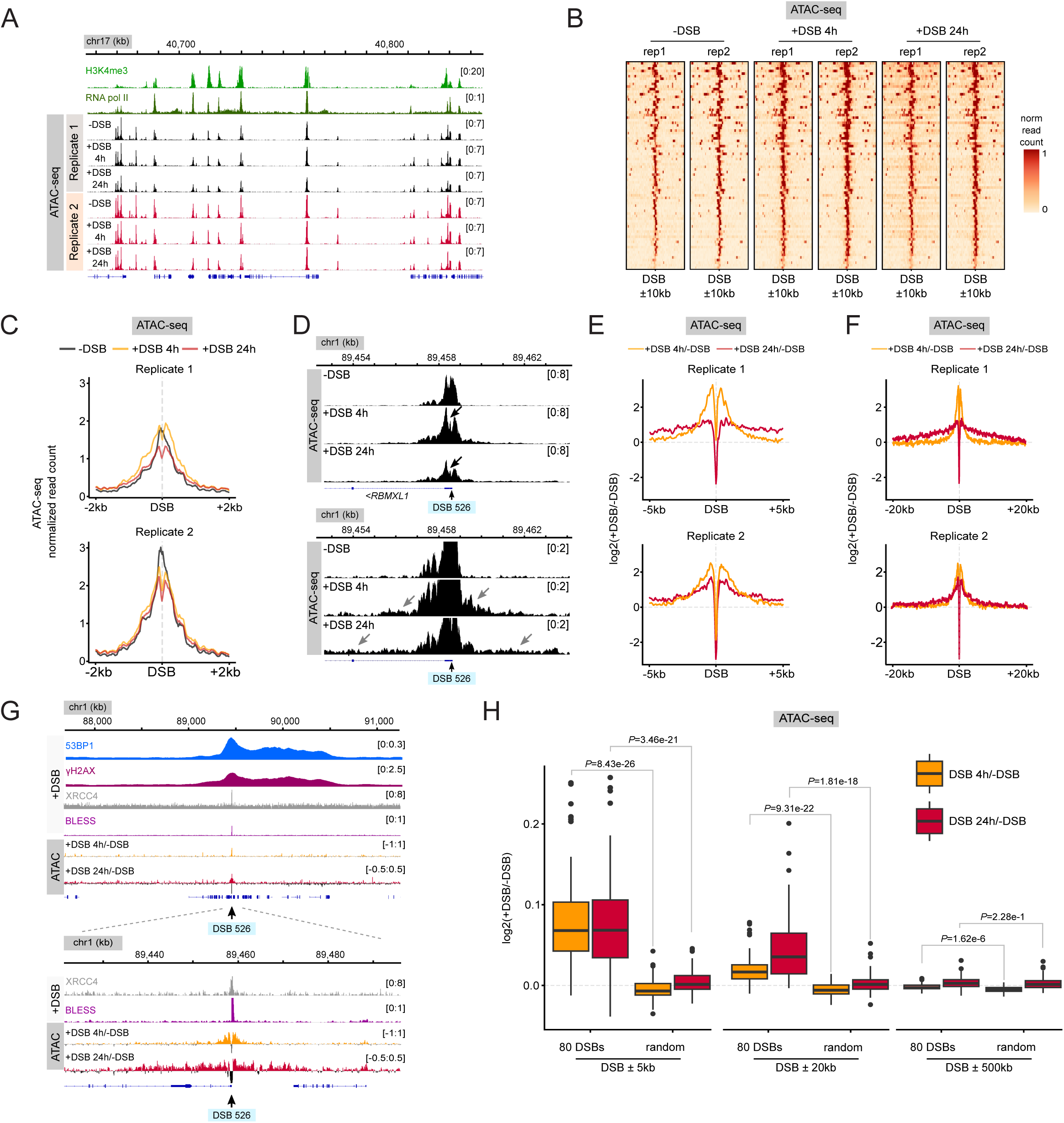
Characterization of chromatin accessibility following DSB induction using ATAC-seq. (A) Genome browser screenshot representing H3K4me3 and RNA pol II ChIP-seq signals in untreated cells (from^82^ and^83^) together with normalized ATAC-seq signals for individual replicates in each condition on a region of chromosome 17. (B) Heatmap representing normalized ATAC-seq signals for 80 DSBs on a 20 kb window (± 10 kb from the DSB) for individual replicates in each condition. (C) Average profiles of normalized ATAC-seq signal at 80 DSBs on a 4 kb window (± 2 kb from the DSB) in untreated cells (dark grey line), after 4 h of DSB induction (yellow line) or 24 h of DSB induction (red line) for replicate 1 (top panel) and replicate 2 (bottom panel). (D) Genome browser screenshot representing normalized ATAC-seq signals for a DSB located on chromosome 17 (DSB 526) at two different magnifications. Black arrows on the upper panel point to the decrease in accessibility proximal to the DSB and grey arrows on the lower panel point to distal increase in accessibility following DSB induction. (E) Average profiles of differential ATAC-seq represented as log2(+DSB/-DSB) after 4 h (yellow line) and 24 h (red line) of DSB induction at 80 DSBs on a 10 kb window (± 5 kb from the DSB) for replicate 1 (top panel) and replicate 2 (bottom panel). (F) Same as (E) on a 40 kb window (± 20 kb from the DSB). (G) Genome browser screenshot representing differential ATAC-seq profiles as log2(+DSB/-DSB) after 4 h or 24 h of DSB induction for a DSB located on chromosome 1 (DSB 526) together with ChIP-seq data for 53BP1 (blue), γH2AX (dark red), XRCC4 (grey) and BLESS (purple) signals (from^43,48^). (H) Boxplot representing the distribution of ATAC-seq signal (Replicate 1) as log2(+DSB/-DSB) at 80 DSBs and 80 control regions after 4 h of DSB induction (yellow) or 24 h of DSB induction (red) on a 10 kb window (± 5 kb from the DSB), 40 kb window (± 20 kb from the DSB), 1 Mb window (± 500 kb from the DSB). Boxes show the interquartile range, center lines represent the median, whiskers extend by 1.5× IQR and dots represent individual outliers. *P* values were obtained from a two-sided Wilcoxon test.

**Supplementary Figure 2:**
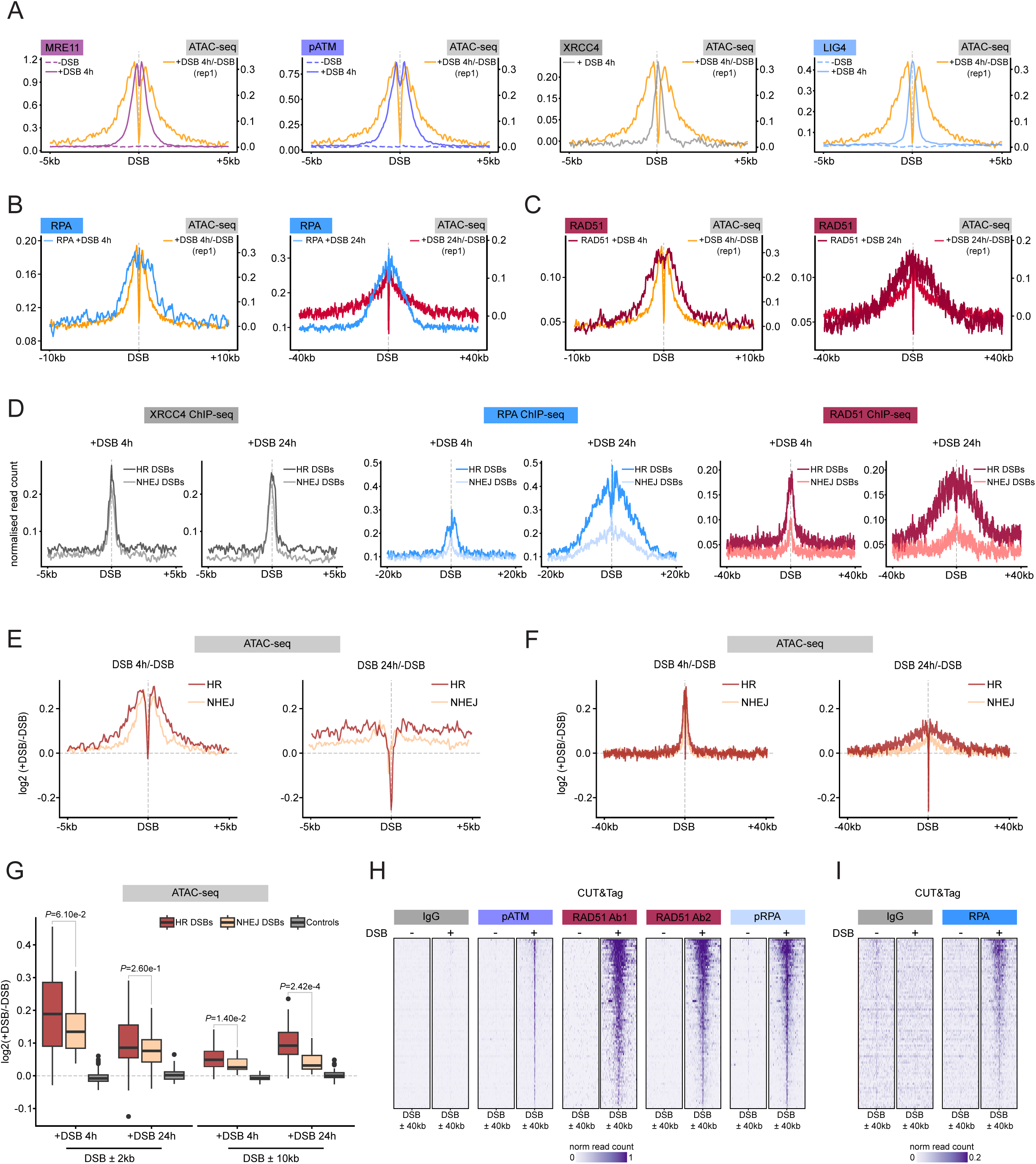
Chromatin accessibility is only increased at a subset of resection-prone DSBs. (A) Average profiles of differential ATAC-seq represented as log2(+DSB/-DSB) after 4 h (yellow lines) of DSB induction, overlayed with normalized ChIP-seq average profiles of MRE11 (dark pink, left panel), pATM (purple, middle left panel, from^47^), XRCC4 (grey, middle right panel, from^48^), and LIG4 (blue, right panel, from^43^) in untreated DIvA cells (dotted line), and after 4 h of DSB induction (solid line) at 80 DSBs on a 10 kb window (± 5 kb from the DSB). (B) Average profiles at 80 DSBs for differential ATAC-seq represented as log2(+DSB/-DSB) after 4 h (left panel, in yellow) and 24 h of DSB induction (right panel, in red), RPA ChIP-seq (blue) after 4 h (left panel) and 24 h (right panel) of DSB induction. Signals are displayed on a 20 kb window (± 10 kb from the DSB) for the 4 h time point (left panel) and 80 kb window (± 40 kb from the DSB) for the 24 h time point (right panel). (C) Average profiles at 80 DSBs for differential ATAC-seq represented as log2(+DSB/-DSB) after 4 h (left panel, in yellow) and 24 h of DSB induction (right panel, in red), RAD51 ChIP-seq (dark red) after 4 h (left panel) and 24 h (right panel) of DSB induction. Signals are displayed on a 20 kb window (± 10 kb from the DSB) for the 4 h time point (left panel) and 80 kb window (± 40 kb from the DSB) for the 24 h time point (right panel). (D) Average profiles for XRCC4, RPA and RAD51 ChIP-seq for 30 HR-prone DSBs and 30 NHEJ-prone DSBs as defined in^43^ after 4 h or 24 h of DSB induction. (E) Average profiles for differential ATAC-seq represented as log2(+DSB/-DSB) after 4 h (left panel) and 24 h of DSB induction (right panel) for 30 HR-prone DSBs (in red) and 30 NHEJ-prone DSBs (in yellow) as defined in^43^. Signals are displayed on a 10 kb window (± 5 kb from the DSB). (F) Same as (E), but displayed on an 80 kb window (± 40 kb from the DSB). (G) Boxplot representing the distribution of ATAC-seq signal as log2(+DSB/-DSB) on a 4 kb window (± 2 kb from the DSB) and 20 kb window (± 10 kb from the DSB) for 30 HR-prone DSBs (in red) and 30 NHEJ-prone DSBs (in yellow) and 80 control regions (in grey) after 4 h and 24 h of DSB induction. Boxes show the interquartile range, center lines represent the median, whiskers extend by 1.5× IQR and dots represent individual outliers. *P* values were obtained from a two-sided Wilcoxon test. (H) Heatmap representation of CUT&Tag signals at 80 DSBs following 24 h of DSB induction (sorted by decreasing levels of differential ATAC-seq signals) on an 80 kb window (± 40 kb from the DSB) for IgG (negative control), ATM S1981P, RAD51 (with 2 different antibodies) and RPA S33P. (I) Same as (H) for IgG and RPA (with a change of scale compared to (E).

**Supplementary Figure 3:**
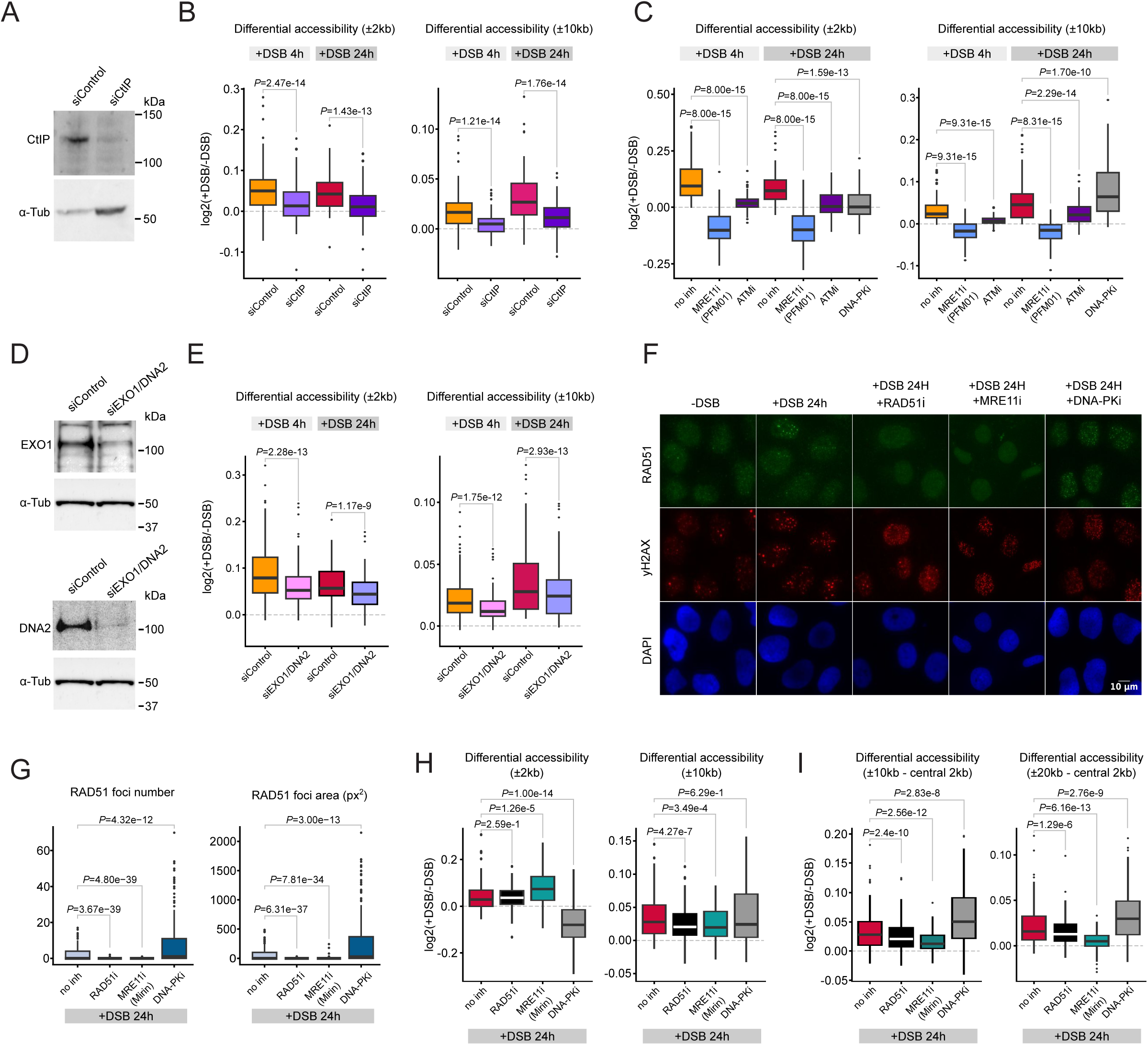
Impact of DNA end resection inhibition on DSB-induced chromatin remodeling. (A) Western blots of CtIP and ɑ-Tubulin after control or CtIP knockdown in U2OS-DIvA cells. (B) Boxplot representing the distribution of differential ATAC-seq profiles as log2(+DSB/-DSB) for 80 DSBs after 4 h or 24 h of DSB induction in siControl- or si-CtIP-treated cells on a 4 kb window (± 2 kb from the DSB, left panel) and 20 kb window (± 10 kb from the DSB, right panel). Boxes show the interquartile range, center lines represent the median, whiskers extend by 1.5× IQR and dots represent individual outliers. *P* values were obtained from a paired two-sided Wilcoxon test. (C) Boxplot representing the distribution of differential ATAC-seq profiles as log2(+DSB/-DSB) for 80 DSBs after 4 h or 24 h of DSB induction and with additional treatment of inhibitors of MRE11 endonuclease activity (PFM01), ATM (KU-55933), and DNA-PKcs (NU7441) on a 4 kb window (± 2 kb from the DSB, left panel) and 20 kb window (± 10 kb from the DSB, right panel). Boxes show the interquartile range, center lines represent the median, whiskers extend by 1.5× IQR and dots represent individual outliers. *P* values were obtained from a paired two-sided Wilcoxon test. (D) Western blots of EXO1 (top panel), DNA2 (bottom panel), and ɑ-Tubulin after control or combined EXO1 and DNA2 knockdown in U2OS-DIvA cells. (E) Boxplot representing the distribution of differential ATAC-seq profiles as log2(+DSB/-DSB) for 80 DSBs after 4 h or 24 h of DSB induction in siControl- or siEXO1+DNA2-treated cells on a 4 kb window (± 2 kb from the DSB, left panel) and 20 kb window (± 10 kb from the DSB, right panel). Boxes show the interquartile range, center lines represent the median, whiskers extend by 1.5× IQR and dots represent individual outliers. *P* values were obtained from a paired two-sided Wilcoxon test. (F) DSBs were induced in DIvA cells for 24 h upon concomitant treatment with the following inhibitors: MRE11i (Mirin, 100 μM), DNA-PKi (NU7441, 2 μM), ATMi (KU-55933, 20 μM) and RAD51i (B02, 25 μM) as indicated. Cells were stained with γH2AX and RAD51 antibodies and representative images are shown. (G) Quantification of RAD51 foci number (left panel) and total foci area (right panel) following treatments described in (I). (H) Boxplot representing the distribution of differential ATAC-seq profiles as log2(+DSB/-DSB) for 80 DSBs after 4 h or 24 h of DSB induction and with additional treatment of inhibitors of RAD51 (B02), MRE11 exonuclease activity (Mirin), and DNA-PKcs (NU7441) on a 4 kb window (± 2 kb from the DSB, left panel) and 20 kb window (± 10 kb from the DSB, right panel). Boxes show the interquartile range, center lines represent the median, whiskers extend by 1.5× IQR and dots represent individual outliers. *P* values were obtained from a paired two-sided Wilcoxon test. (I) Same as (H) on a 10 kb window (± 5 kb from the DSB) with the central 2 kb (± 1 kb from the DSB) removed (left panel), and on a 20 kb window (± 10 kb from the DSB) with the central 2 kb (± 1 kb from the DSB) removes (right panel).

**Supplementary Figure 4:**
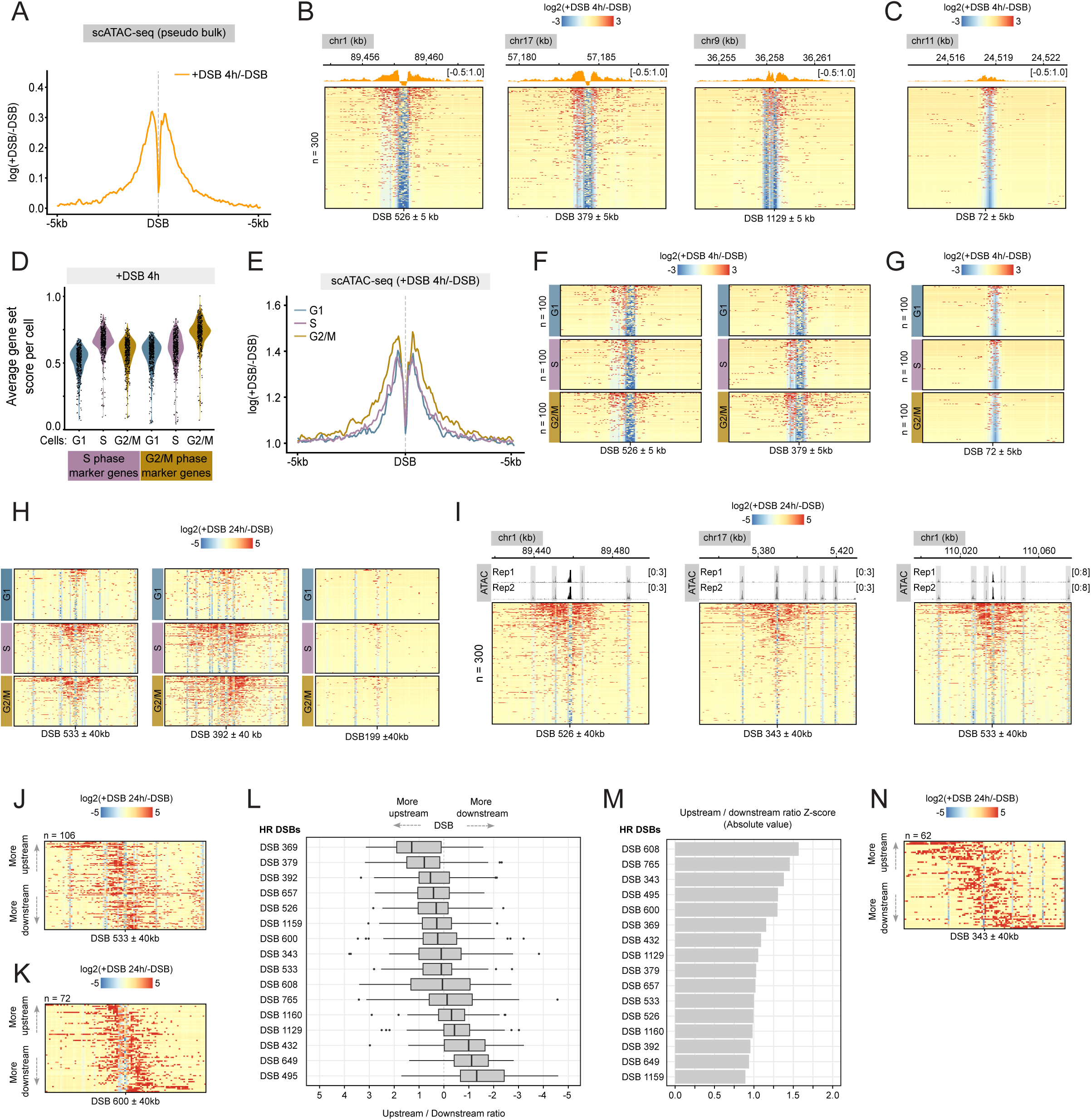
Single-cell DSB-induced chromatin remodeling. (A) Pseudo-bulk average differential ATAC-seq profile as log2(+DSB/-DSB) for 80 DSBs after 4 h of DSB induction on a 10 kb window (± 5 kb from the DSB). (B) Single-cell differential chromatin accessibility profiles as log2(+DSB/-DSB) (n=300 cells) on a 10 kb window (± 5 kb from the DSB) after 4 h of DSB induction for DSBs 526, 379, and 1129. (C) Same as (B), for DSB 72. (D) Violin plots showing average gene set scores per cell for S phase and G2/M phase marker gene sets as defined in Seurat in cells after 4 h of DSB induction. (E) Pseudo-bulk average differential ATAC-seq profile as log2(+DSB/-DSB) for cells in G1 (blue line), S (pink line) or G2/M (gold line) for 80 DSBs after 4 h of DSB induction on a 10 kb window (± 5 kb from the DSB). (F) Single-cell differential chromatin accessibility profiles as log2(+DSB/-DSB) for cells in G1, S or G2/M on a 10 kb window (± 5 kb from the DSB) after 4 h of DSB induction for DSBs 526 and 379. (G) Same as (F), for DSB 72. (H) Single-cell differential chromatin accessibility profiles as log2(+DSB/-DSB) for cells in G1, S or G2/M on a 40 kb window (± 20 kb from the DSB) after 24 h of DSB induction for DSBs 533, 392, and 199 (I) Single-cell differential chromatin accessibility profiles as log2(+DSB/-DSB) (n=300 cells) on a 40 kb window (± 20 kb from the DSB) after 24 h of DSB induction for DSB 526, 343, and 533, with normalized ATAC-seq coverage tracks (black) in untreated cells highlighting accessible genomic regions (in grey). (J) Single-cell differential chromatin accessibility profiles as log2(+DSB/-DSB) for cells after 24 h of DSB induction on an 80 kb window for DSB 533 (n=106 cells). Cells ordered by calculated upstream-to-downstream ratio (see methods). (K) Same as (J), for DSB 600 (n=72 cells). (L) Boxplot representing the upstream-to-downstream ratio (see methods) per cell at individual HR-prone DSBs after 24 h of DSB induction. Boxes show the interquartile range, center lines represent the median, whiskers extend by 1.5× IQR and dots represent individual outliers. (M) Barplot representing the absolute Z-score of the upstream-to-downstream ratio of individual HR-prone DSBs after 24 h of DSB induction. (N) Same as (J), for DSB 343 (n=62 cells).

**Supplementary Figure 5:**
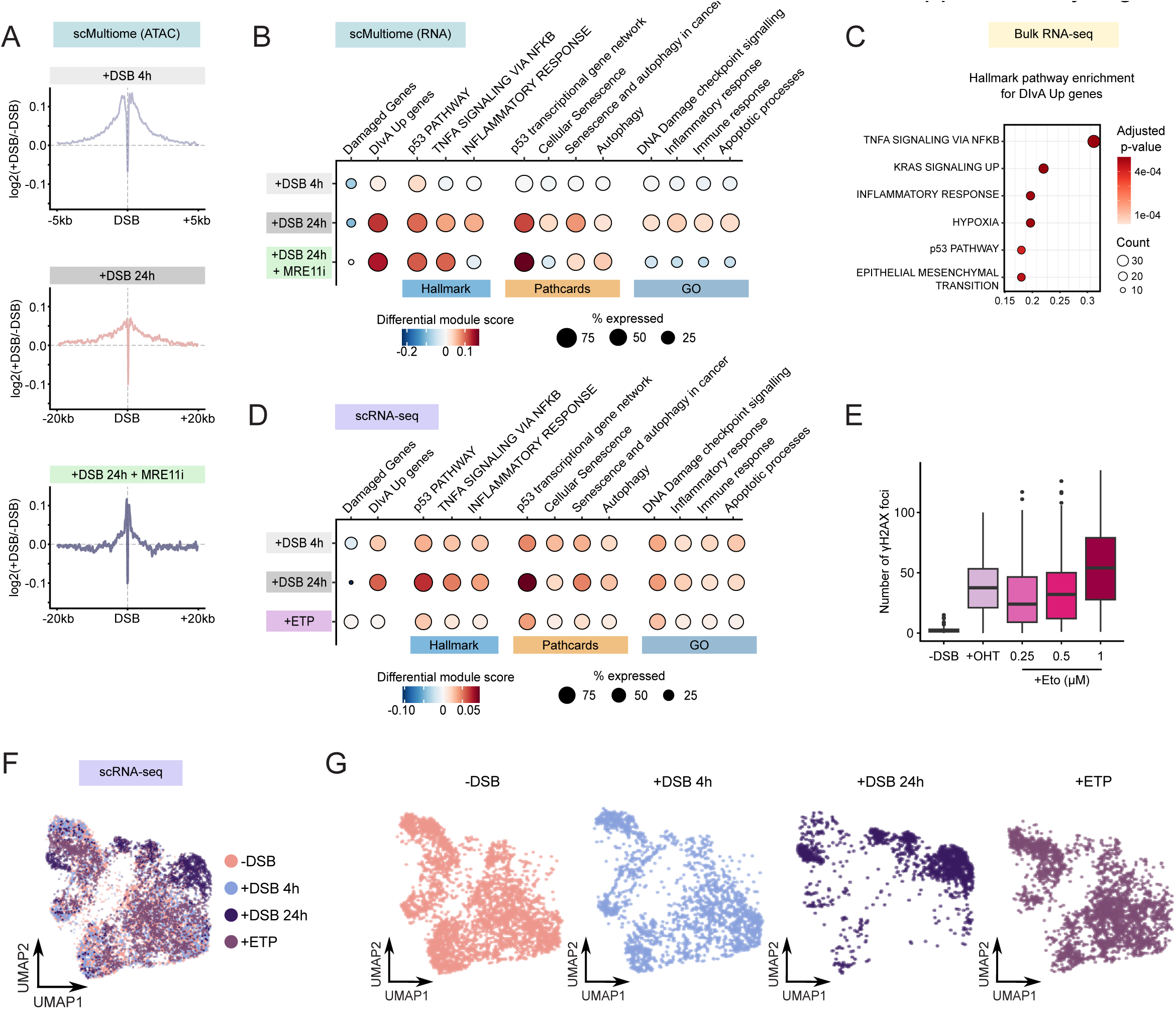
The transcriptional response to genotoxic treatments at single-cell resolution. (A) Pseudo-bulk average differential ATAC-seq profiles from scMultiome as log2(+DSB/-DSB) at 80 DSBs for cells after 4 h (top panel) and 24 h (middle panel) of DSB induction, as well in cells with additional treatment of inhibitors of MRE11 exonuclease activity (Mirin, bottom panel) on a 10 kb window (± 5 kb from the DSB, top panel) and a 40 kb window (± 20 kb from the DSB, middle and bottom panels). (B) Dot plots representing the activity of all assessed Hallmark (H), GO, and Pathcards (PC) gene signatures per treatment from scMultiome. The size of the dot reflects the percentage of cells expressing the genes of interest while the color bar encodes for up- or down-regulation compared with untreated cells. (C) Enrichment of Hallmark gene sets from 286 significantly upregulated genes after 4h of DSB induction identified from bulk RNA-seq experiments^63^. The x-axis shows the gene ratio (number of genes in the term divided by total input genes). The dot size represents the number of genes associated with each term, and the color indicates the adjusted *P* value (Benjamini–Hochberg FDR), with darker colors corresponding to higher significance. (D) Dot plots representing the activity of all assessed Hallmark (H), GO, and Pathcards (PC) gene signatures per treatment from scRNA-seq. The size of the dot reflects the percentage of cells expressing the genes of interest while the color bar encodes for up- or down-regulation compared with untreated cells. (E) Boxplot representing the number of ɣH2AX foci detected by immunofluorescence in U2OS-DIvA cells following the indicated treatment for 4 h. Boxes show the interquartile range, center lines represent the median, whiskers extend by 1.5× IQR and dots represent individual outliers. (F) UMAP projection of 8032 single-cell gene expression profiles from scMultiome. Cells colored according to individual treatments. (G) Same as (F), with individual UMAP projections for each treatment.

**Supplementary Figure 6:**
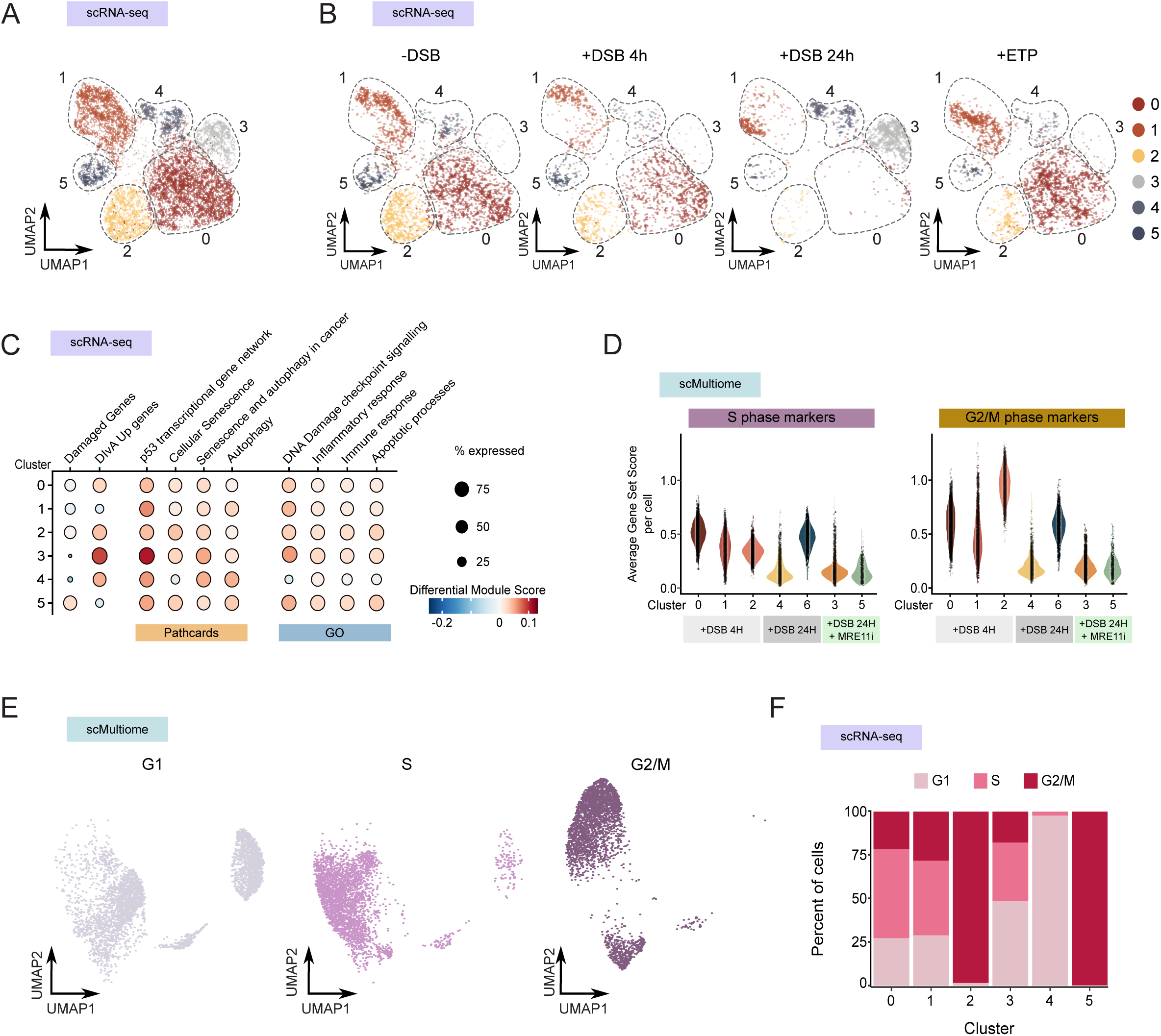
Sustained DNA damage leads to distinct cell fates. (A) UMAP projection of single-cell gene expression profiles from scRNA-seq. Cells colored according to individual clusters. (B) Same as (A), with individual UMAP projections for each treatment. (C) Dot plots representing the activity of assessed GO, and Pathcards (PC) gene signatures per identified cluster relating to a specific treatment from scRNA-seq. The size of the dot reflects the percentage of cells expressing the genes of interest while the color bar encodes for up- or down-regulation compared with untreated cells. (D) Violin plots showing average gene set scores per cell for S phase- (left panel) and G2/M phase-(right panel) marker gene sets as defined in Seurat^60^ in cells from each identified cluster from scMultiome. (E) Individual UMAP projections of cells colored by cell cycle phase from scMultiome. (F) Histogram representing the percentage of cells in each cell cycle phase per identified cluster from scRNA-seq.

**Supplementary Figure 7:**
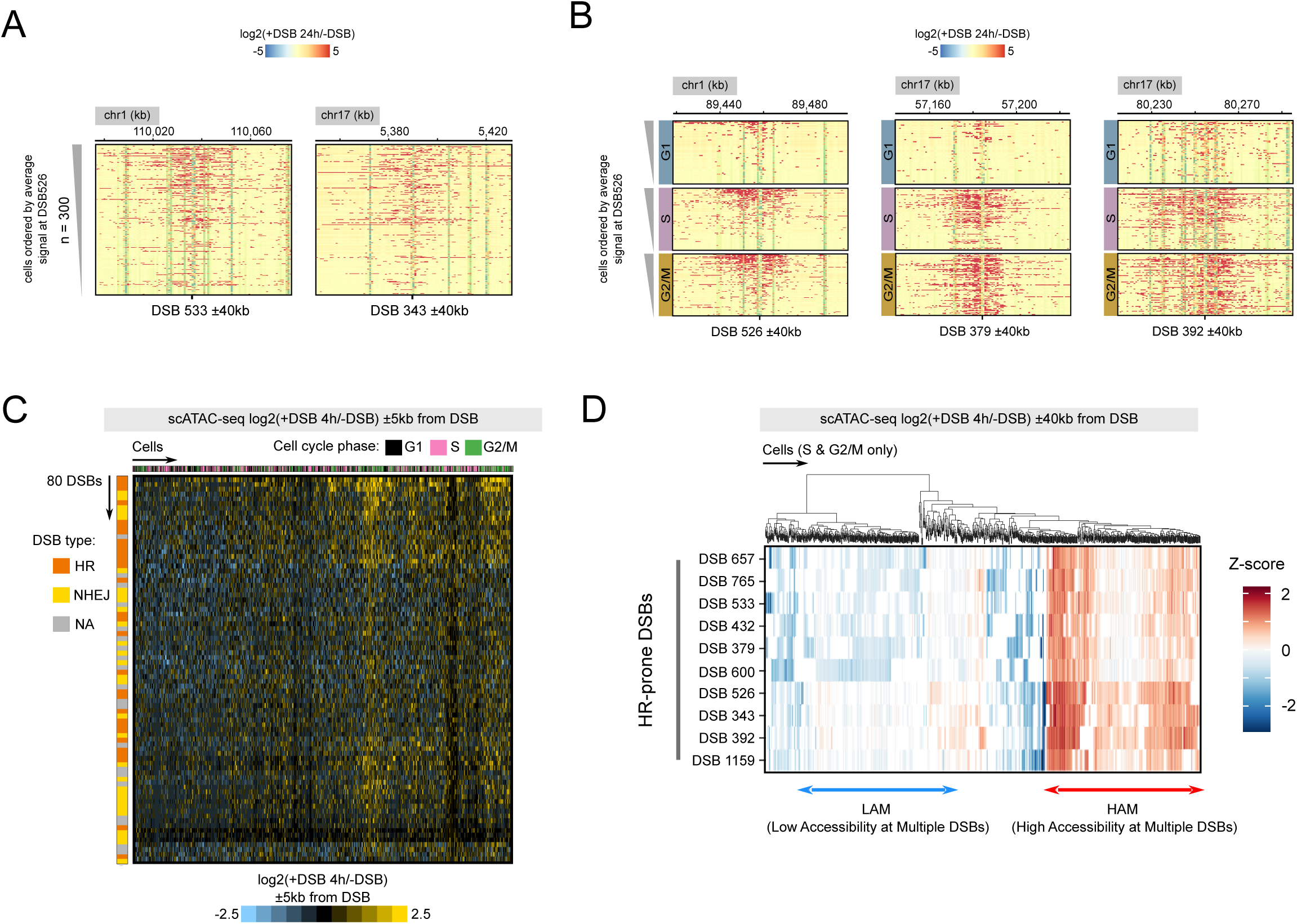
Coordinated DSB-induced chromatin remodeling at specific DSBs after 24 h of DSB induction. (A) Single-cell differential chromatin accessibility profiles as log2(+DSB/-DSB) on an 80 kb window (± 40 kb from the DSB) after 24 h of DSB induction for DSBs 533 and 343. Each heatmap displays 300 cells that are ordered by the mean differential score at DSB 526. (B) Single-cell differential chromatin accessibility profiles as log2(+DSB/-DSB) on an 80 kb window (± 40 kb from the DSB) after 24 h of DSB induction in G1, S and G2/M cells for DSBs 526, 379, and 392. Each heatmap displays 300 cells that are ordered by the mean differential score at DSB 526. (C) Heatmap representing the mean differential log2(+DSB/-DSB) chromatin accessibility scores for cells after 4 h of DSB induction on a 10 kb window (± 5 kb from the DSB). Columns correspond to individual cells colored by cell cycle phase (G1 in black, S in pink, G2/M in green) and ordered by hierarchical clustering. Rows correspond to DSBs colored by preferred repair pathway where applicable (HR in orange, NHEJ in yellow, NA in grey). Color bar refers to the increase (yellow) or decrease (blue) in accessibility at a given DSB in a cell. (D) Hierarchical clustering of cells with a High Accessibility at Multiple DSBs (HAM, in red, n=140) or Low Accessibility at Multiple DSBs (LAM in blue, n=143) in chromatin accessibility among S and G2 cells at a subset of HR-prone DSBs after 4 h of DSB induction. Differential accessibility was computed as log2(+DSB/-DSB) in a single 10 kb bin and transformed into Z-scores (see methods).

**Supplementary Figure 8:**
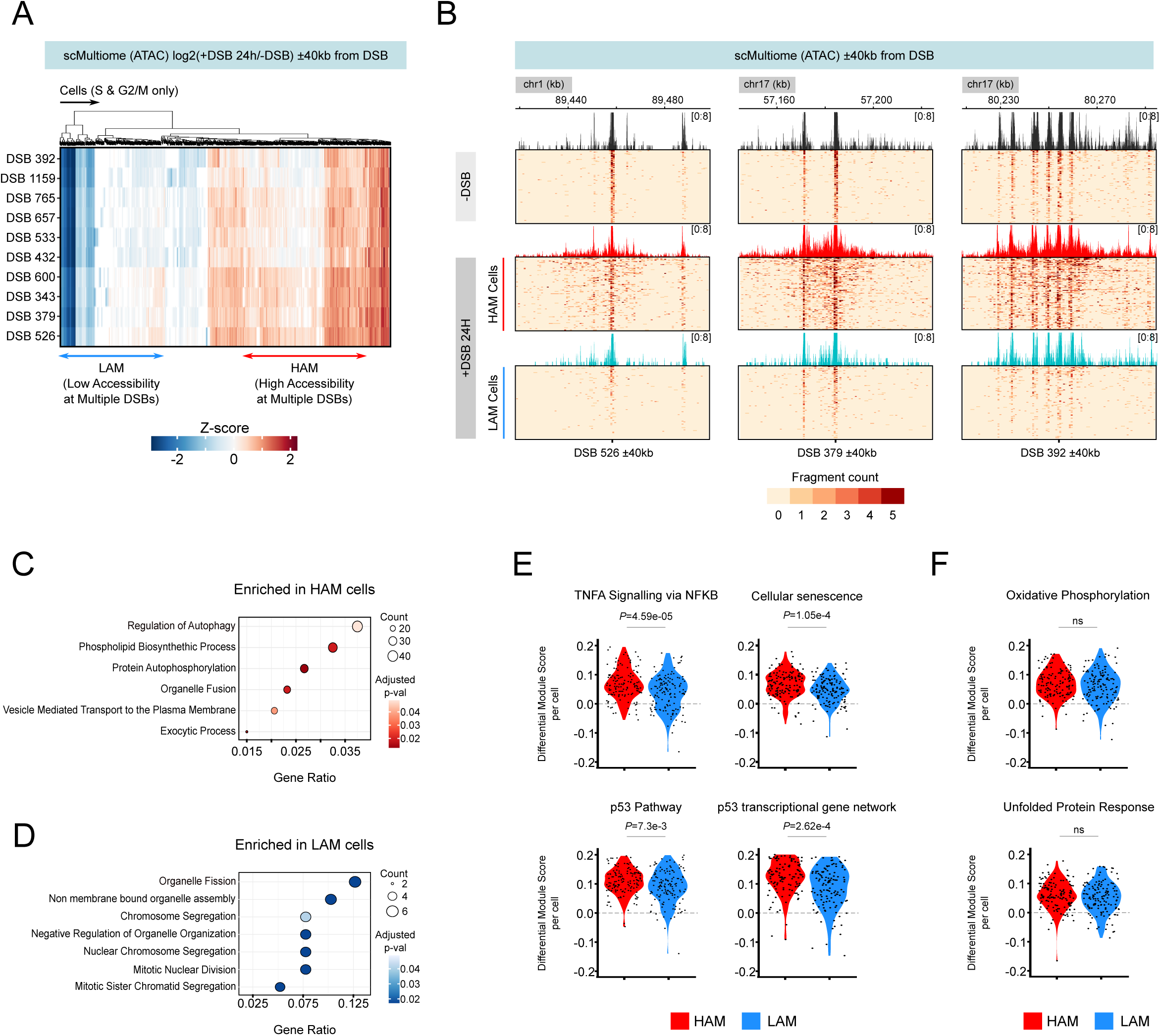
Activation of specific gene expression signatures after DSB induction. (A) Hierarchical clustering of cells from scMultiome with a High Accessibility at Multiple DSBs (HAM, in red, n=140) or Low Accessibility at Multiple DSBs (LAM in blue, n=143) in chromatin accessibility among S and G2 cells at a subset of HR-prone DSBs after 24 h of DSB induction. Differential accessibility was computed as log2(+DSB/-DSB) in a single 10 kb bin and transformed into Z-scores (see methods). (B) Single-cell fragments profiles of non-treated, HAM and LAM sub-populations from scMultiome described in (A) (n=100) on an 80kb window (± 40 kb) at DSBs 526, 379, and 392. Color bar indicates sum of fragments in a given bin. (C) GO (Biological Processes) term enrichment analysis of upregulated genes (n=1611) with a *P* value <0.05 and average log fold change >0.2 in HAM cells compared to LAM cells. The x-axis shows the gene ratio (number of genes in the term divided by total input genes). The dot size represents the number of genes associated with each term, and the color indicates the adjusted p-value (Benjamini–Hochberg FDR), with darker colors corresponding to higher significance. (D) Same as (C), for upregulated genes (n=96) in LAM cells compared to HAM cells. The top 7 GO terms are displayed. (E) Violin plots representing the expression of specific Hallmark and Pathcards gene sets in individual cells in the HAM and LAM sub-populations from S and G2/M cells after 24 h of DSB induction. *P* values were obtained from two-sided Wilcoxon tests. (F) Violin plots representing the expression of specific Hallmark control gene sets in individual cells in the HAM and LAM sub-populations from S and G2/M cells after 24h of DSB induction. *P* values were obtained from two-sided Wilcoxon tests.

