## Supplementary Material for "Single-cell resolution of resection-dependent chromatin accessibility in response to DNA double strand breaks reveals specific gene expression programs"

**Supplementary tables list**

Supplementary Table 1: Cell Ranger quality metrics of single-cell ATAC-seq

Supplementary Table 2: Cell Ranger quality metrics of single-cell Multiome

Supplementary Table 3: Coordinates of damaged genes

Supplementary Table 4: Cell Ranger quality metrics of single-cell RNA-seq

Supplementary Table 5: List of inhibitors used in this study

Supplementary Table 6: List of siRNA used in this study

Supplementary Table 7: List of antibodies used in this study

Supplementary Table 8: Indexing primers for ATAC-seq and CUT&Tag

Supplementary Table 9: List of analyzed gene sets

Supplementary Table 10: List of publicly available datasets used in this study

**Additional Figure 1 :** Single-cell differential chromatin accessibility profiles as log2(+DSB/-DSB) for each of the 80 AsiSI-induced DSBs (n=300 cells).

**Supplementary Table 1 : Cell Ranger quality metrics of single-cell ATAC-seq**

| Condition | NT | OHT 4h | OHT 24h |
|---|---|---|---|
| **Number of cells** | 1525 | 1506 | 1148 |
| **Median fragments per cell** | 26766 | 35759 | 39623 |
| **Total read pairs** | 175825366 | 176631272 | 141759071 |
| **Mean read pairs per cell** | 115295.32 | 117285.04 | 123483.51 |

**Supplementary Table 2 : Cell Ranger quality metrics of single-cell Multiome**

**Multiome-ATAC**

| Condition | NT | OHT 4h | OHT 24h | OHT 24h + Mirin |
|---|---|---|---|---|
| **Number of cells** | 3219 | 4363 | 1713 | 2024 |
| **Median fragments per cell** | 27377 | 17161 | 45938 | 28448 |
| **Total read pairs** | 228846017 | 188460011 | 200875060 | 179514336 |
| **Mean read pairs per cell** | 71092.27 | 43195.05 | 117265.07 | 88692.85 |

**Multiome-RNA**

| Condition | NT | OHT 4h | OHT 24h | OHT 24h + Mirin |
|---|---|---|---|---|
| **Number of cells** | 3217 | 4353 | 1716 | 1999 |
| **Median genes per cell** | 4126 | 3362 | 5760 | 4550 |
| **Total read** | 142541830 | 132950289 | 119311412 | 141205652 |
| **Mean read pairs per cell** | 44308.93 | 30542.22 | 69528.79 | 70638.15 |

**Supplementary Table 3 : Coordinates of damaged genes**

| chromosome | start | end | gene | DSB_gene | gene strand |
|---|---|---|---|---|---|
| chr1 | 9648968 | 9674935 | TMEM201 | SITE495_TMEM201 | + |
| chr1 | 40974438 | 40981710 | EXO5 | SITE516_EXO5 | + |
| chr1 | 89445138 | 89458312 | RBMXL1 | SITE526_RBMXL1 | - |
| chr1 | 110036715 | 110043057 | CYB561D1 | SITE533_CYB561D1 | + |
| chr1 | 204372508 | 204380945 | PPP1R15B | SITE561_PPP1R15B | - |
| chr1 | 223967594 | 224033649 | TP53BP2 | SITE566_TP53BP2 | - |
| chr10 | 3109739 | 3178994 | PFKP | SITE4_PFKP | + |
| chr10 | 94050923 | 94113721 | MARCHF5 | SITE40_MARCHF5 | + |
| chr11 | 24518598 | 25104184 | LUZP2 | SITE72_LUZP2 | + |
| chr11 | 75526254 | 75855276 | UVRAG | SITE101_UVRAG | + |
| chr11 | 85369033 | 85393906 | CREBZF | SITE103_CREBZF | - |
| chr12 | 13153375 | 13157764 | HTR7P1 | SITE133_HTR7P1 | + |
| chr12 | 21950322 | 22094360 | ABCC9 | SITE136_ABCC9 | - |
| chr12 | 121866898 | 122018928 | KDM2B | SITE165_KDM2B | - |
| chr12 | 129556270 | 130388570 | TMEM132D | SITE173_TMEM132D | - |
| chr13 | 114747193 | 114898098 | RASA3 | SITE204_RASA3 | - |
| chr14 | 54941202 | 54955698 | GMFB | SITE219_GMFB | - |
| chr17 | 5389693 | 5394131 | MIS12 | SITE343_MIS12 | + |
| chr17 | 38137059 | 38154212 | PSMD3 | SITE369_PSMD3 | + |
| chr17 | 57060009 | 57184241 | TRIM37 | SITE379_TRIM37 | - |
| chr17 | 61822614 | 61850957 | CCDC47 | SITE380_CCDC47 | - |
| chr17 | 80247921 | 80250690 | LINC01970 | SITE392_LINC01970 | - |
| chr18 | 7567313 | 8406854 | PTPRM | SITE396_PTPRM | + |
| chr18 | 19320752 | 19450914 | MIB1 | SITE404_MIB1 | + |
| chr19 | 2428163 | 2456957 | LMNB2 | SITE432_LMNB2 | - |
| chr19 | 30016324 | 30055458 | VSTM2B | SITE463_VSTM2B | + |
| chr19 | 41903722 | 41930907 | BCKDHA | SITE473_BCKDHA | + |
| chr19 | 42470733 | 42498382 | ATP1A3 | SITE474_ATP1A3 | - |
| chr19 | 45910591 | 45954805 | ERCC1 | SITE477_ERCC1 | - |
| chr2 | 55509454 | 55511608 | PRORSD1P | SITE708_PRORSD1P | + |
| chr2 | 68356936 | 68384659 | DNAAF10 | SITE709_DNAAF10 | - |
| chr2 | 74732169 | 74734822 | PCGF1 | SITE713_PCGF1 | - |
| chr2 | 85822867 | 85824831 | RNF181 | SITE716_RNF181 | + |
| chr2 | 120124552 | 120130119 | DBI | SITE722_DBI | + |
| chr2 | 207938860 | 208031970 | KLF7 | SITE742_KLF7 | - |
| chr20 | 1206687 | 1236520 | RAD21L1 | SITE582_RAD21L1 | + |
| chr20 | 20015011 | 20036690 | CRNKL1 | SITE594_CRNKL1 | - |
| chr20 | 30946133 | 31027122 | ASXL1 | SITE600_ASXL1 | + |
| chr20 | 31995762 | 32031569 | SNTA1 | SITE601_SNTA1 | - |
| chr20 | 37353128 | 37358015 | SLC32A1 | SITE605_SLC32A1 | + |
| chr20 | 42086535 | 42092883 | SRSF6 | SITE608_SRSF6 | + |
| chr21 | 33245332 | 33376377 | HUNK | SITE622_HUNK | + |
| chr21 | 46188494 | 46221735 | UBE2G2 | SITE628_UBE2G2 | - |
| chr22 | 20795805 | 20850082 | KLHL22 | SITE649_KLHL22 | - |
| chr22 | 38864100 | 38879452 | KDELR3 | SITE657_KDELR3 | + |
| chr3 | 52232098 | 52248343 | ALAS1 | SITE765_ALAS1 | + |
| chr3 | 98514784 | 98620539 | DCBLD2 | SITE778_DCBLD2 | - |
| chr3 | 99536705 | 99900576 | CMSS1 | SITE779_CMSS1 | + |

| | | | | | |
|---|---|---|---|---|---|
| chr4 | 83845755 | 83934094 | LIN54 | SITE840_LIN54 | - |
| chr4 | 178351927 | 178363591 | AGA | SITE871_AGA | - |
| chr5 | 68462976 | 68474072 | CCNB1 | SITE899_CCNB1 | + |
| chr5 | 79783918 | 79838382 | FAM151B | SITE903_FAM151B | + |
| chr5 | 142657495 | 142815077 | NR3C1 | SITE919_NR3C1 | - |
| chr6 | 31082576 | 31107869 | PSORS1C1 | SITE953_PSORS1C1 | + |
| chr6 | 37321758 | 37362510 | RNF8 | SITE956_RNF8 | + |
| chr6 | 90341942 | 90348466 | LYRM2 | SITE970_LYRM2 | - |
| chr6 | 135604669 | 135818878 | AHI1 | SITE974_AHI1 | - |
| chr6 | 144606470 | 145174170 | UTRN | SITE975_UTRN | + |
| chr6 | 149887510 | 149912884 | GINM1 | SITE976_GINM1 | + |
| chr7 | 92861678 | 92990435 | VPS50 | SITE1031_VPS50 | + |
| chr7 | 99661459 | 99679346 | ZNF3 | SITE1034_ZNF3 | - |
| chr8 | 66514692 | 66546412 | ARMC1 | SITE1093_ARMC1 | - |
| chr8 | 116420723 | 116681202 | TRPS1 | SITE1107_TRPS1 | - |
| chr8 | 124780678 | 124827692 | FAM91A1 | SITE1110_FAM91A1 | + |
| chr9 | 27937614 | 29213599 | LINGO2 | SITE1123_LINGO2 | - |
| chr9 | 36214437 | 36276975 | GNE | SITE1129_GNE | - |
| chr9 | 127279553 | 127533590 | NR6A1 | SITE1156_NR6A1 | - |
| chr9 | 130683159 | 130693056 | PIP5KL1 | SITE1159_PIP5KL1 | - |
| chr9 | 130882971 | 130890719 | PTGES2 | SITE1160_PTGES2 | - |
| chrX | 1505044 | 1511006 | SLC25A6 | SITE1176_SLC25A6 | - |
| chrX | 45364632 | 45386484 | LINC01204 | SITE1189_LINC01204 | + |
| chrX | 53111548 | 53117722 | TSPYL2 | SITE1191_TSPYL2 | + |
| chrX | 72744110 | 72782921 | MAP2K4P1 | SITE1196_MAP2K4P1 | - |
| chrX | 72782983 | 72906944 | CHIC1 | SITE1196_CHIC1 | + |

**Supplementary Table 4 : Cell Ranger quality metrics of single-cell RNA-seq**

| Condition | NT | OHT 4h | OHT 24h | ETP 4h |
|---|---|---|---|---|
| **Number of cells** | 3061 | 1788 | 1503 | 2399 |
| **Median genes per cell** | 4536 | 5966 | 6390 | 4790 |
| **Total read** | 144107458 | 148985929 | 139116936 | 133153534 |
| **Mean read pairs per cell** | 47079 | 83325 | 92560 | 55504 |

**Supplementary Table 5: List of inhibitors used in this study**

| Name | Target | Reference | Final concentration |
|---|---|---|---|
| KU-55933 | ATM | MedChemExpress (HY-12016) | 20 µM |
| Mirin | MRE11 exonuclease activity | MedChemExpress ( HY-117693) | 100 µM |
| PMF01 | MRE11 endonuclease activity | MedChemExpress (HY-116770) | 100 µM |
| NU7441 | DNA-PKcs | Selleck Chemicals (S2638) | 2 µM |
| B02 | RAD51 | MedChemExpress (HY-101462) | 25 µM |
| Etoposide | TopoII | Sigma (E1383) | 0.5 µM |

**Supplementary Table 6 : List of siRNA used in this study**

| Target | Sequence |
|---|---|
| siCTRL | 5’-CGUACGCGGAAUACUUCGA-3’ |
| siCtIP | 5’-GCUAAAACAGGAACGAAUC-3’ |
| siEXO1 | 5'-CAAGCCUAUUGUCGUAUUUUU-3' |
| siDNA2 | 5'-AAAUAGCCAGUAGUATTCGAU-3' |

**Supplementary Table 7 : List of antibodies used in this study**

| Target | Supplier | Catalog number | Lot Number | Amount | Application |
|---|---|---|---|---|---|
| MRE11 | Genetex | GTX30294 | 822401527 | 2 µl for 200 µg of chromatin | ChIP-seq |
| IgG | Sigma | R9255 | 124M4835V | 2 µg for 200 µg of chromatin | ChIP-seq |
| ATM (S1981P) | Abcam | ab81292 | GR3285525-12 | 0.6 µg for 100000 cells | CUT & Tag |
| RAD51 (ab1) | SantaCruz | sc8349 (H92) | E0616 | 1 µg for 100000 cells | CUT & Tag |
| | | | | 1/200 | IF |
| RAD51 (ab2) | Abcam | ab176458 | GR152117-51 | 1 µg for 100000 cells | CUT & Tag |
| RPA (S33P) | Bethyl | A300-246A | 8 | 1 µg for 100000 cells | CUT & Tag |
| RPA | Abcam | ab10359 | GR3227595-24 | 1 µg for 100000 cells | CUT & Tag |
| IgG | Epicypher | 13-0042 | 20335004-04 | 1 µg for 100000 cells | CUT & Tag |
| CtIP | Active Motif | 61141 | 20217003 | 1/500 | Western-Blot |
| EXO1 | Invitrogen | MA5-12262 | TH2610566 | 1/500 | Western-Blot |
| DNA2 | Abcam | ab96488 | GR260085-1 | 1/500 | Western-Blot |
| γH2AX | Millipore | 05-636, clone JBW301 | 3718325 | 1/1000 | IF |
| alpha-Tubulin | Sigma-Aldrich | T6199 | 00001-43673 | 1/10000 | Western-Blot |

**Supplementary Table 8: Indexing primers for ATAC-seq and CUT&Tag**

| Name | Sequence | Index |
|---|---|---|
| Ad1.1_TAGATCGC | AATGATACGGCGACCACCGAGATCTACACTAGATCGCTCGTCGGCAGCGTCAGATGTGTAT | i5 |
| Ad1.2_CTCTCTAT | AATGATACGGCGACCACCGAGATCTACACCTCTCTATTCGTCGGCAGCGTCAGATGTGTAT | i5 |
| Ad1.3_TATCCTCT | AATGATACGGCGACCACCGAGATCTACACTATCCTCTTCGTCGGCAGCGTCAGATGTGTAT | i5 |
| Ad1.4_AGAGTAGA | AATGATACGGCGACCACCGAGATCTACACAGAGTAGATCGTCGGCAGCGTCAGATGTGTAT | i5 |
| Ad1.5_GTAAGGAG | AATGATACGGCGACCACCGAGATCTACACGTAAGGAGTCGTCGGCAGCGTCAGATGTGTAT | i5 |
| Ad1.6_ACTGCATA | AATGATACGGCGACCACCGAGATCTACACACTGCATATCGTCGGCAGCGTCAGATGTGTAT | i5 |
| Ad1.7_AAGGAGTA | AATGATACGGCGACCACCGAGATCTACACAAGGAGTATCGTCGGCAGCGTCAGATGTGTAT | i5 |
| Ad1.8_CTAAGCCT | AATGATACGGCGACCACCGAGATCTACACCTAAGCCTTCGTCGGCAGCGTCAGATGTGTAT | i5 |
| Ad1.9_TGGAAATC | AATGATACGGCGACCACCGAGATCTACACTGGAAATCTCGTCGGCAGCGTCAGATGTGTAT | i5 |
| Ad1.10_AACATGAT | AATGATACGGCGACCACCGAGATCTACACAACATGATTCGTCGGCAGCGTCAGATGTGTAT | i5 |
| Ad2.1_TAAGGCGA | CAAGCAGAAGACGGCATACGAGATTCGCCTTAGTCTCGTGGGCTCGGAGATGTG | i7 |
| Ad2.2_CGTACTAG | CAAGCAGAAGACGGCATACGAGATCTAGTACGGTCTCGTGGGCTCGGAGATGTG | i7 |
| Ad2.3_AGGCAGAA | CAAGCAGAAGACGGCATACGAGATTTCTGCCTGTCTCGTGGGCTCGGAGATGTG | i7 |
| Ad2.4_TCCTGAGC | CAAGCAGAAGACGGCATACGAGATGCTCAGGAGTCTCGTGGGCTCGGAGATGTG | i7 |
| Ad2.5_GGACTCCT | CAAGCAGAAGACGGCATACGAGATAGGAGTCCGTCTCGTGGGCTCGGAGATGTG | i7 |
| Ad2.6_TAGGCATG | CAAGCAGAAGACGGCATACGAGATCATGCCTAGTCTCGTGGGCTCGGAGATGTG | i7 |
| Ad2.7_CTCTCTAC | CAAGCAGAAGACGGCATACGAGATGTAGAGAGGTCTCGTGGGCTCGGAGATGTG | i7 |
| Ad2.8_CAGAGAGG | CAAGCAGAAGACGGCATACGAGATCCTCTCTGGTCTCGTGGGCTCGGAGATGTG | i7 |
| Ad2.9_GCTACGCT | CAAGCAGAAGACGGCATACGAGATAGCGTAGCGTCTCGTGGGCTCGGAGATGTG | i7 |
| Ad2.10_CGAGGCTG | CAAGCAGAAGACGGCATACGAGATCAGCCTCGGTCTCGTGGGCTCGGAGATGTG | i7 |
| Ad2.11_AAGAGGCA | CAAGCAGAAGACGGCATACGAGATTGCCTCTTGTCTCGTGGGCTCGGAGATGTG | i7 |
| Ad2.12_GTAGAGGA | CAAGCAGAAGACGGCATACGAGATTCCTCTACGTCTCGTGGGCTCGGAGATGTG | i7 |
| Ad2.13_TGGATCTG | CAAGCAGAAGACGGCATACGAGATCAGATCCAGTCTCGTGGGCTCGGAGATGTG | i7 |
| Ad2.14_CCGTTTGT | CAAGCAGAAGACGGCATACGAGATACAAACGGGTCTCGTGGGCTCGGAGATGTG | i7 |

**Supplementary Table 9: List of analysis gene sets**

| HALLMARK Gene Sets | | |
|---|---|---|
| **Gene Set** | **Number of Genes** | **Genes** |
| HYPOXIA | 196 | CXCR7, ADM, ADORA2B, AK4, AKAP12, ALDOA, ALDOB, ALDOC, AMPD3, ANGPTL4, ANKZF1, ANXA2, ATF3, ATP7A, B3GALT6, B4GALNT2, BCAN, BCL2, BGN, BHLHE40, BNIP3L, BRS3, BTG1, CA12, CASP6, CAV1, PTRF, PRKCDBP, CYR61, CTGF, WISP2, CCNG2, CDKN1A, CDKN1B, CDKN1C, CHST2, CHST3, CITED2, COL5A1, CP, CSRP2, CXCR4, DCN, DDIT3, DDIT4, DPYSL4, DTNA, DUSP1, EDN2, EFNA1, EFNA3, EGFR, ENO1, ENO2, ENO3, ERO1L, ERRFI1, ETS1, EXT1, F3, FAM162A, FBP1, FOS, FOSL2, FOXO3, GAA, GALK1, GAPDH, GAPDHS, GBE1, GCK, GCNT2, GLRX, GPC1, GPC3, GPC4, GPI, GRHPR, GYS1, HAS1, HDLBP, HEXA, HK1, HK2, HMOX1, HOXB9, HS3ST1, HSPA5, IDS, IER3, IGFBP1, IGFBP3, IL6, ILVBL, INHA, IRS2, ISG20, JMJD6, JUN, KDM3A, KIF5A, KLF6, KLHL24, LALBA, LARGE, LDHA, LDHC, LOX, LXN, MAFF, MAP3K1, MIF, MT1E, MT2A, MXI1, MYH9, NAGK, NCAN, NDRG1, NDST1, NDST2, NEDD4L, NFIL3, CCRN4L, P4HA1, P4HA2, PAM, PCK1, PDGFB, PDK1, PDK3, PFKFB3, PFKL, PGAM2, PGF, PGK1, PGM1, PGM2, PHKG1, PIM1, PKLR, PKP1, PLAC8, PLAUR, PLIN2, PNRC1, PPARGC1A, PPFIA4, PPP1R15A, PPP1R3C, PRDX5, PRKCA, PYGM, RBPJ, RORA, RRAGD, S100A4, SAP30, SCARB1, SDC2, SDC3, SDC4, SELENBP1, SERPINE1, SIAH2, SLC25A1, SLC2A1, SLC2A3, SLC2A5, SLC37A4, SLC6A6, SRPX, STBD1, STC1, STC2, SULT2B1, TES, TGFB3, TGFBI, TGM2, TIPARP, TKTL1, TMEM45A, TNFAIP3, TPBG, TPD52, TPI1, TPST2, UGP2, VEGFA, VHL, VLDLR, WSB1, XPNPEP1, ZFP36, ZNF292 |
| EPITHELIAL_MESENCHYMAL_TRANSITION | 200 | ABI3BP, ACTA2, ADAM12, ANPEP, APLP1, AREG, BASP1, BDNF, BGN, BMP1, CADM1, CALD1, CALU, CAP2, CAPG, CYR61, CTGF, CD44, CD59, CDH11, CDH2, CDH6, COL11A1, COL12A1, COL16A1, COL1A1, COL1A2, COL3A1, COL4A1, COL4A2, COL5A1, COL5A2, COL5A3, COL6A2, COL6A3, COL7A1, COL8A2, GLT25D1, COMP, COPA, CRLF1, CTHRC1, CXCL1, CXCL12, CXCL6, IL8, DAB2, DCN, DKK1, DPYSL3, DST, ECM1, ECM2, EDIL3, EFEMP2, ELN, EMP3, ENO2, FAP, FAS, FBLN1, FBLN2, FBLN5, FBN1, FBN2, FERMT2, FGF2, FLNA, FMOD, FN1, FOXC2, FSTL1, FSTL3, FUCA1, FZD8, GADD45A, GADD45B, GAS1, GEM, GJA1, GLIPR1, GPC1, GPX7, GREM1, HTRA1, ID2, IGFBP2, IGFBP3, IGFBP4, IL15, IL32, IL6, INHBA, ITGA2, ITGA5, ITGAV, ITGB1, ITGB3, ITGB5, JUN, LAMA1, LAMA2, LAMA3, LAMC1, LAMC2, LGALS1, LOX, LOXL1, LOXL2, LRP1, LRRC15, LUM, MAGEE1, MATN2, MATN3, MCM7, MEST, MFAP5, MGP, MMP1, MMP14, MMP2, MMP3, MSX1, MXRA5, MYL9, MYLK, NID2, NNMT, NOTCH2, NT5E, NTM, OXTR, LEPRE1, PCOLCE, PCOLCE2, PDGFRB, PDLIM4, PFN2, PLAUR, PLOD1, PLOD2, PLOD3, PMEPA1, PMP22, POSTN, PPIB, PRRX1, PRSS2, PTHLH, PTX3, PVR, QSOX1, RGS4, RHOB, SAT1, SCG2, SDC1, SDC4, SERPINE1, SERPINE2, SERPINH1, SFRP1, SFRP4, SGCB, SGCD, SGCG, SLC6A8, SLIT2, SLIT3, SNAI2, SNTB1, SPARC, SPOCK1, SPP1, TAGLN, TFPI2, TGFB1, TGFBI, TGFBR3, TGM2, THBS1, THBS2, THY1, TIMP1, TIMP3, TNC, TNFAIP3, TNFRSF11B, TNFRSF12A, TPM1, TPM2, TPM4, VCAM1, VCAN, VEGFA, VEGFC, VIM, WIPF1, WNT5A |
| HALLMARK_INFLAMMATORY_RESPONSE | 199 | ABCA1, ABI1, ACVR1B, ACVR2A, EMR1, ADM, ADORA2B, ADRM1, AHR, APLNR, AQP9, ATP2A2, ATP2B1, ATP2C1, AXL, BDKRB1, BEST1, BST2, BTG2, C3AR1, C5AR1, CALCRL, CCL17, CCL2, CCL20, CCL22, CCL24, CCL5, CCL7, CCR7, CCRL2, CD14, CD40, CD48, CD55, CD69, CD70, CD82, CDKN1A, CHST2, CLEC5A, CMKLR1, CSF1, CSF3, CSF3R, CX3CL1, CXCL10, CXCL11, CXCL6, IL8, CXCL9, CXCR6, CYBB, EBI3, EDN1, EIF2AK2, EMP3, EREG, F3, FFAR2, FPR1, FZD5, GABBR1, GCH1, GNA15, GNAI3, GP1BA, GPC3, GPR132, GPR183, HAS2, HBEGF, HIF1A, HPN, HRH1, ICAM1, ICAM4, ICOSLG, IFITM1, IFNAR1, IFNGR2, IL10, IL10RA, IL12B, IL15, IL15RA, IL18, IL18R1, IL18RAP, IL1A, IL1B, IL1R1, IL2RB, IL4R, IL6, IL7R, INHBA, IRAK2, IRF1, IRF7, ITGA5, ITGB3, ITGB8, KCNA3, KCNJ2, KCNMB2, KIF1B, KLF6, LAMP3, LCK, LCP2, LDLR, LIF, LPAR1, LTA, LY6E, LYN, MARCO, MEFV, MEP1A, MET, MMP14, MSR1, MXD1, MYC, NAMPT, NDP, NFKB1, NFKBIA, NLRP3, NMI, NMUR1, NOD2, NPFFR2, OLR1, OPRK1, OSM, OSMR, P2RX4, P2RX7, P2RY2, PCDH7, PDE4B, PDPN, PIK3R5, PLAUR, PROK2, PSEN1, PTAFR, PTGER2, PTGER4, PTGIR, PTPRE, PVR, RAF1, RASGRP1, RELA, RGS1, RGS16, RHOG, RIPK2, RNF144B, ROS1, RTP4, SCARF1, SCN1B, SELE, SELS, SELL, SEMA4D, SERPINE1, SGMS2, SLAMF1, SLC11A2, SLC1A2, SLC28A2, SLC31A1, SLC31A2, SLC4A4, SLC7A1, SLC7A2, SPHK1, SRI, STAB1, TACR1, TACR3, TAPBP, TIMP1, TLR1, TLR2, TLR3, TNFAIP6, TNFRSF1B, TNFRSF9, TNFSF10, TNFSF15, TNFSF9, TPBG, VIP |
| APOPTOSIS | 161 | ADD1, AIFM3, ANKH, ANXA1, APP, ATF3, AVPR1A, BAX, BCAP31, BCL10, BCL2L1, BCL2L10, BCL2L11, BCL2L2, BGN, BID, BIK, BIRC3, BMF, BMP2, BNIP3L, BRCA1, BTG2, BTG3, CASP1, CASP2, CASP3, CASP4, CASP6, CASP7, CASP8, CASP9, CAV1, CCNA1, CCND1, CCND2, |

| | | |
|---|---|---|
| | | CD14, CD2, CD38, CD44, CD69, CDC25B, CDK2, CDKN1A, CDKN1B, CFLAR, CLU, CREBBP, CTH, CTNNB1, CYLD, DAP, DAP3, DCN, DDIT3, DFFA, DIABLO, DNAJA1, DNAJC3, DNM1L, DPYD, EBP, EGR3, EMP1, ENO2, ERBB2, ERBB3, EREG, ETF1, F2, F2R, FAS, FASLG, FDXR, FEZ1, GADD45A, GADD45B, GCH1, GNA15, GPX1, GPX3, GPX4, GSN, GSR, GSTM1, GUCY2D, H1F0, HGF, HMGB2, HMOX1, HSPB1, IER3, IFITM3, IFNB1, IFNGR1, IGF2R, IGFBP6, IL18, IL1A, IL1B, IL6, IRF1, ISG20, JUN, KRT18, LEF1, LGALS3, LMNA, LUM, MADD, MCL1, MGMT, MMP2, NEDD9, NEFH, PAK1, PDCD4, PDGFRB, PEA15, PLAT, PLCB2, LPPR4, PMAIP1, PPP2R5B, PPP3R1, PPT1, PRF1, PSEN1, PSEN2, PTK2, RARA, RELA, RETSAT, RHOB, RHOT2, RNASEL, ROCK1, SAT1, SATB1, SC5DL, SLC20A1, SMAD7, SOD1, SOD2, SPTAN1, SQSTM1, TAP1, TGFB2, TGFBR3, TIMP1, TIMP2, TIMP3, TNF, TNFRSF12A, TNFSF10, TOP2A, TSPO, TXNIP, VDAC2, WEE1, XIAP |
| HALLMARK_P53_PATHWAY | 199 | ABAT, ABCC5, ABHD4, ACVR1B, ADA, AEN, AK1, ALOX15B, ANKRA2, APAF1, APP, ATF3, BAIAP2, BAK1, BAX, BLCAP, BMP2, BTG1, BTG2, CASP1, CCND2, CCND3, CCNG1, CCNK, CCP110, CD81, CD82, CDH13, CDK5R1, CDKN1A, CDKN2A, CDKN2AIP, CDKN2B, CEBPA, CGRRF1, CLCA2, ADCK3, CSRNP2, CTSD, CTSF, CYFIP2, DCXR, DDB2, DDIT3, DDIT4, DEF6, DGKA, DNTTIP2, DRAM1, EI24, IKBKAP, EPHA2, EPHX1, EPS8L2, ERCC5, F2R, FAM162A, FAS, FBXW7, FDXR, FGF13, FOS, FOXO3, FUCA1, GADD45A, GLS2, GM2A, GPX2, HIST1H1C, HIST3H2A, H2AFJ, HBEGF, HDAC3, HEXIM1, HINT1, HMOX1, HRAS, HSPA4L, IER3, IER5, IFI30, IL1A, INHBB, IP6K2, LRMP, IRAK1, ISCU, ITGB4, JAG2, JUN, KIF13B, KLF4, KLK8, KRT17, LDHB, LIF, MAPKAPK3, MDM2, MKNK2, MXD1, MXD4, NDRG1, NHLH2, NINJ1, NOL8, NOTCH1, NUDT15, NUPR1, OSGIN1, PCNA, PDGFA, PERP, PHLDA3, PIDD, PITPNC1, PLK2, PLK3, PLXNB2, PMM1, POLH, POM121, PPM1D, PPP1R15A, PRKAB1, PRMT2, PROCR, PTPN14, PTPRE, PVT1, RAB40C, GNB2L1, RAD51C, RAD9A, RALGDS, RAP2B, RB1, RCHY1, RETSAT, RGS16, RHBDF2, RNF19B, RPL18, RPL36, RPS12, RPS27L, RRAD, RRP8, RXRA, S100A10, S100A4, SAT1, SDC1, SEC61A1, SERPINB5, SERTAD3, SESN1, SFN, SLC19A2, SLC35D1, SLC3A2, SLC7A11, SOCS1, SP1, SPHK1, ST14, STEAP3, STOM, TAP1, TAX1BP3, TCHH, TCN2, TGFA, TGFB1, TM4SF1, TM7SF3, TNFSF9, TNNI1, TOB1, TP53, TP63, TPD52L1, TPRKB, TRAF4, TRAFD1, TRIAP1, TRIB3, TSC22D1, TXNIP, UPP1, VAMP8, VDR, VWA5A, WRAP73, WWP1, XPC, ZBTB16, ZFP36L1, ZMAT3, ZNF365 |
| TNFA_SIGNALING_VIA_NFKB | 200 | ABCA1, CXCR7, AREG, ATF3, ATP2B1, B4GALT1, B4GALT5, BCL2A1, BCL3, BCL6, BHLHE40, BIRC2, BIRC3, BMP2, BTG1, BTG2, BTG3, CCL2, CCL20, CCL4, CCL5, CYR61, CCND1, CCNL1, CCRL2, CD44, CD69, CD80, CD83, CDKN1A, CEBPB, CEBPD, CFLAR, CLCF1, CSF1, CSF2, CXCL1, CXCL10, CXCL11, CXCL2, CXCL3, CXCL6, DENND5A, DNAJB4, DRAM1, DUSP1, DUSP2, DUSP4, DUSP5, EDN1, EFNA1, EGR1, EGR2, EGR3, EHD1, EIF1, ETS2, F2RL1, F3, FJX1, FOS, FOSB, FOSL1, FOSL2, FUT4, G0S2, GADD45A, GADD45B, GCH1, GEM, GFPT2, GPR183, HBEGF, HES1, ICAM1, ICOSLG, ID2, IER2, IER3, IER5, IFIH1, IFIT2, IFNGR2, IL12B, IL15RA, IL18, IL1A, IL1B, IL23A, IL6, IL6ST, IL7R, INHBA, IRF1, IRS2, JAG1, JUN, JUNB, KDM6B, KLF10, KLF2, KLF4, KLF6, KLF9, KYNU, LAMB3, LDLR, LIF, LITAF, MAFF, MAP2K3, MAP3K8, MARCKS, MCL1, MSC, MXD1, MYC, NAMPT, NFAT5, NFE2L2, NFIL3, NFKB1, NFKB2, NFKBIA, NFKBIE, NINJ1, NR4A1, NR4A2, NR4A3, OLR1, PANX1, PDE4B, PDLIM5, PER1, PFKFB3, PHLDA1, PHLDA2, PLAU, PLAUR, PLEK, PLK2, PPAP2B, PMEPA1, PNRC1, PPP1R15A, PTGER4, PTGS2, PTPRE, PTX3, RCAN1, REL, RELA, RELB, RHOB, DDX58, RIPK2, RNF19B, SAT1, SDC4, SERPINB2, SERPINB8, SERPINE1, SGK1, SIK1, SLC16A6, SLC2A3, SLC2A6, SMAD3, SNN, SOCS3, SOD2, SPHK1, SPSB1, SQSTM1, STAT5A, TANK, TAP1, TGIF1, TIPARP, TLR2, TNC, TNF, TNFAIP2, TNFAIP3, TNFAIP6, TNFAIP8, TNFRSF9, TNFSF9, TNIP1, TNIP2, TRAF1, TRIB1, TRIP10, TSC22D1, TUBB2A, VEGFA, YRDC, ZBTB10, ZC3H12A, ZFP36 |
| HALLMARK_KRAS_SIGNALING_UP | 199 | ABCB1, ACE, ADAM17, ADAM8, ADAMDEC1, GPR124, ELTD1, AKAP12, AKT2, ALDH1A2, ALDH1A3, AMMECR1, ANGPTL4, ANKH, ANO1, ANXA10, APOD, ARG1, ATG10, AVL9, BIRC3, BMP2, BPGM, BTBD3, BTC, C3AR1, CA2, CAB39L, CBL, CBR4, CBX8, CCL20, CCND2, FAM190B, CD37, CDADC1, CFB, CFH, CFHR2, CIDEA, CLEC4A, CMKLR1, CPE, CROT, CSF2, CSF2RA, CTSS, CXCL10, CXCR4, DNMBP, DOCK2, DUSP6, EMP1, ENG, EPB41L3, EPHB2, EREG, ERO1L, ETS1, ETV1, ETV4, ETV5, EVI5, F13A1, F2RL1, FBXO4, FCER1G, FGF9, FLT4, FUCA1, G0S2, GABRA3, GADD45G, GALNT3, GFPT2, GLRX, GNG11, GPNMB, GPRC5B, GUCY1A3, GYPC, HIST1H2BB, HBEGF, HDAC9, HKDC1, HOXD11, HSD11B1, ID2, IGF2, IGFBP3, IKZF1, IL10RA, IL1B, IL1RL2, IL2RG, IL33, IL7R, INHBA, IRF8, ITGA2, ITGB2, ITGBL1, JUP, KCNN4, KIF5C, KLF4, LAPTM5, LAT2, LCP1, LIF, LY96, MAFB, MALL, MAP3K1, MAP4K1, MAP7, MMD, MMP10, MMP11, MMP9, MPZL2, MTMR10, MYCN, NAP1L2, NGF, NIN, NR0B2, NR1H4, NRP1, PCP4, PCSK1N, PDCD1LG2, PECAM1, PEG3, PIGR, PLAT, PLAU, PLAUR, PLEK2, PLVAP, PPBP, PPP1R15A, PRDM1, SLMO2, PRKG2, PRRX1, PSMB8, PTBP2, PTCD2, PTGS2, PTPRR, RABGAP1L, RBM4, RBP4, RELN, RETN, RGS16, SATB1, SCG3, SCG5, SCN1B, SDCCAG8, SEMA3B, SERPINA3, SLPI, SNAP25, SNAP91, SOX9, SPARCL1, SPON1, SPP1, SPRY2, ST6GAL1, STRN, TFPI, TLR8, TMEM100, |

| | | |
|---|---|---|
| | | TMEM158, TMEM176A, TMEM176B, TNFAIP3, TNFRSF1B, TNNT2, TOR1AIP2, TPH1, TRAF1, TRIB1, TRIB2, TSPAN1, TSPAN13, TSPAN7, USH1C, USP12, VWA5A, WDR33, WNT7A, YRDC, ZNF277, ZNF639 |

### Pathcard Gene Sets

| Gene Set | Number of Genes | Genes |
|---|---|---|
| Autophagy | 62 | CXCR7, ADM, ADORA2B, AK4, AKAP12, ALDOA, ALDOB, ALDOC, AMPD3, ANGPTL4, ANKZF1, ANXA2, ATF3, ATP7A, B3GALT6, B4GALNT2, BCAN, BCL2, BGN, BHLHE40, BNIP3L, BRS3, BTG1, CA12, CASP6, CAV1, PTRF, PRKCDBP, CYR61, CTGF, WISP2, CCNG2, CDKN1A, CDKN1B, CDKN1C, CHST2, CHST3, CITED2, COL5A1, CP, CSRP2, CXCR4, DCN, DDIT3, DDIT4, DPYSL4, DTNA, DUSP1, EDN2, EFNA1, EFNA3, EGFR, ENO1, ENO2, ENO3, ERO1L, ERRFI1, ETS1, EXT1, F3, FAM162A, FBP1, FOS, FOSL2, FOXO3, GAA, GALK1, GAPDH, GAPDHS, GBE1, GCK, GCNT2, GLRX, GPC1, GPC3, GPC4, GPI, GRHPR, GYS1, HAS1, HDLBP, HEXA, HK1, HK2, HMOX1, HOXB9, HS3ST1, HSPA5, IDS, IER3, IGFBP1, IGFBP3, IL6, ILVBL, INHA, IRS2, ISG20, JMJD6, JUN, KDM3A, KIF5A, KLF6, KLHL24, LALBA, LARGE, LDHA, LDHC, LOX, LXN, MAFF, MAP3K1, MIF, MT1E, MT2A, MXI1, MYH9, NAGK, NCAN, NDRG1, NDST1, NDST2, NEDD4L, NFIL3, CCRN4L, P4HA1, P4HA2, PAM, PCK1, PDGFB, PDK1, PDK3, PFKFB3, PFKL, PGAM2, PGF, PGK1, PGM1, PGM2, PHKG1, PIM1, PKLR, PKP1, PLAC8, PLAUR, PLIN2, PNRC1, PPARGC1A, PPFIA4, PPP1R15A, PPP1R3C, PRDX5, PRKCA, PYGM, RBPJ, RORA, RRAGD, S100A4, SAP30, SCARB1, SDC2, SDC3, SDC4, SELENBP1, SERPINE1, SIAH2, SLC25A1, SLC2A1, SLC2A3, SLC2A5, SLC37A4, SLC6A6, SRPX, STBD1, STC1, STC2, SULT2B1, TES, TGFB3, TGFBI, TGM2, TIPARP, TKTL1, TMEM45A, TNFAIP3, TPBG, TPD52, TPI1, TPST2, UGP2, VEGFA, VHL, VLDLR, WSB1, XPNPEP1, ZFP36, ZNF292 |
| Senescence and autophagy in cancer | 104 | ABI3BP, ACTA2, ADAM12, ANPEP, APLP1, AREG, BASP1, BDNF, BGN, BMP1, CADM1, CALD1, CALU, CAP2, CAPG, CYR61, CTGF, CD44, CD59, CDH11, CDH2, CDH6, COL11A1, COL12A1, COL16A1, COL1A1, COL1A2, COL3A1, COL4A1, COL4A2, COL5A1, COL5A2, COL5A3, COL6A2, COL6A3, COL7A1, COL8A2, GLT25D1, COMP, COPA, CRLF1, CTHRC1, CXCL1, CXCL12, CXCL6, IL8, DAB2, DCN, DKK1, DPYSL3, DST, ECM1, ECM2, EDIL3, EFEMP2, ELN, EMP3, ENO2, FAP, FAS, FBLN1, FBLN2, FBLN5, FBN1, FBN2, FERMT2, FGF2, FLNA, FMOD, FN1, FOXC2, FSTL1, FSTL3, FUCA1, FZD8, GADD45A, GADD45B, GAS1, GEM, GJA1, GLIPR1, GPC1, GPX7, GREM1, HTRA1, ID2, IGFBP2, IGFBP3, IGFBP4, IL15, IL32, IL6, INHBA, ITGA2, ITGA5, ITGAV, ITGB1, ITGB3, ITGB5, JUN, LAMA1, LAMA2, LAMA3, LAMC1, LAMC2, LGALS1, LOX, LOXL1, LOXL2, LRP1, LRRC15, LUM, MAGEE1, MATN2, MATN3, MCM7, MEST, MFAP5, MGP, MMP1, MMP14, MMP2, MMP3, MSX1, MXRA5, MYL9, MYLK, NID2, NNMT, NOTCH2, NT5E, NTM, OXTR, LEPRE1, PCOLCE, PCOLCE2, PDGFRB, PDLIM4, PFN2, PLAUR, PLOD1, PLOD2, PLOD3, PMEPA1, PMP22, POSTN, PPIB, PRRX1, PRSS2, PTHLH, PTX3, PVR, QSOX1, RGS4, RHOB, SAT1, SCG2, SDC1, SDC4, SERPINE1, SERPINE2, SERPINH1, SFRP1, SFRP4, SGCB, SGCD, SGCG, SLC6A8, SLIT2, SLIT3, SNAI2, SNTB1, SPARC, SPOCK1, SPP1, TAGLN, TFPI2, TGFB1, TGFBI, TGFBR3, TGM2, THBS1, THBS2, THY1, TIMP1, TIMP3, TNC, TNFAIP3, TNFRSF11B, TNFRSF12A, TPM1, TPM2, TPM4, VCAM1, VCAN, VEGFA, VEGFC, VIM, WIPF1, WNT5A |
| Cellular senescence | 196 | CDK4, CDK6, LOC102724334, CDKN2A, CDKN2B, CDKN2C, CDKN2D, H4C16, H3C14, H2BC26, FOS, H2BC1, H2AC8, H2AC7, H2AX, H2BC5, H2BC3, H3-3A, H3-3B, H3C15, JUN, H2AB1, H4C15, H2AJ, MAPK1, MAPK3, RPS27A, H3C13, H2AC19, UBA52, UBB, UBC, H4C9, H2AC14, H2AC6, H2AC4, H2AC18, H2AC20, H2BC8, H2BC13, H2BC15, H2BC14, H2BC7, H2BC6, H2BC9, H2BC10, H2BC4, H2BC17, H2BC21, H3C1, H3C4, H3C3, H3C6, H3C11, H3C8, H3C12, H3C10, H3C2, H4C1, H4C4, H4C6, H4C12, H4C11, H4C3, H4C8, H4C2, H4C5, H4C13, H4C14, H2BC12, H3C7, H2BC11, H2AZ2, CDK2, CDKN1A, CDKN1B, ANAPC10, CEBPB, EHMT2, UBE2C, ANAPC16, CDC26, ANAPC15, VENTX, UBE2S, ANAPC2, ANAPC4, IGFBP7, IL1A, IL6, CXCL8, NFKB1, FZR1, ANAPC5, ANAPC7, ANAPC11, MAPK7, RELA, RPS6KA1, RPS6KA2, RPS6KA3, ANAPC1, STAT3, UBE2D1, UBE2E1, EHMT1, CDC23, CDC16, CCNA2, CCNA1, CDC27, COMMD3-BMI1, MAPK14, E2F1, E2F2, E2F3, PHC1, PHC2, AGO3, AGO4, EZH2, SCMH1, TNIK, TNRC6B, KDM6B, CBX6, SUZ12, AGO1, TNRC6A, IFNB1, MDM2, MDM4, MAP3K5, MOV10, MINK1, MAPK8, MAPK11, MAPK9, MAPK10, MAP2K3, MAP2K6, MAP2K7, CBX8, TNRC6C, RBBP4, RBBP7, RING1, RNF2, MAP2K4, BMI1, TFDP1, TFDP2, TP53, TXN, MAPKAPK3, PHC3, CBX2, CBX4, MAPKAPK5, EED, MAPKAPK2, MAP4K4, RAD50, KAT5, ERF, ETS1, ETS2, CABIN1, ASF1A, POT1, TINF2, UBN1, H1-0, H1-2, H1-3, H1-4, H1-5, H1-1, HMGA1, ID1, LMNB1, MRE11, NBN, ATM, TERF2IP, EP400, RB1, ACD, SP1, TERF1, TERF2, HIRA, HMGA2, H3-4, CCNE1, CCNE2 |
| p53 transcriptional gene network | 96 | LOC105376156, CDK2, CDKN1A, IRF9, SIVA1, SMR3B, CPT1C, ADORA2B, E2F7, DDB2, GADD45A, ERCC5, SLC7A11, MTOR, FUCA1, BBC3, GLS2, |

| | | |
|---|---|---|
| | | SESN1, SFN, GPX1, APAF1, ICAM1, FAS, FASLG, IRF5, LIF, MIR145, MIR200C, MIR34A, MIR34B, MIR34C, MGMT, MLH1, MSH2, NCF2, NOTCH1, RRM2B, SERPINE1, PCNA, PRKAG2, POLK, SERPINB5, PRKAG3, PMAIP1, PML, POLH, DDIT4, DRAM1, PIDD1, PRKAA1, PRKAA2, PRKAB1, PRKAB2, PRKAG1, RPRM, TIGAR, PTEN, ADGRB1, RPTOR, BAX, SAT1, CCL2, CX3CL1, TP53AIP1, PERP, MLST8, ZMAT3, DEPTOR, SLC2A1, AURKA, THBS1, TNF, TSC2, XPC, XRCC5, BTG2, NANOG, ULBP2, ULBP1, SESN2, ULK1, ACAD11, AKT1S1, ALDH4A1, TNFRSF10D, TNFRSF10B, CCNE1, CCNG1, TP53INP1, TP53I3, TRAF4, ISG15, ULK2, CDC25C, SCO2 |

### Gene Ontology (GO) Gene Sets

| Gene Set | Number of Genes | Genes |
|---|---|---|
| DNA damage checkpoint signalling GO:0000077 | 125 | ATM, ATR, BARD1, CCND1, BLM, BRCA1, BRCA2, CASP2, CCNG1, CDK1, CDC5L, CDK2, CDKN1A, CDKN1B, CHEK1, FOXN3, PLK3, ATF2, CRY1, MAPK14, E2F1, ERCC6, FANCD2, GML, H2AX, HUS1, MDM2, FOXO4, MRE11, MSH2, MUC1, NBN, PLK1, PML, PPP1R10, PRKDC, PTPN11, RAD1, RAD9A, RAD17, RAD51, RPA2, RPL26, TP53, TP53BP1, XPC, CUL4A, CDC14B, MBTPS1, CRADD, IER3, TAOK2, CLOCK, BABAM2, MDC1, TTI1, TELO2, FEM1B, PLK2, TOPBP1, CHEK2, TREX1, UFL1, INTS7, SYF2, HINFP, GIGYF2, FBXO6, FBXO4, PRPF19, BABAM1, RPA4, DONSON, NOP53, RPS27L, MRNIP, WAC, FZR1, TAOK3, MBTPS2, TRIAP1, GTSE1, DTL, UIMC1, MAP3K20, ETAA1, GNB1L, TIPIN, RFWD3, PIDD1, TRIM39, TAOK1, USP28, CCAR2, RINT1, CLSPN, STK33, BRCC3, RNASEH2B, FBXO31, NEK11, WDR76, TTI2, MUS81, CEP63, CDK5RAP3, PARP9, RHNO1, BRIP1, ATRIP, ABRAXAS1, DOT1L, BRSK1, TICRR, ZNF830, PRAP1, HUS1B, E2F7, RAD9B, EME1, DTX3L, SDE2, EME2, TIPRL, EIF2AK4 |
| Inflammatory response GO:0006954 | 892 | A2M, SERPINA3, ABCF1, ACP5, ACVR1, ADA, ADAM8, ADCY1, ADCY7, ADCY8, ADM, ADORA1, ADORA2A, ADORA2B, ADORA3, PARP4, AGER, AGT, AGTR1, AGTR2, AHR, AHSG, AIF1, AKT1, ABCD1, ABCD2, ALOX5, ALOX5AP, ALOX15, ANXA1, AOAH, APCS, BIRC2, BIRC3, XIAP, APOA1, APOD, APOE, APP, ARNT, ASS1, ATM, AXL, AZU1, BCL6, BCR, BDKRB1, BDKRB2, BMP2, BMP6, BMPR1B, BPGM, BST1, BTK, C1QA, C3, C3AR1, C4A, C4B, C5, C5AR1, TMEM258, DAGLA, CALCA, CAMK4, CASP1, CASP4, CASP5, CD5L, CD6, CD14, CD28, CD36, CD40, CD40LG, CD44, CD47, CD68, CD81, ADGRE5, CDH5, CDO1, CEBPA, CEBPB, CTSC, CHI3L1, CHUK, CLU, CMA1, CCR1, CCR3, CCR4, CCR5, CCR6, CCR7, ACKR2, CMKLR1, LTB4R, CNR1, CNR2, CNTF, COL6A1, CRH, CRHBP, CRP, MAPK14, CSF1, CSF1R, CSNK1A1, CX3CR1, CYBA, CYBB, CYLD, CYP19A1, DDT, DDX3X, DHX9, DNASE1, DNASE1L3, DPEP1, ECM1, S1PR3, EDNRB, EGFR, EPHA2, CELA1, ELANE, ELF3, ELF4, EPHB2, EPO, ESR1, ETS1, EXT1, EXTL3, EZH2, F2, F2R, F2RL1, F3, F8, F12, FABP4, FANCA, FANCD2, FASN, FCER1A, FCGR1A, FCGR1BP, FCGR2A, FCGR2B, FCGR3A, FCGR3B, FGR, FOXF1, FN1, FOLR2, FOS, FOSL2, FPR1, FPR2, FPR3, MTOR, FUT4, FUT7, ACKR1, FYN, GATA3, GBA1, GBP2, GGT1, GGT3P, B4GALT1, GGT5, GHSR, GNAT2, GPR4, XCR1, CXCR3, GPR17, GPER1, GPR31, GPR32, GPR32P1, GPR33, FFAR3, FFAR2, GPS2, GPX1, GRN, CXCL1, CXCL2, CXCL3, GSTP1, HCK, HGF, HIF1A, HK1, HLA-DRB1, HLA-E, HMGB1, HMGB2, HMOX1, NR4A1, HP, HRH1, HSPA8, HSPG2, NDST1, TNC, HYAL1, IFI16, IFI35, IFNA2, IFNG, IFNGR1, IFNGR2, IGF1, IGHE, IGHG1, RBPJ, IKBKB, IL1A, IL1B, IL1R1, IL1RAP, IL1RN, IL2, IL2RA, IL4, IL4R, IL5, IL5RA, IL6, IL6R, IL6ST, CXCL8, IL9, CXCR2, IL10, IL10RA, IL10RB, IL12B, IL13, IL15, IL16, IL17A, IL18, IDO1, CXCL10, INS, IRAK2, IRF3, IRF5, ISL1, ITGAL, ITGAM, ITGB1, ITGB2, ITGB6, ITIH4, JAK2, KARS1, KCNJ8, KIT, KLKB1, KNG1, KRT1, KRT16, LBP, LDLR, LEP, LGALS1, LGALS2, LGALS9, LIPA, LPL, LRP1, LTA, CD180, LY75, LYN, LYZ, SMAD1, SMAD3, MAPT, MAS1, MBL2, MDK, MEFV, MEP1B, MGST2, MIF, CXCL9, MMP3, MMP8, MMP9, CD200, ABCC1, MVK, MYD88, NAGLU, NAIP, NDP, NFATC3, NFATC4, NFE2L2, NFKB1, NFKB2, NFKBIA, NFKBIB, NFX1, NINJ1, NKG7, NOS2, NOTCH1, NOTCH2, CCN3, NPY5R, NT5E, OLR1, ORM1, ORM2, OSM, P2RX1, P2RX7, FURIN, SERPINE1, REG3A, PDE2A, ENPP3, PER1, PF4, PF4V1, SERPINA1, PIK3CD, PIK3CG, PLA2G2A, PLCG1, PLCG2, SERPINF2, PLP1, PLSCR1, POLB, PPARA, PPARD, PPARG, PPBP, PPP2CA, PRCP, PRKCD, PRKD1, PRKCQ, PRKCZ, MAPK7, MAPK8, MAPK9, MAPK13, MAP2K3, EIF2AK2, PROC, PSEN1, PSG9, PSMA1, PSMA6, PSMB4, PTAFR, PTGDR, PTGER1, PTGER2, PTGER3, PTGER4, PTGFR, PTGIR, PTGIS, PTGS2, PTN, PTPN2, PTPN6, PTPRC, PTX3, RAC1, RARRES2, RB1, REL, RELA, RELB, RORA, RPS19, S100A8, S100A9, S100A12, SAA1, SAA2, SAA4, SCN9A, SCNN1B, CCL1, CCL2, CCL3, CCL3L1, CCL4, CCL5, CCL7, CCL8, CCL11, CCL13, CCL14, CCL15, CCL16, CCL17, CCL18, CCL19, CCL20, CCL21, CCL22, CCL23, CCL24, CCL25, CXCL6, CXCL11, CXCL5, XCL1, CX3CL1, SELE, SELP, SLAMF1, SLC11A1, SLC18A2, SMO, SIGLEC1, SNCA, SOD1, SPN, SRC, STAT3, STAT5B, VAMP7, XCL2, SYK, TAC1, TACR1, MAP3K7, TBXA2R, PRDX2, TEK, TFR2, TFRC, TGFB1, THBS1, TIMP1, TLR1, TLR2, TLR3, TLR4, TLR5, TMSB4X, TNF, TNFAIP3, TNFAIP6, TNFRSF1A, |

| | | |
|---|---|---|
| | | TNFRSF1B, TNFSF4, TNFRSF4, TYRO3, TYROBP, SCGB1A1, UMOD, VCAM1, TRPV1, WNT5A, YES1, ZFP36, ZP3, IL1R2, CXCR4, SCG2, PLA2G7, CSRP3, FOSL1, FXR1, GPR68, BAP1, HYAL3, PLA2G10, CUL3, ATRN, TPST1, SEMA7A, IKBKG, CST7, CHST1, AP3B1, TNFSF11, PLA2G4C, AOC3, VAMP8, HYAL2, TRADD, SNX4, RIPK1, RIPK2, SNAP23, TNFRSF11A, IL18RAP, IL1RL2, IL18R1, CCN4, IER3, VNN1, SPHK1, RPS6KA4, PGLYRP1, SOCS3, CCRL2, PSTPIP1, FCGR2C, NMI, IL1RL1, OSMR, LARGE1, PNMA1, RPS6KA5, AIMP1, MFHAS1, MAPKAPK2, KLF4, CD163, ADIPOQ, PLAA, CHST2, AIM2, LY86, THEMIS2, PTGES, NR1D1, CLOCK, SOCS5, HDAC9, HDAC4, SPATA2, TRIM14, ZEB2, TRIL, PJA2, AREL1, NR1H4, NR1D2, HDAC5, NR1H3, TSPAN2, RASGRP1, LPCAT3, CHST4, KLRG1, CD96, TCIRG1, TNIP1, TLR6, CCL26, NOD1, AGR2, CXCL13, TXNIP, CELF1, CXCR6, TRAF3IP2, PLK2, CYSLTR1, HNRNPA0, TMED2, LILRB4, LIAS, TRIM31, DUSP10, USP18, TREX1, SCN11A, PARK7, TUSC2, MGLL, NLRP1, CARD8, SBNO2, SIRT2, TAB2, KDM6B, STAB1, SYT11, UFL1, BRD4, CD2AP, MKRN2, KPNA6, LY96, PLD3, SHPK, IL17RA, APOL2, RHBDD3, PRDX5, FBXL2, LETMD1, DHRS7B, APPL1, ABHD12, PTPN22, STAP1, PLA2G2D, NUPR1, IL36RN, NOX1, LAT, FOXP1, EIF2AK1, CHIA, IL36B, IL37, IL36A, IL17C, IL17B, C5AR2, PDCD4, SMPDL3B, RABGEF1, STK39, NKIRAS2, GIT1, DROSHA, PYCARD, TBK1, BLNK, POMT2, PLA2G2E, ADGRE2, PLA2G3, NOX4, IL20, IL22, F11R, FOXP3, ZNF580, TLR7, TLR8, VPS54, IL23A, GPRC5B, GHRL, IL17D, IL20RB, SETD4, TLR9, TREM2, TOLLIP, UGT1A1, PXK, ODAM, APPL2, ACER3, CAMK2N1, FEM1A, POMGNT1, NLRP2, KDM4D, VPS35, SELENOS, PLGRKT, ASH1L, PBK, IL36G, MMP26, CYP26B1, SUCNR1, SLAMF8, DHX33, LXN, CTNNBIP1, CAMK1D, SCYL3, PBXIP1, SCYL1, TMIGD3, MAVS, MARK4, HAMP, WFDC1, NLRC4, IL22RA1, IL21, CARD18, ACE2, SIGIRR, HRH4, TRPV4, PROK2, GPSM3, SLC39A8, NOD2, CARD9, NFKBIZ, MMP25, CLEC7A, IL25, FNDC4, LRRC19, WNK4, CHID1, CUEDC2, FKRP, TNIP2, BRCC3, TNFAIP8L2, MCPH1, RHBDF2, NLRX1, GSDMD, SETD6, TNIP3, ATAT1, ZC3H12A, TRIM45, SCUBE1, AKNA, NDFIP1, CPTP, APOL3, ZBP1, TRIM11, ADAMTS12, TLR10, SHARPIN, JAM3, ITCH, WDR83, TTBK1, AFAP1L2, IL1F10, SPINK7, TRIM55, LOXL3, CREB3L3, NFKBID, IL17RC, HAVCR2, ZDHHC12, TSLP, SIGLEC10, TSPAN18, OTULIN, MACIR, DUOXA1, IL33, NLRP12, MYLK3, IL17F, NLRP3, TIRAP, CARD16, PGLYRP2, C1QTNF3, GBP5, IL22RA2, PIK3AP1, LRRK2, PLD4, NLRP13, NLRP8, NLRP5, REG3G, CD200R1, IL17RE, IL31RA, NEK7, SIRPA, LACC1, ARMH4, LRFN5, C2CD4A, IL34, NLRP4, TICAM1, IL23R, NFAM1, PYDC2, DAB2IP, NLRP6, ANO6, NLRC3, NLRP7, TRIM65, UNC13D, NLRP11, DAGLB, NAPEPLD, DEFB114, IL27, RICTOR, CERS6, TAC4, H2BC1, NCR3, MRGPRX1, NEAT1, METRNL, CXCL17, TAFA3, NLRP9, NLRP10, NLRP14, FFAR4, STING1, CD200R1L, IRGM, TICAM2, LILRA5, USP50, NRROS, C2CD4B, CCL4L1, C1QTNF12, RAB44, DUOXA2, MIRLET7G, MIR105-1, MIR125A, MIR126, MIR128-1, MIR129-1, MIR130B, MIR135A1, MIR136, MIR138-1, MIR140, MIR141, MIR142, MIR144, MIR145, MIR146A, MIR149, MIR15A, MIR15B, MIR16-1, MIR17, MIR181A2, MIR181B1, MIR181C, MIR187, MIR195, MIR197, MIR199A1, MIR19A, MIR19B1, MIR20A, MIR203A, MIR204, MIR205, MIR206, MIR21, MIR22, MIR221, MIR222, MIR223, MIR26A1, MIR30C2, MIR31, MIR92A1, MIR93, MIR98, CCL3L3, MIR324, MIR338, MIR361, MIR378A, MIR488, STMP1, NCF1, MSMP, MIR590, MIR657, GGT2P, CCR2, ACOD1, MIR766, MIR675, MIR920, PLA2G4B, MIR302E, MIR4286, MIR3909, CASP12, MIR6869, PYDC5 |
| Immune response GO:0006955 | 2045 | A2M, ABL1, ACP5, ACTG1, ADA, ADAM8, ADAR, ADARB1, ADCY7, ADORA2B, PARP1, AP1G1, AGER, JAG1, AHR, AIF1, AKT1, ALCAM, ALOX5, ALOX15, AMBP, ANG, ANGPT1, ANXA1, ANXA3, APCS, AIRE, BIRC2, BIRC3, XIAP, APOA1, APOA2, APOA4, APOE, APP, KLK3, FAS, FASLG, AQP4, TRIM23, ARG1, ARG2, ARRB2, ASCL2, ASS1, ATP1B1, ATP7A, AXL, AZGP1, AZU1, B2M, ADGRB1, BAX, BCL2, BCL3, BCL6, TNFRSF17, BCR, CFB, CEACAM1, PRDM1, BLK, CXCR5, BMI1, BMP6, BMPR1A, BMX, POLR3D, BPI, BRAF, BST1, BST2, BTK, BTN1A1, C1QBP, SERPING1, C1QA, C1QB, C1QC, C1R, C1S, C2, C3, C3AR1, C4A, C4B, C4BPA, C4BPB, C5, C5AR1, C6, C7, C8A, C8B, C8G, C9, CACNB3, CACNB4, CALM1, CAMK4, CAMK2A, CAMP, CASP1, CASP4, CASP6, CASP8, CAV1, CBL, CBLB, CD1A, CD1B, CD1C, CD1D, CD1E, CD2, CD3D, CD3E, CD3G, CD247, CD4, CD5L, CD6, CD7, CD8A, CD8B, CD8B2, CD14, CD19, MS4A1, CD22, CD27, CD28, CD80, CD86, TNFSF8, CD33, CD36, CD38, CD40, CD40LG, CD47, CD58, CD59, CD68, CD70, CD74, CD79A, CD79B, CD81, ADGRE5, CDC42, CDH17, CEBPB, CEBPG, CTSC, CEACAM8, CHGA, CHIT1, LYST, CHUK, CLC, CLU, CCR1, CCR3, CCR4, CCR5, CCR6, CCR7, CCR8, ACKR2, CMKLR1, LTB4R, CNR1, CNR2, COL3A1, CR1, CR1L, CR2, CREBBP, CRHR1, CRIP1, CRK, CRKL, CRP, MAPK14, CSF1, CSF1R, CSF2, CSF2RB, CSF3, CSK, CSNK1A1, CTLA4, CTSG, CTSH, CTSK, CTSL, CTSV, CTSS, CTSW, CX3CR1, CYBA, CYBB, CYLD, CYP11B1, CYP27B1, CD55, DAPK1, DAPK3, DDX1, DDX3X, DHX9, DHX15, DEFA1, DEFA3, DEFA4, DEFA5, DEFA6, DEFB1, DEFB4A, CFD, GSDME, NQO1, DMBT1, DNASE1, DNASE1L3, DNASE2, DOCK2, DRD2, DUSP3, GPR183, ECM1, EDA, EDN1, EIF2B1, ELANE, ELF1, EMP2, ADGRE1, EP300, EPHB2, EPRS1, ERCC1, EREG, ESR1, ETS1, EXT1, F2, |

F2RL1, F12, PTK2B, FAU, FCAR, FCER1A, MS4A2, FCER1G, FCER2, FCGR1A, FCGR1BP, FCGR2A, FCGR2B, FCGR3A, FCGR3B, FCGRT, FCN1, FCN2, FER, FES, FGA, FGB, FGL1, FGR, FKBP1A, FOXF1, FOXJ1, FLNB, FOSL2, FPR1, FPR2, FPR3, FRK, MTOR, FTH1, FUT7, FYB1, FYN, IFI6, XRCC6, GAPDH, GATA1, GATA2, GATA3, GATA6, GBP1, GBP2, GBP3, GCH1, GEM, GFER, GFI1, GNL1, GP2, GPI, GPLD1, CCR10, XCR1, CXCR3, GPR17, GPER1, GPR31, GPR32, GPR32P1, GPR33, FFAR3, FFAR2, GPS2, GPX1, GRB2, GRN, CXCL1, CXCL2, CXCL3, GRP, PDIA3, GSN, MSH6, GZMA, GZMB, GZMM, HCK, NCKAP1L, CFH, HFE, CFHR1, CFHR2, UBE2K, HK1, HLA-A, HLA-B, HLA-C, HLA-DMA, HLA-DMB, HLA-DOA, HLA-DOB, HLA-DPA1, HLA-DPB1, HLA-DQA1, HLA-DQA2, HLA-DQB1, HLA-DQB2, HLA-DRA, HLA-DRB1, HLA-DRB3, HLA-DRB4, HLA-DRB5, HLA-E, HLA-F, HLA-G, HLA-H, MR1, HLX, HMGB1, HMGB2, HMGB3, HMGN2, HPRT1, HPX, HRAS, HRG, HRH2, HSPA1A, HSPA1B, HSPA8, HSP90AA1, HSPD1, HTN3, ICAM1, IRF8, CFI, IFI16, IFI27, IFI35, IFIT2, IFIT1, IFIT3, IFNA1, IFNA2, IFNA4, IFNA5, IFNA6, IFNA7, IFNA8, IFNA10, IFNA13, IFNA14, IFNA16, IFNA17, IFNA21, IFNAR1, IFNAR2, IFNB1, IFNG, IFNGR1, IFNGR2, IFNW1, IGF1R, IGHA1, IGHA2, IGHD, IGHE, IGHG1, IGHG2, IGHG3, IGHG4, IGHM, JCHAIN, IGKC, RBPJ, IGLC1, IGLC2, IGLC3, IGLC6, IGLL1, IKBKB, IL1A, IL1B, IL1R1, IL1RAP, IL1RN, IL2, IL2RA, IL2RB, IL2RG, IL3, IL4, IL4R, IL5, IL5RA, IL6, IL6R, IL6ST, IL7, IL7R, CXCL8, CXCR1, IL9, CXCR2, IL9R, IL10, IL10RB, IL12A, IL12B, IL12RB1, IL13, IL13RA2, IL15, IL16, IL17A, IL18, IDO1, CXCL10, INS, INPP5D, INPPL1, IRAK1, IRAK2, IRF1, IRF3, IRF4, IRF5, IRF7, ISG20, ITGA4, ITGAD, ITGAL, ITGAM, ITGB2, ITGB6, ITGB7, ITGB8, ITK, JAK1, JAK2, JAK3, JUNB, KARS1, KCNJ8, KCNN4, KIF5B, KIR2DL1, KIR2DL3, KIR2DL4, KIR2DS1, KIR2DS5, KIR3DL1, KIR3DS1, KIT, KLRB1, KLRC1, KLRC2, KLRC3, KLRD1, IPO5, KRT1, KRT6A, KRT16, LAG3, LAIR1, LAMP1, LBP, LCK, LCN2, LCP1, LCP2, LEP, LFNG, LGALS1, LGALS3, LGALS4, LGALS9, LIF, LIG4, LIMK1, LIPA, LRCH4, LRP1, LTA, LTB, LTBR, LTF, LY9, CD180, LY75, LYN, SH2D1A, LYZ, SMAD3, SMAD6, SMAD7, MBL2, MBP, CD46, MDK, MEF2C, MEFV, MAP3K1, MAP3K5, MEN1, MFNG, CIITA, MICB, MIF, CXCL9, MLH1, KMT2A, MMP3, MMP12, MNDA, MOG, CD200, MPL, MRC1, CITED1, MSH2, MST1R, MUC7, MX1, MX2, MYB, MYD88, MYO1C, NAGLU, NAIP, NBN, NCF2, NCK1, NEDD4, NFATC2, NFE2L2, NFIL3, NFKB1, NFKB2, NFKBIA, NFKBIL1, NINJ1, NKG7, NMB, NMBR, NONO, NOS2, NOTCH1, NOTCH2, PNP, NPY5R, OAS1, OAS2, OAS3, OPRD1, OPRK1, OSM, P2RX7, FURIN, PRDX1, PAK1, PAK2, PAK3, REG3A, PAWR, PCBP2, PCK1, PDCD1, PDE4B, PDE4D, ENPP1, ENPP2, ENPP3, PDPK1, PECAM1, PF4, PF4V1, CFP, PGC, PHB1, PI3, SERPINB9, PIGR, PIK3CA, PIK3CD, PIK3CG, PIK3R1, PIK3R2, PLA2G1B, PLA2G4A, PLA2G5, PLCG1, PLCG2, PLD2, PLEC, PLSCR1, PML, EXOSC9, PMS2, POU2AF1, POU2F2, PPARG, MED1, PPBP, PPP2CA, PPP3CB, PPP6C, PRF1, PRG2, PRKCB, PRKCD, PRKCE, PRKCH, PKN1, PRKD1, PRKCQ, PRKCZ, PRKDC, MAPK1, MAPK8, MAPK9, MAPK10, MAP2K6, MAP2K7, EIF2AK2, PRNP, PRSS2, PRSS3, MASP1, KLK7, PRTN3, PSEN1, PSG9, PSMA1, PSMB4, PSMB10, PTAFR, PTGDR, PTGDS, PTGER4, PTK2, PTK6, PTPN1, PTPN2, PTPN6, PTPN11, PTPRC, PTPRD, PTPRJ, PTPRS, PTX3, PVR, NECTIN1, NECTIN2, MAP4K2, RAB27A, RAC2, RAF1, RAG1, RAG2, RAP1GAP, RARA, RARRES2, REL, REG1A, REG1B, RELA, RELB, TRIM27, RFPL1, RFX1, RGS1, RNASE2, RNASE3, RNASE4, RNASE6, BRD2, RORA, RORC, RPL30, RPL39, RPS3, RPS6KA3, RPS6KB1, RPS19, S100A7, S100A8, S100A9, S100A12, S100A13, SERPINB4, SCNN1B, CCL1, CCL2, CCL3, CCL3L1, CCL4, CCL5, CCL7, CCL8, CCL11, CCL13, CCL14, CCL15, CCL16, CCL17, CCL18, CCL19, CCL20, CCL21, CCL22, CCL23, CCL24, CCL25, CXCL6, CXCL11, CXCL5, XCL1, CX3CL1, CXCL12, SEC14L1, SECTM1, SEMG1, SEMG2, MAP2K4, SFPQ, SFTPD, SHB, SHMT2, ST6GAL1, ST3GAL1, SIPA1, SKP2, SLAMF1, SLC11A1, SLC15A2, SLC18A2, SLC22A5, SLPI, SMPD1, SNCA, SOS1, SP2, SP100, UAP1, SPI1, SPN, SPRR2A, SRC, SRMS, SRPK1, SRPK2, TRIM21, STAT1, STAT2, STAT3, STAT4, STAT5B, STAT6, STK11, STX4, STXBP1, STXBP2, STXBP3, SUPT6H, VAMP2, VAMP7, XCL2, SYK, ADAM17, MAP3K7, TAP1, TAP2, TARBP2, TCF7, TCF12, TRA, TRAV6, TRB, TRGC1, TRGC2, TRGV1, TRGV2, TRGV3, TRGV4, TRGV5, TRGV8, TRGV9, TRGV10, TRGV11, CRIPTO, TEC, TF, TFE3, TFRC, TGFB1, TGFB2, TGFB3, TGFBR3, THBS1, THY1, TLR1, TLR2, TLR3, TLR4, TLR5, TSPAN6, TNF, TNFAIP1, TNFAIP3, TNFRSF1B, TP53, TP53BP1, TRAF2, TRAF3, TRAF6, TSC1, TTC4, TWIST1, TNFSF4, TNFRSF4, TXK, TYK2, TYRO3, TYROBP, UBA7, UBE2N, UFD1, UMOD, UNG, NR1H2, VAV1, VAV2, EZR, VIM, VPREB1, VTN, WAS, LAT2, NSD2, WNT5A, XBP1, XRCC5, YES1, YWHAE, YWHAZ, ZAP70, TRIM25, TRIM26, ZP3, ZYX, LRP8, LAPTM5, PXDN, IL1R2, CXCR4, FZD5, MAPKAPK3, BAG6, LST1, TFEB, NR4A3, FOSL1, SLC7A5, AKAP1, MADCAM1, KDM5D, EPX, DYSF, KLRC4, EOMES, H2BC8, H2BC7, H2BC6, H2BC10, H2BC4, H2BC21, PLA2G6, STX7, DHX16, GPR65, SEMA7A, PIK3R3, IKBKG, IFITM1, DGKZ, CST7, APOL1, AP3B1, FCN3, PRKRA, TNFSF11, SKAP1, RNASET2, OASL, VAMP8, BECN1, MARCO, S1PR4, SNX4, GBF1, TNFSF14, TNFSF13, TNFSF12, TNFSF10, TNFSF9, ADAM15, CD164, TNFRSF14, RIPK2, FADD, SNAP23, RIOK3, TNFRSF11A, IL18RAP, IL1RL2, IL18R1, BANF1, CD84, APLN, VNN1, SQSTM1,

TAX1BP1, EIF2B4, EIF2B3, EIF2B2, EIF2B5, ENDOU, BCL10, RAB29, KYNU, H2BC11, PGLYRP1, TNFSF18, MAP3K14, SOCS3, CH25H, CCRL2, PSTPIP1, CLDN1, USP14, FCGR2C, NMI, ATG12, EXO1, EBAG9, IL1RL1, ARHGEF2, DDX21, AURKB, FCMR, IL32, PNMA1, UBE2L6, MFHAS1, MAPKAPK2, CD83, VAMP3, SLC22A13, LPXN, NCR1, AIM2, LY86, ITM2A, IL27RA, EIF4E2, THEMIS2, ATG5, STX8, LITAF, CXCL14, BCAR1, GTPBP1, NR1D1, APOBEC3B, TRAF4, ISG15, IKBKE, SOCS5, PHF14, N4BP1, LRRC14, PUM1, TBKBP1, HDAC4, MATR3, TRIM14, TESPA1, GAB2, TRIL, PJA2, TOMM70, G3BP2, USP15, TNFSF15, NR1H4, TANK, PARP3, NR1H3, IL18BP, PQBP1, CERT1, ACTR3, ACTR2, TRIM10, RASGRP1, OPTN, G3BP1, EBI3, TRIM28, CHST4, CNIH1, TENM1, TRIM13, FLOT1, PRG4, KLRG1, CD96, CALCOCO2, IGSF6, AKAP8, LILRB2, TCIRG1, TNIP1, CRISP3, CLEC4M, TLR6, CCL26, TRIM22, TUBB4B, BTN3A3, BTN2A2, NOD1, PRG3, WFDC2, SPAG11B, IFITM3, RAPGEF3, SPON2, RBM14, CDC42EP2, VAV3, MAD2L2, CLEC10A, TRIM38, SYNCRIP, SEMA4D, SEMA3C, KAT5, IPO7, UBD, BATF, IFI44, CXCL13, GNLY, IFITM2, COLEC10, SH2B2, TRIM3, HEXIM1, RBCK1, POLR3F, POLR3G, POLR3C, KHDRBS1, CXCR6, CD226, TNFSF13B, CNPY3, RFPL3, RFPL2, MASP2, TRAF3IP2, PLK2, CYSLTR1, CCR9, CPLX2, CCL27, LILRB1, ARID5A, USP20, HCST, FGL2, CFHR4, CFHR3, MALT1, TRAFD1, BTNL3, SERINC3, IFI44L, LILRB5, SPINK5, LILRB4, TMED1, LILRA1, LILRB3, LILRA3, LILRA2, RIPK3, MID2, RAPGEF4, TRIM31, ADAMTS13, BTN3A2, BTN3A1, BTN2A1, CD160, POLR3A, CDC37, HHLA2, CORO1A, IRAK3, DUSP10, PTGDR2, USP18, TREX1, SCN11A, CD300A, PARK7, VSIG4, PHB2, RAB11FIP2, NLRP1, ZBTB1, CARD8, SBNO2, KLRK1, SIRT2, SCAP, TRIM32, PAXIP1, ENDOD1, SWAP70, TRIM35, SARM1, RAP1GAP2, KDM6B, N4BP3, FCHO1, GRAMD4, TTLL12, RFTN1, PLEKHM2, DTX4, PLCL2, ICOSLG, PUM2, SIRT1, BRD4, TNFRSF13B, MORC3, RPL13A, CLCF1, PIK3R5, ATP6V0A2, LILRA4, PADI4, CDC42EP4, RIGI, CLEC5A, CD2AP, MKRN2, LY96, TRIM29, CADM1, IL17RA, IFIT5, PANX1, SLC39A6, TTLL1, KLK5, FBXL2, PRKD2, TRIM58, ZDHHC5, SAMHD1, SIN3A, TKFC, TMEM98, ANKRD17, GIGYF2, APPL1, LSM14A, PTPN22, STAP1, CLEC4E, FBXO9, PLA2G2D, PPP1R14B, IL36RN, LAT, LAMP3, FOXP1, IL36B, IL37, IL36A, IL17B, C5AR2, SIT1, TNFRSF21, SMPDL3B, CRCP, ADAMDEC1, RABGEF1, APOBEC3C, IGKV1-5, IGHV8-51-1, IGHV7-81, IGHV6-1, IGHV5-10-1, IGHV5-51, IGHV4-38-2, IGHV4-61, IGHV4-59, IGHV4-39, IGHV4-34, IGHV4-31, IGHV4-30-4, IGHV4-28, IGHV4-4, IGHV3-38-3, IGHV3-74, IGHV3-73, IGHV3-72, IGHV3-66, IGHV3-64, IGHV3-53, IGHV3-49, IGHV3-48, IGHV3-43, IGHV3-38, IGHV3-35, IGHV3-33, IGHV3-30, IGHV3-23, IGHV3-21, IGHV3-20, IGHV3-16, IGHV3-15, IGHV3-13, IGHV3-11, IGHV3-9, IGHV3-7, IGHV2-70, IGHV2-26, IGHV2-5, IGHV1-69-2, IGHV1-38-4, IGHV1-69, IGHV1-58, IGHV1-45, IGHV1-24, IGHV1-18, IGHV1-8, IGHV1-3, IGHJ1, IGHD1-1, DLL1, TRDV3, TRDV2, TRDV1, TRDJ1, TRDD1, TRDC, TRBV30, TRBV29-1, TRBV28, TRBV27, TRBV25-1, TRBV24-1, TRBV23-1, TRBV20-1, TRBV19, TRBV18, TRBV17, TRBV16, TRBV14, TRBV13, TRBV12-5, TRBV12-4, TRBV12-3, TRBV11-3, TRBV11-2, TRBV11-1, TRBV10-3, TRBV10-2, TRBV10-1, TRBV9, TRBV7-9, TRBV7-8, TRBV7-7, TRBV7-6, TRBV7-4, TRBV7-3, TRBV7-2, TRBV7-1, TRBV6-9, TRBV6-8, TRBV6-7, TRBV6-6, TRBV6-5, TRBV6-4, TRBV6-3, TRBV6-1, TRBV5-8, TRBV5-7, TRBV5-6, TRBV5-5, TRBV5-4, TRBV5-3, TRBV5-1, TRBV4-3, TRBV4-2, TRBV4-1, TRBV3-1, TRBV2, TRBJ2-7, TRBJ2-6, TRBJ2-5, TRBJ2-4, TRBJ2-3, TRBJ2-2, TRBJ2-1, TRBJ1-6, TRBJ1-5, TRBJ1-4, TRBJ1-3, TRBJ1-2, TRBJ1-1, TRBD1, TRBC2, TRBC1, TRAV41, TRAV40, TRAV39, TRAV38-2DV8, TRAV38-1, TRAV36DV7, TRAV35, TRAV34, TRAV30, TRAV29DV5, TRAV27, TRAV26-2, TRAV26-1, TRAV25, TRAV24, TRAV23DV6, TRAV22, TRAV21, TRAV20, TRAV19, TRAV18, TRAV17, TRAV16, TRAV14DV4, TRAV13-2, TRAV13-1, TRAV12-3, TRAV12-2, TRAV12-1, TRAV10, TRAV9-2, TRAV9-1, TRAV8-6, TRAV8-4, TRAV8-3, TRAV8-2, TRAV8-1, TRAV7, TRAV5, TRAV4, TRAV3, TRAV2, TRAV1-2, TRAV1-1, TRAJ42, TRAJ31, TRAJ3, TRAC, IGLV11-55, IGLV10-54, IGLV9-49, IGLV8-61, IGLV7-46, IGLV7-43, IGLV6-57, IGLV5-52, IGLV5-48, IGLV5-45, IGLV5-39, IGLV5-37, IGLV4-69, IGLV4-60, IGLV4-3, IGLV3-32, IGLV3-27, IGLV3-25, IGLV3-22, IGLV3-21, IGLV3-19, IGLV3-16, IGLV3-12, IGLV3-10, IGLV3-9, IGLV3-1, IGLV2-33, IGLV2-23, IGLV2-18, IGLV2-14, IGLV2-11, IGLV2-8, IGLV1-51, IGLV1-50, IGLV1-47, IGLV1-44, IGLV1-40, IGLV1-36, IGLJ1, IGLC7, IGKV6D-41, IGKV6D-21, IGKV3D-20, IGKV3D-15, IGKV3D-11, IGKV3D-7, IGKV2D-30, IGKV2D-29, IGKV2D-28, IGKV2D-26, IGKV2D-24, IGKV1D-43, IGKV1D-42, IGKV1D-39, IGKV1D-37, IGKV1D-33, IGKV1D-17, IGKV1D-13, IGKV1D-12, IGKV1D-8, IGKV6-21, IGKV5-2, IGKV4-1, IGKV3-20, IGKV3-15, IGKV3-7, IGKV2-40, IGKV2-30, IGKV2-29, IGKV2-28, IGKV2-24, IGKV1-39, IGKV1-37, IGKV1-27, IGKV1-17, IGKV1-16, IGKV1-13, IGKV1-12, IGKV1-9, IGKV1-8, IGKV1-6, IGKJ1, RGCC, DBNL, SETD2, PHPT1, PYCARD, TBK1, CD274, BLNK, ZDHHC1, VPREB3, ICOS, CNOT7, MYL11, HCFC2, IL19, UBQLN1, NOP53, TBX21, ADGRE2, CD209, STOML2, PLA2G3, IL21R, TRAT1, CLEC4A, FOXP3, EXOSC3, ZBTB7B, KMT5B, IRAK4, LEF1, HERC5, MARCHF2, C1RL, TLR7, ECSIT, SLC15A3, BPIFA1, TLR8, SPG21, YTHDF2, UBE2J1, ACKR4, IL23A, CYRIB, NUB1, GPRC5B, POLR3K, MSRB1, CD244, ERAP1, UBASH3A, IL20RB, TRIM34, TLR9, H2BC12L, TREM2, TREM1, SASH3,

TOLLIP, RBM47, APBB1IP, SHLD2, RC3H2, PARP14, BTN2A3P, OTUD4, TRPM4, SAMD9, DPP8, ANKHD1, LAX1, DUS2, LIME1, RNF125, PPP2R3C, MARCHF1, BANK1, SUSD4, RNF31, AKIRIN2, TRIM68, TMEM33, RIF1, APPL2, ARL8B, ATAD3A, TRIM62, WDR41, PSPC1, UBE2W, SHFL, LGR4, NAGK, DDX60, OTUB1, DOCK10, LYAR, RAB20, NLRP2, NPLOC4, POLR3B, POLR3E, INAVA, PAG1, SELENOS, ITFG2, ERBIN, BTNL2, CRTAM, CTNNBL1, IL36G, METTL3, CCL28, SUCNR1, ZC3HAV1, IFNK, SLAMF8, SPHK2, TCIM, WRNIP1, SPIRE1, DHX33, GPR108, SPPL2B, DUSP22, HMCES, OTUD7B, ACKR3, ENTPD7, TRIM49, FERRY3, CYSLTR2, PGLYRP4, CD177, RTN4, PELI1, ZNFX1, SLC39A10, MCOLN1, IGHV7-4-1, IGHV3-30-3, SENP7, AICDA, S100A14, MAVS, PDP2, WDFY1, USP29, EPG5, MARK4, HAMP, SLAMF7, HMHB1, ZP4, SLC46A2, CXCL16, NLRC4, CACTIN, IL21, PLEKHA1, FAM3A, BACH2, APOBEC3G, GPATCH3, MYO1G, SAMSN1, VSIR, NOD2, IFIH1, ERAP2, CARD9, SEMA4A, RAB17, NFKBIZ, AZI2, DCLRE1C, CLEC7A, PLA2G2F, ILRUN, IL25, LRRC19, SLC26A6, WNK1, CHID1, COLEC11, CUEDC2, PVRIG, TRIM48, DHX58, TNIP2, LILRA6, BRCC3, TMEM43, FCRL2, CYBC1, ULBP3, MUL1, TNFAIP8L2, RHBDF2, NLRX1, VTCN1, GSDMD, ZDHHC11, BTNL8, ATAD5, TNIP3, ATAT1, SVEP1, ZC3H12A, RNF34, CCDC92, ALPK1, TASL, CEP63, ULBP2, ULBP1, TRAF3IP3, PDCD1LG2, CD276, PRR7, NDFIP1, CPTP, ZBP1, COLEC12, CFHR5, FBXO38, TRIM11, CLPB, TRIM8, UNC93B1, DEFB126, TRIM7, RNF170, TLR10, TRIM56, FCRL5, FCRL4, PARP9, JAM3, ITCH, FCAMR, USP44, PRAM1, NLRC5, SLA2, TMEM126A, ZDHHC18, RNF135, CARD11, SANBR, IL1F10, USP38, RNASE7, LOXL3, CRACR2A, TRIM51, KMT5C, NFKBID, FCRLA, HAVCR2, ORAI1, ZDHHC12, SPPL2A, RAB2B, ZNRF1, UBL7, H2BC12, TRIM5, ZCCHC3, TSLP, GALP, TRIM4, IGHV3-30-5, SIGLEC10, KLHL6, TRIM15, SLAMF9, OTULIN, IL33, TRIM41, RNF185, RSAD2, TIFA, BPIFB1, PKHD1L1, NEURL3, CLEC6A, GARIN5A, IL17F, VPS26B, TMEM106A, NLRP3, TIRAP, ERMAP, PGLYRP2, PGLYRP3, SLAMF6, CGAS, FCRL3, GBP4, GBP5, TNFRSF13C, RNF166, CLNK, LEAP2, SH2D1B, MRGPRX2, DEFB118, TRIM6, EXOSC6, PIK3AP1, TRIM64, TMEM45B, TRIM51G, SLC15A4, SPPL3, PLD4, RNASE8, CMTM3, SPNS2, TMIGD2, LYPLAL1, RNF19B, FCRLB, WFDC12, CST9LP1, CST9L, CST9, TRIM43, REG3G, CD200R1, OTOP1, IL31RA, GPR151, IRAK1BP1, SDHAF4, RAET1E, TRIM40, DOCK11, APOBEC3D, DEFB104A, NEK7, WFDC3, SMCR8, ROMO1, WFDC10A, DEFB127, SIRPA, LACC1, IL34, CD300LF, PIK3R6, NLRP4, TICAM1, ZNRF4, GCSAML, RC3H1, DCST1, IL23R, PYHIN1, WFDC5, SHLD1, NFAM1, DTX3L, BTLA, PYDC2, DAB2IP, BTNL9, RAET1L, NKX2-3, ZFPM1, GBP6, DENND1B, IFNLR1, WFDC13, APOBEC3H, RNF168, SLC30A8, CLEC4C, DHX36, NLRP6, POLR3H, PLPP4, PIANP, NLRC3, FYB2, APOBEC3A, APOBEC3F, APLF, TRIM65, UNC13D, PDE12, GAPT, TUBB, TRIML2, MPEG1, MARCHF8, DEFB105A, DEFB106A, DEFB107A, DEFB108B, DEFB110, DEFB112, DEFB113, DEFB114, DEFB115, DEFB116, DEFB119, DEFB121, DEFB123, DEFB124, DEFB125, DEFB128, PELI3, IL27, STXBP4, YTHDF3, BPIFC, MCOLN2, SERINC5, ZNF683, GCSAM, NCR3, WFDC11, WFDC9, IL4I1, PYDC1, WFDC10B, IFNL2, IFNL3, IFNL1, TRIM49B, MILR1, HMSD, SSC5D, TREML4, NLRP2B, TRIM59, NLRP9, NLRP10, CLEC4D, IFNE, RAB7B, RAB43, CLEC4G, TRIML1, STING1, TREML1, RFPL4A, FCRL6, IRGM, RAET1G, KAAG1, TICAM2, LILRA5, BPIFB3, USP50, NCR3LG1, NRROS, USP17L2, THEMIS, GPR15LG, CLEC2A, CLEC12B, SCIMP, CCL4L1, GBP7, USP27X, TRIM77, TRIM49D1, DEFB132, RAB44, SMIM30, PLPP6, DEFB133, MIRLET7B, MIR105-1, MIR136, MIR140, MIR146A, MIR149, MIR17, MIR18A, MIR181B1, MIR19A, MIR19B1, MIR20A, MIR200B, MIR200C, MIR21, MIR302A, MIR34A, CCL3L3, EIF2AK4, TARM1, DEFB107B, DEFB104B, DEFB108A, DEFB106B, DEFB105B, MIR520E, MIR520B, DEFB135, DEFB136, DEFB134, SCART1, TRIM64B, TRIM49C, DEFB131A, TRIM64C, STMP1, TRIM43B, NCF1, SPAG11A, DEFA1B, CCR2, TRIM49D2, RFPL4AL1, ACOD1, MIR758, MIR708, BTNL10P, DEFB131B, KIR2DS2, CD24, C17orf99, MIR302E, IGLL5, KLRF2, MICA, KLRC4-KLRK1, IFNL4, MIR6869, IGHV2-70D, IGHV1-69D, IGHV3-64D, IGHV3-43D, PYDC5, SHLD3

**Supplementary Table 10 : List of publicly available datasets used in this study**

| Dataset | Cell Line | Accession Number | Reference |
|---|---|---|---|
| 53BP1 ChIP-seq | U2OS-DIvA | E-MTAB-5817 | Clouaire et al., 2018 |
| LIG4 ChIP-seq | U2OS-DIvA | E-MTAB-5817 | Clouaire et al., 2018 |
| BLESS | U2OS-DIvA | E-MTAB-5817 | Clouaire et al., 2018 |
| RAD51 ChIP-seq 4h | U2OS-DIvA | E-MTAB-1241 | Aymard et al., 2014 |
| XRCC4 ChIP-seq | U2OS-DIvA | E-MTAB-1241 | Aymard et al., 2014 |
| yH2AX ChIP-seq | U2OS-DIvA | E-MTAB-8851 | Arnould et al., 2021 |
| p-ATM ChIP-seq | U2OS-DIvA | E-MTAB-8851 | Arnould et al., 2021 |
| RAD51 ChIP-seq 24h | U2OS-DIvA | E-MTAB-11592 | Cohen, Guenolé, Lazar et al., 2022 |
| RPA ChIP-seq | U2OS-DIvA | E-MTAB-13197 | Saur, Lesage et al., 2025 |
| RNA Pol II ChIP-seq | U2OS-DIvA | E-MTAB-13197 | Saur, Lesage et al., 2025 |
| RNA-seq | U2OS-DIvA | E-MTAB-6318 | Cohen et al., 2018 |
| H3K4me3 | U2OS-DIvA | E-MTAB-15639 (private link available upon request) | Gong et al, 2017 |

**Additional Figure 1 :** Single-cell differential chromatin accessibility profiles as log2(+DSB/-DSB) for each of the 80 AsiSI-induced DSBs (n=300 cells).

Left panels represent differential accessibility on a 10 kb window (± 5 kb from the DSB) after 4 h of DSB induction. Right panels represent differential accessibility on an 80 kb window (± 40 kb from the DSB) after 24 h of DSB induction.

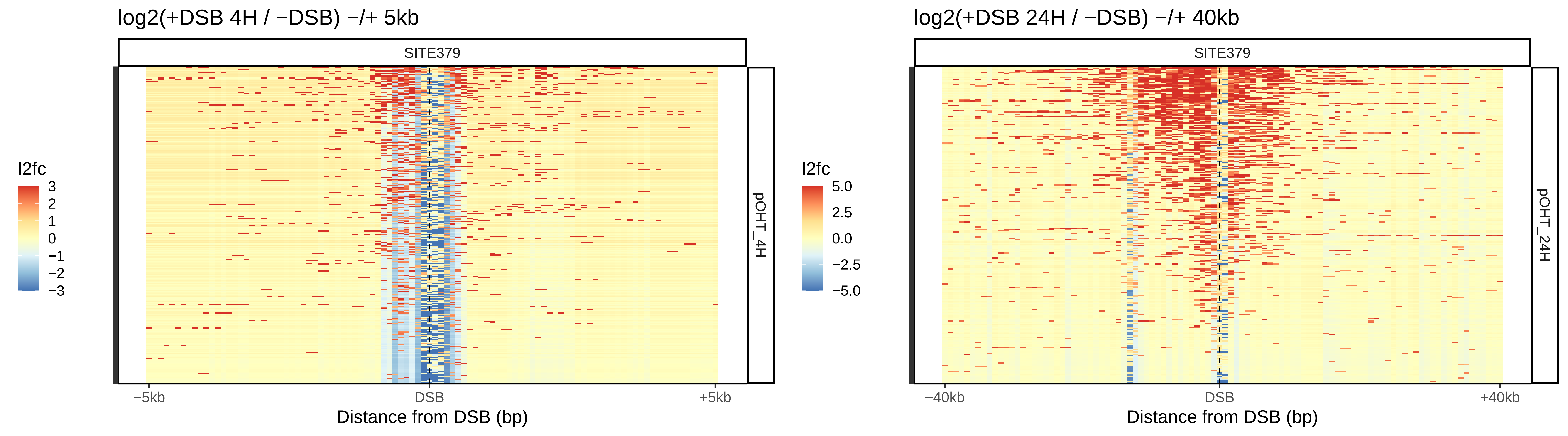

log2(+DSB 4H / −DSB) −/+ 5kb
SITE379
l2fc
3
2
1
0
−1
−2
−3
pOHT_4H
−5kb
DSB
+5kb
Distance from DSB (bp)
log2(+DSB 24H / −DSB) −/+ 40kb
SITE379
l2fc
5.0
2.5
0.0
−2.5
−5.0
pOHT_24H
−40kb
DSB
+40kb
Distance from DSB (bp)

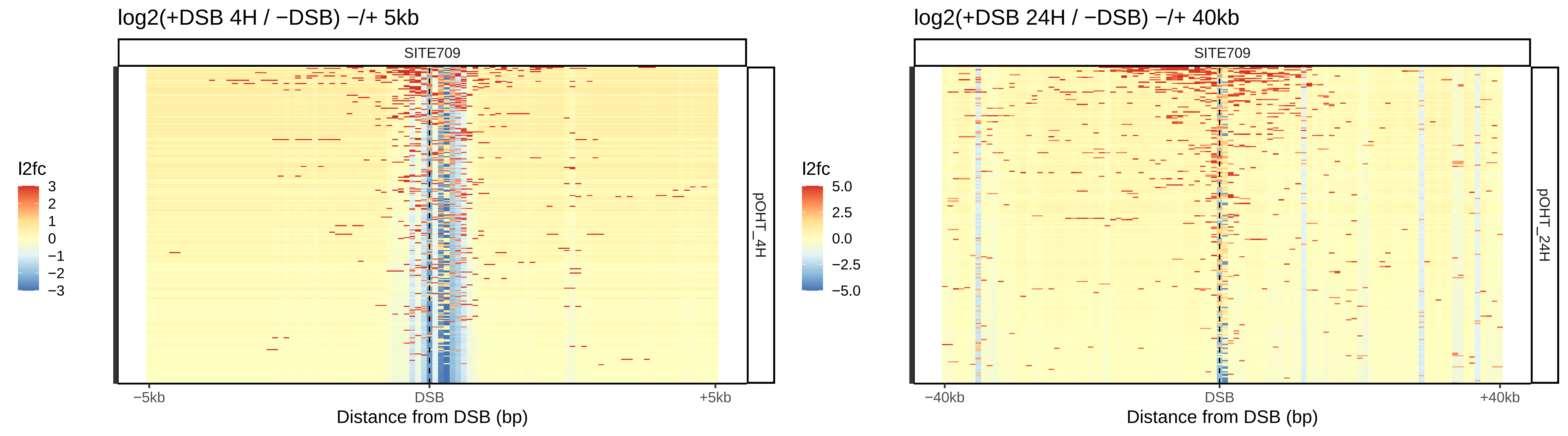

log2(+DSB 4H / −DSB) −/+ 5kb
SITE709
l2fc
3
2
1
0
−1
−2
−3
pOHT_4H
−5kb
DSB
+5kb
Distance from DSB (bp)
log2(+DSB 24H / −DSB) −/+ 40kb
SITE709
l2fc
5.0
2.5
0.0
−2.5
−5.0
pOHT_24H
−40kb
DSB
+40kb
Distance from DSB (bp)

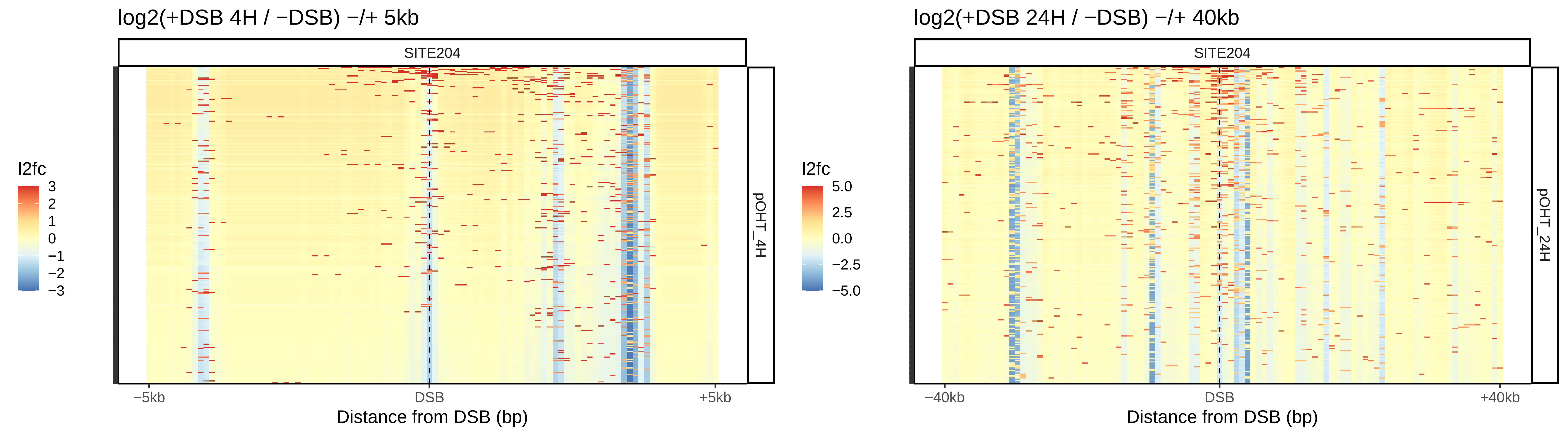

log2(+DSB 4H / −DSB) −/+ 5kb
SITE204
pOHT_4H
l2fc
3
2
1
0
−1
−2
−3
−5kb
DSB
+5kb
Distance from DSB (bp)
log2(+DSB 24H / −DSB) −/+ 40kb
SITE204
pOHT_24H
l2fc
5.0
2.5
0.0
−2.5
−5.0
−40kb
DSB
+40kb
Distance from DSB (bp)

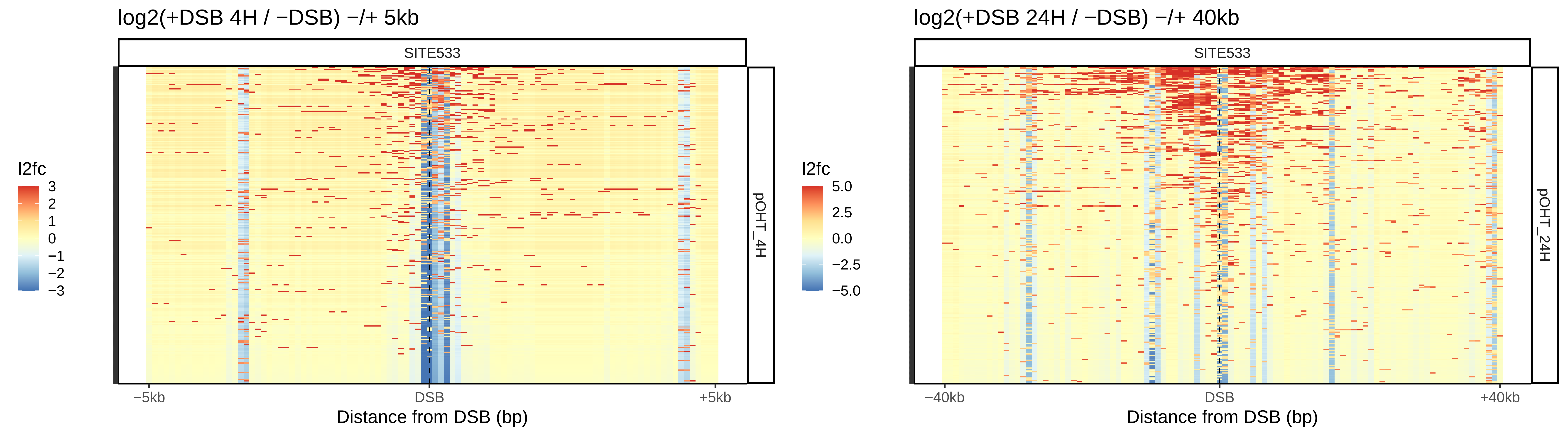

log2(+DSB 4H / −DSB) −/+ 5kb
SITE533
l2fc
3
2
1
0
−1
−2
−3
pOHT_4H
−5kb
DSB
+5kb
Distance from DSB (bp)
log2(+DSB 24H / −DSB) −/+ 40kb
SITE533
l2fc
5.0
2.5
0.0
−2.5
−5.0
pOHT_24H
−40kb
DSB
+40kb
Distance from DSB (bp)

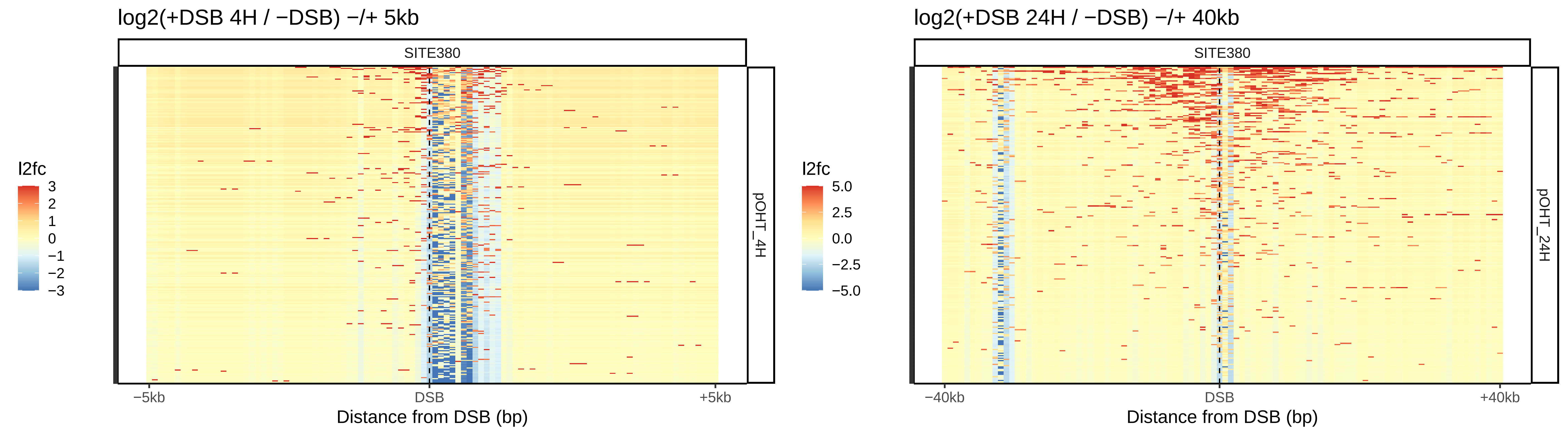

log2(+DSB 4H / −DSB) −/+ 5kb
SITE380
pOHT_4H
l2fc
3
2
1
0
−1
−2
−3
−5kb
DSB
+5kb
Distance from DSB (bp)
log2(+DSB 24H / −DSB) −/+ 40kb
SITE380
pOHT_24H
l2fc
5.0
2.5
0.0
−2.5
−5.0
−40kb
DSB
+40kb
Distance from DSB (bp)

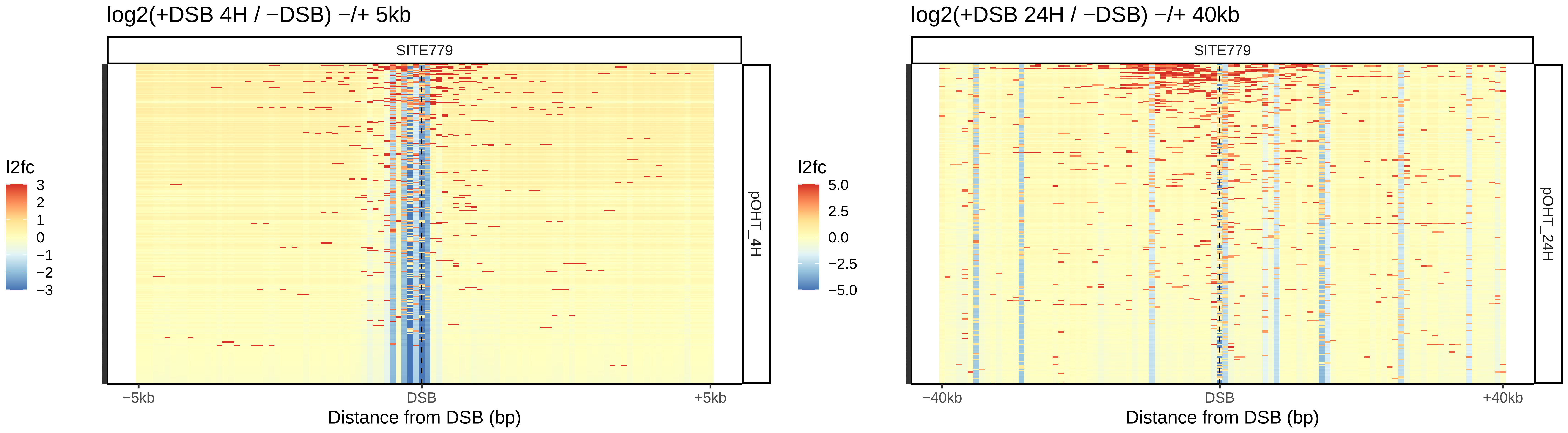
log2(+DSB 4H / −DSB) −/+ 5kb
SITE779
l2fc
3
2
1
0
−1
−2
−3
pOHT_4H
−5kb
DSB
+5kb
Distance from DSB (bp)
log2(+DSB 24H / −DSB) −/+ 40kb
SITE779
l2fc
5.0
2.5
0.0
−2.5
−5.0
pOHT_24H
−40kb
DSB
+40kb
Distance from DSB (bp)

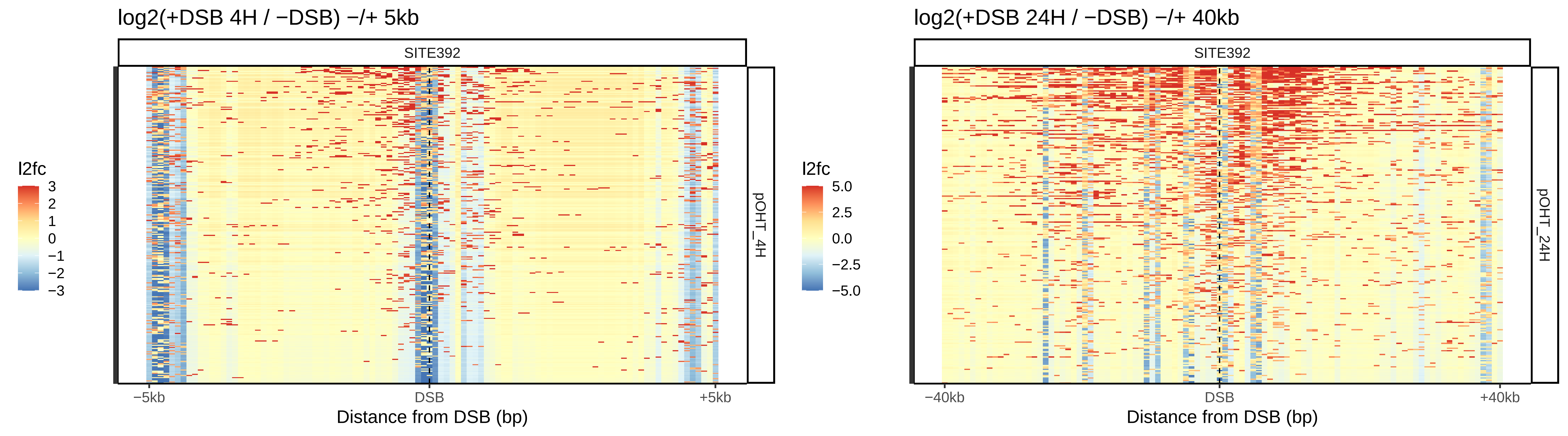
log2(+DSB 4H / −DSB) −/+ 5kb
SITE392
l2fc
3
2
1
0
−1
−2
−3
pOHT_4H
−5kb
DSB
+5kb
Distance from DSB (bp)
log2(+DSB 24H / −DSB) −/+ 40kb
SITE392
l2fc
5.0
2.5
0.0
−2.5
−5.0
pOHT_24H
−40kb
DSB
+40kb
Distance from DSB (bp)

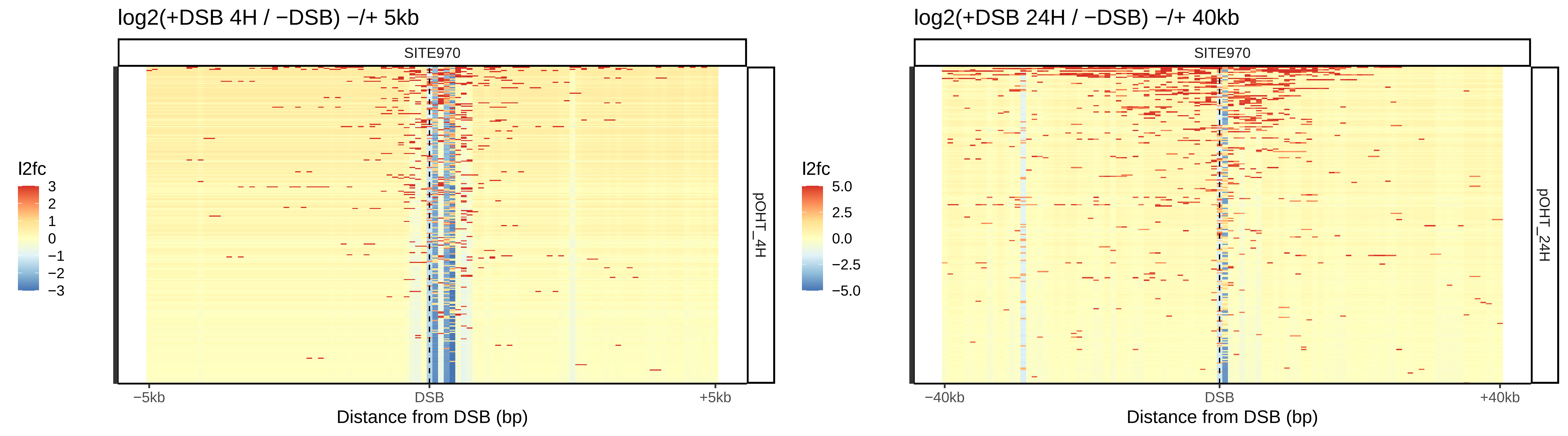

log2(+DSB 4H / −DSB) −/+ 5kb
SITE970
l2fc
3
2
1
0
−1
−2
−3
pOHT_4H
−5kb
DSB
+5kb
Distance from DSB (bp)
log2(+DSB 24H / −DSB) −/+ 40kb
SITE970
l2fc
5.0
2.5
0.0
−2.5
−5.0
pOHT_24H
−40kb
DSB
+40kb
Distance from DSB (bp)

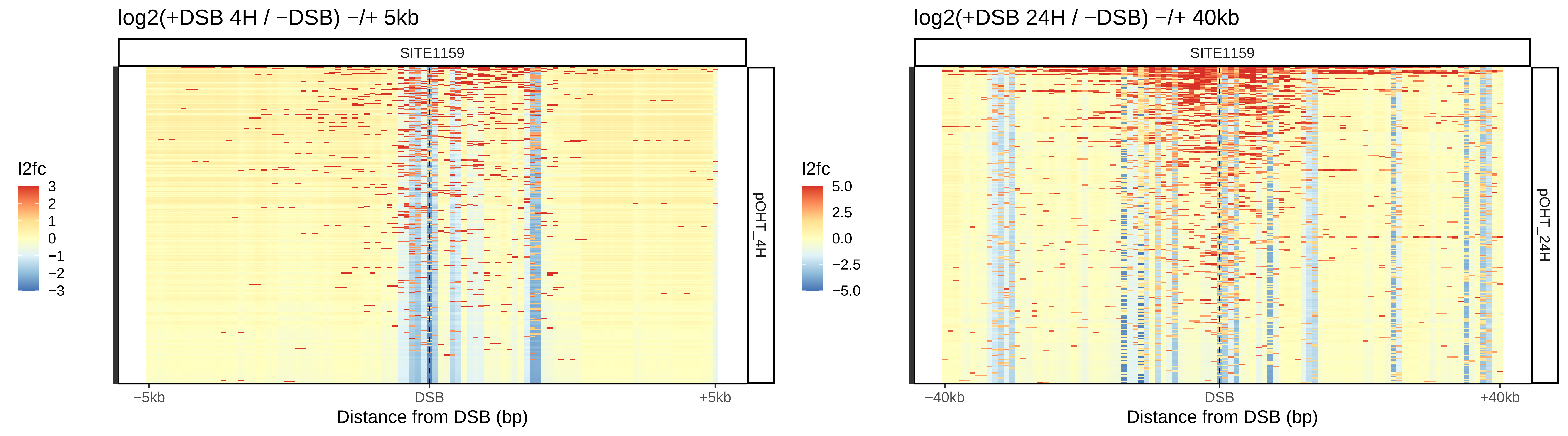
log2(+DSB 4H / −DSB) −/+ 5kb
SITE1159
l2fc
3
2
1
0
−1
−2
−3
pOHT_4H
−5kb
DSB
+5kb
Distance from DSB (bp)
log2(+DSB 24H / −DSB) −/+ 40kb
SITE1159
l2fc
5.0
2.5
0.0
−2.5
−5.0
pOHT_24H
−40kb
DSB
+40kb
Distance from DSB (bp)

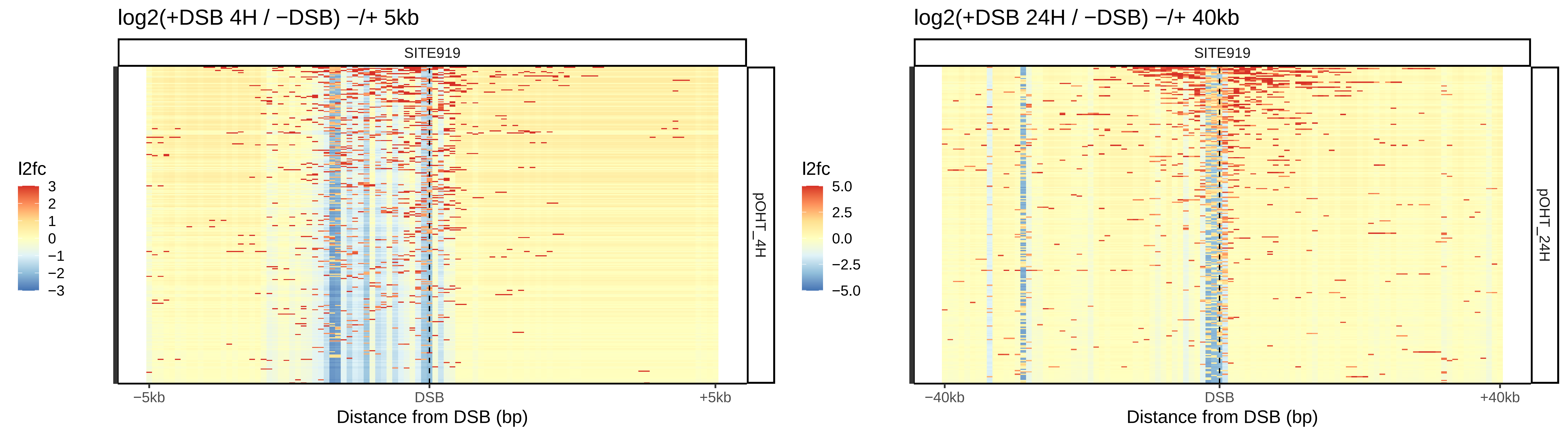
log2(+DSB 4H / −DSB) −/+ 5kb
SITE919
l2fc
3
2
1
0
−1
−2
−3
pOHT_4H
−5kb
DSB
+5kb
Distance from DSB (bp)
log2(+DSB 24H / −DSB) −/+ 40kb
SITE919
l2fc
5.0
2.5
0.0
−2.5
−5.0
pOHT_24H
−40kb
DSB
+40kb
Distance from DSB (bp)

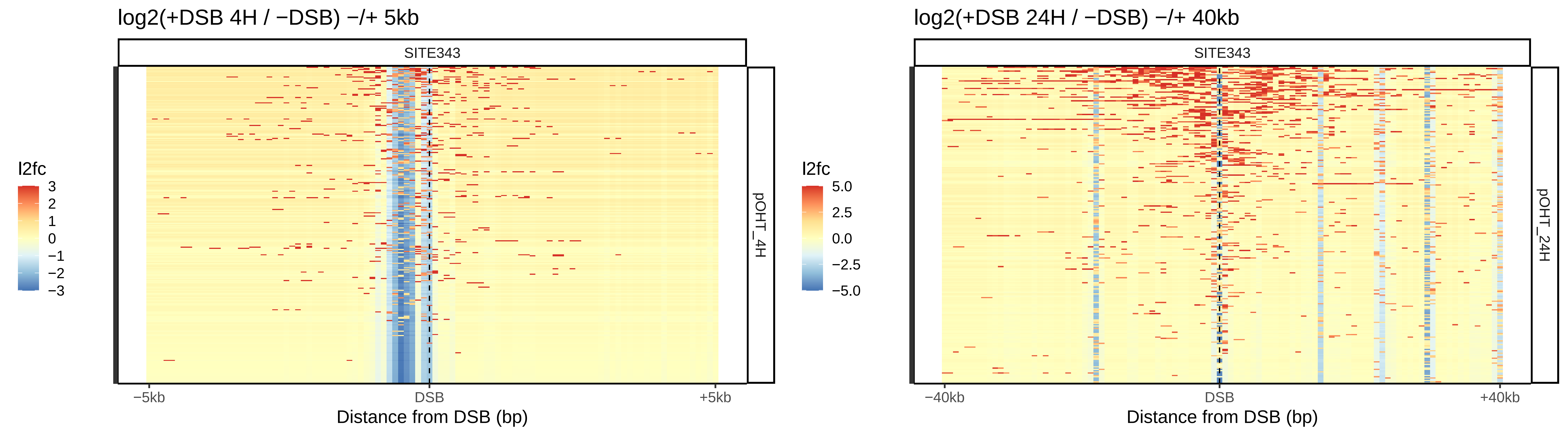

log2(+DSB 4H / −DSB) −/+ 5kb
SITE343
l2fc
3
2
1
0
−1
−2
−3
pOHT_4H
−5kb
DSB
+5kb
Distance from DSB (bp)
log2(+DSB 24H / −DSB) −/+ 40kb
SITE343
l2fc
5.0
2.5
0.0
−2.5
−5.0
pOHT_24H
−40kb
DSB
+40kb
Distance from DSB (bp)

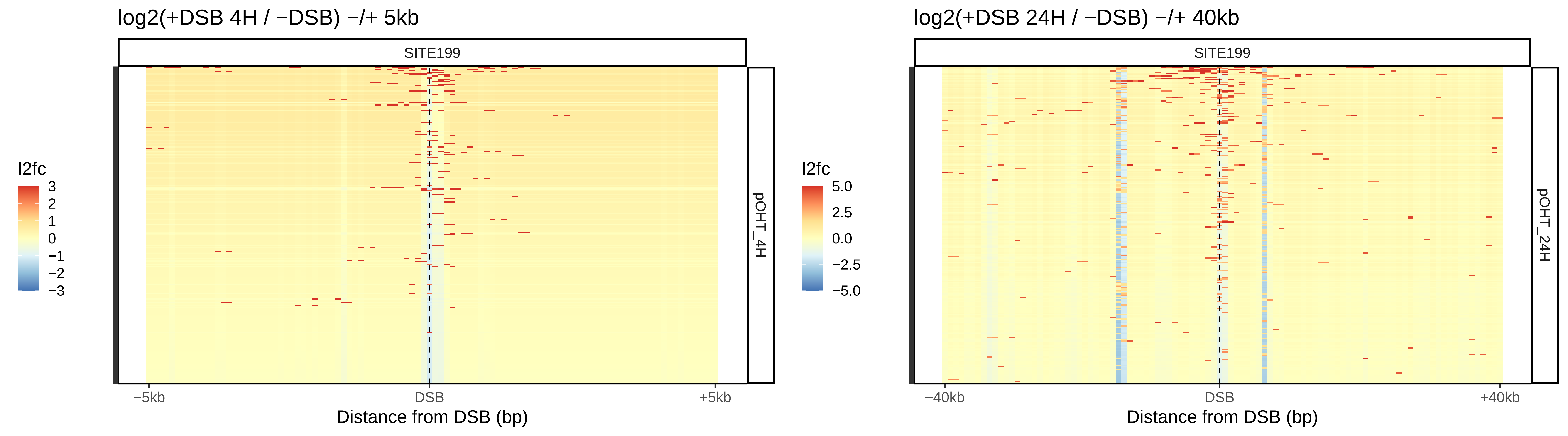

log2(+DSB 4H / −DSB) −/+ 5kb
SITE199
pOHT_4H
l2fc
3
2
1
0
−1
−2
−3
−5kb
DSB
+5kb
Distance from DSB (bp)
log2(+DSB 24H / −DSB) −/+ 40kb
SITE199
pOHT_24H
l2fc
5.0
2.5
0.0
−2.5
−5.0
−40kb
DSB
+40kb
Distance from DSB (bp)

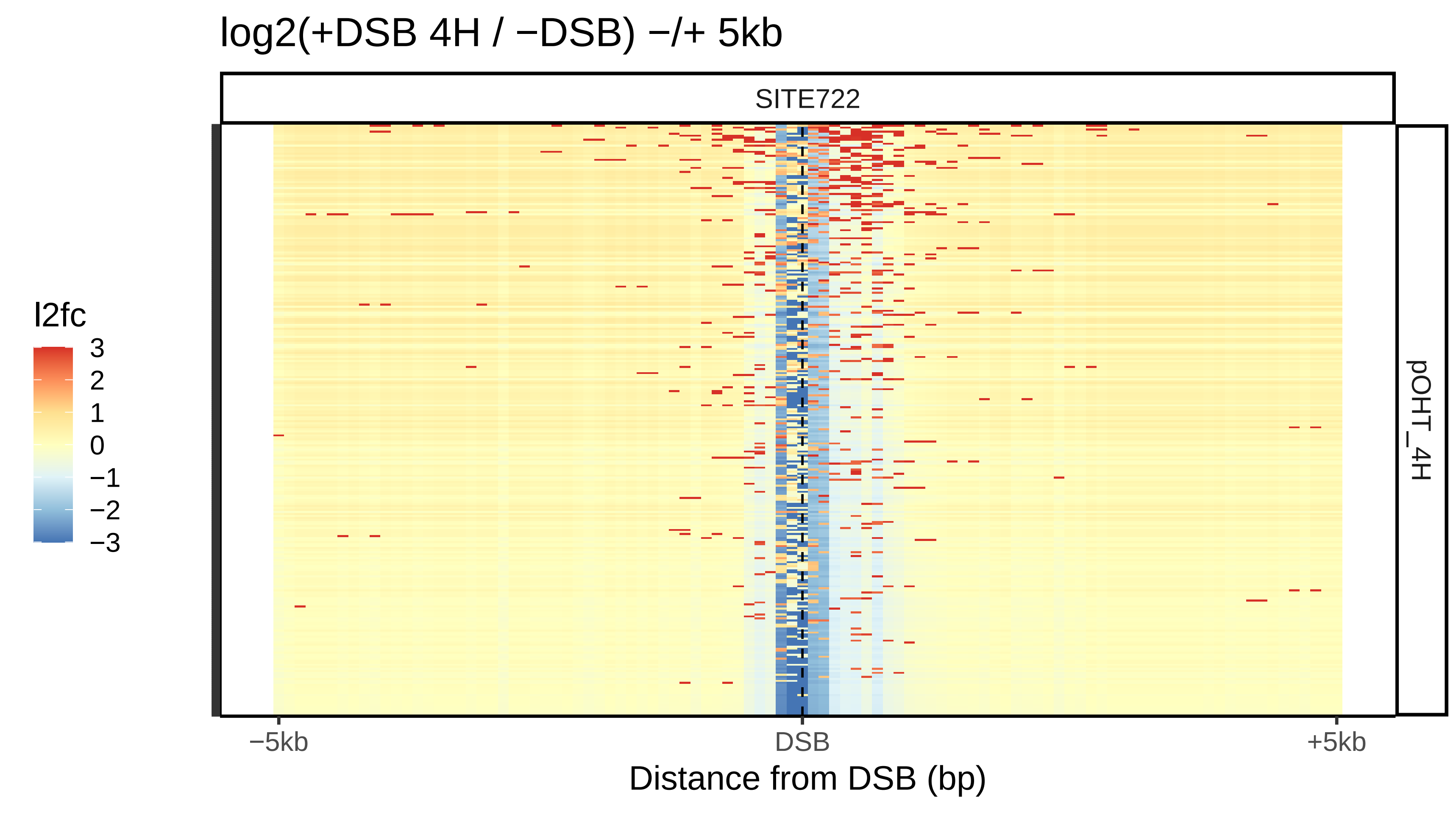

log2(+DSB 4H / −DSB) −/+ 5kb
SITE722
pOHT_4H
l2fc
3
2
1
0
−1
−2
−3
−5kb
DSB
+5kb
Distance from DSB (bp)


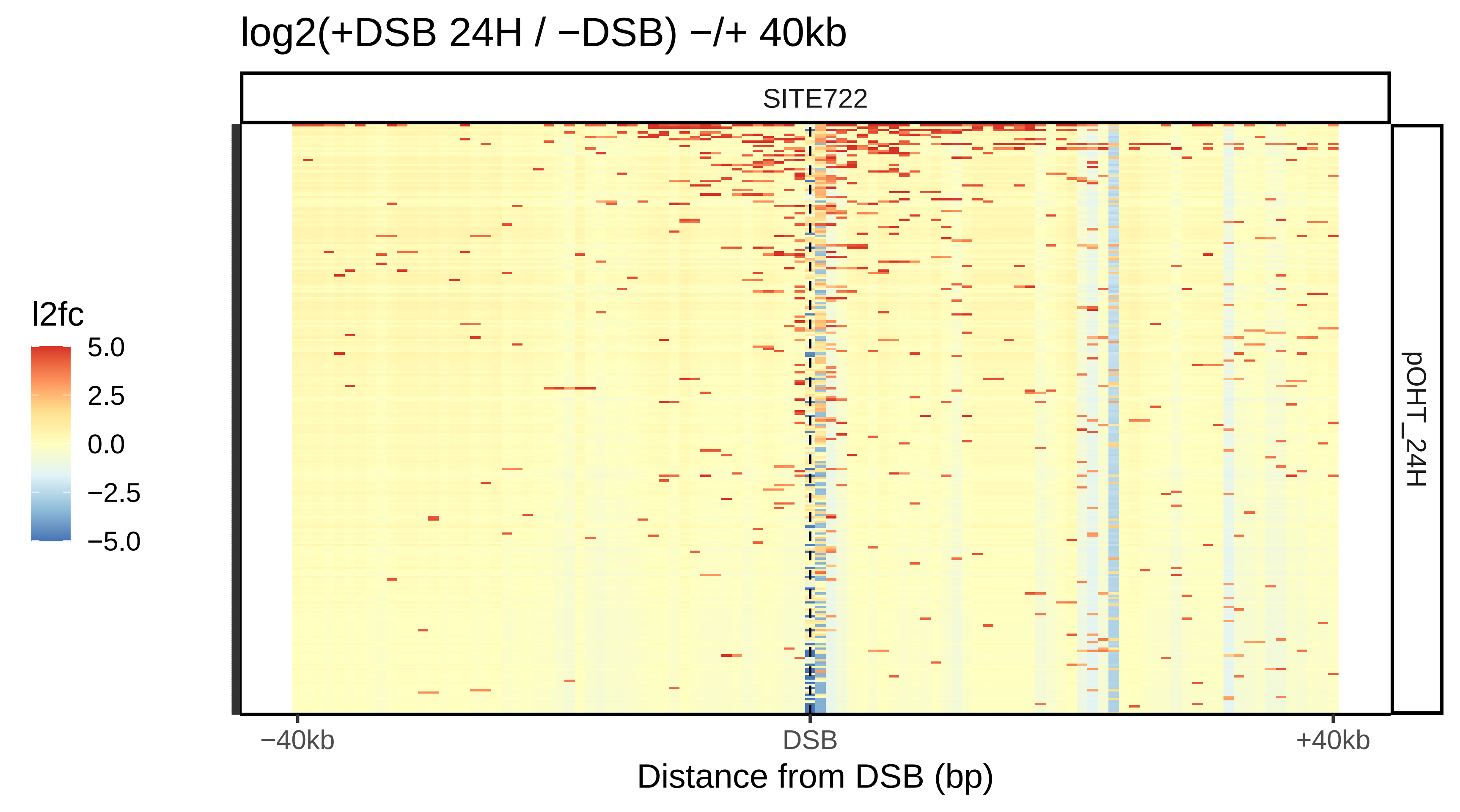

log2(+DSB 24H / −DSB) −/+ 40kb
SITE722
pOHT_24H
l2fc
5.0
2.5
0.0
−2.5
−5.0
−40kb
DSB
+40kb
Distance from DSB (bp)

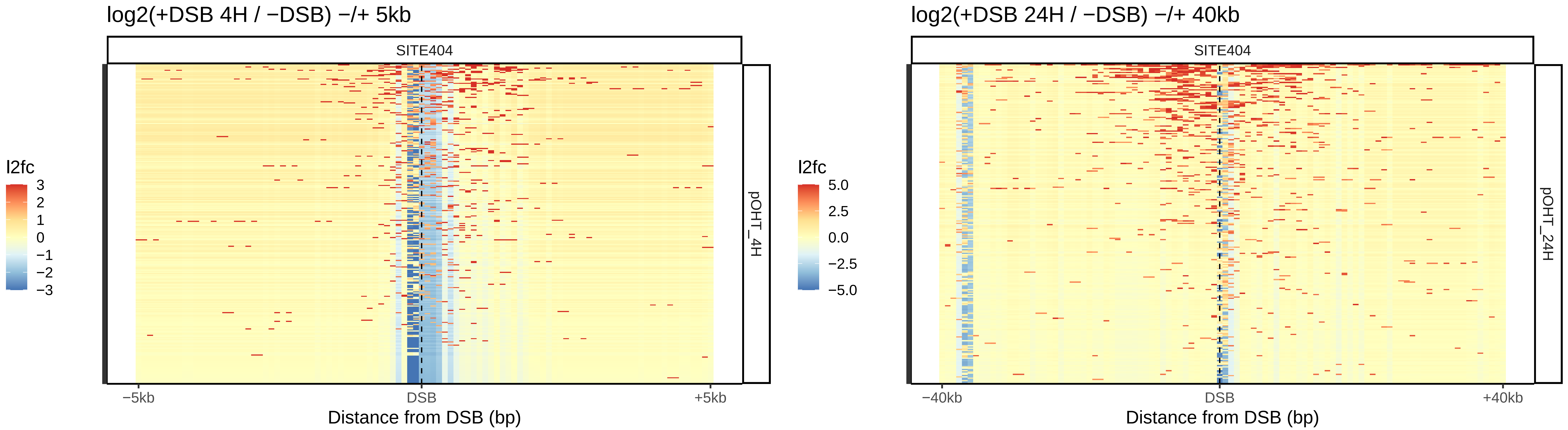

log2(+DSB 4H / −DSB) −/+ 5kb
SITE404
l2fc
3
2
1
0
−1
−2
−3
pOHT_4H
−5kb
DSB
+5kb
Distance from DSB (bp)
log2(+DSB 24H / −DSB) −/+ 40kb
SITE404
l2fc
5.0
2.5
0.0
−2.5
−5.0
pOHT_24H
−40kb
DSB
+40kb
Distance from DSB (bp)

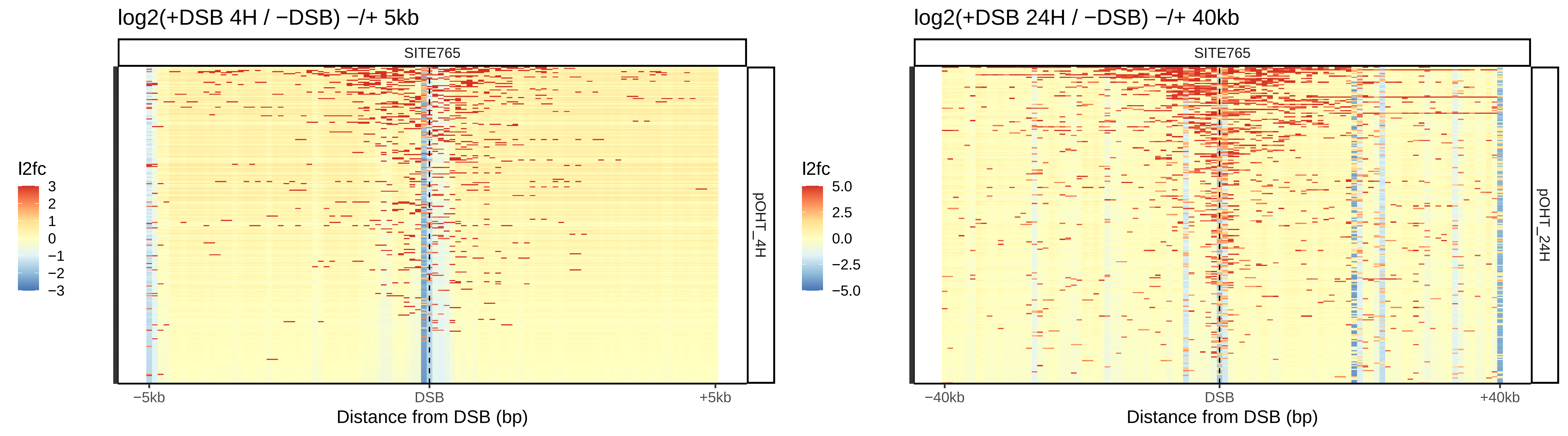
log2(+DSB 4H / −DSB) −/+ 5kb
SITE765
pOHT_4H
l2fc
3
2
1
0
−1
−2
−3
−5kb
DSB
+5kb
Distance from DSB (bp)
log2(+DSB 24H / −DSB) −/+ 40kb
SITE765
pOHT_24H
l2fc
5.0
2.5
0.0
−2.5
−5.0
−40kb
DSB
+40kb
Distance from DSB (bp)

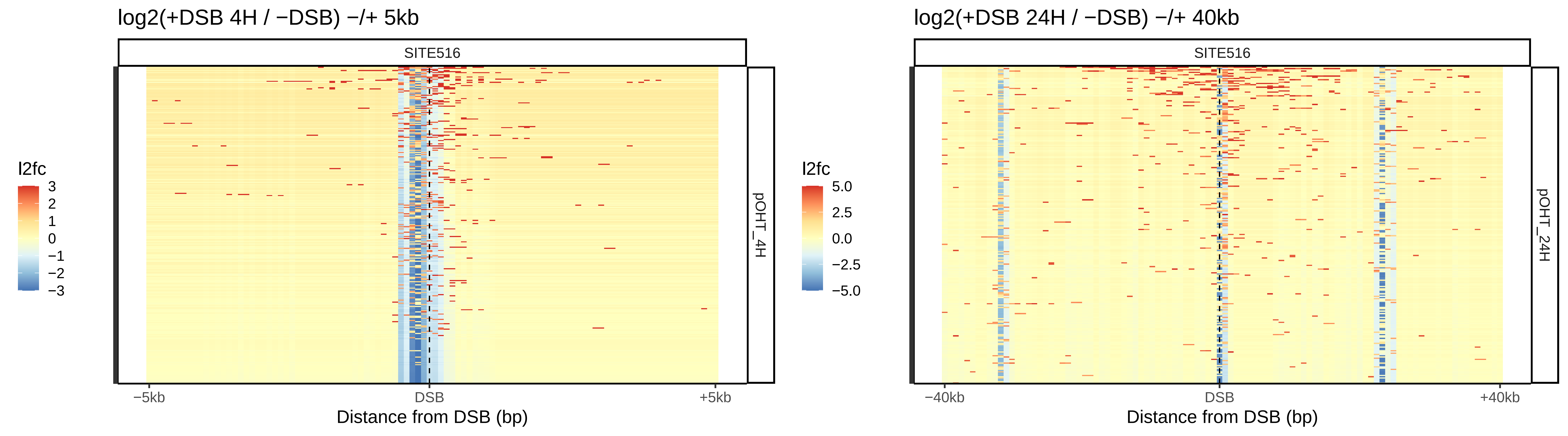

log2(+DSB 4H / −DSB) −/+ 5kb
SITE516
pOHT_4H
l2fc
3
2
1
0
−1
−2
−3
−5kb
DSB
+5kb
Distance from DSB (bp)
log2(+DSB 24H / −DSB) −/+ 40kb
SITE516
pOHT_24H
l2fc
5.0
2.5
0.0
−2.5
−5.0
−40kb
DSB
+40kb
Distance from DSB (bp)

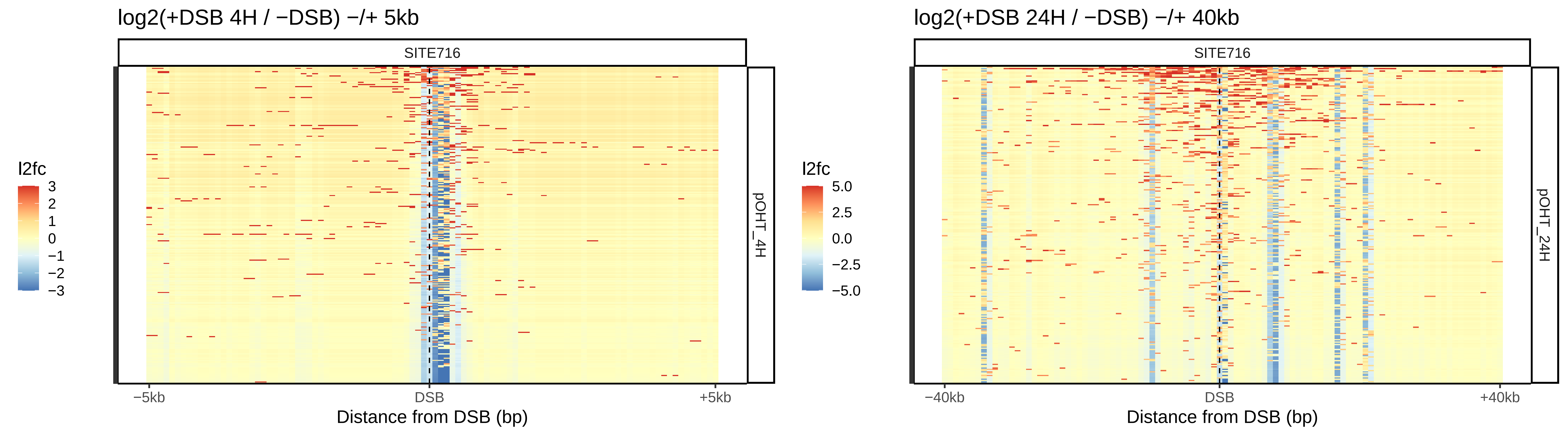

log2(+DSB 4H / −DSB) −/+ 5kb
SITE716
pOHT_4H
l2fc
3
2
1
0
−1
−2
−3
−5kb
DSB
+5kb
Distance from DSB (bp)
log2(+DSB 24H / −DSB) −/+ 40kb
SITE716
pOHT_24H
l2fc
5.0
2.5
0.0
−2.5
−5.0
−40kb
DSB
+40kb
Distance from DSB (bp)

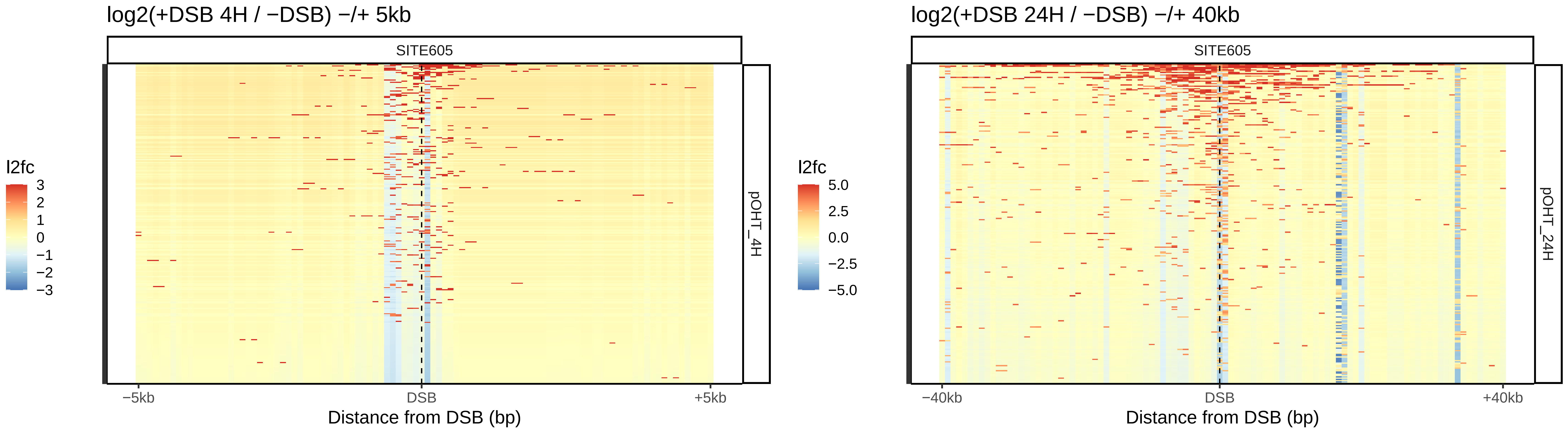
log2(+DSB 4H / −DSB) −/+ 5kb
SITE605
pOHT_4H
l2fc
3
2
1
0
−1
−2
−3
−5kb
DSB
+5kb
Distance from DSB (bp)
log2(+DSB 24H / −DSB) −/+ 40kb
SITE605
pOHT_24H
l2fc
5.0
2.5
0.0
−2.5
−5.0
−40kb
DSB
+40kb
Distance from DSB (bp)

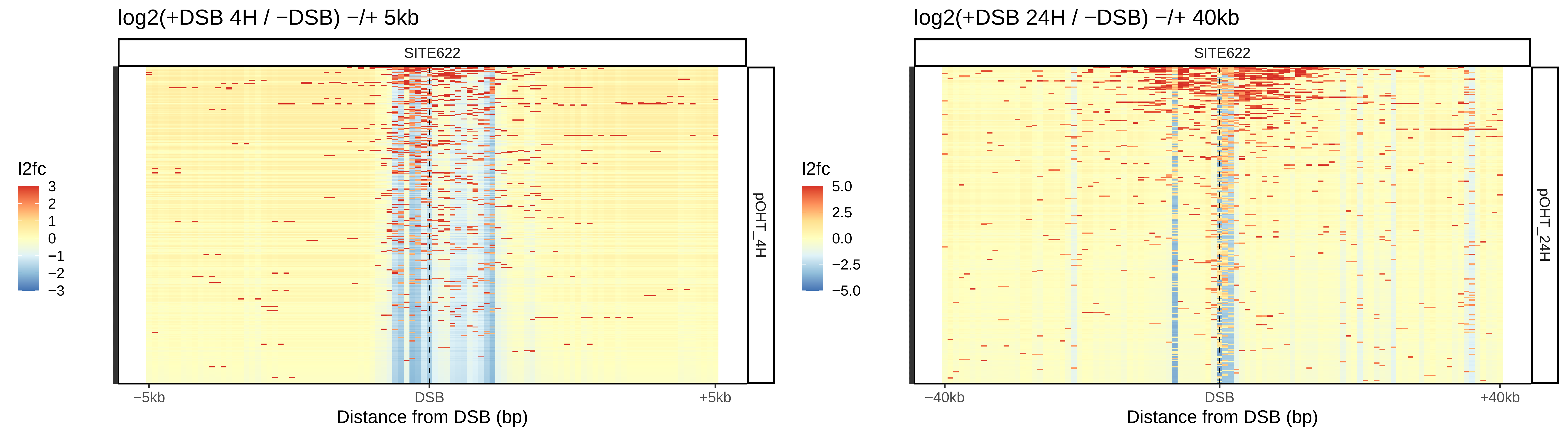

log2(+DSB 4H / −DSB) −/+ 5kb
SITE622
l2fc
3
2
1
0
−1
−2
−3
pOHT_4H
−5kb
DSB
+5kb
Distance from DSB (bp)
log2(+DSB 24H / −DSB) −/+ 40kb
SITE622
l2fc
5.0
2.5
0.0
−2.5
−5.0
pOHT_24H
−40kb
DSB
+40kb
Distance from DSB (bp)

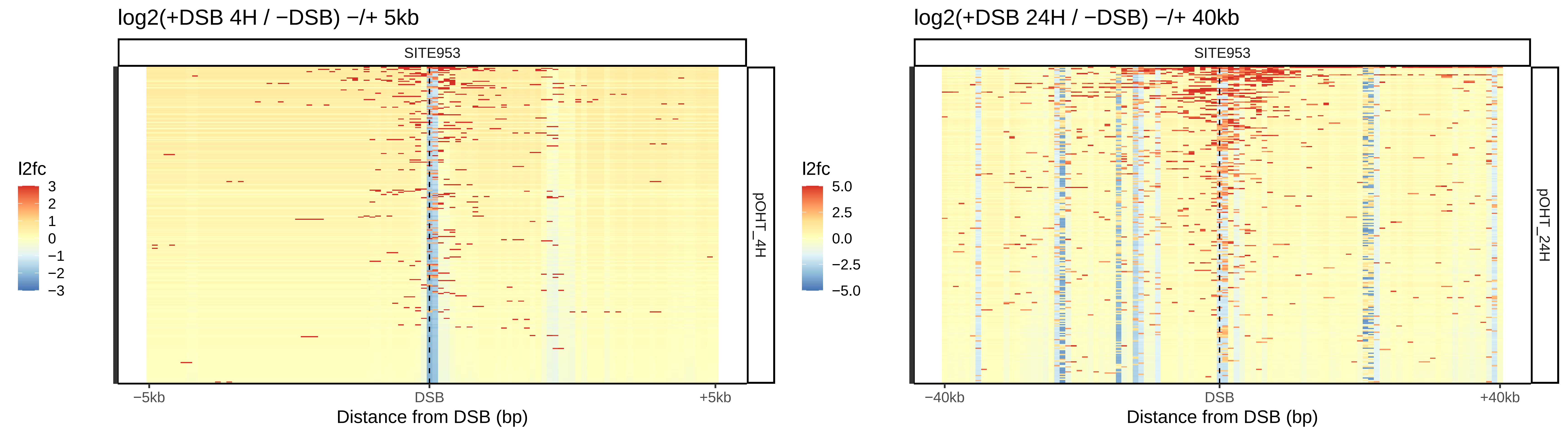
log2(+DSB 4H / −DSB) −/+ 5kb
SITE953
l2fc
3
2
1
0
−1
−2
−3
pOHT_4H
−5kb
DSB
+5kb
Distance from DSB (bp)
log2(+DSB 24H / −DSB) −/+ 40kb
SITE953
l2fc
5.0
2.5
0.0
−2.5
−5.0
pOHT_24H
−40kb
DSB
+40kb
Distance from DSB (bp)

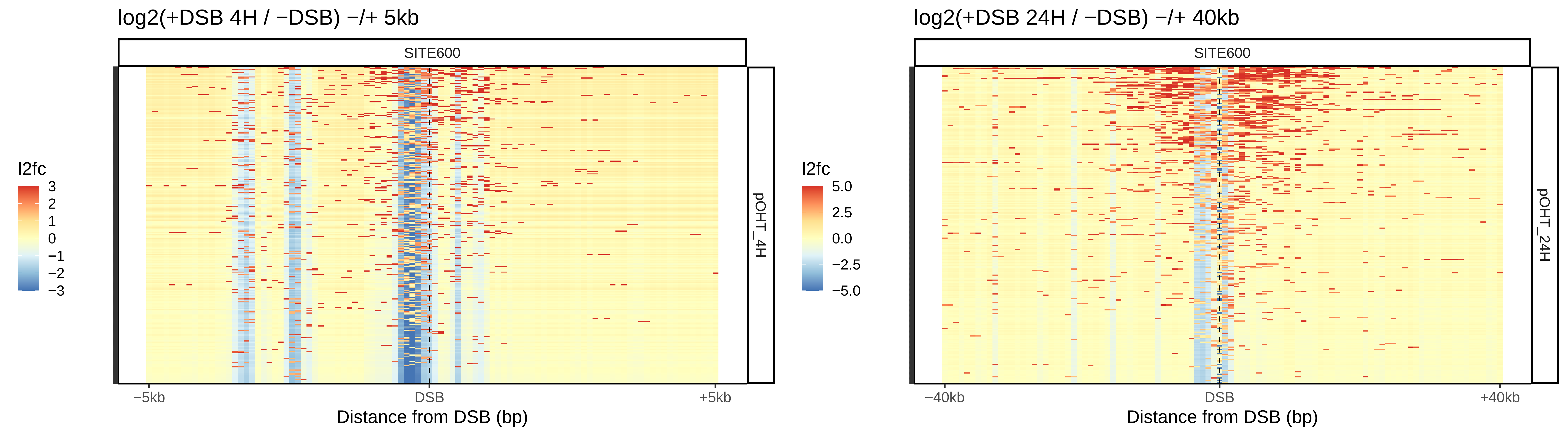
log2(+DSB 4H / −DSB) −/+ 5kb
SITE600
l2fc
3
2
1
0
−1
−2
−3
pOHT_4H
−5kb
DSB
+5kb
Distance from DSB (bp)
log2(+DSB 24H / −DSB) −/+ 40kb
SITE600
l2fc
5.0
2.5
0.0
−2.5
−5.0
pOHT_24H
−40kb
DSB
+40kb
Distance from DSB (bp)

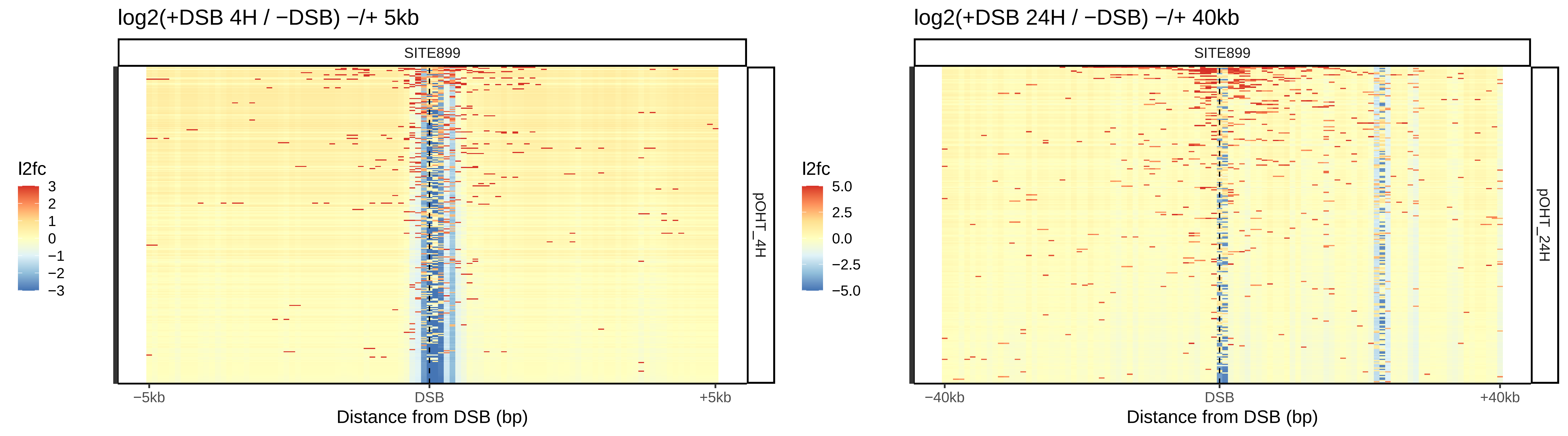

log2(+DSB 4H / −DSB) −/+ 5kb
SITE899
l2fc
3
2
1
0
−1
−2
−3
pOHT_4H
−5kb
DSB
+5kb
Distance from DSB (bp)
log2(+DSB 24H / −DSB) −/+ 40kb
SITE899
l2fc
5.0
2.5
0.0
−2.5
−5.0
pOHT_24H
−40kb
DSB
+40kb
Distance from DSB (bp)

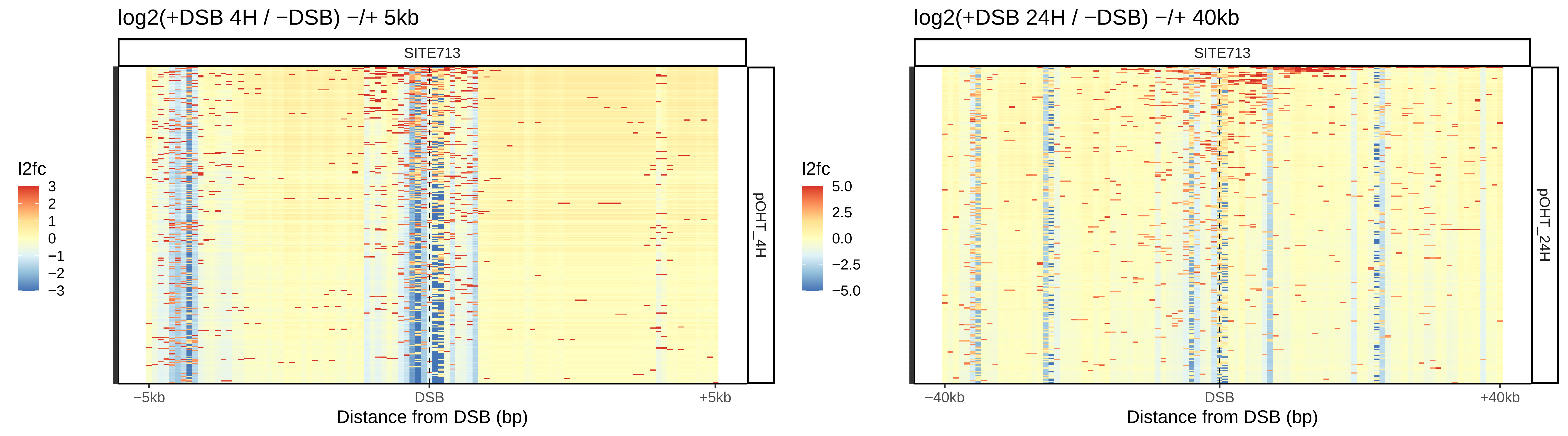
log2(+DSB 4H / −DSB) −/+ 5kb
SITE713
pOHT_4H
l2fc
3
2
1
0
−1
−2
−3
−5kb
DSB
+5kb
Distance from DSB (bp)
log2(+DSB 24H / −DSB) −/+ 40kb
SITE713
pOHT_24H
l2fc
5.0
2.5
0.0
−2.5
−5.0
−40kb
DSB
+40kb
Distance from DSB (bp)

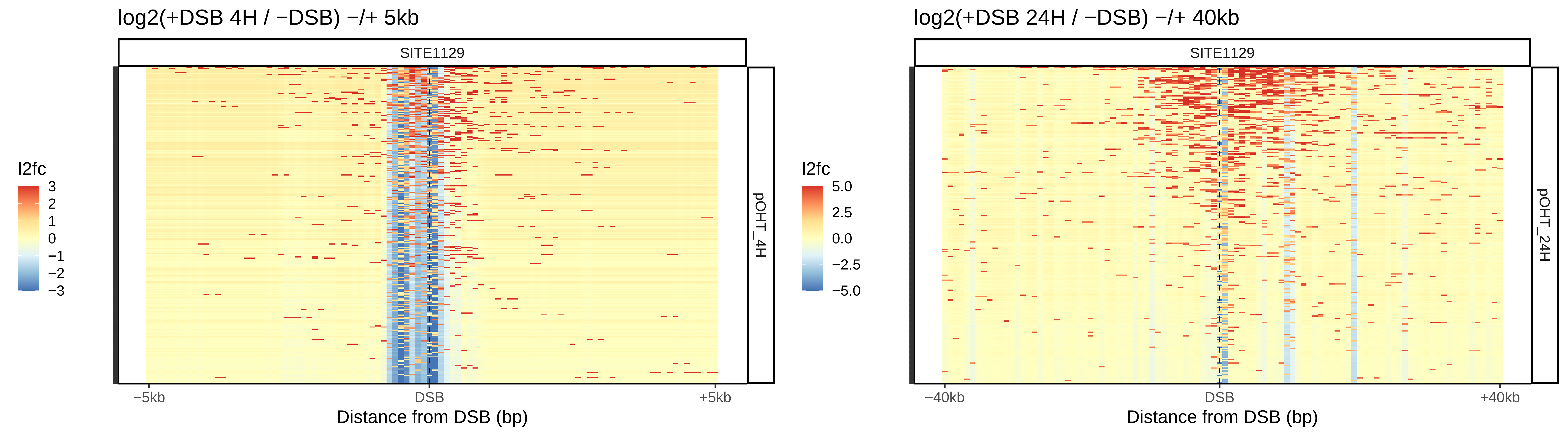
log2(+DSB 4H / −DSB) −/+ 5kb
SITE1129
l2fc
3
2
1
0
−1
−2
−3
pOHT_4H
−5kb
DSB
+5kb
Distance from DSB (bp)
log2(+DSB 24H / −DSB) −/+ 40kb
SITE1129
l2fc
5.0
2.5
0.0
−2.5
−5.0
pOHT_24H
−40kb
DSB
+40kb
Distance from DSB (bp)

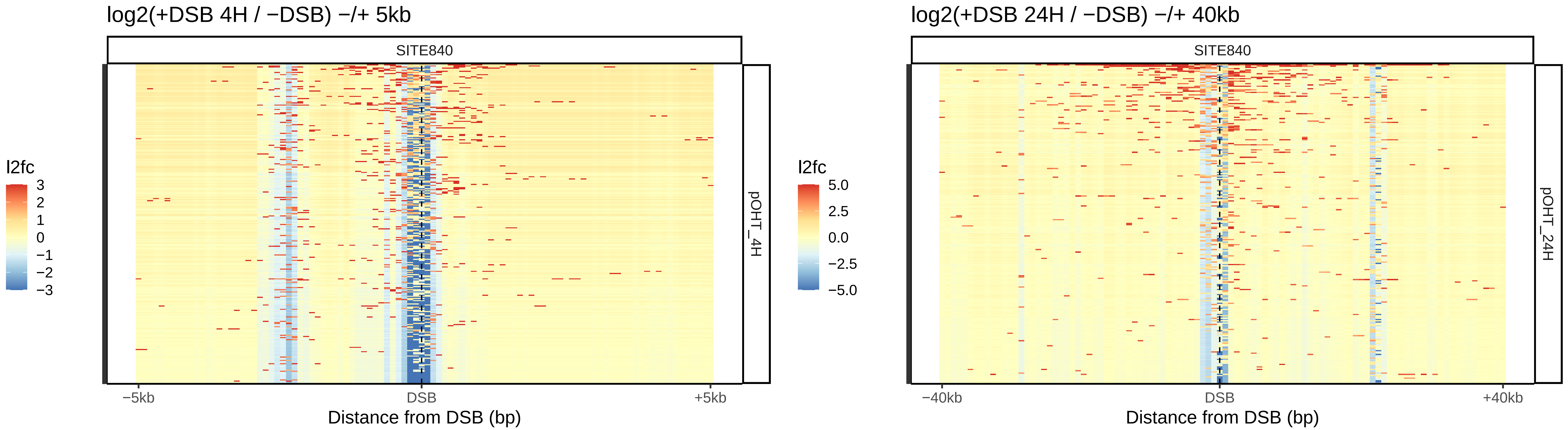
log2(+DSB 4H / −DSB) −/+ 5kb
SITE840
l2fc
3
2
1
0
−1
−2
−3
pOHT_4H
−5kb
DSB
+5kb
Distance from DSB (bp)
log2(+DSB 24H / −DSB) −/+ 40kb
SITE840
l2fc
5.0
2.5
0.0
−2.5
−5.0
pOHT_24H
−40kb
DSB
+40kb
Distance from DSB (bp)

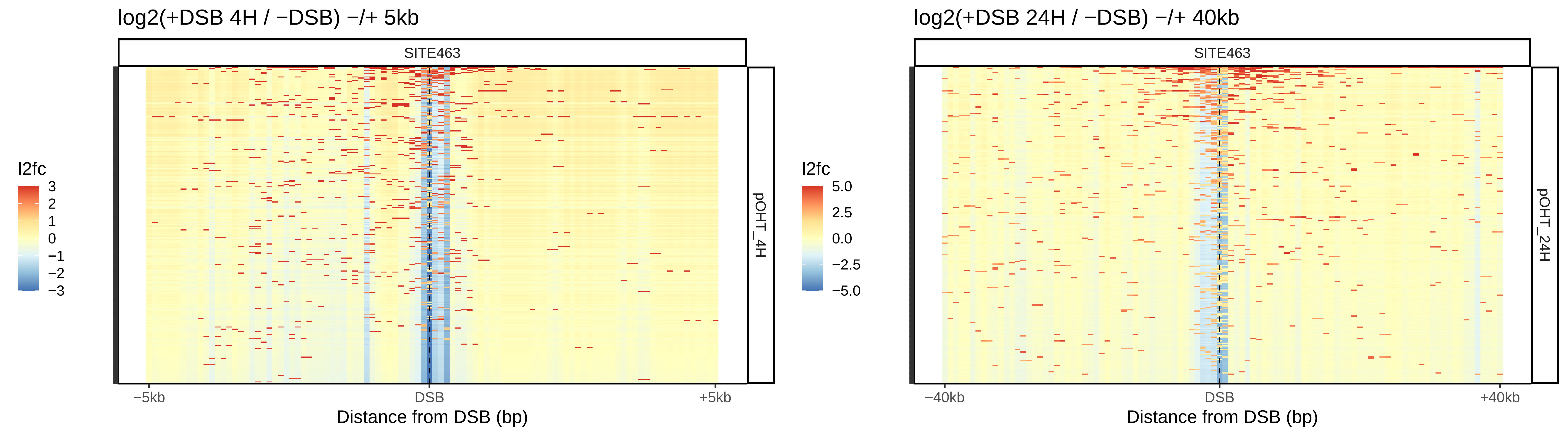
log2(+DSB 4H / −DSB) −/+ 5kb
SITE463
pOHT_4H
l2fc
3
2
1
0
−1
−2
−3
−5kb
DSB
+5kb
Distance from DSB (bp)
log2(+DSB 24H / −DSB) −/+ 40kb
SITE463
pOHT_24H
l2fc
5.0
2.5
0.0
−2.5
−5.0
−40kb
DSB
+40kb
Distance from DSB (bp)

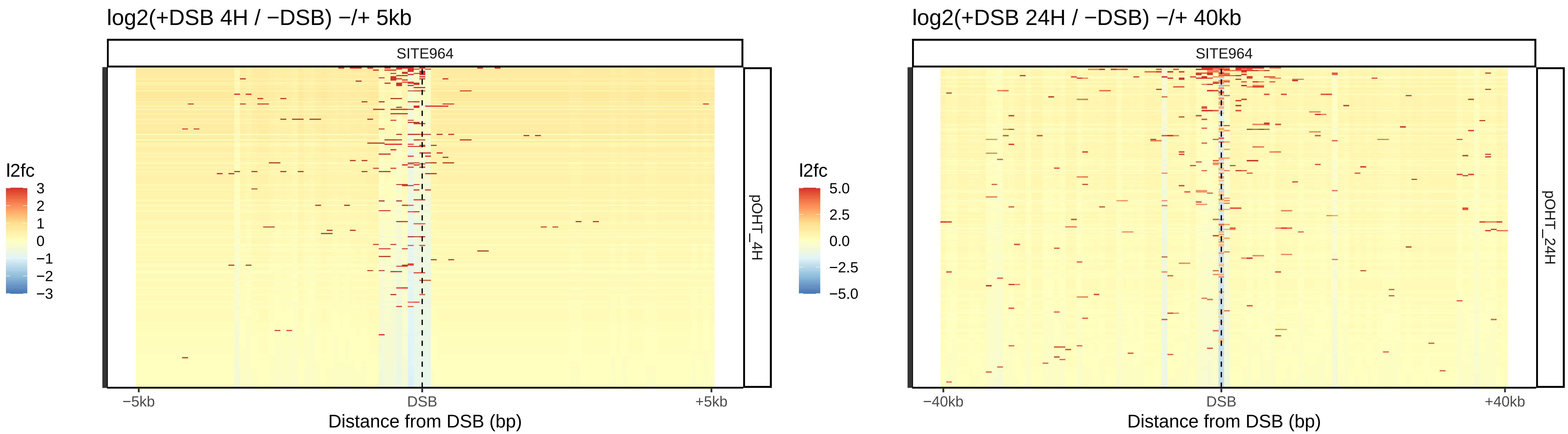

log2(+DSB 4H / −DSB) −/+ 5kb
SITE964
pOHT_4H
l2fc
3
2
1
0
−1
−2
−3
−5kb
DSB
+5kb
Distance from DSB (bp)
log2(+DSB 24H / −DSB) −/+ 40kb
SITE964
pOHT_24H
l2fc
5.0
2.5
0.0
−2.5
−5.0
−40kb
DSB
+40kb
Distance from DSB (bp)

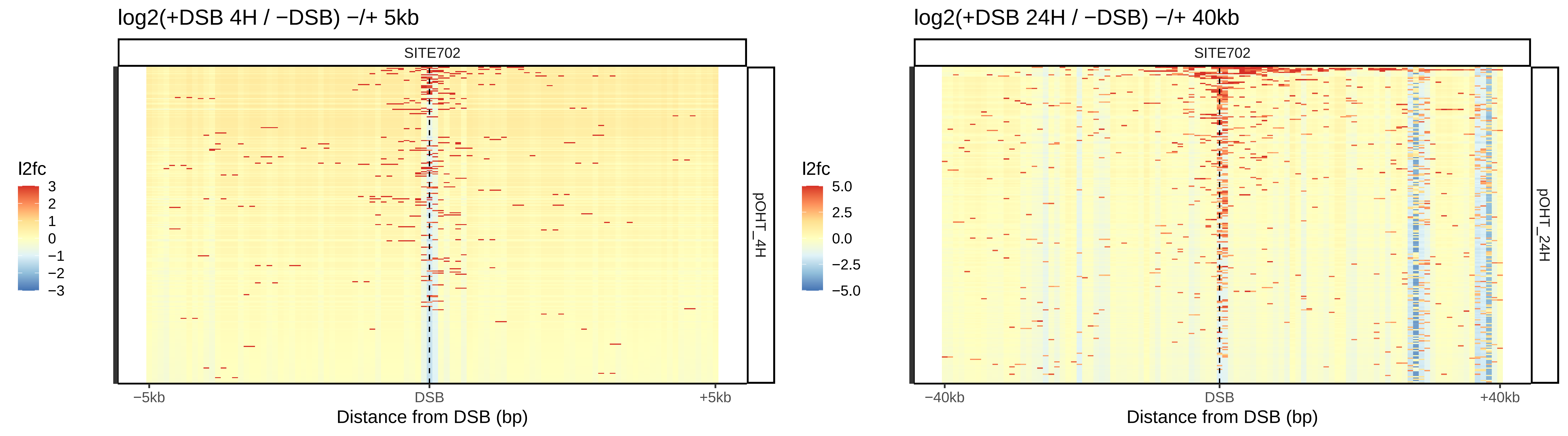
log2(+DSB 4H / −DSB) −/+ 5kb
SITE702
l2fc
3
2
1
0
−1
−2
−3
pOHT_4H
−5kb
DSB
+5kb
Distance from DSB (bp)
log2(+DSB 24H / −DSB) −/+ 40kb
SITE702
l2fc
5.0
2.5
0.0
−2.5
−5.0
pOHT_24H
−40kb
DSB
+40kb
Distance from DSB (bp)

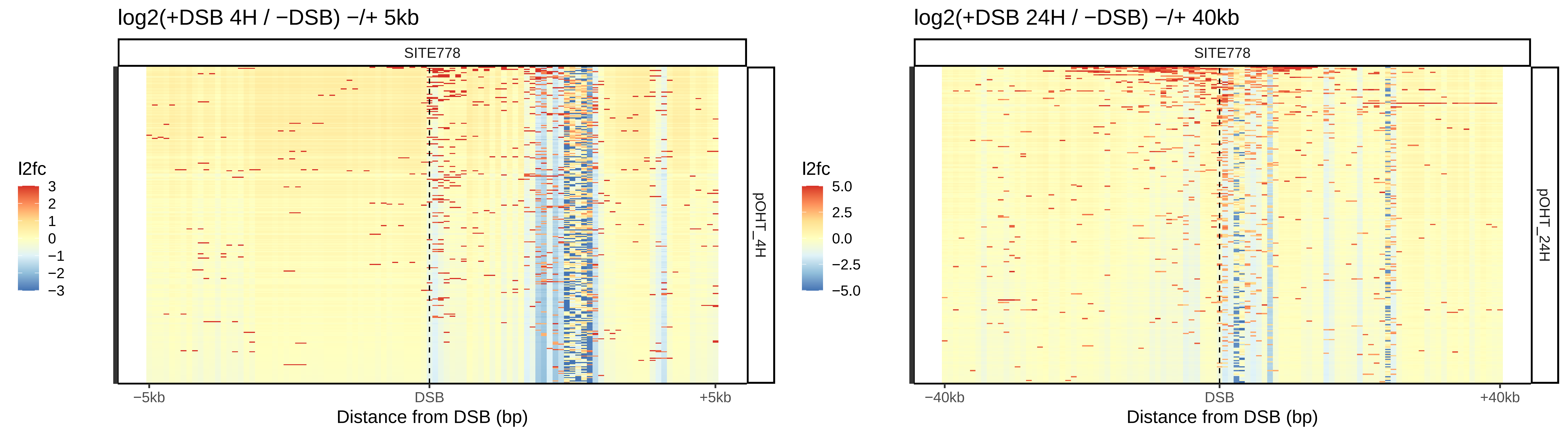

log2(+DSB 4H / −DSB) −/+ 5kb
SITE778
pOHT_4H
l2fc
3
2
1
0
−1
−2
−3
−5kb
DSB
+5kb
Distance from DSB (bp)
log2(+DSB 24H / −DSB) −/+ 40kb
SITE778
pOHT_24H
l2fc
5.0
2.5
0.0
−2.5
−5.0
−40kb
DSB
+40kb
Distance from DSB (bp)

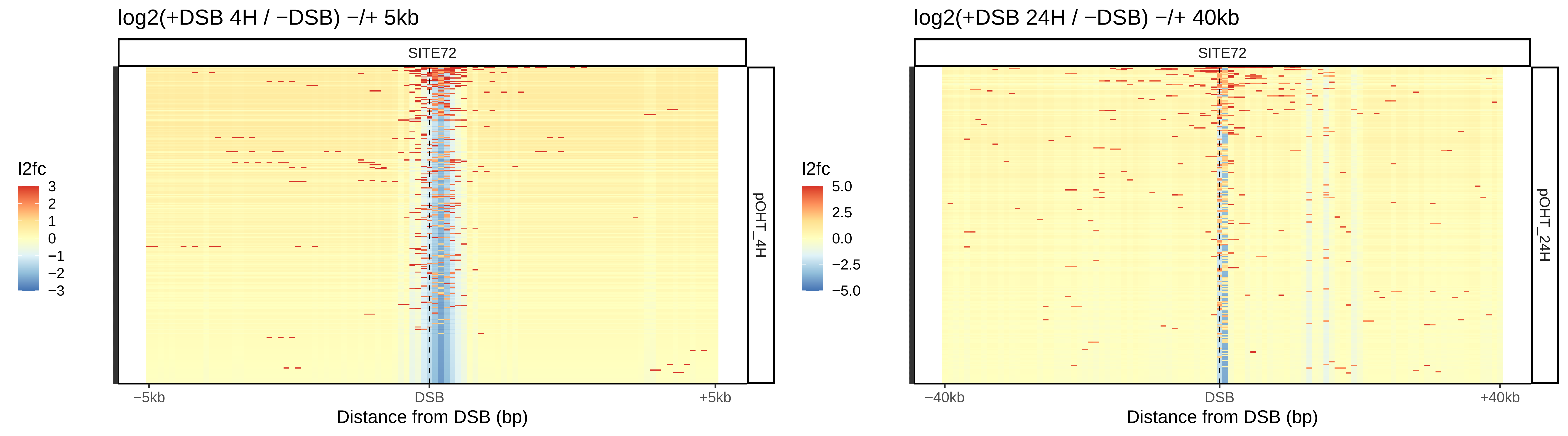

log2(+DSB 4H / −DSB) −/+ 5kb
SITE72
pOHT_4H
l2fc
3
2
1
0
−1
−2
−3
−5kb
DSB
+5kb
Distance from DSB (bp)
log2(+DSB 24H / −DSB) −/+ 40kb
SITE72
pOHT_24H
l2fc
5.0
2.5
0.0
−2.5
−5.0
−40kb
DSB
+40kb
Distance from DSB (bp)

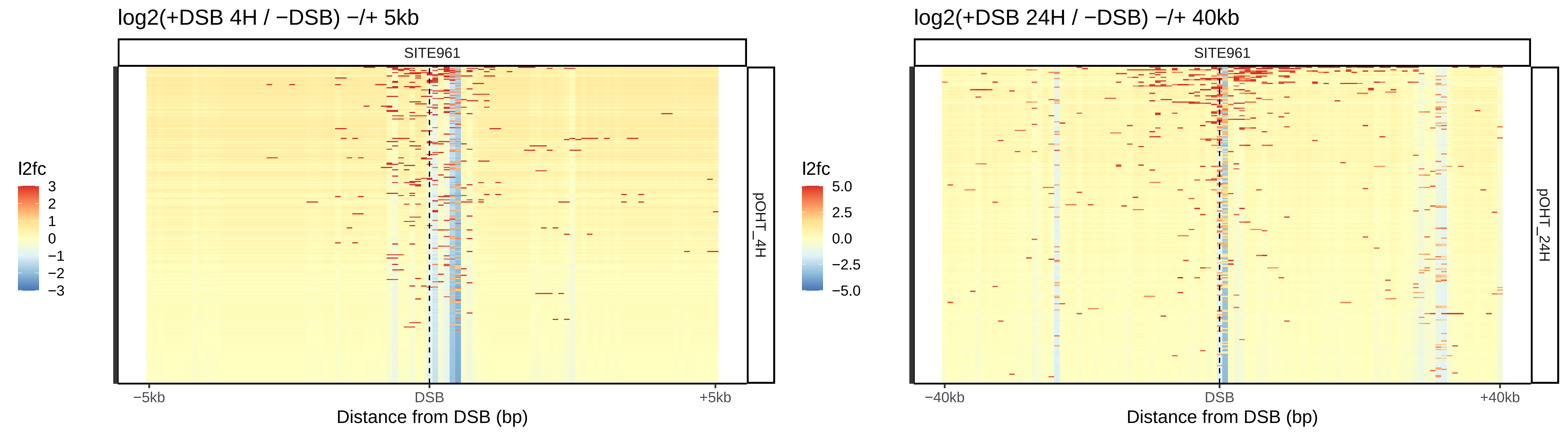
log2(+DSB 4H / −DSB) −/+ 5kb
SITE961
l2fc
3
2
1
0
−1
−2
−3
pOHT_4H
−5kb
DSB
+5kb
Distance from DSB (bp)
log2(+DSB 24H / −DSB) −/+ 40kb
SITE961
l2fc
5.0
2.5
0.0
−2.5
−5.0
pOHT_24H
−40kb
DSB
+40kb
Distance from DSB (bp)

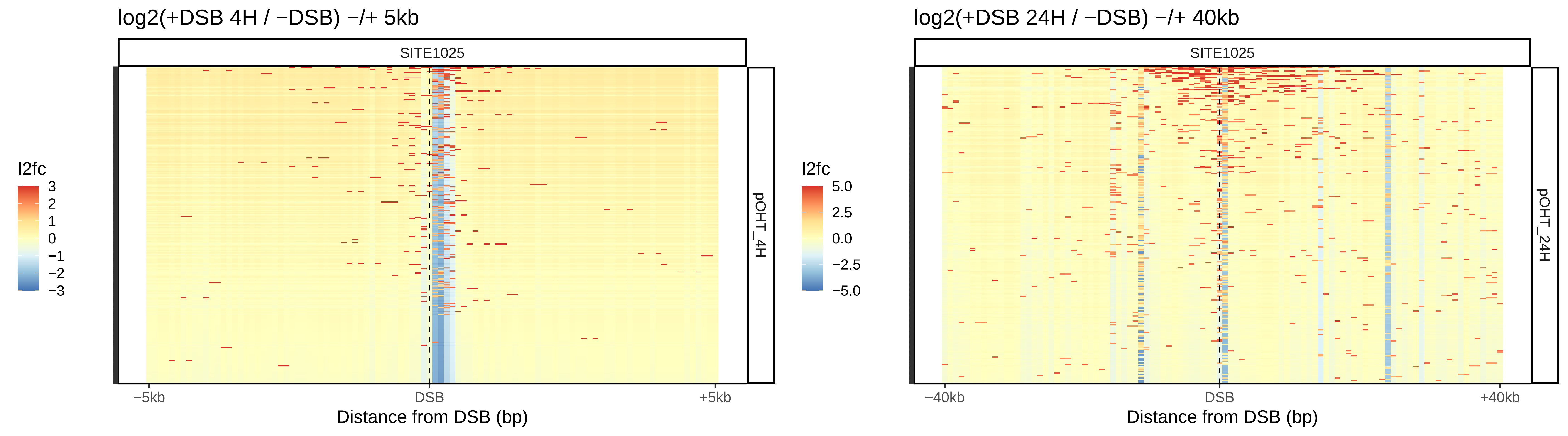

log2(+DSB 4H / −DSB) −/+ 5kb
SITE1025
pOHT_4H
l2fc
3
2
1
0
−1
−2
−3
−5kb
DSB
+5kb
Distance from DSB (bp)
log2(+DSB 24H / −DSB) −/+ 40kb
SITE1025
pOHT_24H
l2fc
5.0
2.5
0.0
−2.5
−5.0
−40kb
DSB
+40kb
Distance from DSB (bp)

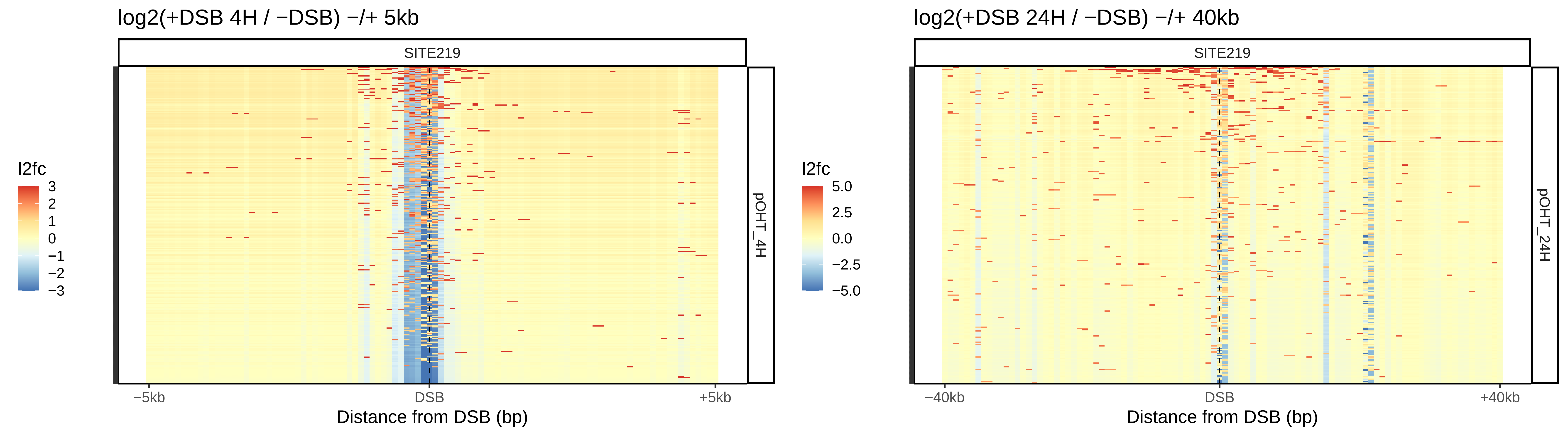
log2(+DSB 4H / −DSB) −/+ 5kb
SITE219
l2fc
3
2
1
0
−1
−2
−3
pOHT_4H
−5kb
DSB
+5kb
Distance from DSB (bp)
log2(+DSB 24H / −DSB) −/+ 40kb
SITE219
l2fc
5.0
2.5
0.0
−2.5
−5.0
pOHT_24H
−40kb
DSB
+40kb
Distance from DSB (bp)

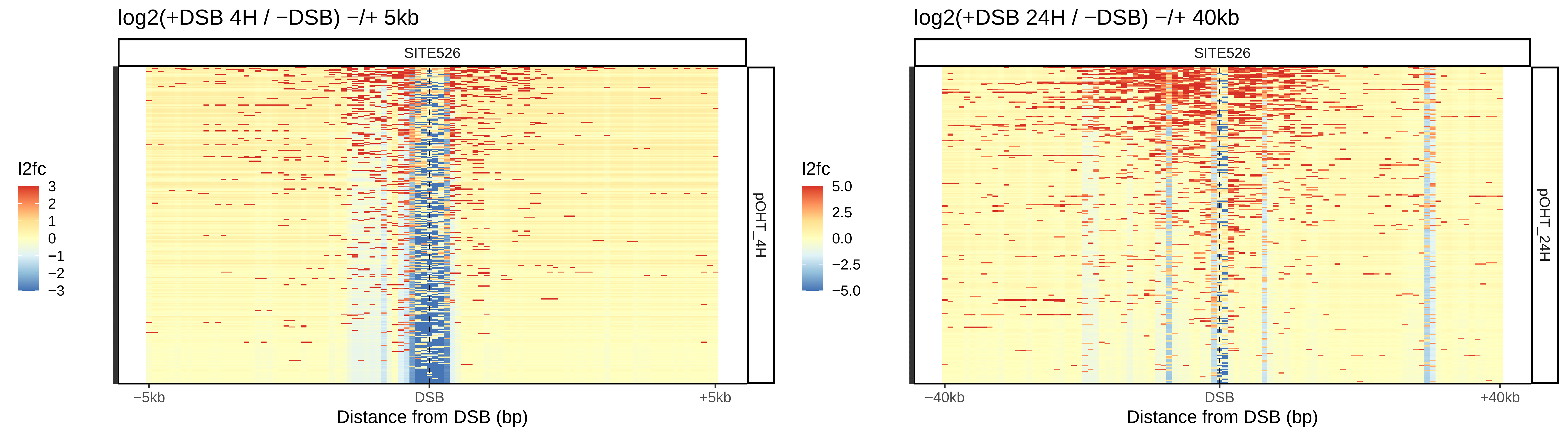

log2(+DSB 4H / −DSB) −/+ 5kb
SITE526
pOHT_4H
l2fc
3
2
1
0
−1
−2
−3
−5kb
DSB
+5kb
Distance from DSB (bp)
log2(+DSB 24H / −DSB) −/+ 40kb
SITE526
pOHT_24H
l2fc
5.0
2.5
0.0
−2.5
−5.0
−40kb
DSB
+40kb
Distance from DSB (bp)

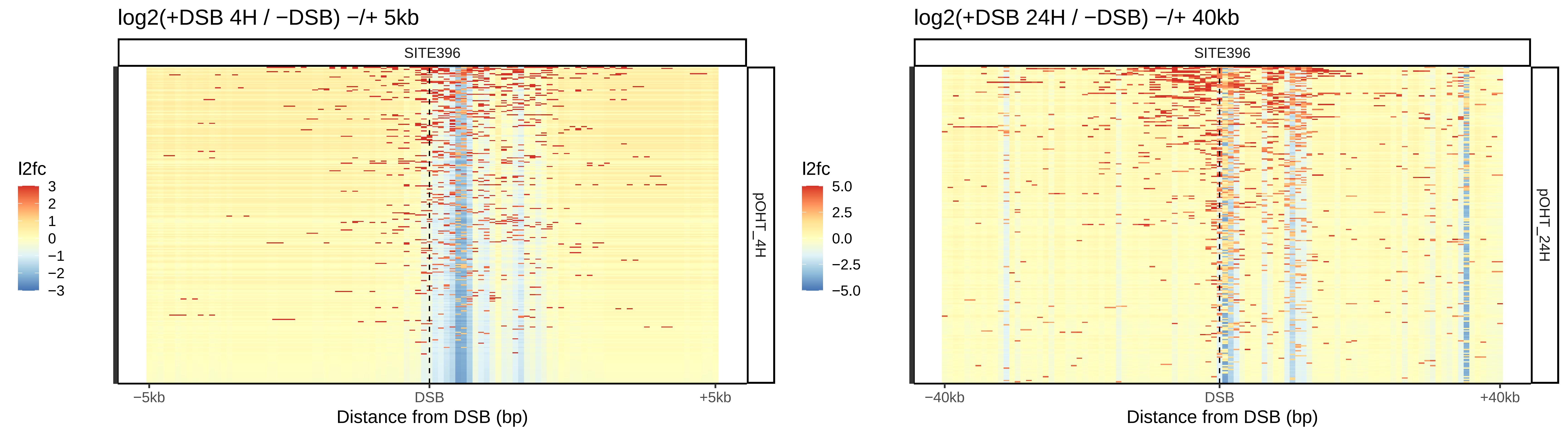

log2(+DSB 4H / −DSB) −/+ 5kb
SITE396
pOHT_4H
l2fc
3
2
1
0
−1
−2
−3
−5kb
DSB
+5kb
Distance from DSB (bp)
log2(+DSB 24H / −DSB) −/+ 40kb
SITE396
pOHT_24H
l2fc
5.0
2.5
0.0
−2.5
−5.0
−40kb
DSB
+40kb
Distance from DSB (bp)

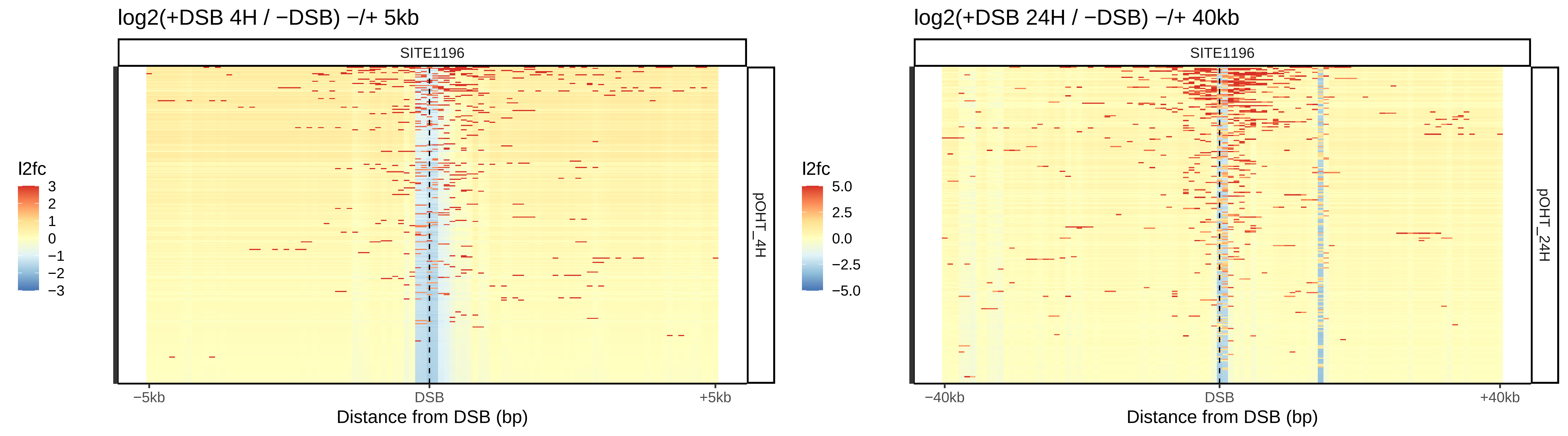
log2(+DSB 4H / −DSB) −/+ 5kb
SITE1196
l2fc
3
2
1
0
−1
−2
−3
pOHT_4H
−5kb
DSB
+5kb
Distance from DSB (bp)
log2(+DSB 24H / −DSB) −/+ 40kb
SITE1196
l2fc
5.0
2.5
0.0
−2.5
−5.0
pOHT_24H
−40kb
DSB
+40kb
Distance from DSB (bp)

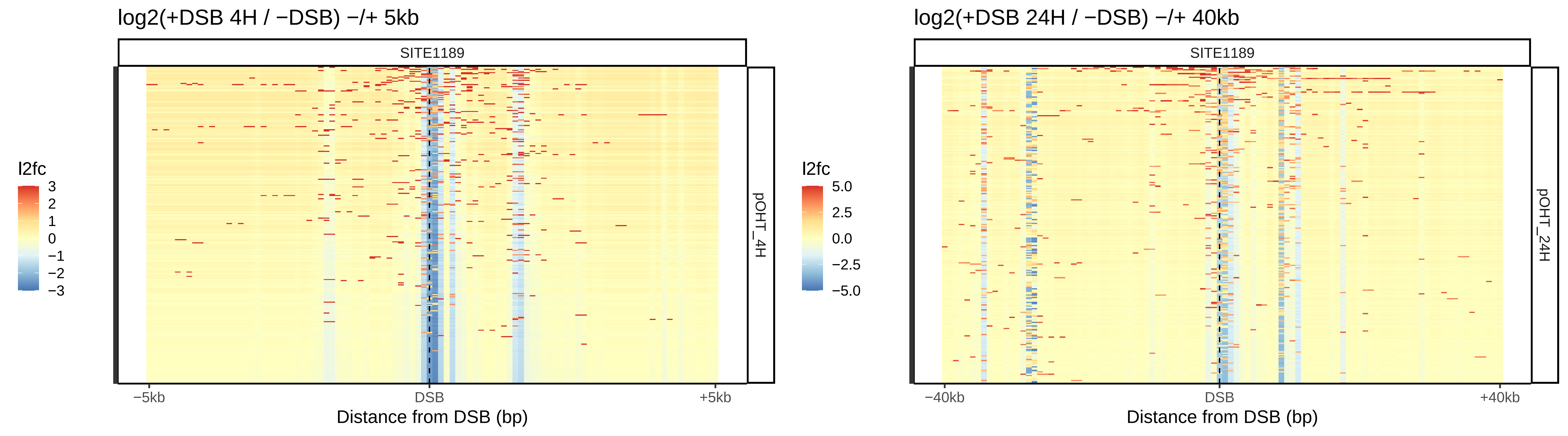
log2(+DSB 4H / −DSB) −/+ 5kb
SITE1189
l2fc
3
2
1
0
−1
−2
−3
pOHT_4H
−5kb
DSB
+5kb
Distance from DSB (bp)
log2(+DSB 24H / −DSB) −/+ 40kb
SITE1189
l2fc
5.0
2.5
0.0
−2.5
−5.0
pOHT_24H
−40kb
DSB
+40kb
Distance from DSB (bp)

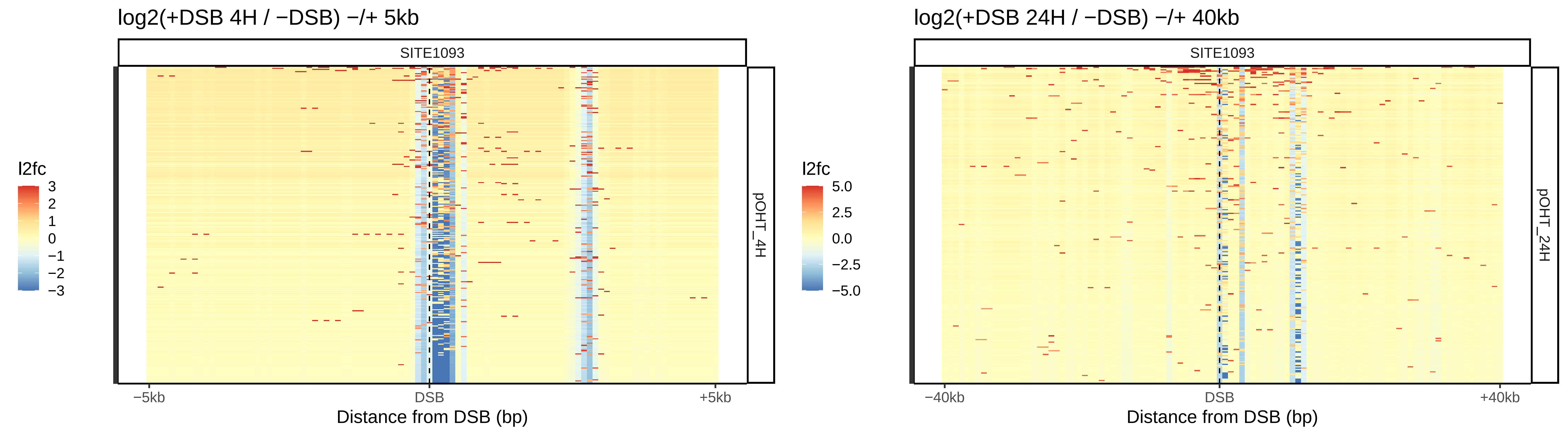

log2(+DSB 4H / −DSB) −/+ 5kb
SITE1093
pOHT_4H
l2fc
3
2
1
0
−1
−2
−3
−5kb
DSB
+5kb
Distance from DSB (bp)
log2(+DSB 24H / −DSB) −/+ 40kb
SITE1093
pOHT_24H
l2fc
5.0
2.5
0.0
−2.5
−5.0
−40kb
DSB
+40kb
Distance from DSB (bp)

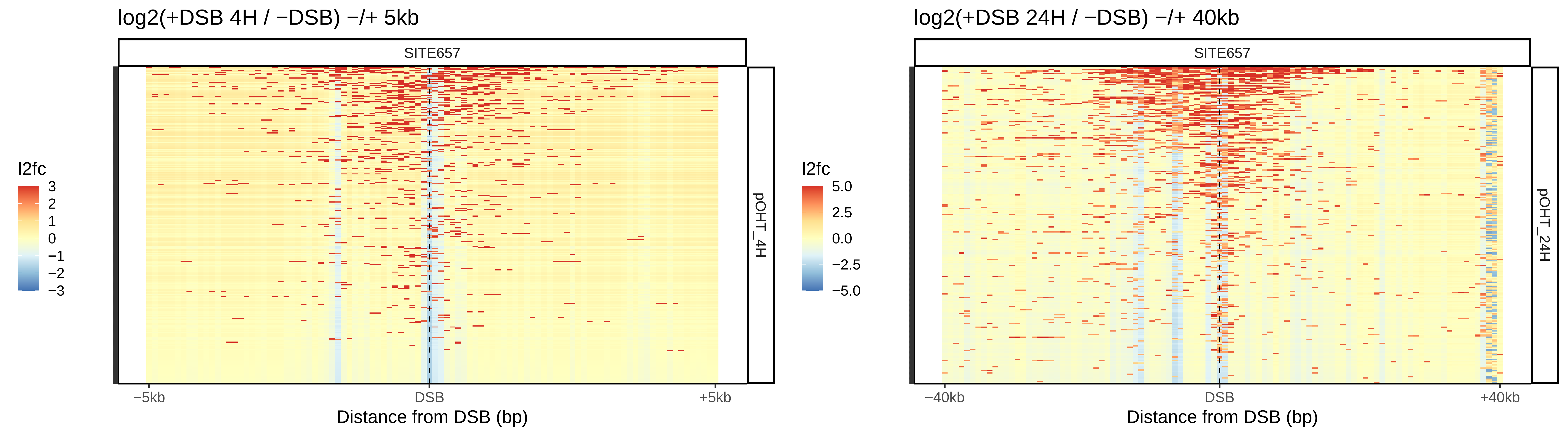
log2(+DSB 4H / −DSB) −/+ 5kb
SITE657
l2fc
3
2
1
0
−1
−2
−3
pOHT_4H
−5kb
DSB
+5kb
Distance from DSB (bp)
log2(+DSB 24H / −DSB) −/+ 40kb
SITE657
l2fc
5.0
2.5
0.0
−2.5
−5.0
pOHT_24H
−40kb
DSB
+40kb
Distance from DSB (bp)

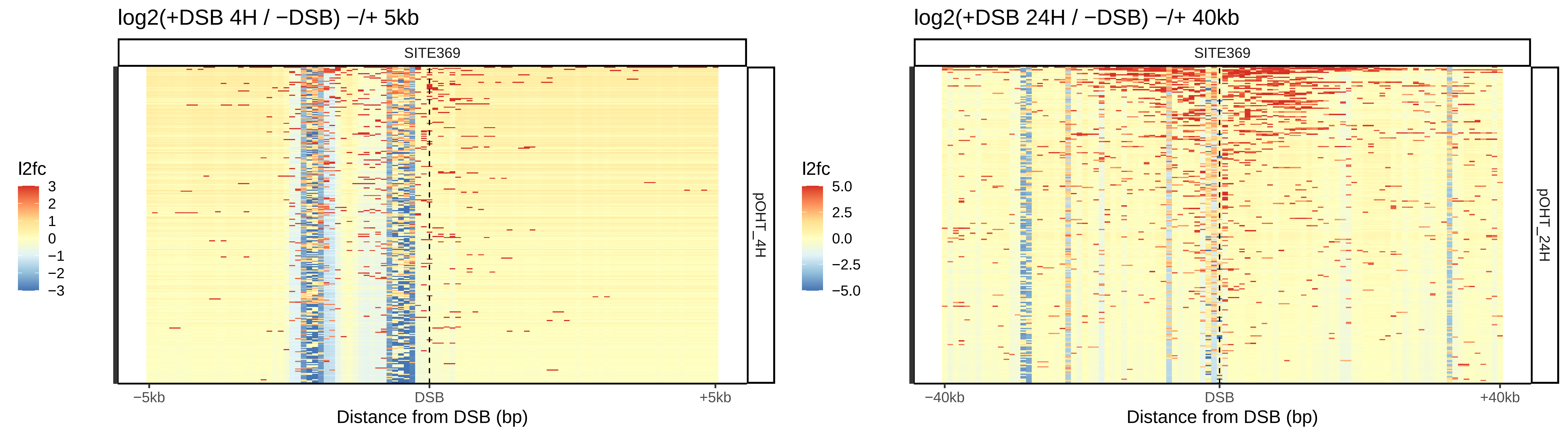

log2(+DSB 4H / −DSB) −/+ 5kb
SITE369
l2fc
3
2
1
0
−1
−2
−3
pOHT_4H
−5kb
DSB
+5kb
Distance from DSB (bp)
log2(+DSB 24H / −DSB) −/+ 40kb
SITE369
l2fc
5.0
2.5
0.0
−2.5
−5.0
pOHT_24H
−40kb
DSB
+40kb
Distance from DSB (bp)

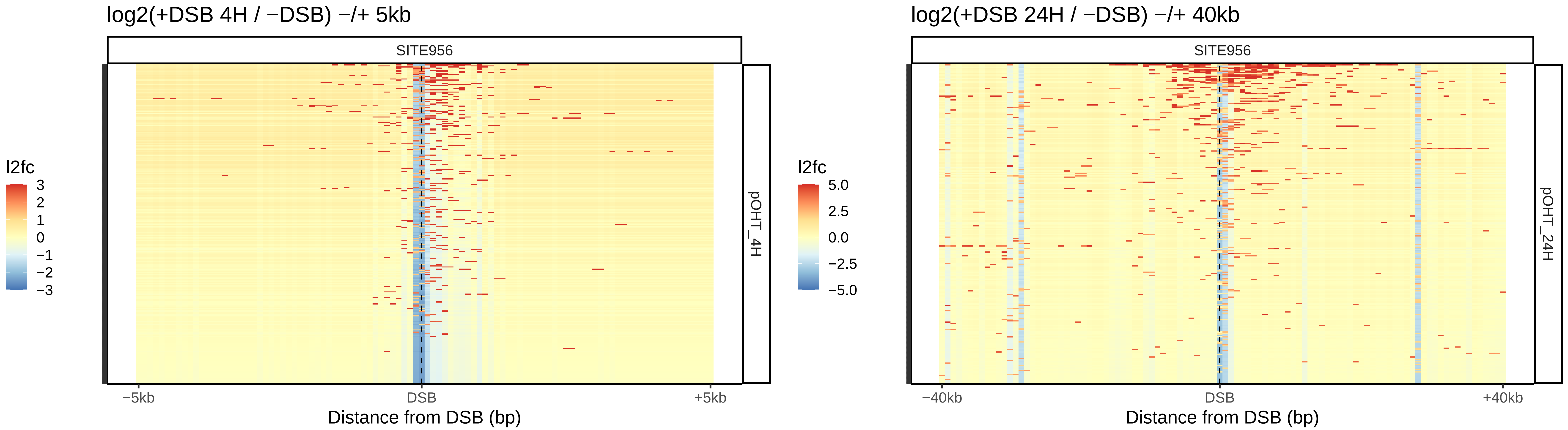
log2(+DSB 4H / −DSB) −/+ 5kb
SITE956
l2fc
3
2
1
0
−1
−2
−3
pOHT_4H
−5kb
DSB
+5kb
Distance from DSB (bp)
log2(+DSB 24H / −DSB) −/+ 40kb
SITE956
l2fc
5.0
2.5
0.0
−2.5
−5.0
pOHT_24H
−40kb
DSB
+40kb
Distance from DSB (bp)

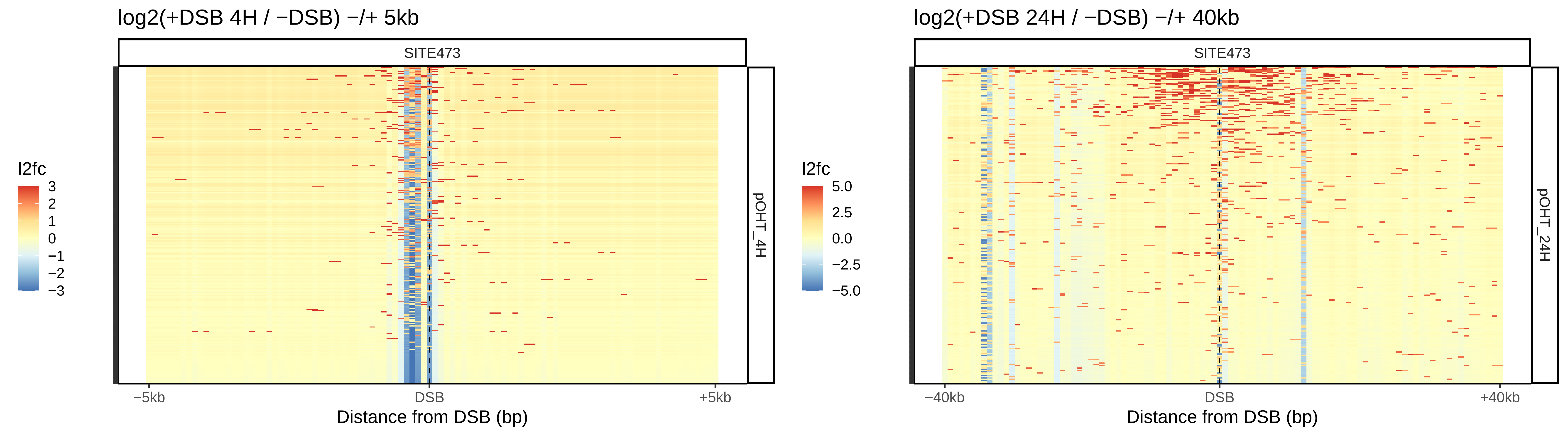
log2(+DSB 4H / −DSB) −/+ 5kb
SITE473
pOHT_4H
l2fc
3
2
1
0
−1
−2
−3
−5kb
DSB
+5kb
Distance from DSB (bp)
log2(+DSB 24H / −DSB) −/+ 40kb
SITE473
pOHT_24H
l2fc
5.0
2.5
0.0
−2.5
−5.0
−40kb
DSB
+40kb
Distance from DSB (bp)

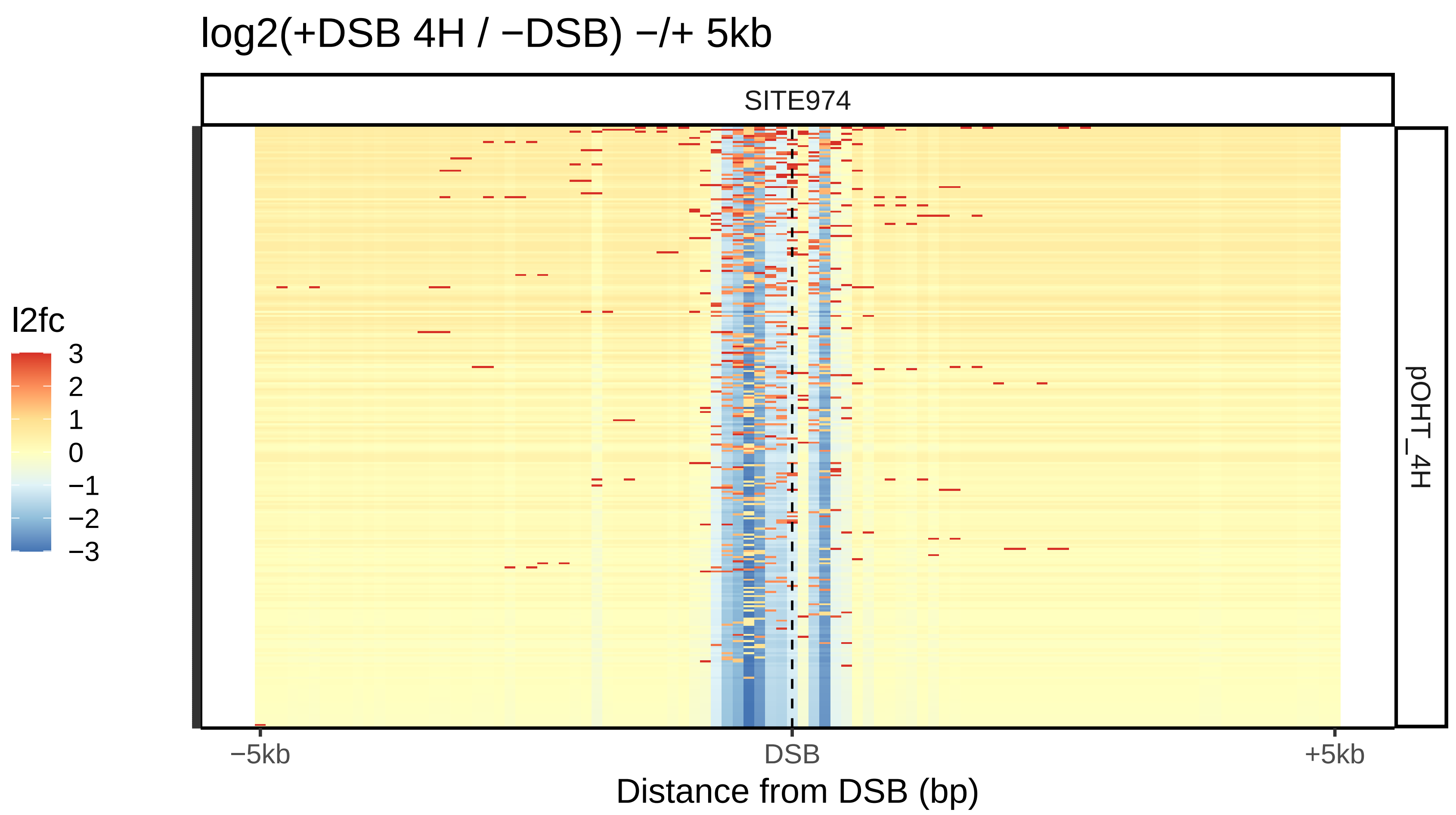
log2(+DSB 4H / −DSB) −/+ 5kb
SITE974
l2fc
3
2
1
0
−1
−2
−3
pOHT_4H
−5kb
DSB
+5kb
Distance from DSB (bp)
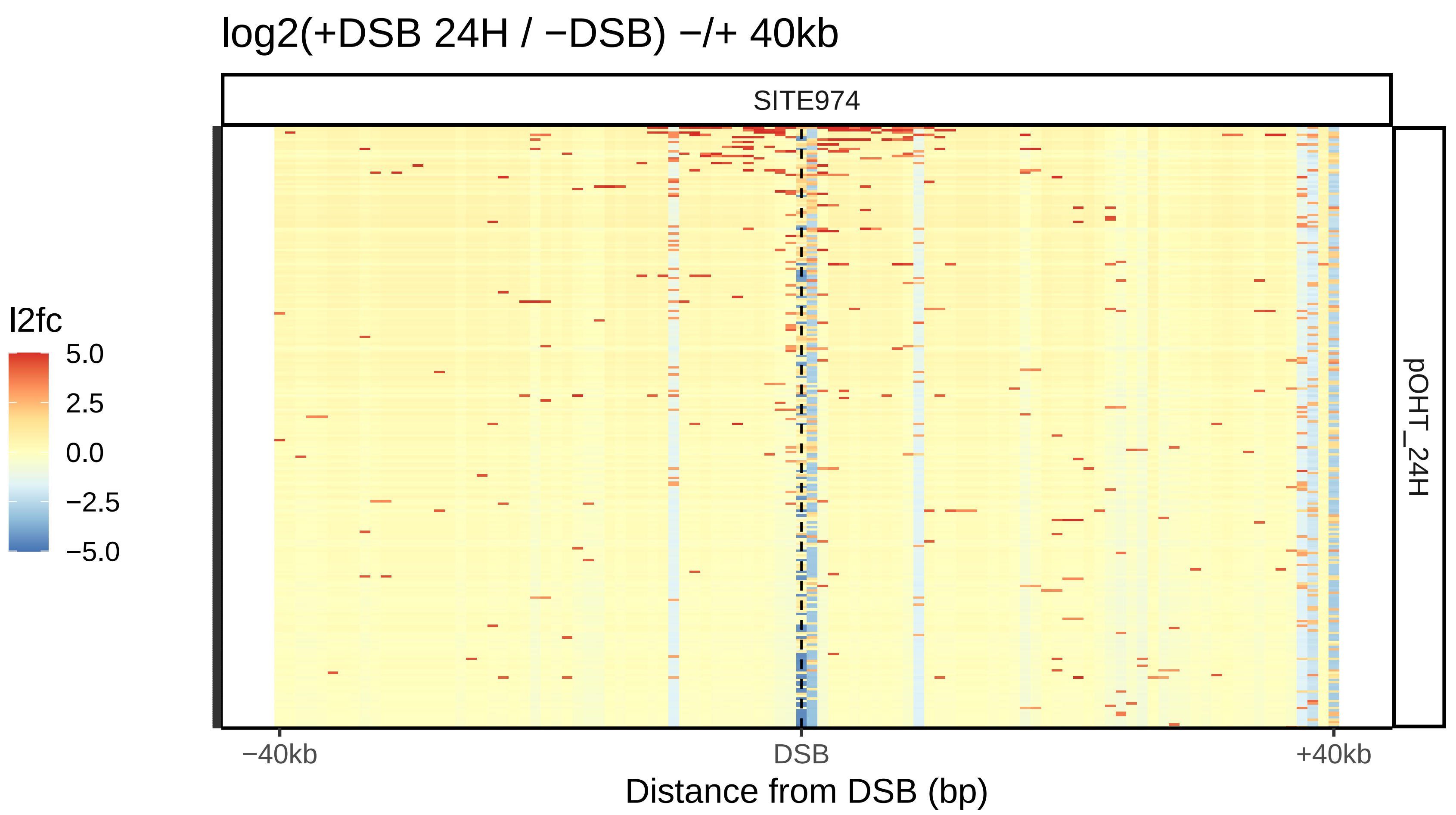
log2(+DSB 24H / −DSB) −/+ 40kb
SITE974
l2fc
5.0
2.5
0.0
−2.5
−5.0
pOHT_24H
−40kb
DSB
+40kb
Distance from DSB (bp)

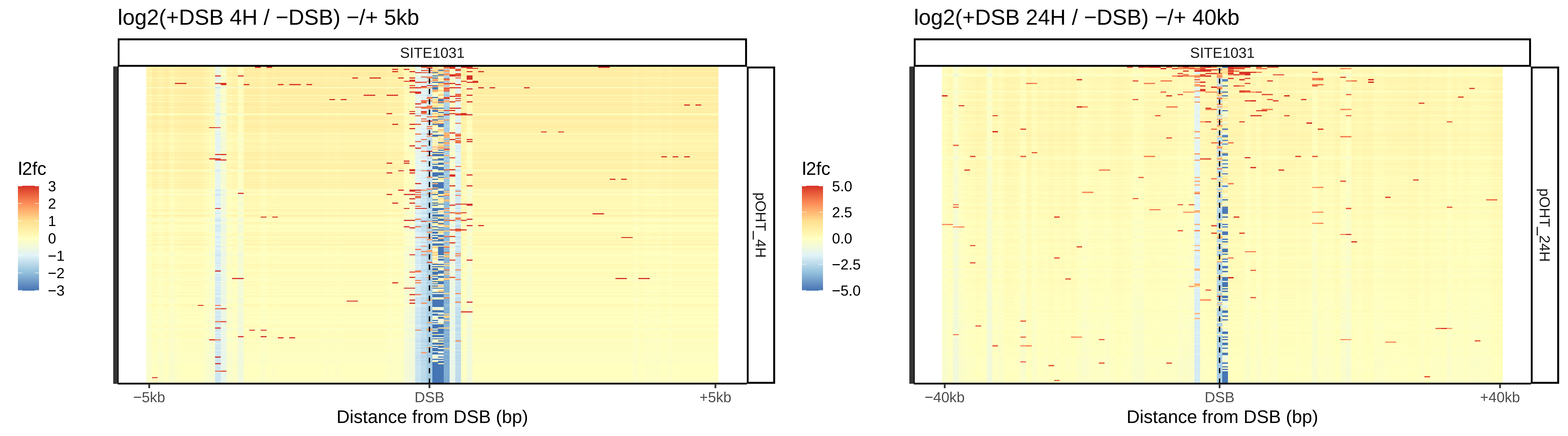
log2(+DSB 4H / −DSB) −/+ 5kb
SITE1031
l2fc
3
2
1
0
−1
−2
−3
pOHT_4H
−5kb
DSB
+5kb
Distance from DSB (bp)
log2(+DSB 24H / −DSB) −/+ 40kb
SITE1031
l2fc
5.0
2.5
0.0
−2.5
−5.0
pOHT_24H
−40kb
DSB
+40kb
Distance from DSB (bp)

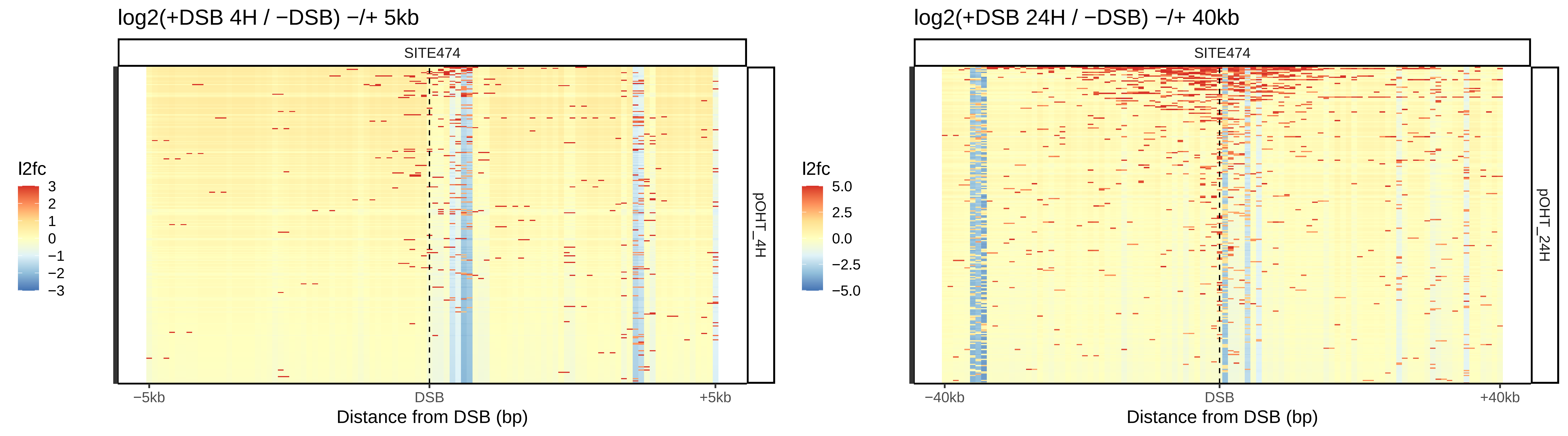

log2(+DSB 4H / −DSB) −/+ 5kb
SITE474
pOHT_4H
l2fc
3
2
1
0
−1
−2
−3
−5kb
DSB
+5kb
Distance from DSB (bp)
log2(+DSB 24H / −DSB) −/+ 40kb
SITE474
pOHT_24H
l2fc
5.0
2.5
0.0
−2.5
−5.0
−40kb
DSB
+40kb
Distance from DSB (bp)

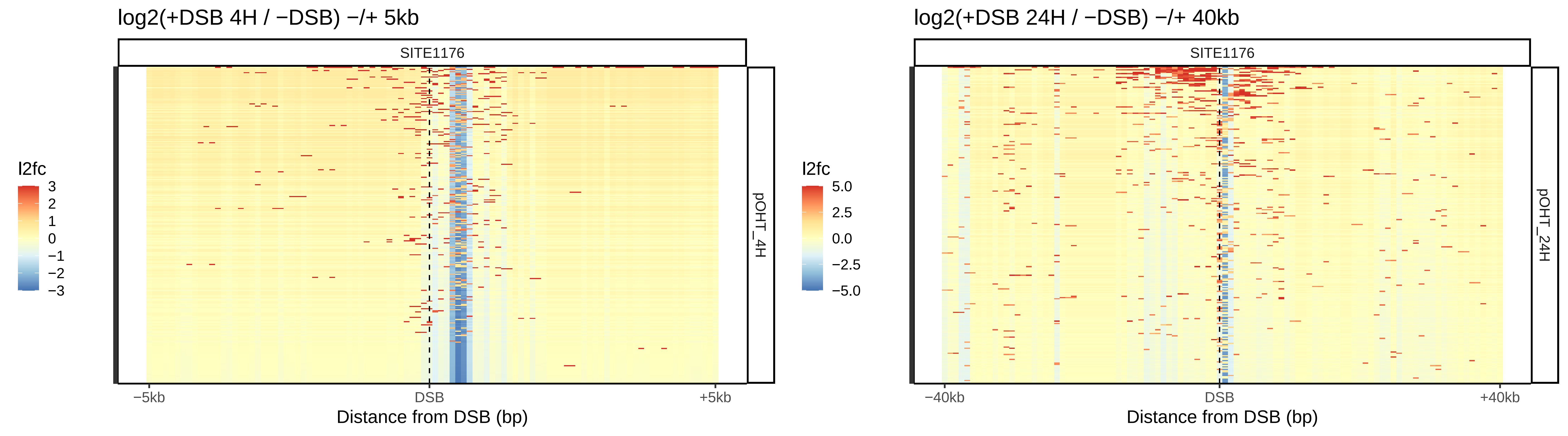
log2(+DSB 4H / −DSB) −/+ 5kb
SITE1176
l2fc
3
2
1
0
−1
−2
−3
pOHT_4H
−5kb
DSB
+5kb
Distance from DSB (bp)
log2(+DSB 24H / −DSB) −/+ 40kb
SITE1176
l2fc
5.0
2.5
0.0
−2.5
−5.0
pOHT_24H
−40kb
DSB
+40kb
Distance from DSB (bp)

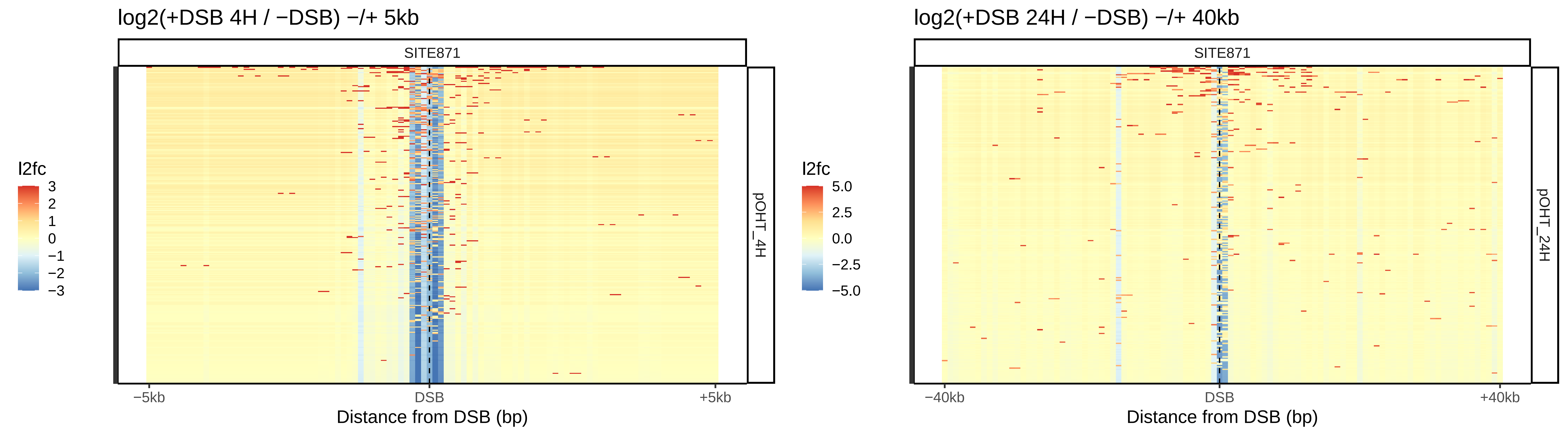

log2(+DSB 4H / −DSB) −/+ 5kb
SITE871
pOHT_4H
l2fc
3
2
1
0
−1
−2
−3
−5kb
DSB
+5kb
Distance from DSB (bp)
log2(+DSB 24H / −DSB) −/+ 40kb
SITE871
pOHT_24H
l2fc
5.0
2.5
0.0
−2.5
−5.0
−40kb
DSB
+40kb
Distance from DSB (bp)

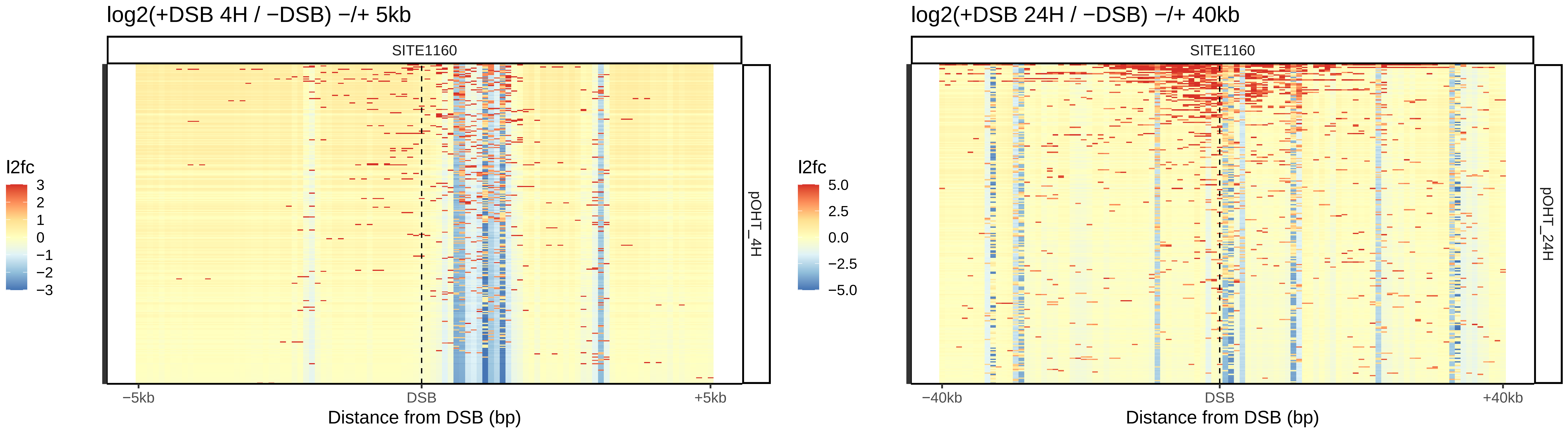

log2(+DSB 4H / −DSB) −/+ 5kb
SITE1160
l2fc
3
2
1
0
−1
−2
−3
pOHT_4H
−5kb
DSB
+5kb
Distance from DSB (bp)
log2(+DSB 24H / −DSB) −/+ 40kb
SITE1160
l2fc
5.0
2.5
0.0
−2.5
−5.0
pOHT_24H
−40kb
DSB
+40kb
Distance from DSB (bp)

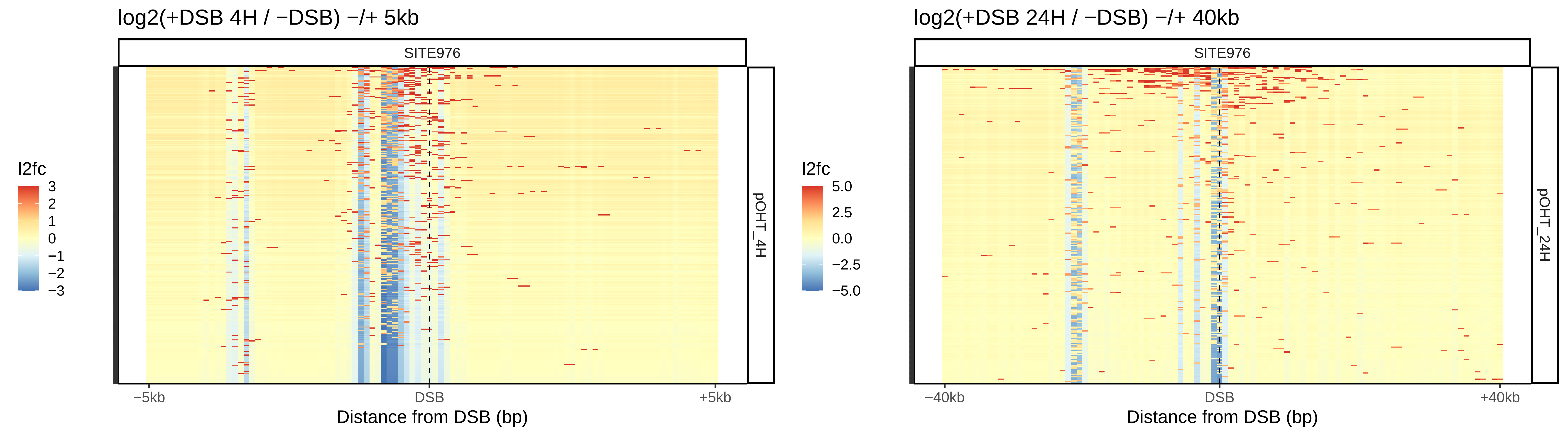

log2(+DSB 4H / −DSB) −/+ 5kb
SITE976
l2fc
3
2
1
0
−1
−2
−3
pOHT_4H
−5kb
DSB
+5kb
Distance from DSB (bp)
log2(+DSB 24H / −DSB) −/+ 40kb
SITE976
l2fc
5.0
2.5
0.0
−2.5
−5.0
pOHT_24H
−40kb
DSB
+40kb
Distance from DSB (bp)

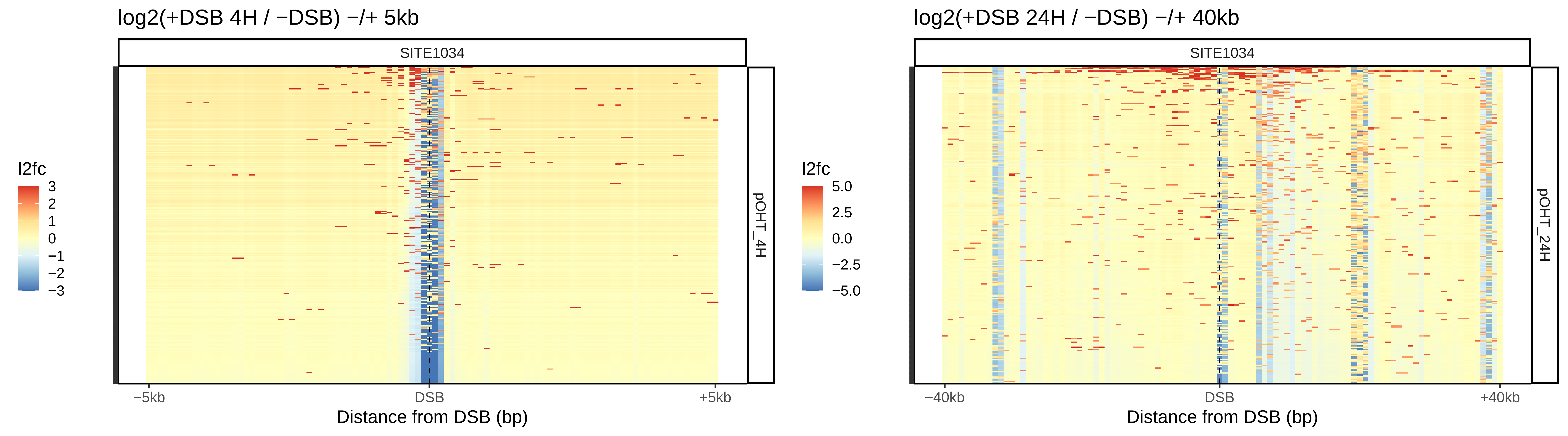

log2(+DSB 4H / −DSB) −/+ 5kb
SITE1034
pOHT_4H
l2fc
3
2
1
0
−1
−2
−3
−5kb
DSB
+5kb
Distance from DSB (bp)
log2(+DSB 24H / −DSB) −/+ 40kb
SITE1034
pOHT_24H
l2fc
5.0
2.5
0.0
−2.5
−5.0
−40kb
DSB
+40kb
Distance from DSB (bp)

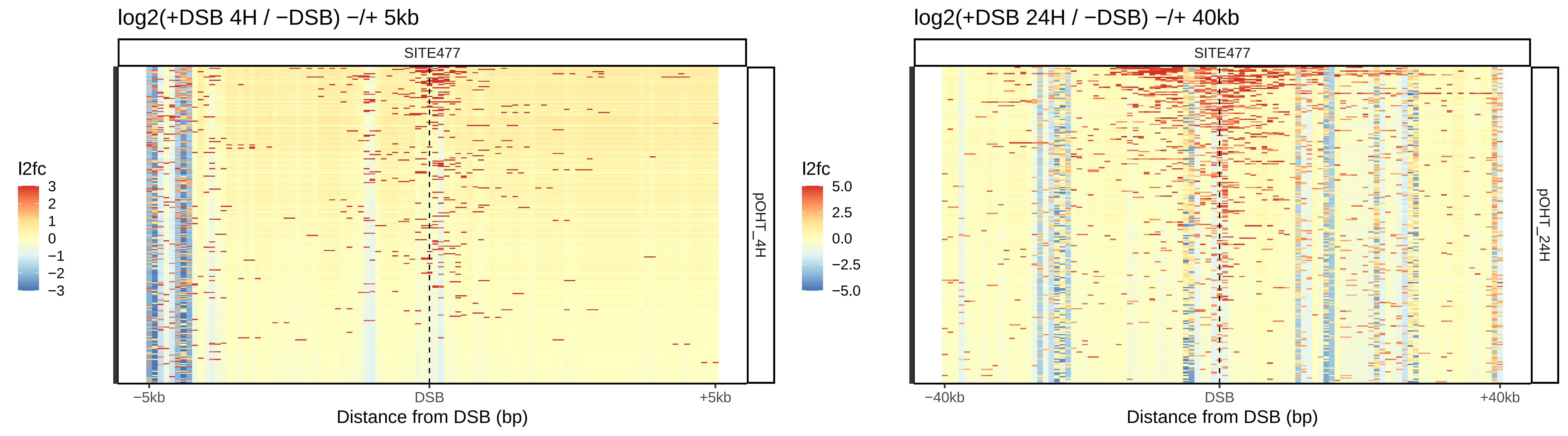

log2(+DSB 4H / −DSB) −/+ 5kb
SITE477
pOHT_4H
l2fc
3
2
1
0
−1
−2
−3
−5kb
DSB
+5kb
Distance from DSB (bp)
log2(+DSB 24H / −DSB) −/+ 40kb
SITE477
pOHT_24H
l2fc
5.0
2.5
0.0
−2.5
−5.0
−40kb
DSB
+40kb
Distance from DSB (bp)

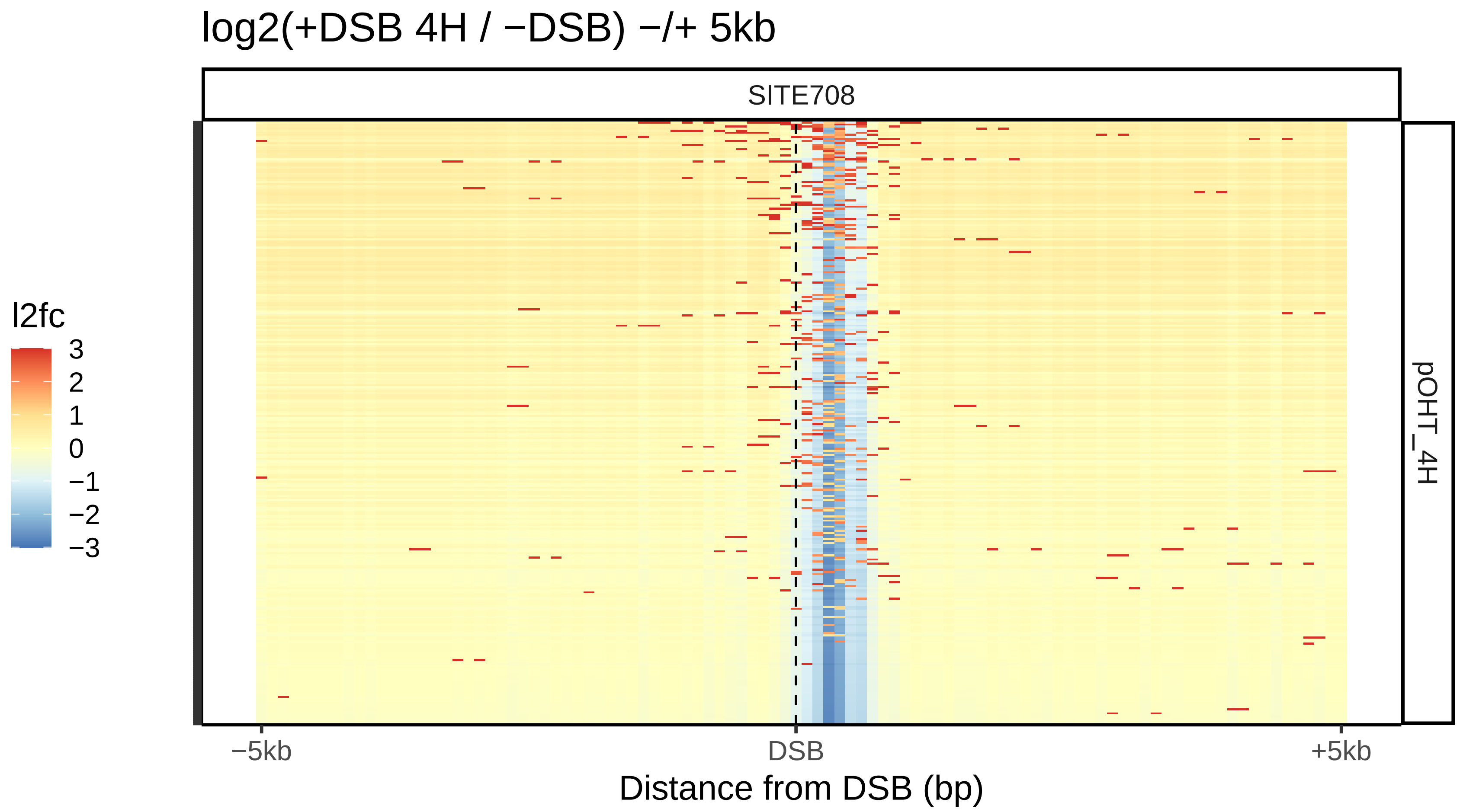

log2(+DSB 4H / −DSB) −/+ 5kb
SITE708
l2fc
3
2
1
0
−1
−2
−3
pOHT_4H
−5kb
DSB
+5kb
Distance from DSB (bp)


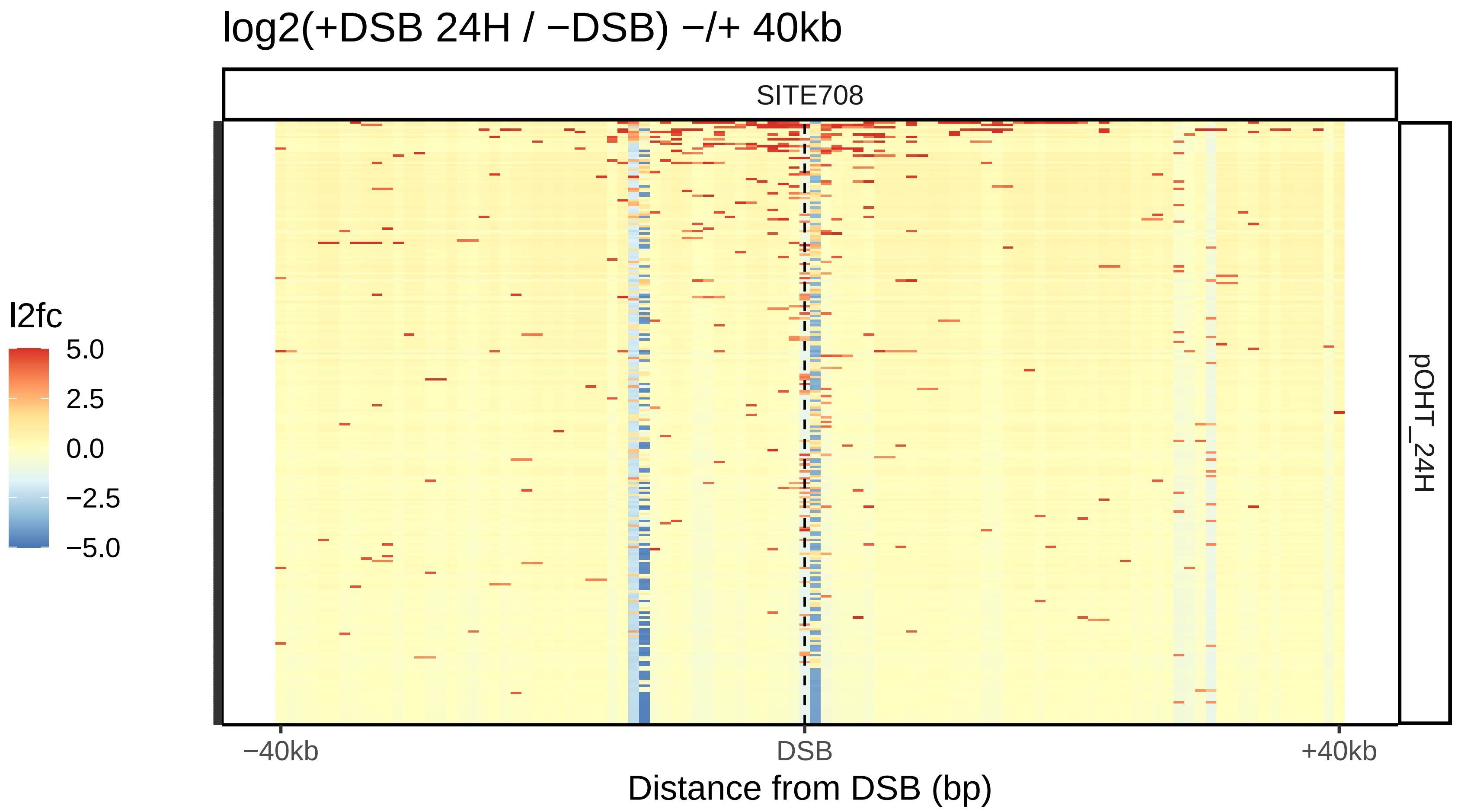

log2(+DSB 24H / −DSB) −/+ 40kb
SITE708
l2fc
5.0
2.5
0.0
−2.5
−5.0
pOHT_24H
−40kb
DSB
+40kb
Distance from DSB (bp)

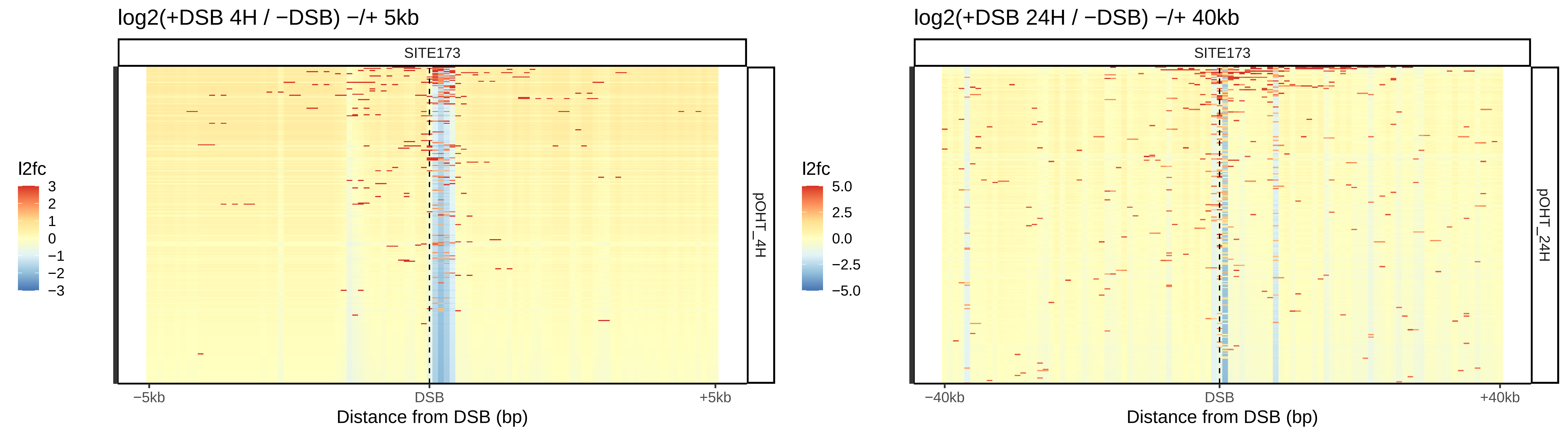

log2(+DSB 4H / −DSB) −/+ 5kb
SITE173
pOHT_4H
l2fc
3
2
1
0
−1
−2
−3
−5kb
DSB
+5kb
Distance from DSB (bp)
log2(+DSB 24H / −DSB) −/+ 40kb
SITE173
pOHT_24H
l2fc
5.0
2.5
0.0
−2.5
−5.0
−40kb
DSB
+40kb
Distance from DSB (bp)

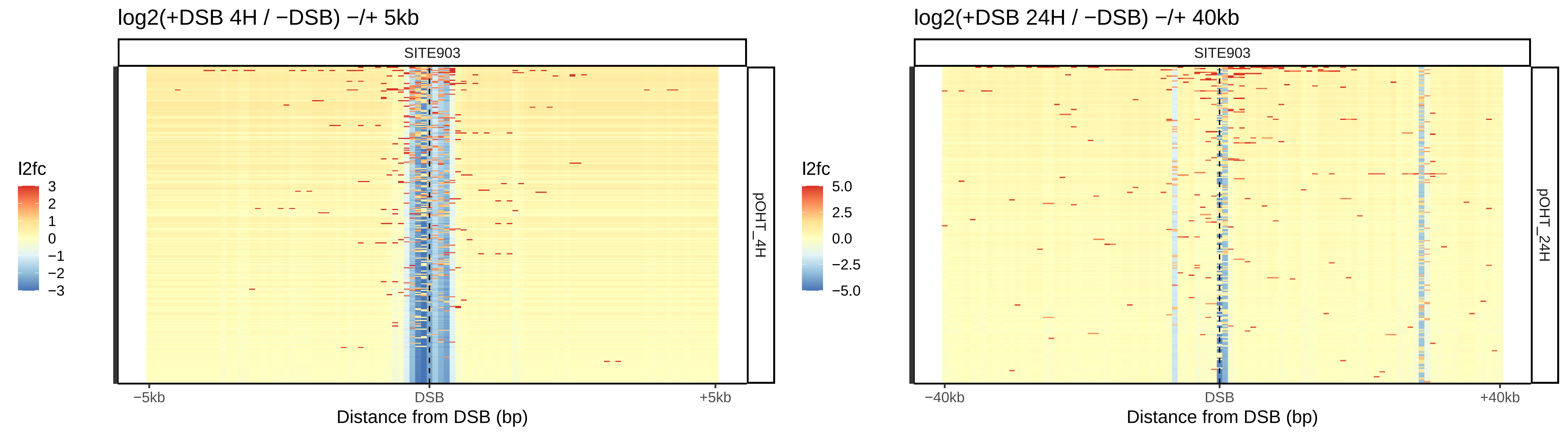

log2(+DSB 4H / −DSB) −/+ 5kb
SITE903
pOHT_4H
l2fc
3
2
1
0
−1
−2
−3
−5kb
DSB
+5kb
Distance from DSB (bp)
log2(+DSB 24H / −DSB) −/+ 40kb
SITE903
pOHT_24H
l2fc
5.0
2.5
0.0
−2.5
−5.0
−40kb
DSB
+40kb
Distance from DSB (bp)

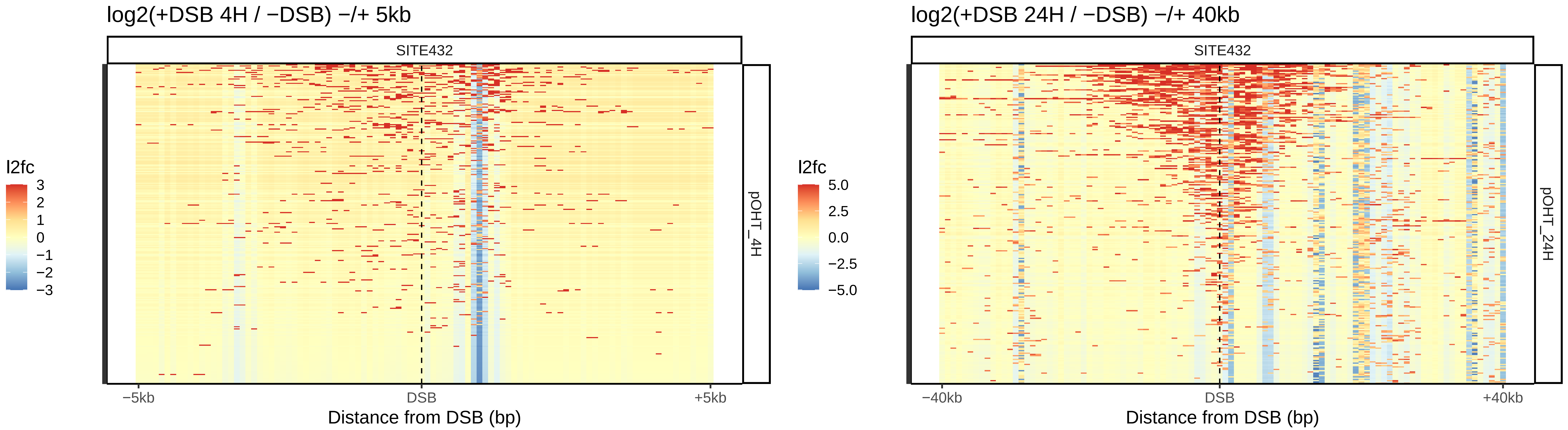

log2(+DSB 4H / −DSB) −/+ 5kb
SITE432
l2fc
3
2
1
0
−1
−2
−3
pOHT_4H
−5kb
DSB
+5kb
Distance from DSB (bp)
log2(+DSB 24H / −DSB) −/+ 40kb
SITE432
l2fc
5.0
2.5
0.0
−2.5
−5.0
pOHT_24H
−40kb
DSB
+40kb
Distance from DSB (bp)

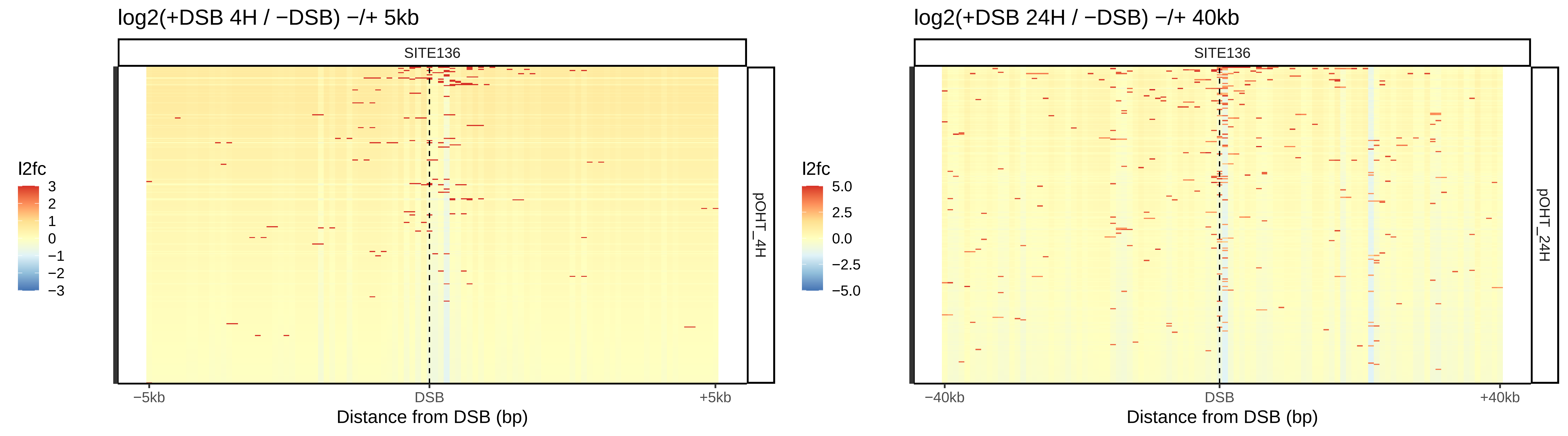

log2(+DSB 4H / −DSB) −/+ 5kb
SITE136
pOHT_4H
l2fc
3
2
1
0
−1
−2
−3
−5kb
DSB
+5kb
Distance from DSB (bp)
log2(+DSB 24H / −DSB) −/+ 40kb
SITE136
pOHT_24H
l2fc
5.0
2.5
0.0
−2.5
−5.0
−40kb
DSB
+40kb
Distance from DSB (bp)

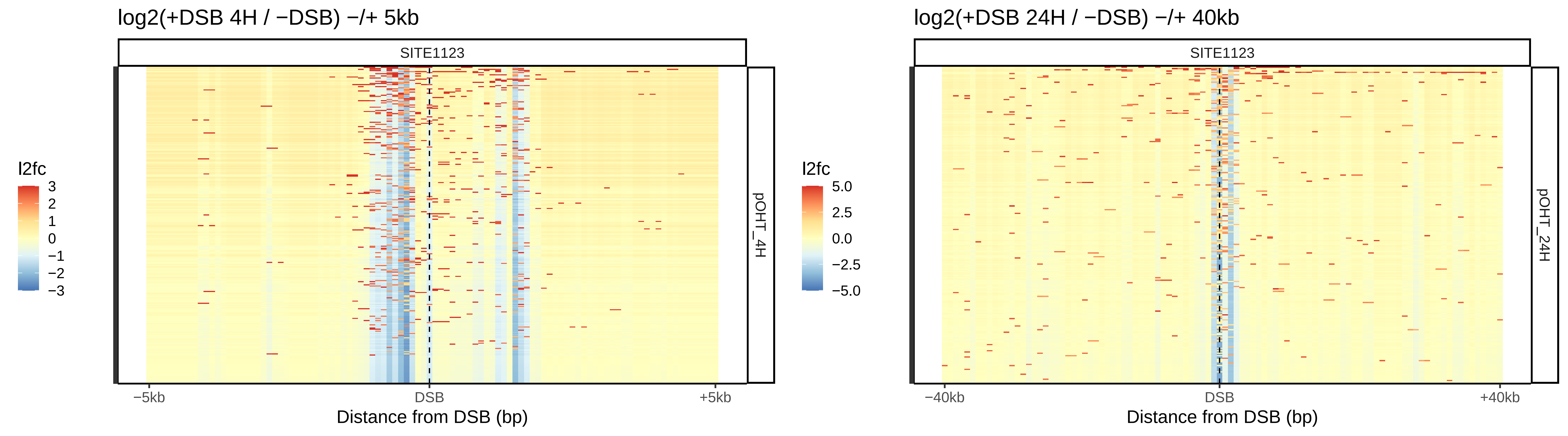

log2(+DSB 4H / −DSB) −/+ 5kb
SITE1123
l2fc
3
2
1
0
−1
−2
−3
pOHT_4H
−5kb
DSB
+5kb
Distance from DSB (bp)
log2(+DSB 24H / −DSB) −/+ 40kb
SITE1123
l2fc
5.0
2.5
0.0
−2.5
−5.0
pOHT_24H
−40kb
DSB
+40kb
Distance from DSB (bp)

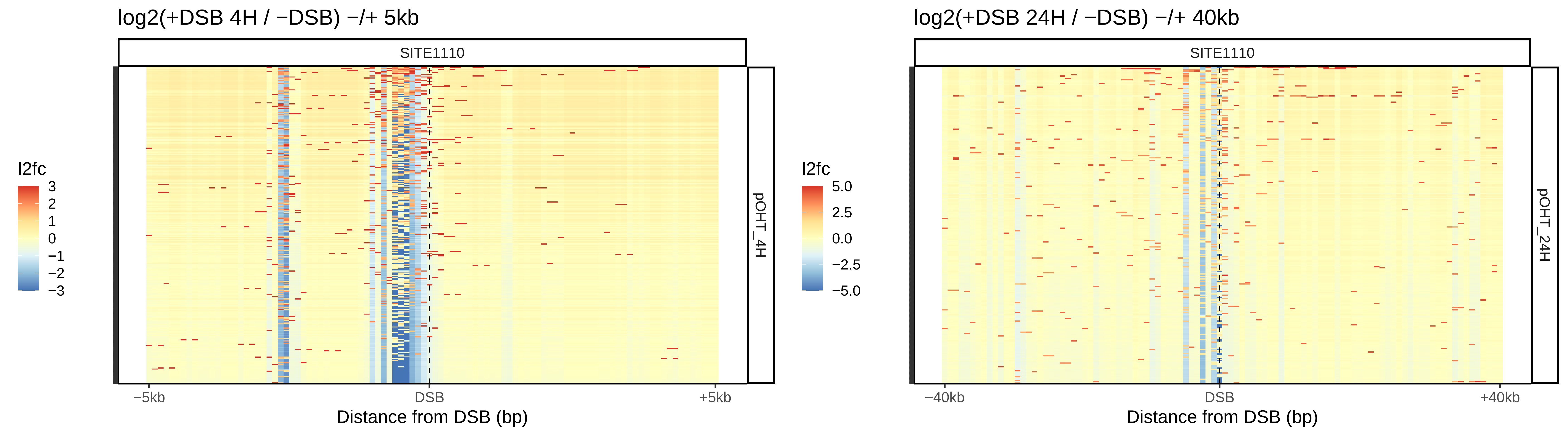
log2(+DSB 4H / −DSB) −/+ 5kb
SITE1110
pOHT_4H
l2fc
3
2
1
0
−1
−2
−3
−5kb
DSB
+5kb
Distance from DSB (bp)
log2(+DSB 24H / −DSB) −/+ 40kb
SITE1110
pOHT_24H
l2fc
5.0
2.5
0.0
−2.5
−5.0
−40kb
DSB
+40kb
Distance from DSB (bp)

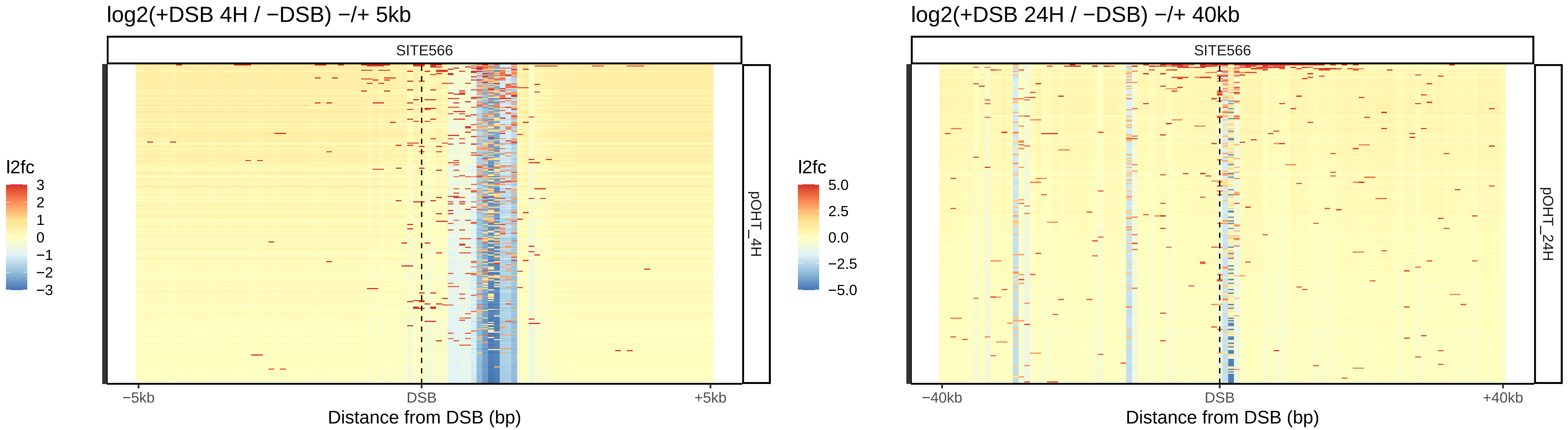

log2(+DSB 4H / −DSB) −/+ 5kb
SITE566
pOHT_4H
l2fc
3
2
1
0
−1
−2
−3
−5kb
DSB
+5kb
Distance from DSB (bp)
log2(+DSB 24H / −DSB) −/+ 40kb
SITE566
pOHT_24H
l2fc
5.0
2.5
0.0
−2.5
−5.0
−40kb
DSB
+40kb
Distance from DSB (bp)

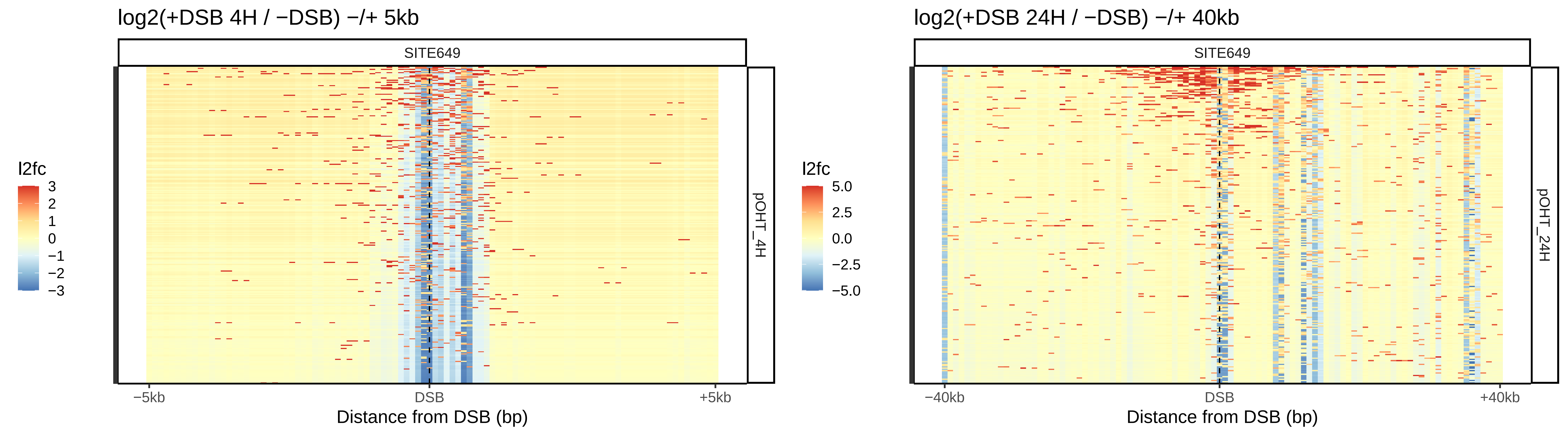
log2(+DSB 4H / −DSB) −/+ 5kb
SITE649
pOHT_4H
l2fc
3
2
1
0
−1
−2
−3
−5kb
DSB
+5kb
Distance from DSB (bp)
log2(+DSB 24H / −DSB) −/+ 40kb
SITE649
pOHT_24H
l2fc
5.0
2.5
0.0
−2.5
−5.0
−40kb
DSB
+40kb
Distance from DSB (bp)

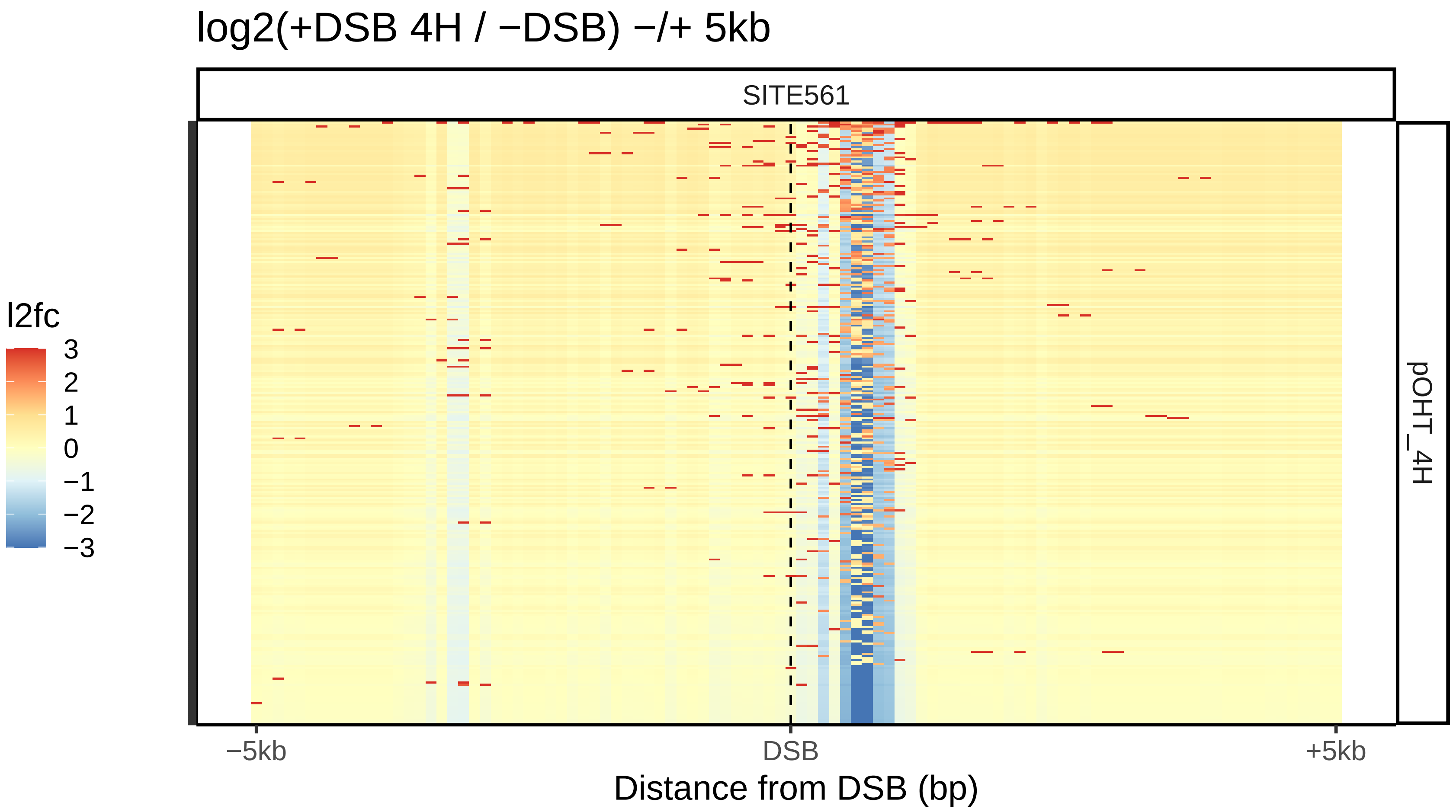

log2(+DSB 4H / −DSB) −/+ 5kb
SITE561
pOHT_4H
l2fc
3
2
1
0
−1
−2
−3
−5kb
DSB
+5kb
Distance from DSB (bp)


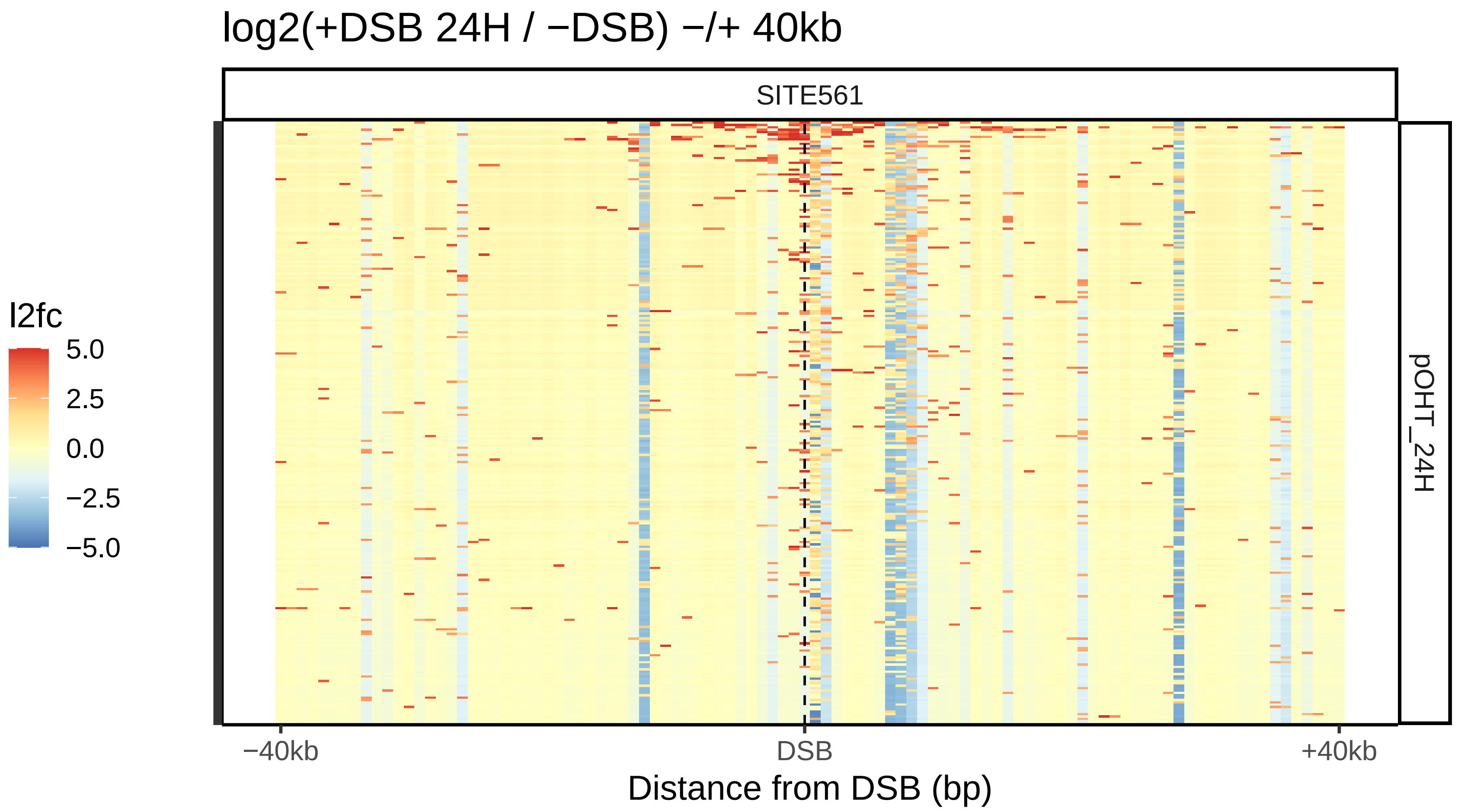

log2(+DSB 24H / −DSB) −/+ 40kb
SITE561
pOHT_24H
l2fc
5.0
2.5
0.0
−2.5
−5.0
−40kb
DSB
+40kb
Distance from DSB (bp)

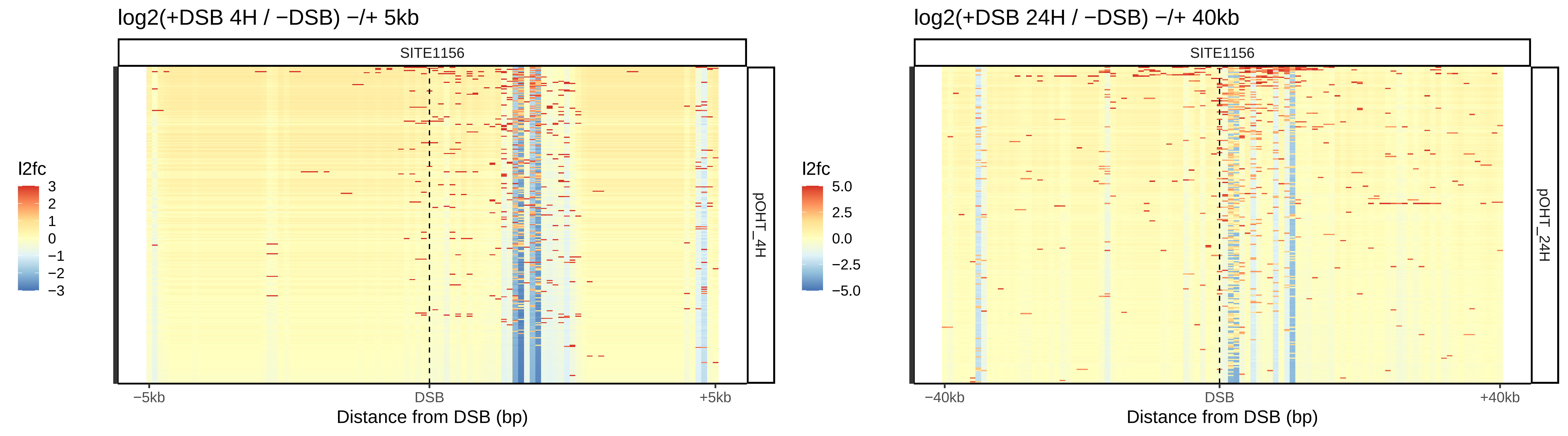

log2(+DSB 4H / −DSB) −/+ 5kb
SITE1156
l2fc
3
2
1
0
−1
−2
−3
pOHT_4H
−5kb
DSB
+5kb
Distance from DSB (bp)
log2(+DSB 24H / −DSB) −/+ 40kb
SITE1156
l2fc
5.0
2.5
0.0
−2.5
−5.0
pOHT_24H
−40kb
DSB
+40kb
Distance from DSB (bp)

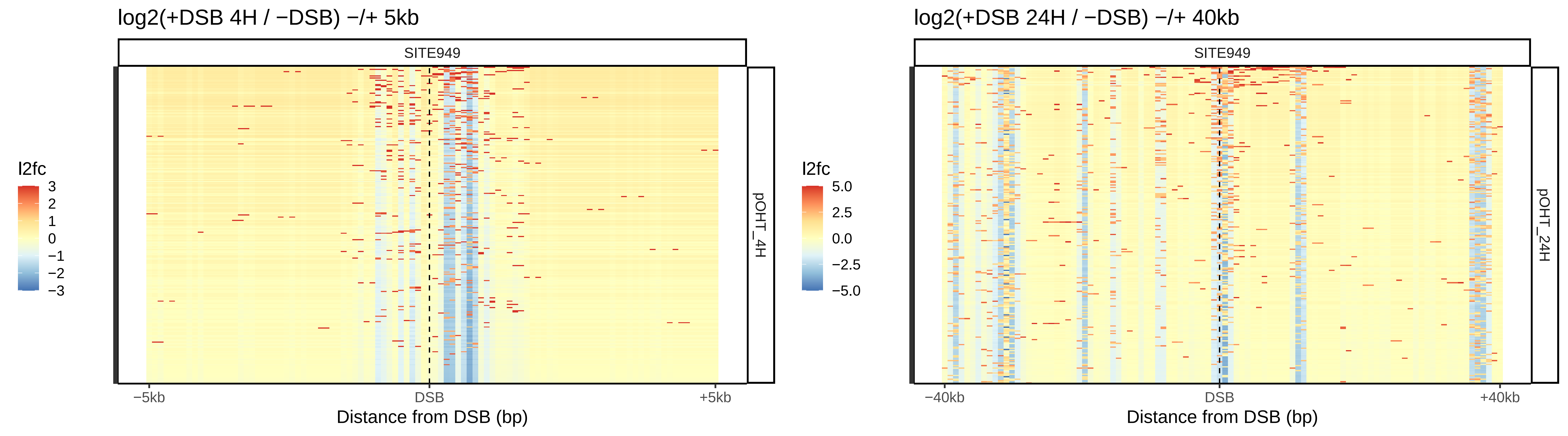
log2(+DSB 4H / −DSB) −/+ 5kb
SITE949
l2fc
3
2
1
0
−1
−2
−3
pOHT_4H
−5kb
DSB
+5kb
Distance from DSB (bp)
log2(+DSB 24H / −DSB) −/+ 40kb
SITE949
l2fc
5.0
2.5
0.0
−2.5
−5.0
pOHT_24H
−40kb
DSB
+40kb
Distance from DSB (bp)

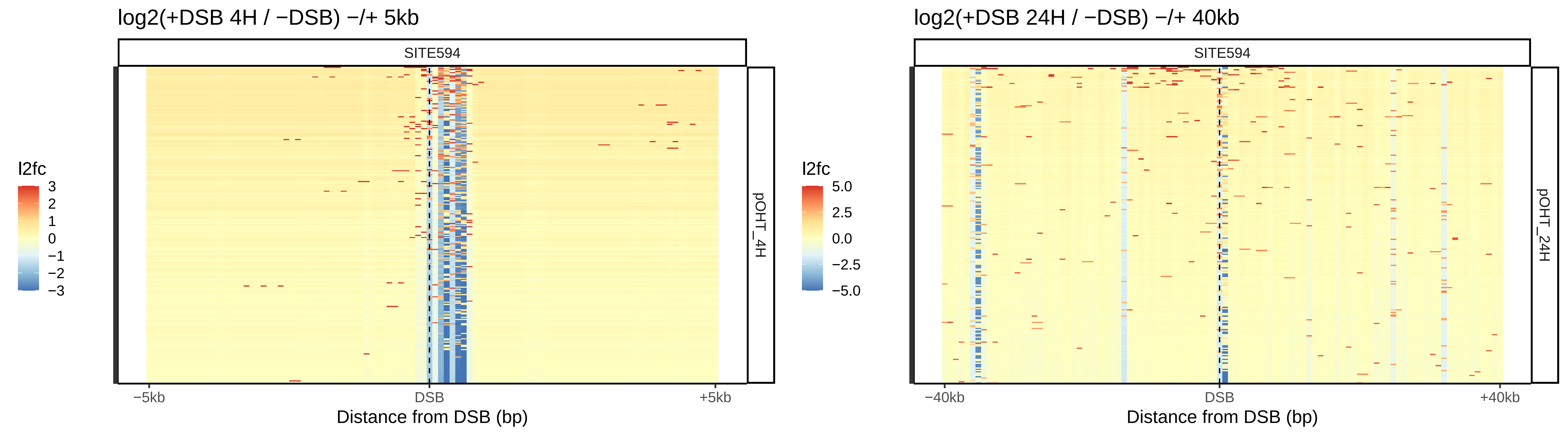

log2(+DSB 4H / −DSB) −/+ 5kb
SITE594
l2fc
3
2
1
0
−1
−2
−3
pOHT_4H
−5kb
DSB
+5kb
Distance from DSB (bp)
log2(+DSB 24H / −DSB) −/+ 40kb
SITE594
l2fc
5.0
2.5
0.0
−2.5
−5.0
pOHT_24H
−40kb
DSB
+40kb
Distance from DSB (bp)

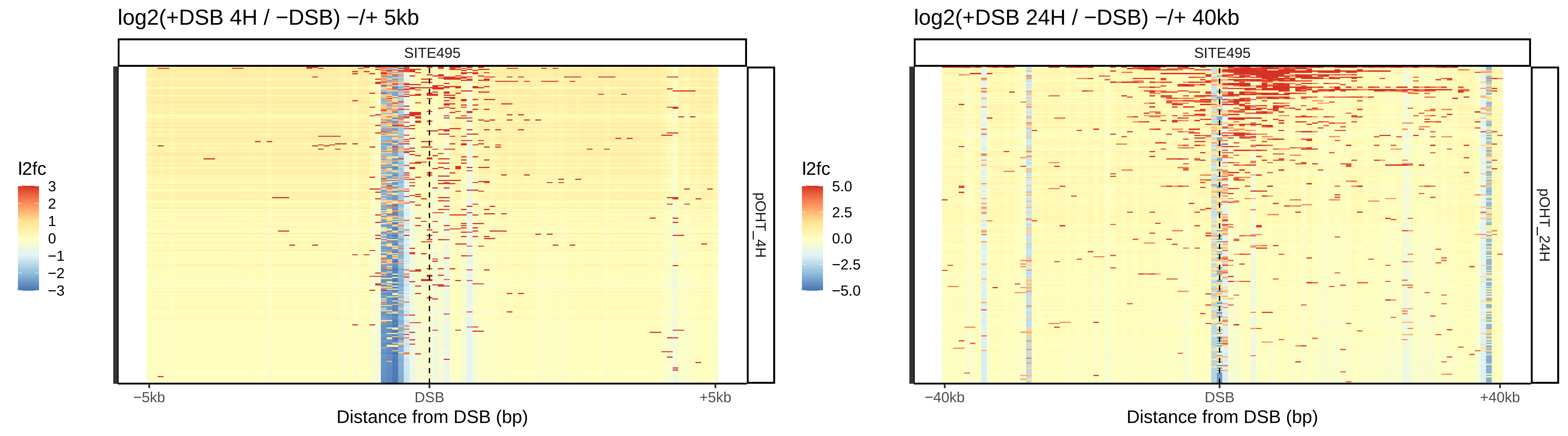
log2(+DSB 4H / −DSB) −/+ 5kb
SITE495
l2fc
3
2
1
0
−1
−2
−3
pOHT_4H
−5kb
DSB
+5kb
Distance from DSB (bp)
log2(+DSB 24H / −DSB) −/+ 40kb
SITE495
l2fc
5.0
2.5
0.0
−2.5
−5.0
pOHT_24H
−40kb
DSB
+40kb
Distance from DSB (bp)

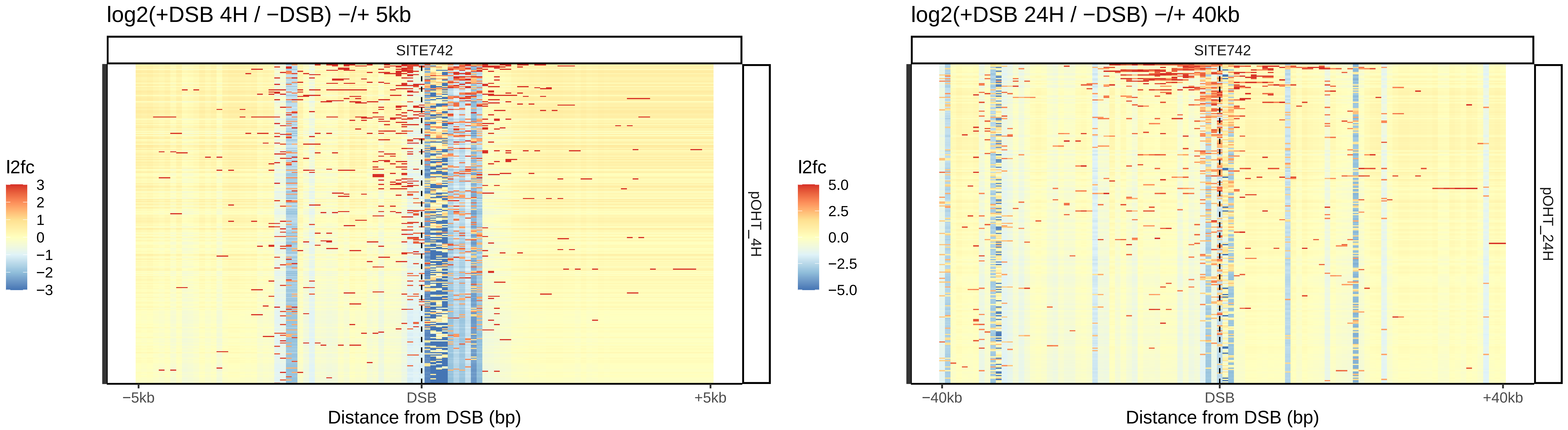

log2(+DSB 4H / −DSB) −/+ 5kb
SITE742
pOHT_4H
l2fc
3
2
1
0
−1
−2
−3
−5kb
DSB
+5kb
Distance from DSB (bp)
log2(+DSB 24H / −DSB) −/+ 40kb
SITE742
pOHT_24H
l2fc
5.0
2.5
0.0
−2.5
−5.0
−40kb
DSB
+40kb
Distance from DSB (bp)

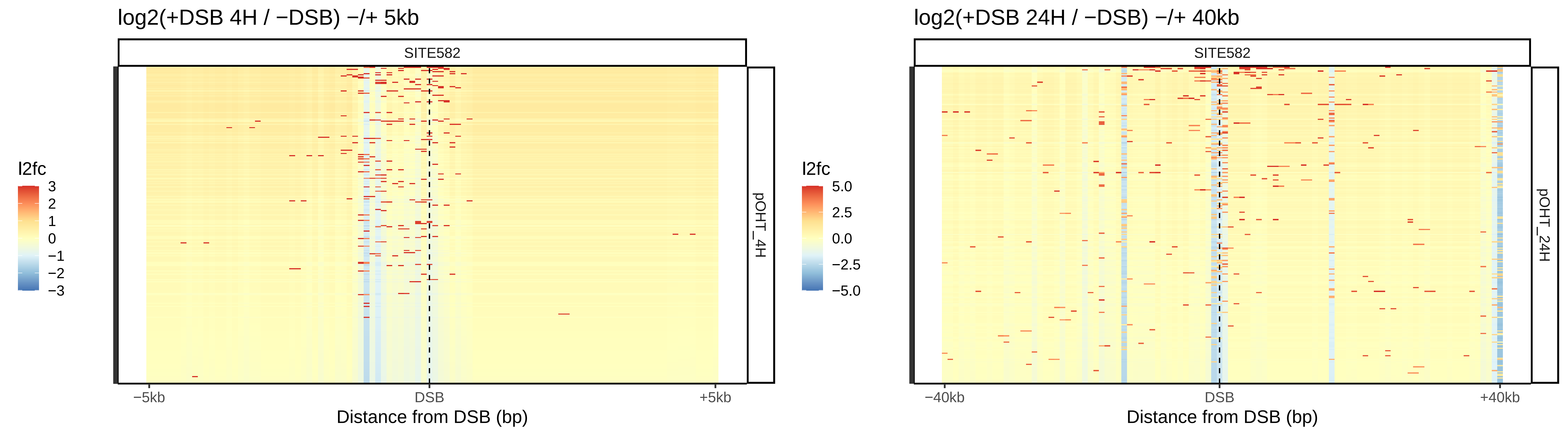

log2(+DSB 4H / −DSB) −/+ 5kb
SITE582
l2fc
3
2
1
0
−1
−2
−3
pOHT_4H
−5kb
DSB
+5kb
Distance from DSB (bp)
log2(+DSB 24H / −DSB) −/+ 40kb
SITE582
l2fc
5.0
2.5
0.0
−2.5
−5.0
pOHT_24H
−40kb
DSB
+40kb
Distance from DSB (bp)

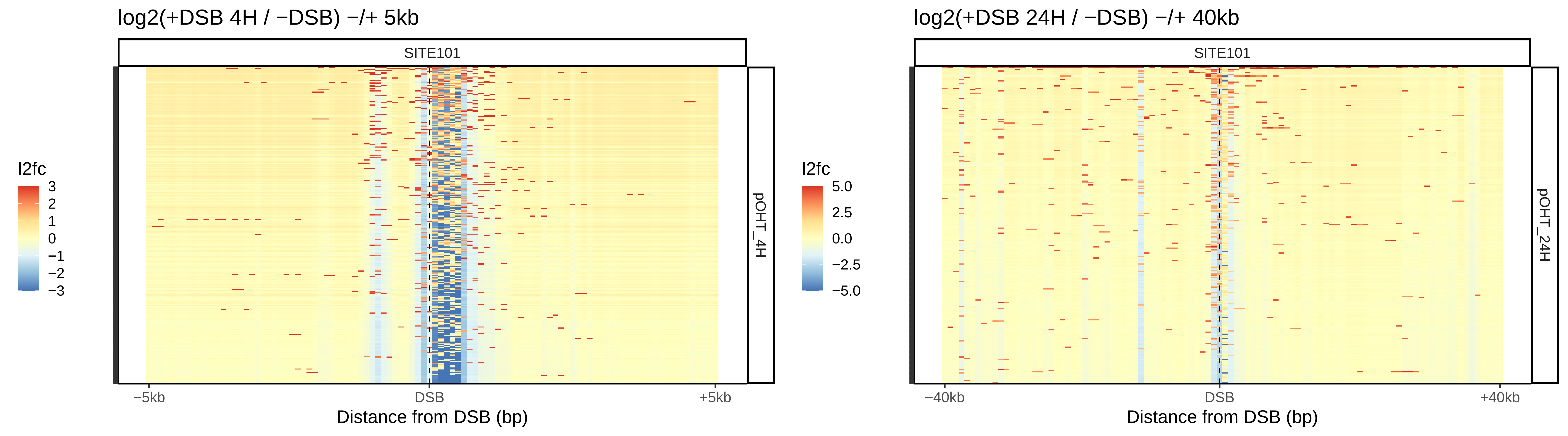

log2(+DSB 4H / −DSB) −/+ 5kb
SITE101
pOHT_4H
l2fc
3
2
1
0
−1
−2
−3
−5kb
DSB
+5kb
Distance from DSB (bp)
log2(+DSB 24H / −DSB) −/+ 40kb
SITE101
pOHT_24H
l2fc
5.0
2.5
0.0
−2.5
−5.0
−40kb
DSB
+40kb
Distance from DSB (bp)

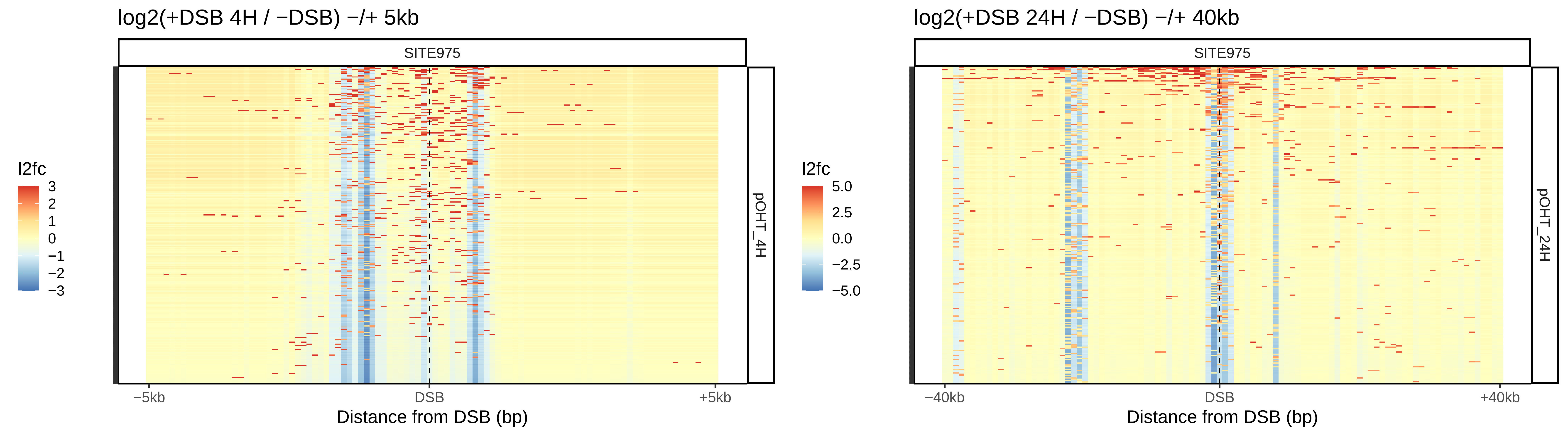

log2(+DSB 4H / −DSB) −/+ 5kb
SITE975
pOHT_4H
l2fc
3
2
1
0
−1
−2
−3
−5kb
DSB
+5kb
Distance from DSB (bp)
log2(+DSB 24H / −DSB) −/+ 40kb
SITE975
pOHT_24H
l2fc
5.0
2.5
0.0
−2.5
−5.0
−40kb
DSB
+40kb
Distance from DSB (bp)

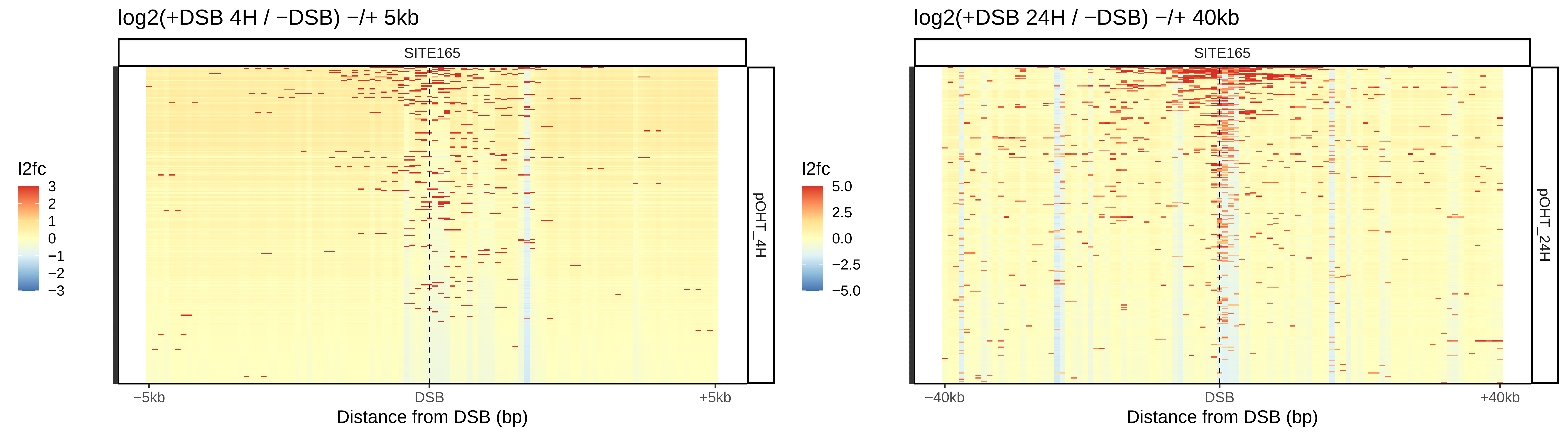

log2(+DSB 4H / −DSB) −/+ 5kb
SITE165
pOHT_4H
l2fc
3
2
1
0
−1
−2
−3
−5kb
DSB
+5kb
Distance from DSB (bp)
log2(+DSB 24H / −DSB) −/+ 40kb
SITE165
pOHT_24H
l2fc
5.0
2.5
0.0
−2.5
−5.0
−40kb
DSB
+40kb
Distance from DSB (bp)

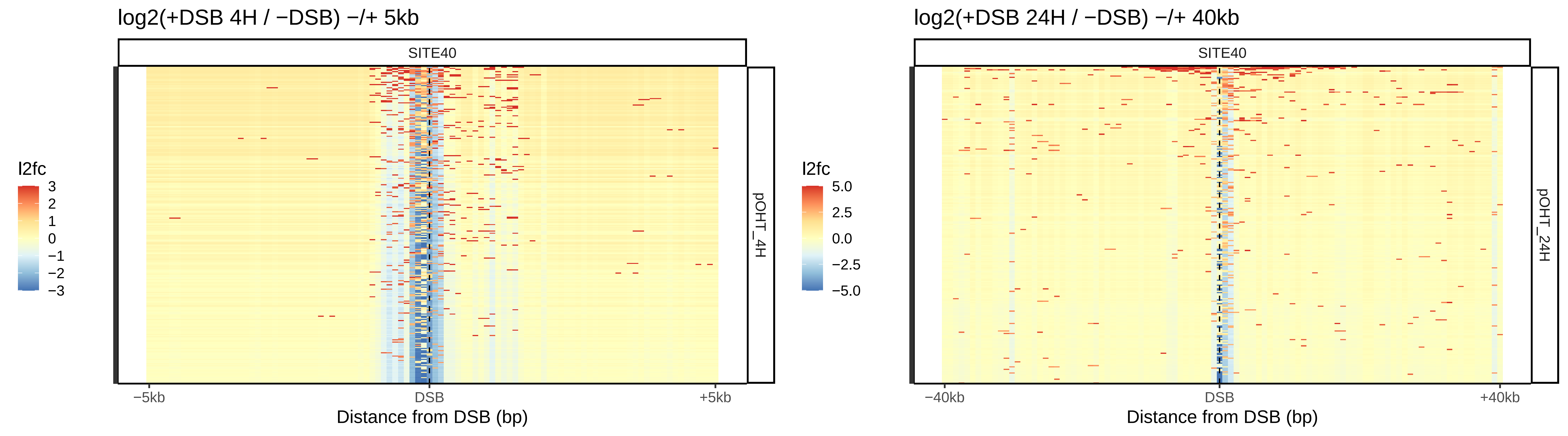

log2(+DSB 4H / −DSB) −/+ 5kb
SITE40
pOHT_4H
l2fc
3
2
1
0
−1
−2
−3
−5kb
DSB
+5kb
Distance from DSB (bp)
log2(+DSB 24H / −DSB) −/+ 40kb
SITE40
pOHT_24H
l2fc
5.0
2.5
0.0
−2.5
−5.0
−40kb
DSB
+40kb
Distance from DSB (bp)

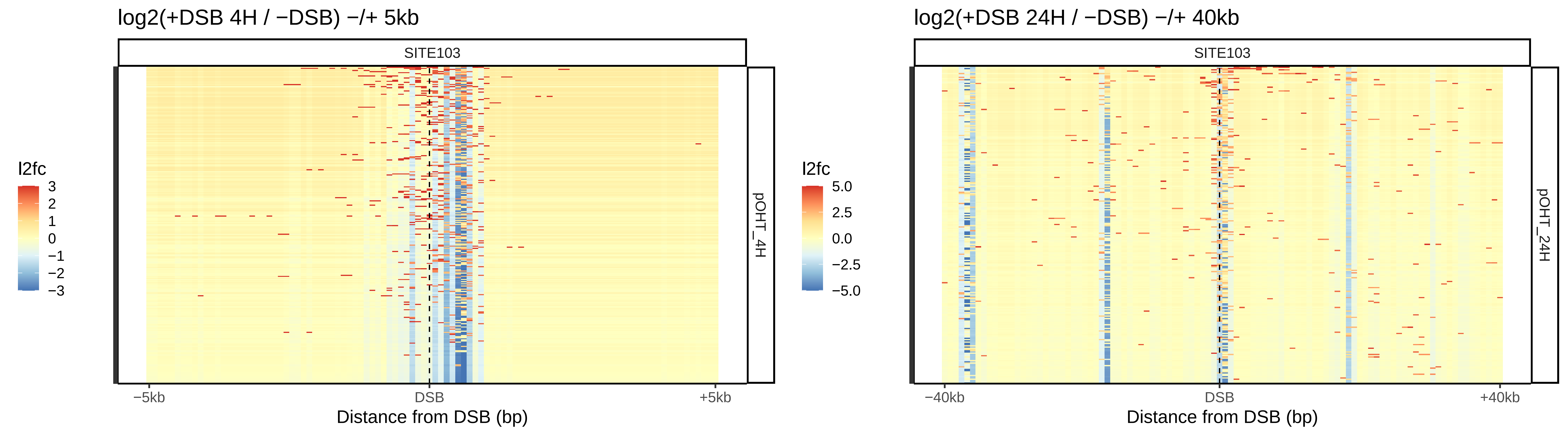

log2(+DSB 4H / −DSB) −/+ 5kb
SITE103
l2fc
3
2
1
0
−1
−2
−3
pOHT_4H
−5kb
DSB
+5kb
Distance from DSB (bp)
log2(+DSB 24H / −DSB) −/+ 40kb
SITE103
l2fc
5.0
2.5
0.0
−2.5
−5.0
pOHT_24H
−40kb
DSB
+40kb
Distance from DSB (bp)

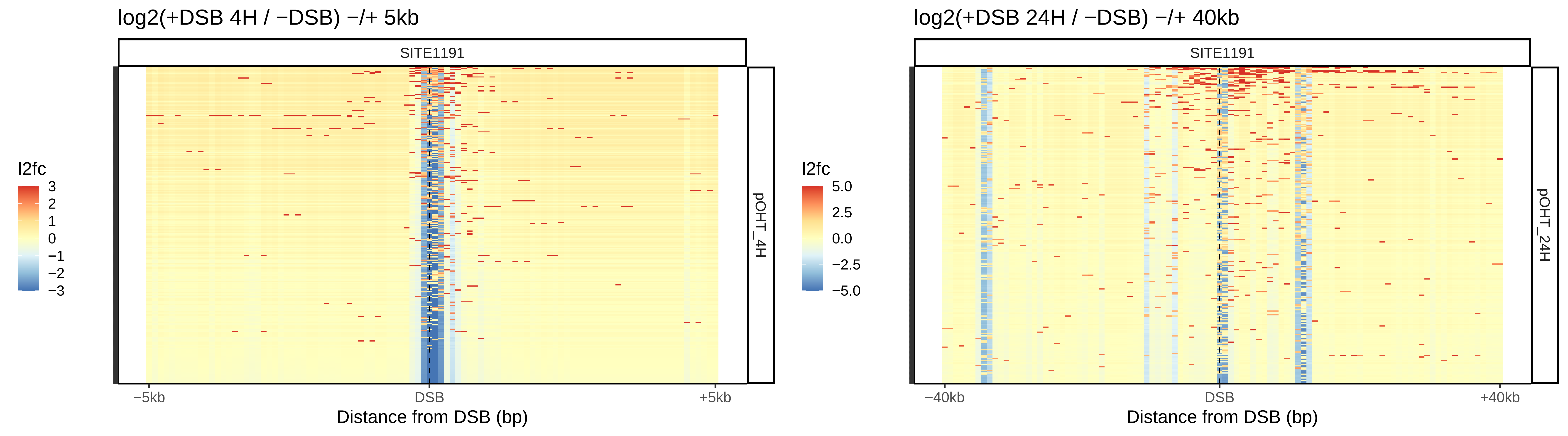

log2(+DSB 4H / −DSB) −/+ 5kb
SITE1191
l2fc
3
2
1
0
−1
−2
−3
pOHT_4H
−5kb
DSB
+5kb
Distance from DSB (bp)
log2(+DSB 24H / −DSB) −/+ 40kb
SITE1191
l2fc
5.0
2.5
0.0
−2.5
−5.0
pOHT_24H
−40kb
DSB
+40kb
Distance from DSB (bp)

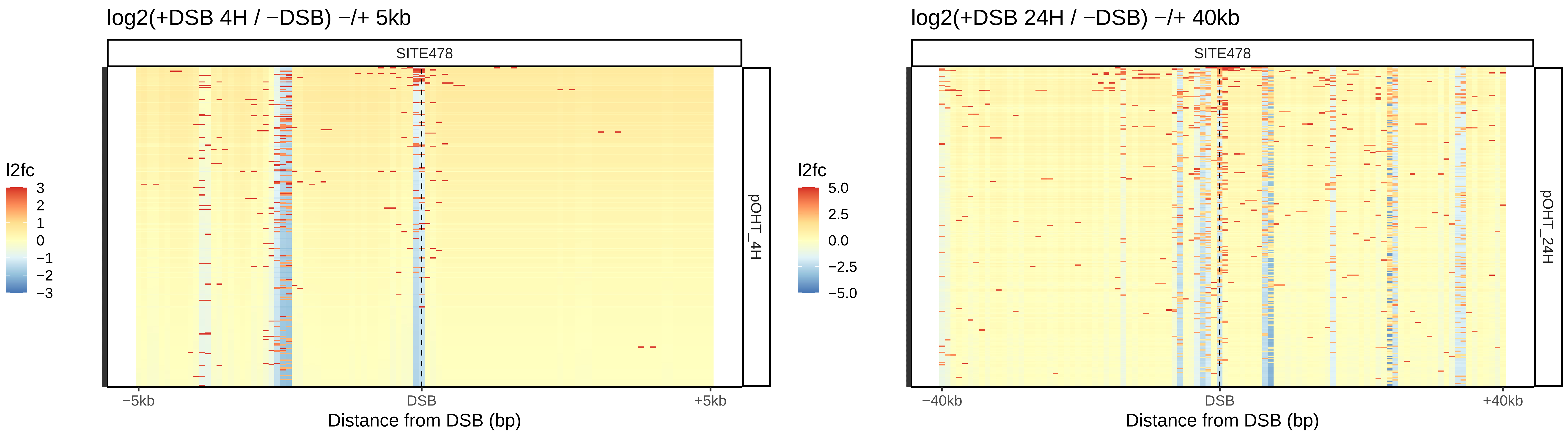
log2(+DSB 4H / −DSB) −/+ 5kb
SITE478
l2fc
3
2
1
0
−1
−2
−3
pOHT_4H
−5kb
DSB
+5kb
Distance from DSB (bp)
log2(+DSB 24H / −DSB) −/+ 40kb
SITE478
l2fc
5.0
2.5
0.0
−2.5
−5.0
pOHT_24H
−40kb
DSB
+40kb
Distance from DSB (bp)

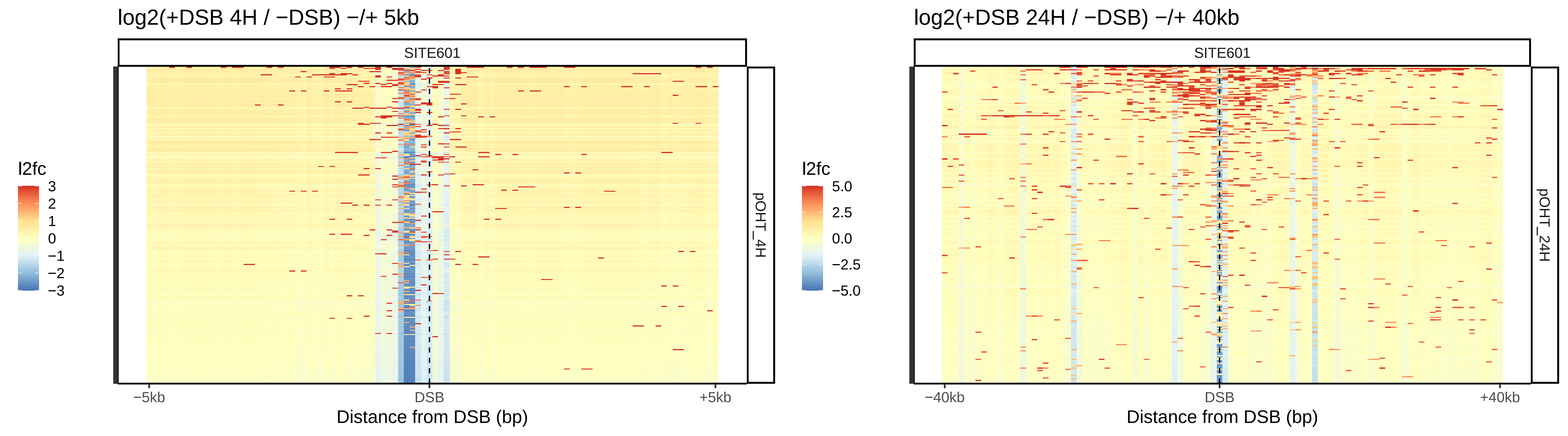

log2(+DSB 4H / −DSB) −/+ 5kb
SITE601
pOHT_4H
l2fc
3
2
1
0
−1
−2
−3
−5kb
DSB
+5kb
Distance from DSB (bp)
log2(+DSB 24H / −DSB) −/+ 40kb
SITE601
pOHT_24H
l2fc
5.0
2.5
0.0
−2.5
−5.0
−40kb
DSB
+40kb
Distance from DSB (bp)

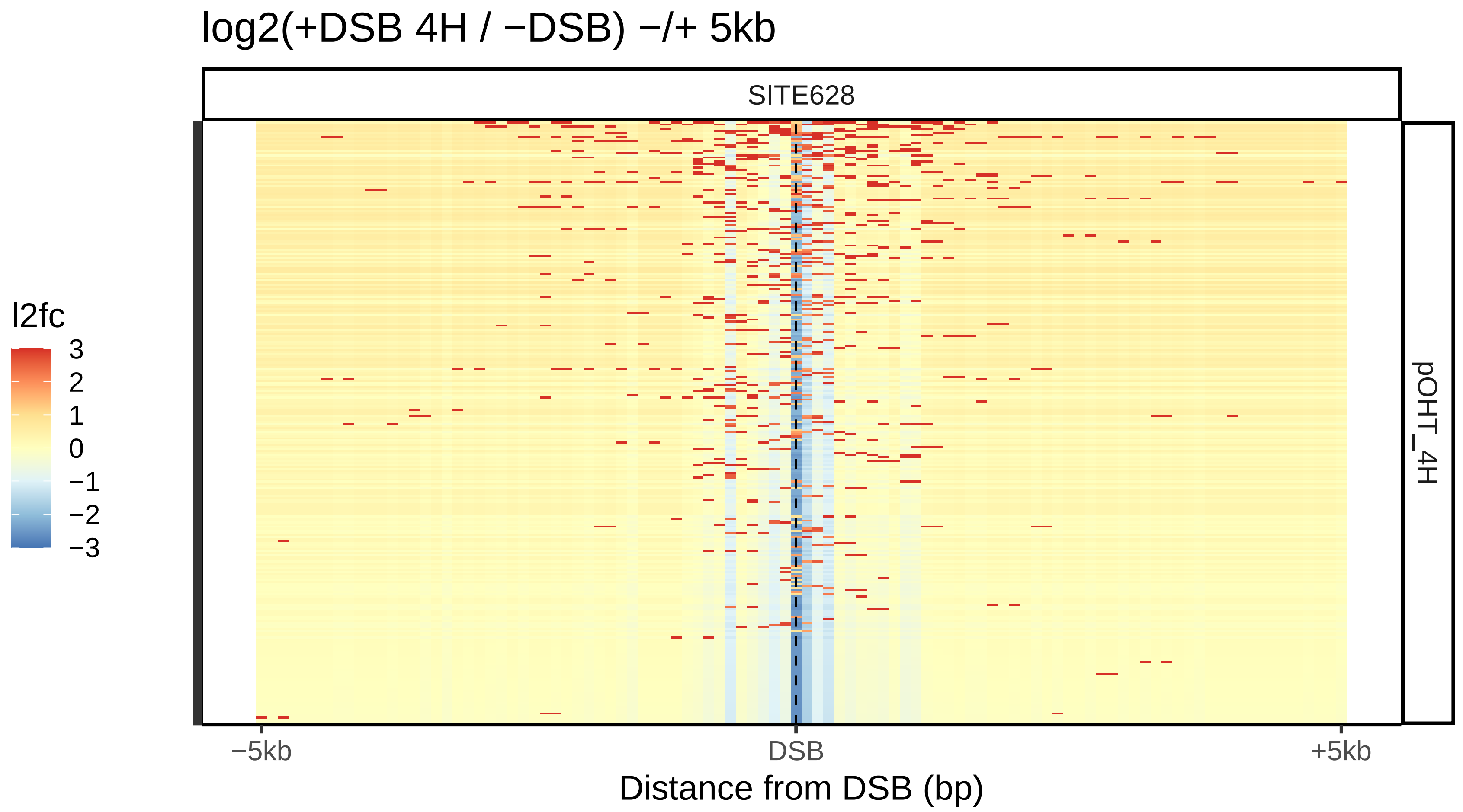
log2(+DSB 4H / −DSB) −/+ 5kb
SITE628
l2fc
3
2
1
0
−1
−2
−3
pOHT_4H
−5kb
DSB
+5kb
Distance from DSB (bp)

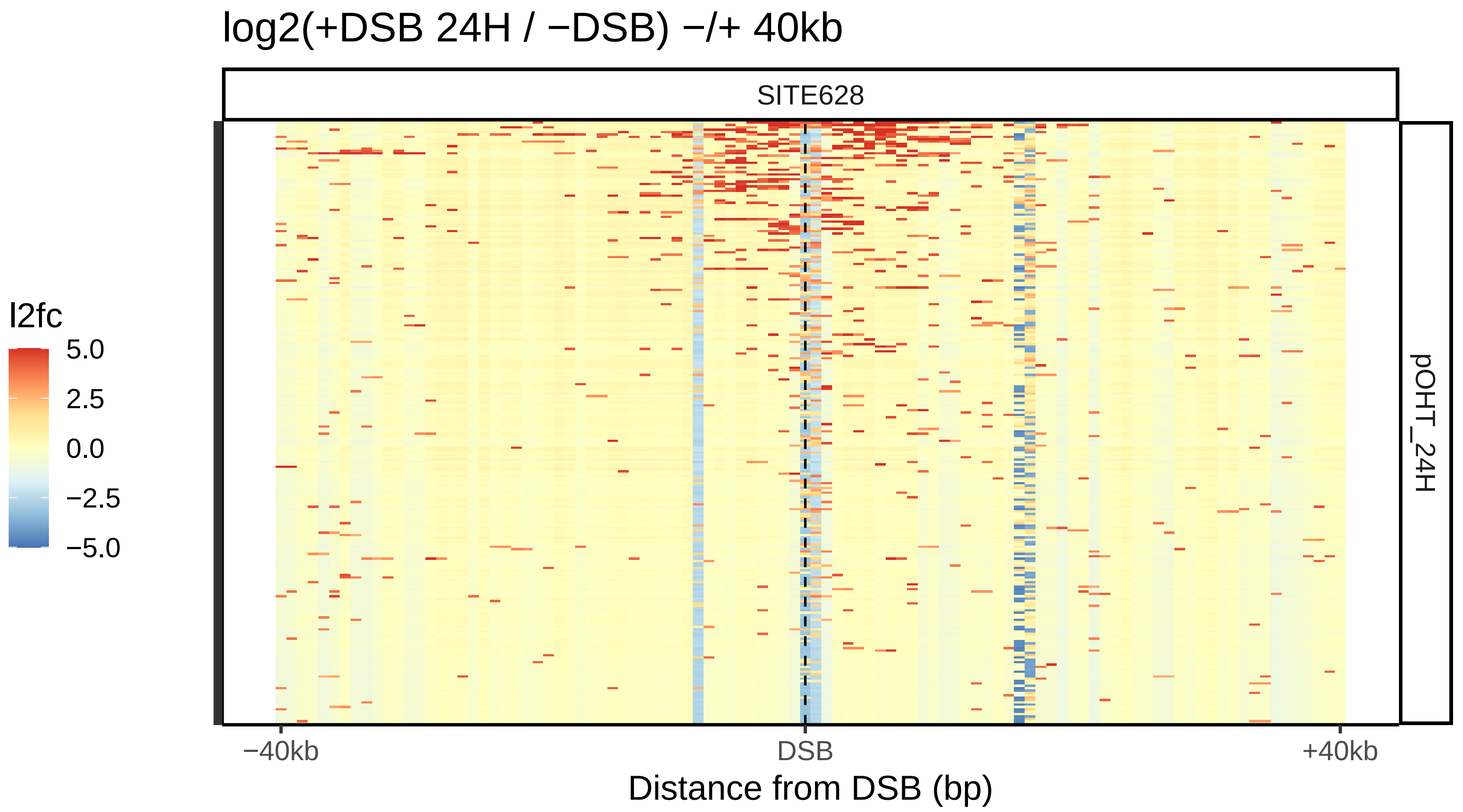
log2(+DSB 24H / −DSB) −/+ 40kb
SITE628
l2fc
5.0
2.5
0.0
−2.5
−5.0
pOHT_24H
−40kb
DSB
+40kb
Distance from DSB (bp)

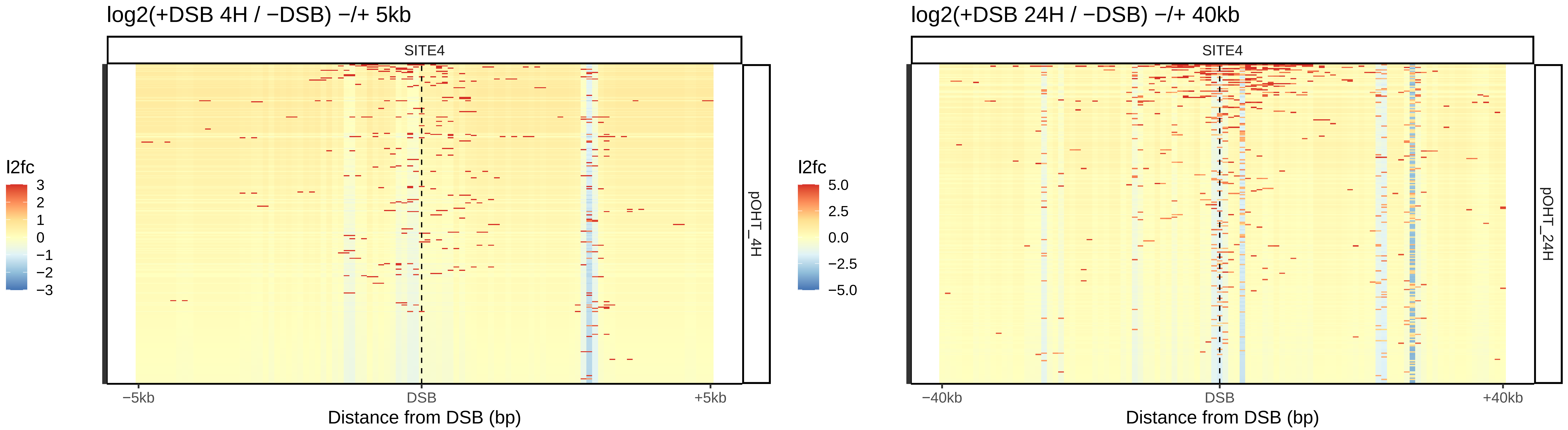
log2(+DSB 4H / −DSB) −/+ 5kb
SITE4
l2fc
3
2
1
0
−1
−2
−3
pOHT_4H
−5kb
DSB
+5kb
Distance from DSB (bp)
log2(+DSB 24H / −DSB) −/+ 40kb
SITE4
l2fc
5.0
2.5
0.0
−2.5
−5.0
pOHT_24H
−40kb
DSB
+40kb
Distance from DSB (bp)

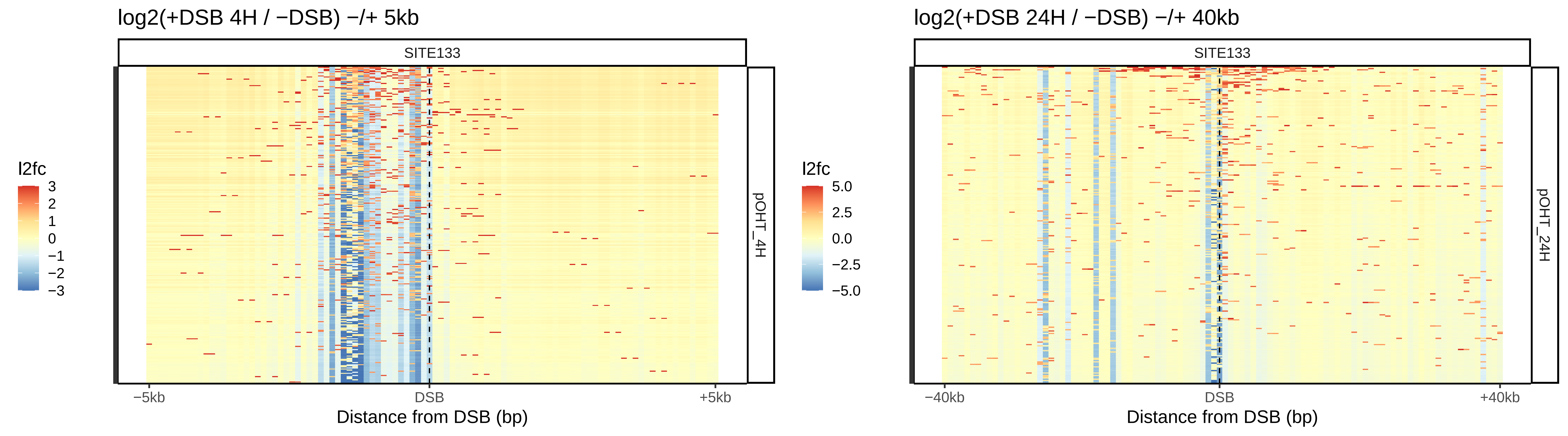
log2(+DSB 4H / −DSB) −/+ 5kb
SITE133
pOHT_4H
l2fc
3
2
1
0
−1
−2
−3
−5kb
DSB
+5kb
Distance from DSB (bp)
log2(+DSB 24H / −DSB) −/+ 40kb
SITE133
pOHT_24H
l2fc
5.0
2.5
0.0
−2.5
−5.0
−40kb
DSB
+40kb
Distance from DSB (bp)

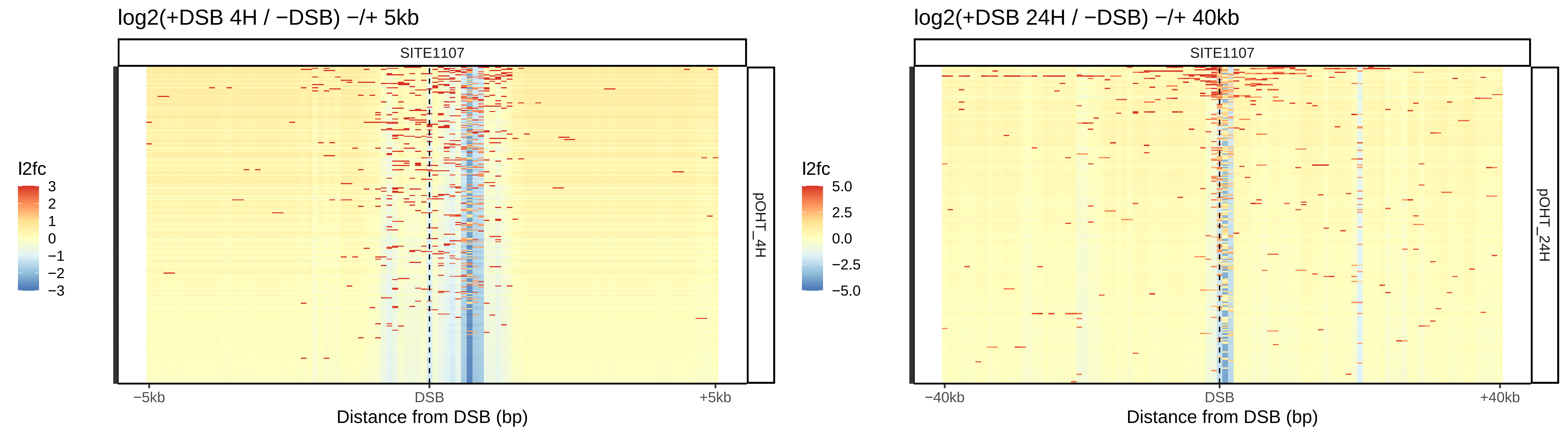

log2(+DSB 4H / −DSB) −/+ 5kb
SITE1107
l2fc
3
2
1
0
−1
−2
−3
pOHT_4H
−5kb
DSB
+5kb
Distance from DSB (bp)
log2(+DSB 24H / −DSB) −/+ 40kb
SITE1107
l2fc
5.0
2.5
0.0
−2.5
−5.0
pOHT_24H
−40kb
DSB
+40kb
Distance from DSB (bp)

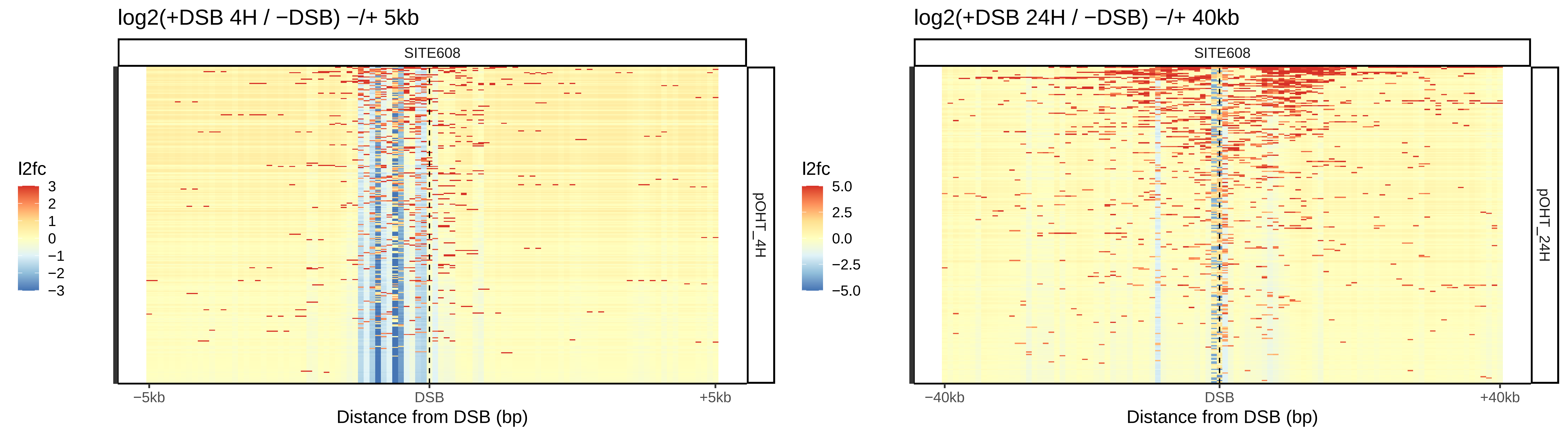
log2(+DSB 4H / −DSB) −/+ 5kb
SITE608
l2fc
3
2
1
0
−1
−2
−3
pOHT_4H
−5kb
DSB
+5kb
Distance from DSB (bp)
log2(+DSB 24H / −DSB) −/+ 40kb
SITE608
l2fc
5.0
2.5
0.0
−2.5
−5.0
pOHT_24H
−40kb
DSB
+40kb
Distance from DSB (bp)
